# Identifiable inverse optimal transport for single-cell state transitions

**DOI:** 10.64898/2026.10.01.756056

**Authors:** Han Dong, Yongqiang Gong, Yin Liu

## Abstract

Signed effects of target-state programmes are unidentifiable in balanced optimal transport because fixed endpoint distributions make their contributions constant across feasible plans. We introduce UOT-IOT, a semi-relaxed inverse transport framework that restores measurable target-mass responses and provides a rank condition for local identifiability. It recovers coefficients in ground-truth benchmarks (correlation 0.997 versus 0.081 for balanced transport), transfers frozen directions to six unseen samples with specificity controls, and achieves the lowest transition error against seven methods under matched evaluation conditions. GH-IOT separates shared directions from context and lineage-history offsets, revealing opposing history-associated effects. IFN/HLA is associated with patient treatment response, whereas EMT relates to drug sensitivity and perturbation-induced state changes. PERSIST-IOT improves future population-composition prediction in an independent experiment. Together, these modules distinguish current state, transition propensity and future population outcomes while supporting testable biological interpretations.

---

Single-cell sequencing measures molecular states at sampled times, whereas many biological questions concern subsequent changes in resistance, invasion or immune visibility. Pseudotime, RNA velocity and lineage tracing provide ordering, local direction and ancestry information, respectively.^1–12^ Similar transcriptional states can nevertheless arise from different histories and show different transition propensities across tissues, patients or treatments. Interpreting these changes requires determining whether the signed effects of biological programmes can be uniquely and stably recovered from observations.

Optimal transport (OT) models population-level state transitions. Waddington-OT links temporal snapshots with proliferation information, neural optimal transport learns perturbation responses, and moscot maps cells across time, space and modalities.^13–15^ Continuous-flow and gene-dynamics models extend prediction to unobserved times and perturbations.^16–19^ To interpret programmes as promoting or suppressing transitions, their signed coefficients must be identifiable from the observations and testable in independent data.

When both source and target marginals are fixed, a feature depending only on the target state contributes equally to every feasible transport plan. Its coefficient is a gauge degree of freedom, along which the plan-based loss is flat.^20,21^ Inverse OT theory characterises fixed-marginal identifiability, while unbalanced and entropy-regularised formulations provide mechanisms for relaxing marginal constraints.^22–26^ This ambiguity affects target-state programmes commonly interpreted biologically, including proliferation, epithelial–mesenchymal transition (EMT), stemness and immune visibility.^27–29^ Stable reconstruction of a transport plan therefore does not establish the signed effects of these programmes.

We address this ambiguity with three related modules (Fig. 1; Table 1). UOT-IOT fixes the source marginal and uses a finite Kullback–Leibler penalty on the target marginal to restore measurable responses and diagnose local identifiability. GH-IOT separates shared transition structure from context and lineage-history offsets. PERSIST-IOT propagates learned dynamics to future population composition and persistence detection. Using prespecified frozen protocols, we test coefficient recovery, external transfer with specificity controls, comparisons with seven methods, history- and treatment-associated programmes, and prospective prediction. These analyses evaluate numerical identifiability, biological interpretation and prediction separately.

**Fig. 1.**
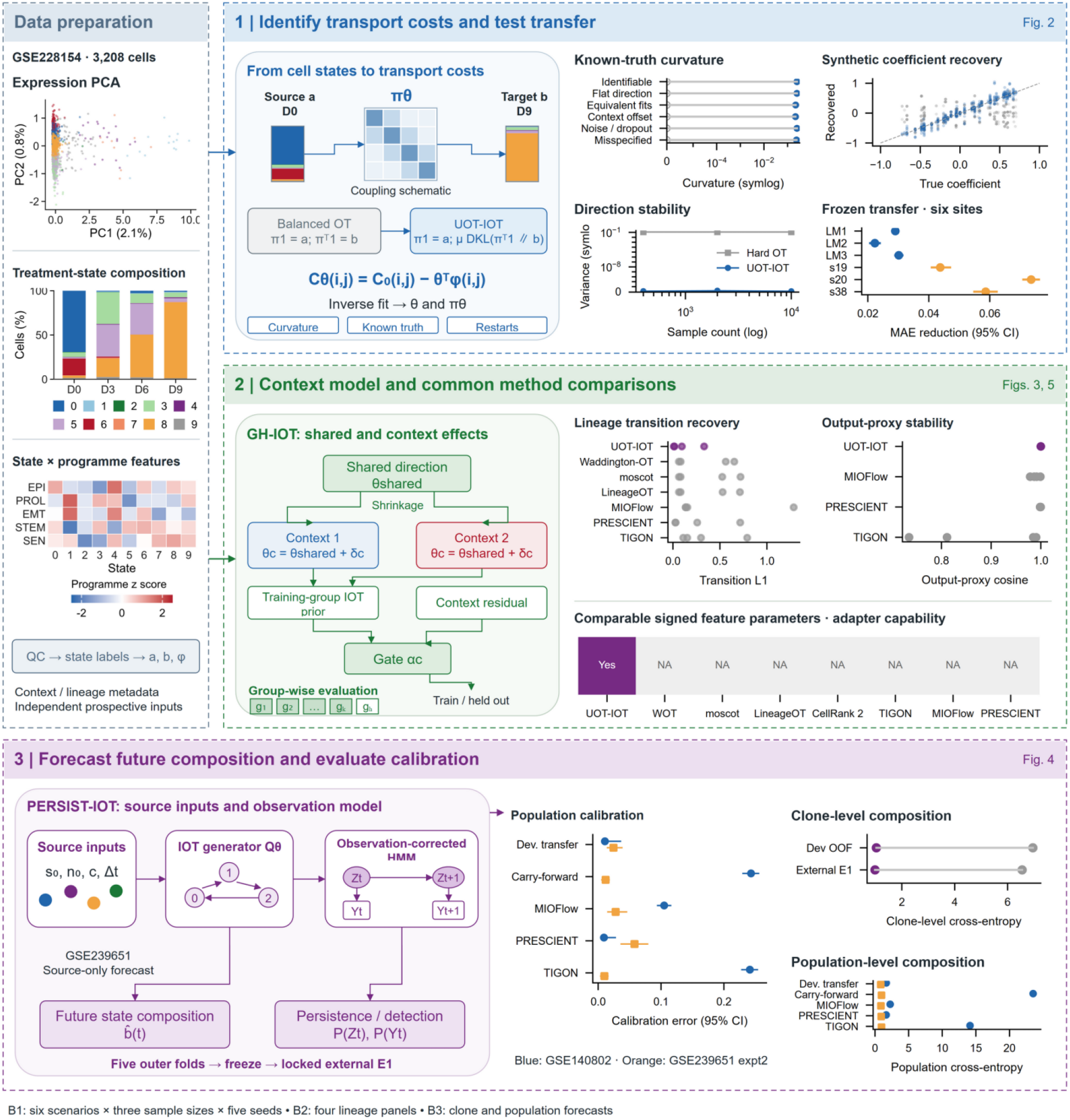
An identifiable inverse optimal transport framework for cell-state transitions. Figure 1 summarises the three modules: UOT-IOT learns and diagnoses identifiable directions; GH-IOT separates shared transition structure from lineage-history and context offsets; and PERSIST-IOT uses the learned operator to predict future population composition and persistence.

**Table 1.** Public datasets and validation design.

| Analysis | Data source | Included observations | Validation design |
| --- | --- | --- | --- |
| Temporal states and identifiability | GSE228154<sup>30</sup> | 3,208 cells; 4 time points; 10 states | D0, D3, D6 and D9; empirical curvature and state-programme diagnostics |
| Frozen external direction transfer | Training: GSE163558<sup>31</sup>; external: GSE246662<sup>32</sup> and GSE183904<sup>33</sup> | Training: 40,672 cells; 6 transition samples | Frozen costs and standardisation; LM1–LM3, s19, s20 and s38 |
| Lineage-transition benchmark | GSE140802<sup>9</sup>, GSE239651<sup>34</sup>, macsGESTALT<sup>35</sup> | 4 panels; 4–7 states | Comparison with 7 external method families under a common information boundary; 5 frozen seeds |
| Lineage history | GSE171940<sup>36</sup> | TENA: 5 mice; Cdh: 3 mice | Ever-EMT and epithelial lineages paired within each mouse |
| Patient response | ARTEMIS<sup>37</sup> | 79 patients (45 pCR, 34 RD); 39,426 tumour cells | Patient-aggregated scores; 10,000 patient bootstrap resamples; prespecified testing family |
| Drug sensitivity | GDSC<sup>38,39</sup> | 5 drugs; 513–520 cell lines per drug | Per-drug Spearman correlation between EMT score and $\ln(\text{IC}_{50})$ |
| PRRX1 perturbation | GSE164488<sup>40</sup> | 3 biological replicate pairs | Day 4 under TGFβ; siPRRX1 versus siCTR paired by replicate |
| Future composition and persistence | GSE239651<sup>34</sup> (CellTag<sup>41</sup>) | 2 independent experiments; 4,867/7,267 composition-prediction cases | Fivefold development grouped by source clone family; experiment 2 as locked external E1 |

Sizes refer to the analysis subsets used here. Statistical units vary by task, and a dataset may contribute to multiple tasks. pCR, pathological complete response; RD, residual disease.

## Results

### A semi-relaxed operator makes transition directions identifiable

Fixed-marginal transport generally cannot identify target-state programmes: such a programme contributes a constant to every feasible plan, so its coefficient does not affect the fit. UOT-IOT resolves this by fixing the source marginal while anchoring the target marginal with a finite Kullback–Leibler penalty. Target mass then has a closed-form sensitivity to each coefficient, the associated linear operator is invertible at finite penalty strength, and—after removing constant-feature and fixed-source gauge freedoms—the rank of the effective feature Jacobian establishes local identifiability (Methods). Directional identification thus becomes a computable property of the model rather than an assumption.

We tested this mechanism in a ground-truth benchmark spanning six scenario classes, three sample sizes (400, 2,000 and 10,000) and five seeds, for 90 paired runs with run-level diagnostics (Fig. 2a–c). Under balanced transport, curvature along pure-column directions is numerically zero (scenario means from ™2.7 × 10^−10^ to +3.0 × 10^−11^), whereas semi-relaxed fitting restores curvature to 0.217–0.250. Under the equivalent-fit setting, the correlation between true and recovered coefficients is 0.997 for the semi-relaxed fit versus 0.081 for balanced transport. Mean paired improvements across scenarios are +0.67 in coefficient cosine (worst paired lower bound +0.31), +0.24 in pure-column curvature (worst +0.18) and +0.12 in restart-direction variance (worst +0.07), based on 10,000 bootstrap resamples (Fig. 2b,c). Restart-direction variance falls from 0.12 under balanced transport to 1.5 × 10^−13^–2.5 × 10^−13^. The maximum discrepancy between the closed-form sensitivity and finite differences is 1.4 × 10^−10^. Across eight penalty values, the sensitivity norm decreases monotonically over the converged range (0.695 to 0.200), while the working point μ = 0.5 retains a sensitivity of 0.622 (Extended Data Fig. 1).

**Fig. 2.**
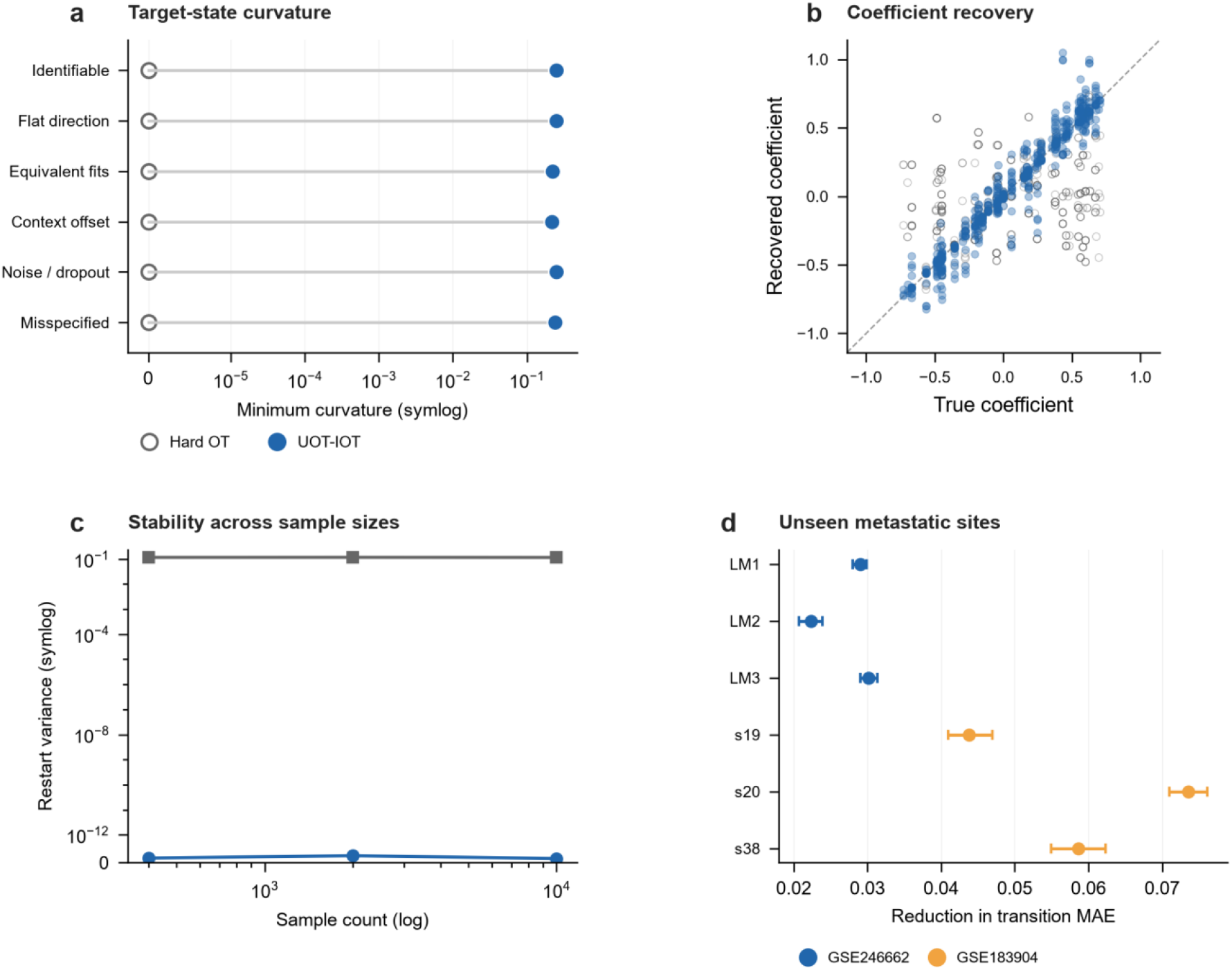
Identifiability and frozen-direction transfer. a, Empirical curvature for five pure-column target-state directions across six ground-truth scenarios; the horizontal axis uses a symlog scale with a linear threshold of 10^−5^. Open grey circles indicate balanced transport, filled blue circles indicate semi-relaxed inverse transport, and grey lines connect the two fits within each scenario. b, True versus recovered coefficients in an eight-state synthetic system, pooling all benchmark runs; open grey circles indicate balanced transport, filled blue circles indicate semi-relaxed fitting, and the dashed line indicates equality. c, Restart-direction variance across sample sizes (400, 2,000 and 10,000), with a symlog linear threshold of 10^−12^; grey squares indicate balanced transport and blue circles indicate semi-relaxed fitting. d, Improvement in conditional-transition MAE relative to the independent-marginal baseline in six unseen transition samples after training on GSE163558 and freezing model parameters and standardisation; points show improvement and horizontal bars show 95% intervals from 200 bootstrap resamples per site. Blue denotes GSE246662 and orange denotes GSE183904.

### Identifiable directions transfer with validated specificity

The frozen-model analysis asks whether a learned cost is a property of the transition itself or of a single sample. We trained once on 40,672 cells from GSE163558, fixed the cost parameters and feature standardisation, and evaluated conditional-transition mean absolute error in six transition samples that were never used for training: LM1–LM3 from GSE246662 and s19, s20 and s38 from GSE183904 (Fig. 2d). Every sample outperformed the independent-marginal baseline, with MAE improvements of 0.0223–0.0736 (relative reductions of 16.2–46.1%); all six site-level 95% bootstrap intervals excluded zero.

Specificity was tested with paired controls that reused exactly the same bootstrap resamples for the observed direction and every null direction (Extended Data Fig. 5). Matched-norm Gaussian directions yielded gains close to zero (site means −0.001 to 0.008), and none of 100 random directions produced positive gains at all six sites simultaneously (joint P = 0.0099). Reversing the frozen direction produced negative gains (−0.005 to −0.022). The transfer benefit is therefore specific to the learned direction: it disappears for random directions of the same magnitude and reverses when the direction is inverted.

### A unified benchmark tests transition recovery, stability and interpretability

Cross-method comparison is meaningful only when data, information boundaries and metrics are aligned. Our benchmark gives every method the same panel inputs, restricts each method to information allowed by its contract and scores all methods on common metrics across five frozen seeds (Methods). Version pins are reported in Extended Data Table 2 and complete run-level results in Extended Data Table 3. Seven externally developed method families passed the eligibility criteria—Waddington-OT, moscot, LineageOT, CellRank 2, TIGON, MIOFlow and PRESCIENT—alongside UOT-IOT and two simple baselines.

Across four lineage panels, UOT-IOT yields weighted transition-matrix L1 errors of 0.011– 0.332, the lowest in every panel, compared with 0.064–1.285 for comparator methods (Fig. 3a). Coupling error is competitive (UOT-IOT 0.80–1.00; comparators 0.93–1.01). Direction estimates are exceptionally stable: pairwise cosines are 1.000 in all four panels and the minimum cosine for real-data parameter directions is 0.996, compared with 0.735–0.999 for comparators (Fig. 3b). UOT-IOT is also the fastest method, with a median runtime of 1.1 s per run versus 3.4–33.0 s for the others (Fig. 3f). It is the only method that directly returns gauge-normalised signed feature coefficients; methods without such an output are recorded as structurally unavailable rather than unstable. Under the prespecified non-inferiority protocol (5% relative margins for coupling and transition metrics; a 3% absolute margin for top-1 accuracy; 10,000 paired bootstrap resamples), all three benchmark suites meet the primary criterion. Fate correlations and top-1 endpoints are reported in full as secondary outcomes (Extended Data Table 4), and directional availability is summarised in Extended Data Table 5.

**Fig. 3.**
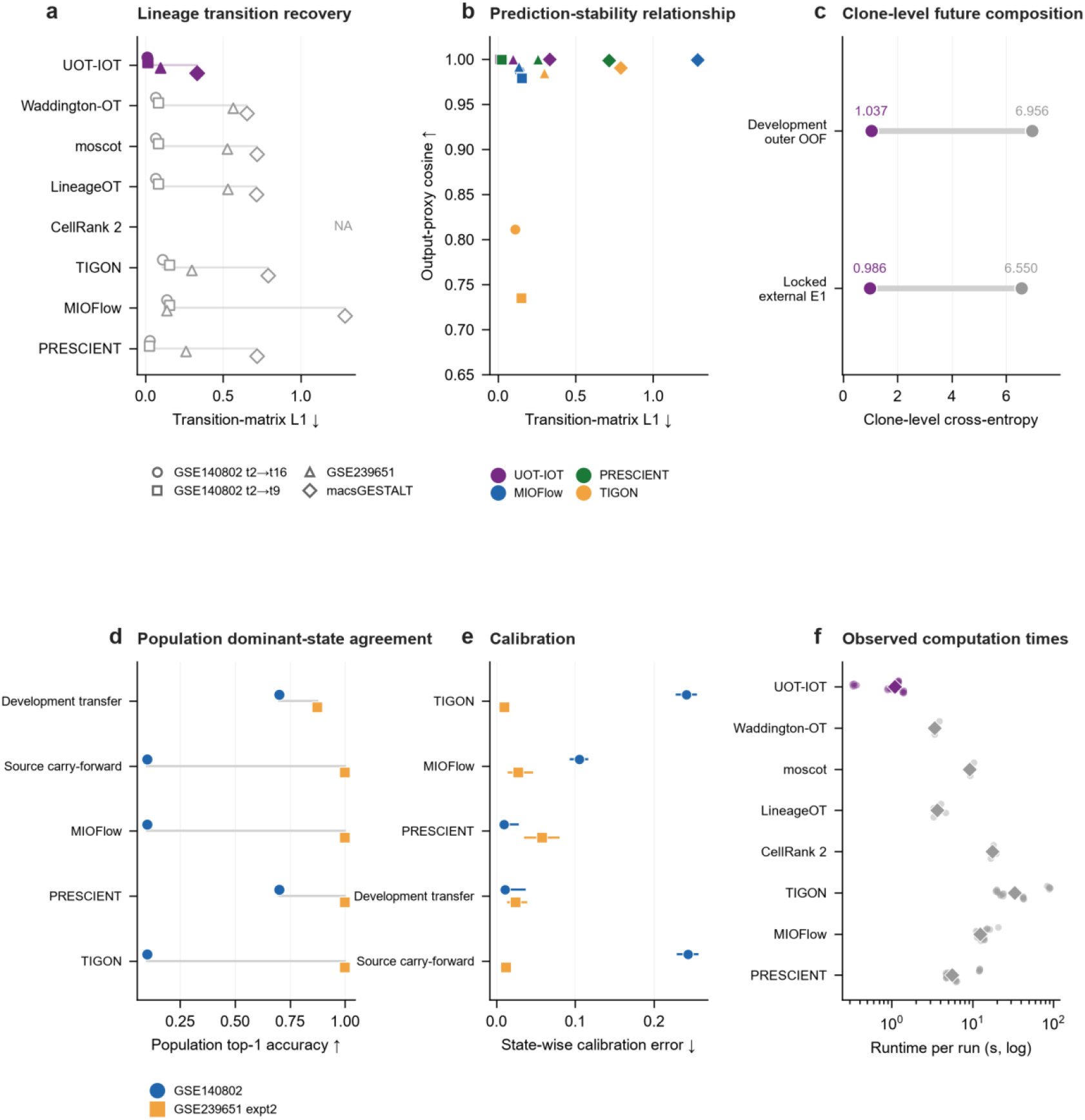
Benchmark comparison and prospective prediction. a, Weighted transition-matrix L1 error for each method across four lineage panels; symbols distinguish datasets, and purple lines and symbols denote UOT-IOT. b, Relationship between transition-matrix L1 error and proxy directional stability (pairwise cosine across seeds), shown only for the four methods with comparable outputs; colours indicate methods and shapes indicate datasets. c, Clone-level future-state cross-entropy for PERSIST-IOT and the no-transport ablation in development OOF prediction and the independent experiment; points show values and lines show paired comparisons. d, Population-level dominant-state agreement for five methods in the GSE140802 leave-one-animal panels and the independent GSE239651 experiment; symbols distinguish datasets. e, State-level expected calibration error with 95% intervals from 10,000 held-out-panel cluster bootstrap resamples. f, Runtime per run on a logarithmic axis; each point is one run and diamonds show method medians. NA indicates that a method does not provide the corresponding output.

### An identifiable operator predicts future population composition

PERSIST-IOT estimates time-varying state dynamics from training histories and combines them with source observations and treatment intervals to output future state composition and a separate persistence/detection probability (Methods). Trained on one CellTag experiment and evaluated in held-out folds and the independent GSE239651 experiment, it reduces clone-level future-state cross-entropy relative to the no-transport ablation of the same model from 6.956 to 1.037 in development (4,867 prediction cases) and from 6.550 to 0.986 in the independent experiment (7,267 cases)—reductions of 85.1% and 84.9%, respectively. Dominant-state agreement increases from 49.7% to 64.4% and from 47.5% to 68.2% (+14.7 and +20.7 percentage points; Fig. 3c; Supplementary Table 1). Cross-entropy uses a probability floor of 10^−12^, which strongly penalises the zero-probability predictions of the no-transport ablation (Methods).

Population-level composition—the endpoint applicable to external dynamics methods—is evaluated in the GSE140802 leave-one-animal panels and the independent GSE239651 experiment together with state-level calibration intervals (Fig. 3d,e; Extended Data Fig. 2). Across these panels, the strongest methods achieve composition errors of 0.86–1.68 and calibration errors of 0.009–0.024, providing an external reference for the clone-level results. Detection is reported as a secondary endpoint together with its uncertainty intervals and ranking metrics (Supplementary Table 1; Extended Data Table 6).

### The same operator provides an auditable derived time coordinate

The transition operator can also yield a one-dimensional relative time coordinate through expected random-walk hitting time to a specified root state. We treat this coordinate as a derived readout of the operator rather than independent evidence of direction. In a synthetic chain with known progression, it recovers the true ordering (mean Spearman ρ = 0.93) and is perfectly stable across restarts and sample sizes (pairwise ρ = 1.00). In real data, it remains stable across restarts (0.99–1.00) and leave-one-site resampling (0.97), and agrees with a diffusion-pseudotime baseline (ρ = 0.80). Its credibility therefore depends on the underlying transition operator. We report marginal-prevalence ordering, permuted ground truth, shuffled target labels, shuffled coupling and random-root controls in parallel (Extended Data Fig. 4); the same diagnostics that establish directional identifiability also determine whether this time coordinate can be attributed to a specific biological programme.

### Lineage history makes similar states point in different directions

Mathematical identifiability and external transfer show that a direction can be estimated, but not whether it carries biological information beyond the static expression state. GH-IOT therefore represents a transition relation as the sum of a direction shared across samples and a history/context offset, and uses permanent lineage labels to test whether history provides information beyond the current state. In GSE171940, the ever-EMT label retains whether cells have previously passed through partial or complete EMT; within each mouse, we compare ever-EMT and epithelial lineages using the same six-branch representation. The invasive-EMT branch points in opposite directions in the two labelling systems: −0.508 in TENA (95% CI −0.859 to −0.157, paired P = 0.0159, five mice) and +1.497 in Cdh (0.500 to 2.493, P = 0.0231, three mice; Fig. 4; Supplementary Table 2). The six-branch history-effect vectors have a cosine similarity of −0.796, and full-pipeline label permutations that preserve preprocessing, prior construction and held-group order give one-sided P = 0.0130 and P = 0.0300. These results show that observed expression primarily describes where a cell is now, whereas lineage history provides an additional condition on where it tends to go next: similar measured states do not imply a unique transition direction.

**Fig. 4.**
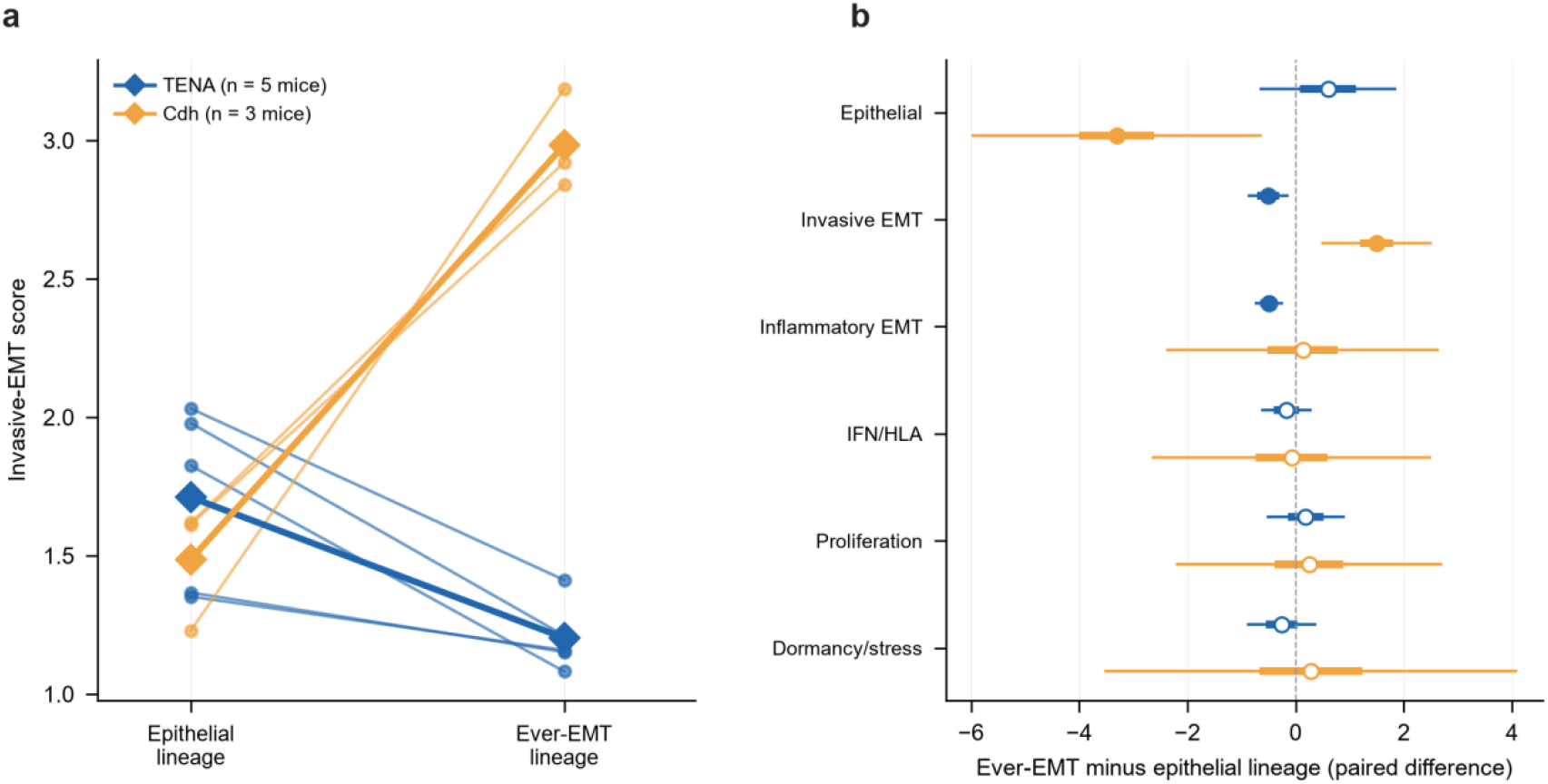
Lineage history makes similar states point in different directions. a, Paired invasive-EMT scores for ever-EMT and epithelial lineages within each mouse in GSE171940; small symbols and connecting lines denote individual mice (five TENA and three Cdh), and large diamonds denote group means. b, Mean differences across six branches (ever-EMT minus epithelial lineage) with paired 95% confidence intervals; thin bars show paired-t intervals and thick bars show mean ±1 s.e.; circles denote TENA, squares denote Cdh and filled symbols indicate raw paired P < 0.05. The statistical unit is the mouse. The cosine similarity between the six-branch direction vectors is −0.796; one-sided full-pipeline permutation tests (1,000 permutations) give P = 0.0130 for TENA and P = 0.0300 for Cdh.

### Different programmes link transition directions to distinct levels of treatment phenotype

We next asked whether biological programmes along identifiable directions have reproducible phenotypic meanings with clearly defined boundaries. In the ARTEMIS pretreatment cohort (79 patients; 45 pCR and 34 RD), IFN/HLA activity is higher in patients who achieve pCR (mean difference +0.260, patient-bootstrap 95% interval 0.104–0.404, Benjamini– Hochberg q = 0.0122, single-branch AUROC 0.697), whereas invasive EMT has little patient-level discriminatory value (+0.003, q = 0.821, AUROC 0.534; Fig. 5; Supplementary Table 2). This latter result does not imply that the EMT programme lacks biological meaning. Two orthogonal datasets test it at different levels: across five EGFR tyrosine kinase inhibitors in GDSC, EMT scores are positively correlated with ln(IC_50_) (median Spearman ρ = 0.2535; 513– 520 cell lines per drug; all P < 10^−5^), and in MDCK cells, PRRX1 knockdown shifts epithelial and invasive-EMT scores in opposite directions (paired P = 0.0206 and P = 0.0160, three biological replicate pairs; Extended Data Fig. 3). Different programmes therefore play different interpretive roles: IFN/HLA corresponds to an immune-visibility programme associated with patient treatment response, whereas EMT is better viewed as an axis of drug sensitivity and state change than as a universal patient-level classifier.

**Fig. 5.**
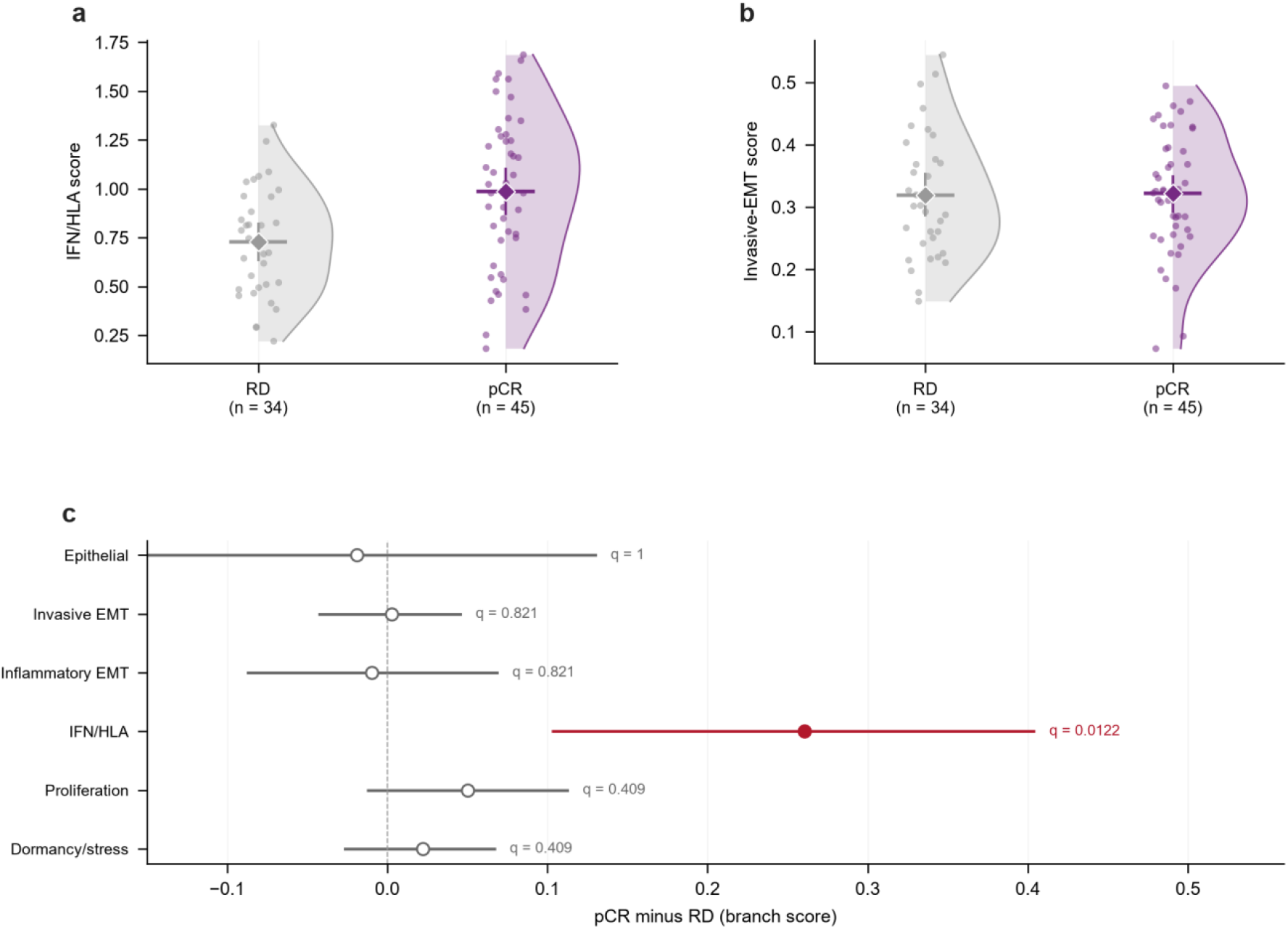
Biological programmes link transition directions to treatment response. a, Patient-level IFN/HLA scores for pCR and RD patients in ARTEMIS; points denote patients, half-violins show within-group distributions, and diamonds with error bars show means and 95% intervals from 10,000 patient bootstrap resamples (pCR − RD = +0.260; q = 0.0122; AUROC = 0.697). b, Corresponding invasive-EMT scores (pCR − RD = +0.003; q = 0.821; AUROC = 0.534). c, Patient-level mean differences (pCR − RD) and bootstrap intervals for six predefined branches; Benjamini–Hochberg adjustment retains the prespecified testing family. IFN/HLA is the only branch that separates the two outcome groups.

## Discussion

UOT-IOT makes the identifiability of signed programme coefficients testable. Relaxing the target marginal converts a fixed-marginal gauge direction into a measurable target-mass response. Closed-form sensitivity, feature-rank conditions, empirical curvature and restart stability provide complementary diagnostics, while ground-truth recovery and controlled frozen transfer test whether estimated directions are recoverable and transferable. Biological interpretation remains conditional on these checks and the chosen feature representation.

Under matched information boundaries, the framework achieves strong transition recovery, directional stability and runtime while meeting prespecified predictive non-inferiority criteria. Its gauge-normalised signed coefficients provide an output distinct from transport plans and prospective predictions; each requires separate validation.

The biological analyses associate lineage history with different, sometimes opposing, programme directions and link different programmes to distinct endpoints. In ARTEMIS, IFN/HLA is associated with pathological complete response. EMT shows little patient-level discrimination in that cohort but remains associated with drug sensitivity and PRRX1 perturbation. These findings support context-dependent programme interpretation and do not establish causal effects of history or programme activity.

PERSIST-IOT improves future composition prediction in held-out and independent experiments, whereas persistence-detection gains vary by metric and partition. Cross-entropy improvements must also be interpreted in light of the probability floor and the no-transport ablation’s zero-probability predictions. The outputs describe population composition and probabilistic persistence or detection, rather than deterministic individual-cell fate. Together, the framework and its diagnostics distinguish current state, transition propensity and future population outcome, providing parameters for population comparisons and experimentally testable hypotheses.

## Online Methods

### Study design and data units

We computationally reanalyse public data, using synthetic systems for coefficient and coupling recovery and public single-cell, lineage and patient data for model evaluation; no new biological samples were generated. All analyses use explicitly defined statistical units (cells, mice, patients, cell lines or prediction cases), and the same dataset may contribute to multiple tasks; complete definitions of tasks, units and information boundaries are provided in Extended Data Table 1. GSE228154 contains 3,208 cells across four afatinib time points (D0, D3, D6 and D9) and ten states. GSE163558 provides 40,672 training cells; frozen-direction transfer is evaluated on LM1–LM3 from GSE246662 and s19, s20 and s38 from GSE183904. The lineage-transition benchmark uses four panels: two time spans from GSE140802, the GSE239651 transition panel and the macsGESTALT panel. Prospective prediction uses two independent CellTag experiments from GSE239651. Lineage-history analysis uses GSE171940 (five TENA and three Cdh mice). After filtering for pCR/RD outcomes and tumour annotation, the ARTEMIS pretreatment cohort contributes 79 patients (45 pCR and 34 RD; 39,426 tumour cells). GDSC provides matched expression and drug-sensitivity records for five EGFR tyrosine kinase inhibitors (513–520 cell lines per drug), and GSE164488 provides three paired MDCK replicates at day 4 under TGFβ.

### State representations and features

States follow the existing cell groupings, state centroids and metadata in each task. Features combine expression similarity, source and target programme scores and specified interaction terms; standardisation statistics are fixed during training and reused for external samples. Programme interpretation uses frozen gene sets for six branches: epithelial, invasive EMT, inflammatory EMT, IFN/HLA, proliferation and dormancy/stress. Gene coverage in ARTEMIS is 8/8, 17/17, 15/15, 17/17, 12/12 and 12/12, respectively. Programme definitions are fixed before outcome comparisons, and patient scores aggregate tumour-cell scores. Quantitative conclusions come from the cost and prediction tasks; the expression principal components and state heat maps in Fig. 1 summarise saved inputs and features.

### Semi-relaxed transport and cost learning

Let *a* and *b* be the observed source and target distributions, with cost features *ϕ*_*ij*_ ∈ ℝ^*F*^ We parameterise the cost as

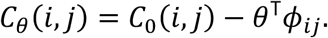

Thus, conditional on feature standardisation and the remaining coefficients, a positive coefficient lowers the transition cost as its feature increases. The semi-relaxed forward problem is

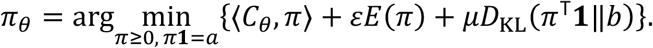

where *E*(*π*) = Σ_*ij*_ *π*_*ij*_ (log*π*_*ij*_ – 1),ε > 0, and 0 ≤ *µ* < ∞. The generalized Kullback–Leibler divergence follows its standard definition.^42^

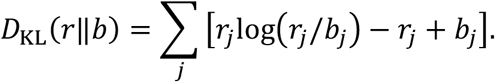

Fixing the source marginal through *a* fixes total mass, whereas target relaxation adjusts relative state composition; absolute proliferation, death and sampling intensity require additional count and observation models.

The transport plan defines row-conditional transition probabilities *T*_*θ*_ (*i,j*) = *π*_*θ*_(*i,j*)/*a*. Given observed transitions 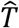, the outer fit minimises cross-entropy between them and*T*_*θ*_, with task-specific parameter regularisation. Multiple observations are fitted jointly according to the training design. Expression-derived observation plans and measured lineage labels retain separate records of their information sources. The forward solution uses accelerated fixed-point iteration, and parameter gradients follow implicit differentiation of the converged equations; numerical precision, initialisation and stopping criteria follow the corresponding analysis configurations, and closed-form checks use double precision.

### Pure-column equivalence and sensitivity

For feature *f* with *ϕ*_*ij,f*_ = *ν*_*j*_ balanced constraints *π***1** = *a* and *π*^⊤^ **1** = *b* imply

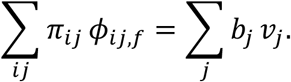

The right-hand side is the feature moment of the fixed target marginal. All values of *θ*_*f*_ therefore yield the same optimal plan; the forward optimal value changes linearly with that coefficient, whereas the plan-based outer loss remains constant. Pure-row additive costs have the same equivalence structure under a fixed source marginal.

At a semi-relaxed solution, let *r* = *π*^⊤^ **1**, *s*_*i,j*_ = *π*_*ij*_/*a*_*i*_, and define

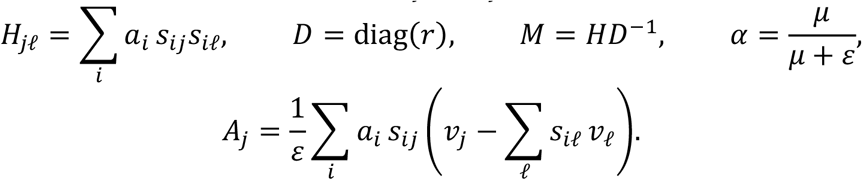

The target-mass response along a pure-column direction is

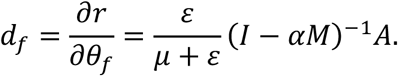

Under positive support and smoothness of the forward solution, *M* is similar to a row-stochastic matrix and has spectral radius 1. Finite *μ* gives *α* < *1*, so *I* − *αM* is invertible, and *A* ≠ *0* implies *d*_*f*_ ≠ 0; a constant direction gives *A* = *0* . For multiple coefficients, local identification additionally requires the observation Jacobian to have full column rank after gauge freedoms are removed. Figure 2 reports directional curvature, coefficient recovery and restart-stability diagnostics; Extended Data Fig. 1 reports the penalty scan.

### Ground-truth benchmark (B1)

B1 comprises six scenario classes: an identifiable soft-marginal construction; a hard-marginal pure-column construction with a provably flat direction; equivalent fits in which different parameter vectors produce similar target distributions; shared structure plus context offsets; observation noise and dropout; and misspecification with nonlinear or hidden confounding. Each scenario is evaluated at three sample sizes (400, 2,000 and 10,000) and five frozen seeds, giving 90 paired runs per method. Metrics are the minimum curvature along pure-column directions, correlation between true and recovered coefficients, held-out coupling error and restart-direction variance. Paired comparisons use 10,000 bootstrap resamples; the flat-direction and equivalent-fit settings form the basis of the parameter-recovery criteria. Results are reported as scenario means and paired intervals across matched runs.

### External-method benchmark (B2) and information boundaries

B2 evaluates all methods on the same lineage panels. Each method receives the panel, permitted time points and any information not prohibited by its contract; prospective tasks prohibit target observations, target marginals and future clone graphs. Adapters call fixed versions of Waddington-OT, moscot, LineageOT, CellRank 2, TIGON, MIOFlow and PRESCIENT, with versions and commits recorded. Metrics are weighted transition-matrix L1 error, relative Frobenius coupling error, fate Pearson/Spearman correlation, top-1 fate accuracy, runtime and peak memory. Metrics not produced by a method are recorded as NA and are not imputed. The non-inferiority protocol fixes a 5% relative margin for coupling and transition metrics and a 3% absolute margin for top-1 accuracy, evaluated against each external method using 10,000 paired bootstrap resamples. Primary B2 decisions use coupling and transition metrics; fate correlation and top-1 comparisons are secondary outcomes.

### Prospective prediction and observation model

PERSIST-IOT estimates a continuous-time generator *O* from training histories. Off-diagonal entries are parameterised with softplus, and each diagonal entry is the negative sum of the remaining entries in that row, ensuring zero row sums and non-negative off-diagonal entries. The transition reference over interval *Δt* = exp(*OΔt*).^43^ The generator is fitted using future-composition cross-entropy and parameter regularisation. For training source marginal *ā*, the prospective target prior is

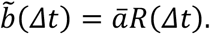

A balanced Sinkhorn projection onto a and b_Δt gives the conditional transition matrix used to propagate each clone’s source composition. This projection uses the generator-derived target prior, whereas the semi-relaxed snapshot problem uses the observed target distribution. Prediction inputs comprise source observations, treatment and the specified interval; future observations are used only for evaluation.

The persistence/detection observation model contains an active latent state and an absorbing inactive state. Source features determine the hazard of exiting the active state; count observations follow a zero-inflated negative binomial distribution, and the inactive-state background rate is fixed from training.^44^ A two-state forward recursion computes sequence likelihoods at time points with observations.^45^ The model reports latent persistence probability and observed detection probability separately. Detection metrics use observed detection labels, and calibration follows the locked run configuration.

### Context and composition evaluation

GH-IOT represents the cost in context *c* as *θ*_*c*_ = *θ*_shared_+ *δ*_*c*_. Context offsets have shrinkage regularisation selected by inner evaluation within training groups. Gate configurations combine IOT prior predictions with supervised residuals; the global gate uses a shared weight, and the Dirichlet–multinomial configuration uses a composition likelihood.^46^ The common metric is per-cell Dirichlet–multinomial negative log-likelihood; methods are ranked within each task and ranks are averaged across tasks. History-direction comparisons use frozen six-branch representations and within-mouse paired differences. The overall history test uses 1,000 label permutations that preserve preprocessing, prior construction and held-group order; empirical P values use the plus-one correction and a one-sided tail in the prespecified direction.^47^

### Pseudotime readout and evaluation

The row-normalised state-transition operator defines a random walk over states. For a specified root, let *o* denote the root state, *F* the remaining states *W* and the stochastic matrix used to compute hitting times; then

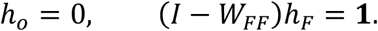

Solving this system yields a relative pseudotime coordinate. The root is chosen as the state with the largest source mass, and root sensitivity is reported as a control. Evaluation uses the correlation between state-level order and known time information, restart/seed/leave-one-unit-out stability, and a diffusion-pseudotime baseline (kNN graph, k = 15; lazy diffusion α = 0.5; hitting time to the root cell). Marginal-prevalence ordering, permuted ground truth, shuffled target labels, shuffled coupling and random-root controls are all reported. The synthetic chain uses latent progression as ground truth; GSE228154 uses frozen inverse-OT observations (source distributions, feature tensor and observed transition operator); lineage panels use the couplings from each benchmarked method. Acceptance criteria were frozen before analysis; complete coordinates, baselines and controls are reported in Extended Data Fig. 4 and the source data.

### Statistics and reproducibility

Patient-level mean differences and intervals use 10,000 patient bootstrap resamples, with Benjamini–Hochberg adjustment retaining the prespecified testing family.^48,49^ Mouse-lineage and MDCK comparisons use two-sided paired t tests with t-based 95% intervals; the statistical unit is the mouse or biological replicate pair. GDSC uses per-drug Spearman correlation, and cell-line sets may overlap across drugs. External transfer uses 200 bootstrap resamples per site; negative controls reuse exactly the same resamples for the frozen direction, 100 matched-norm Gaussian directions, 100 coefficient-permutation directions, the sign-reversed direction and the zero direction. Empirical two-sided P values are derived from the direction-sampling distribution. Because several cost features are correlated, permutation nulls can retain some information; matched-norm Gaussian directions are therefore the primary control. Future-state composition is evaluated by cross-entropy, dominant-state agreement and state-level calibration error, with a 10^−12^ probability floor for cross-entropy. Brier differences for detection use 10,000 paired clone bootstrap resamples, keeping related cases from the same clone within the same resampling unit; AUPRC is reported separately.^50,51^ Calibration intervals use 10,000 bootstrap resamples clustered by held-out panel with 10 bins. The best supervised comparator is selected separately by partition and metric.^52^ Benchmark protocols, seeds, method-version pins and gating definitions are frozen in hashed protocol files; every figure panel maps to a machine-readable source table with input hashes. Release gates G1–G9 cover method coverage, the three benchmarks, non-inferiority, parameter recovery, directional stability, external directional validity, figure reproduction, public release and the pseudotime readout.

## Data availability

GEO data are available under accession numbers GSE228154, GSE163558, GSE246662, GSE183904, GSE173958, GSE171940, GSE164488, GSE239651 and GSE140802. Public ARTEMIS data are available in the CZI CELLxGENE TNBC collection (SRA project PRJNA1041570). GDSC drug-sensitivity and cell-line data are available from Genomics of Drug Sensitivity in Cancer. Figure source data, frozen panels and derived statistical tables are provided with the reproducibility package (see Code availability), with statistical units matching the figure legends.

## Code availability

The benchmark package, including frozen panels, method adapters, review tables and figure-generation scripts, is released under the MIT licence at https://github.com/XTSgreen/IOT-single-cell-paper. The core implementation is distributed as bioIOT (version 0.2.1) on PyPI and in the bioIOT-py GitHub repository, providing semi-relaxed inverse optimal transport, cost fitting, implicit gradients, transition matrices, pseudotime and AnnData interfaces. A versioned archive is released with the paper.

## Notes

Funding: This work was supported by the Tianjin Undergraduate Teaching Quality & Reform Research Program for Higher Education (B251005526).

### Competing Interest Statement

The authors have declared no competing interest.

