## Supplemental materials for "Identifiable inverse optimal transport for single-cell state transitions"

Extended Data Figures 1–5, Supplementary Tables 1–2 and Extended Data Tables 1–6.

### Extended Data Figures

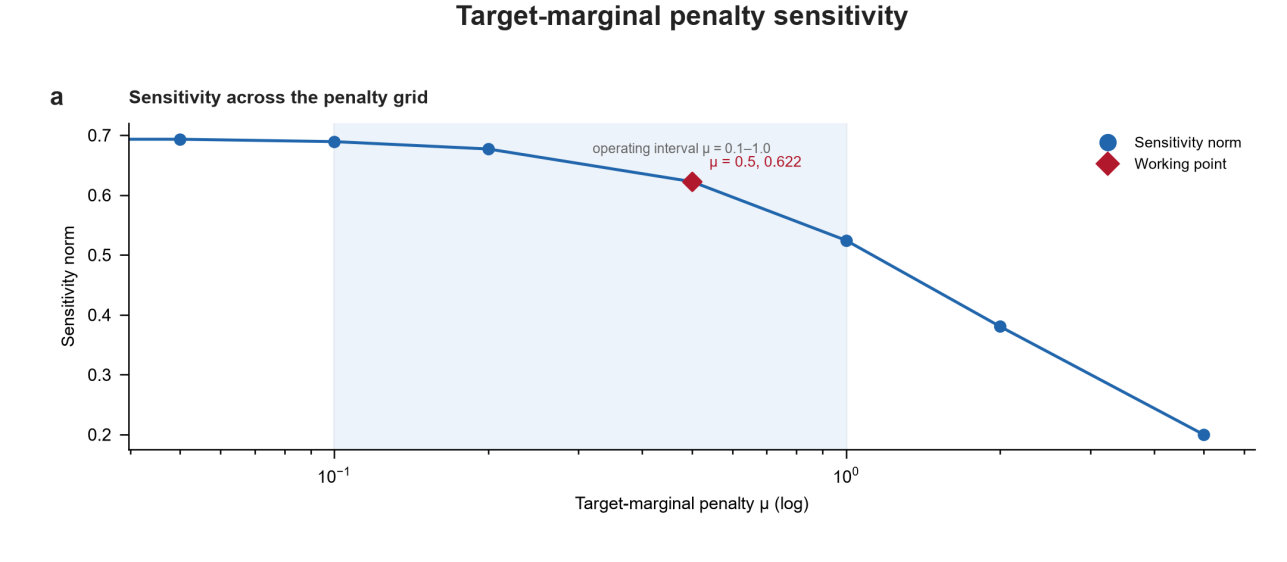

Extended Data Fig. 1 | Target-marginal penalty sensitivity scan.

Sensitivity norm along the pure-column direction in the synthetic system as a function of target-marginal penalty μ (eight penalty values). The red star marks the working point μ = 0.5 (sensitivity norm 0.622); the curve decreases monotonically over the converged range (0.695 to 0.200).

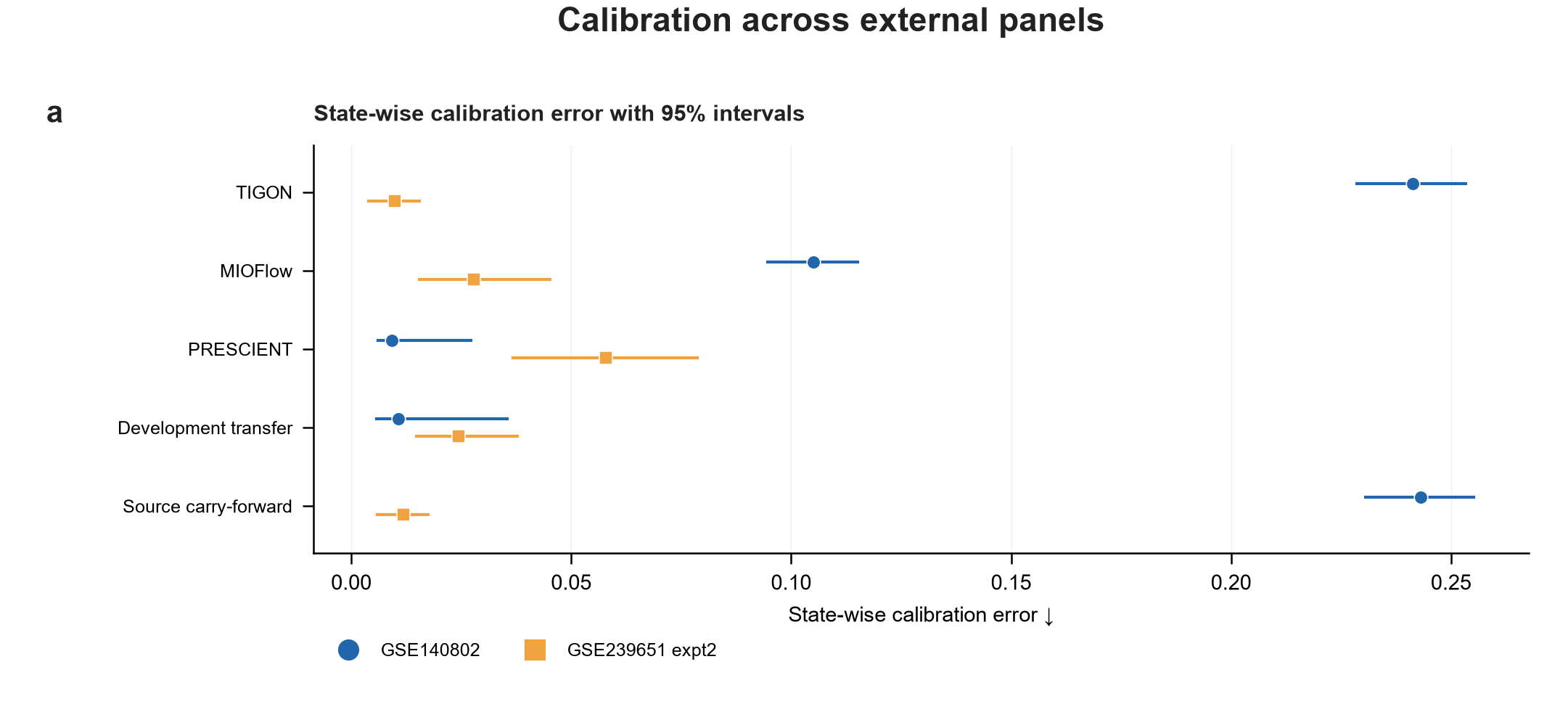

Extended Data Fig. 2 | External-panel calibration.

State-level expected calibration error for each method on GSE140802 and GSE239651 expt2, with 95% intervals from 10,000 held-out-panel cluster bootstrap resamples; blue denotes GSE140802 and orange denotes GSE239651 expt2.

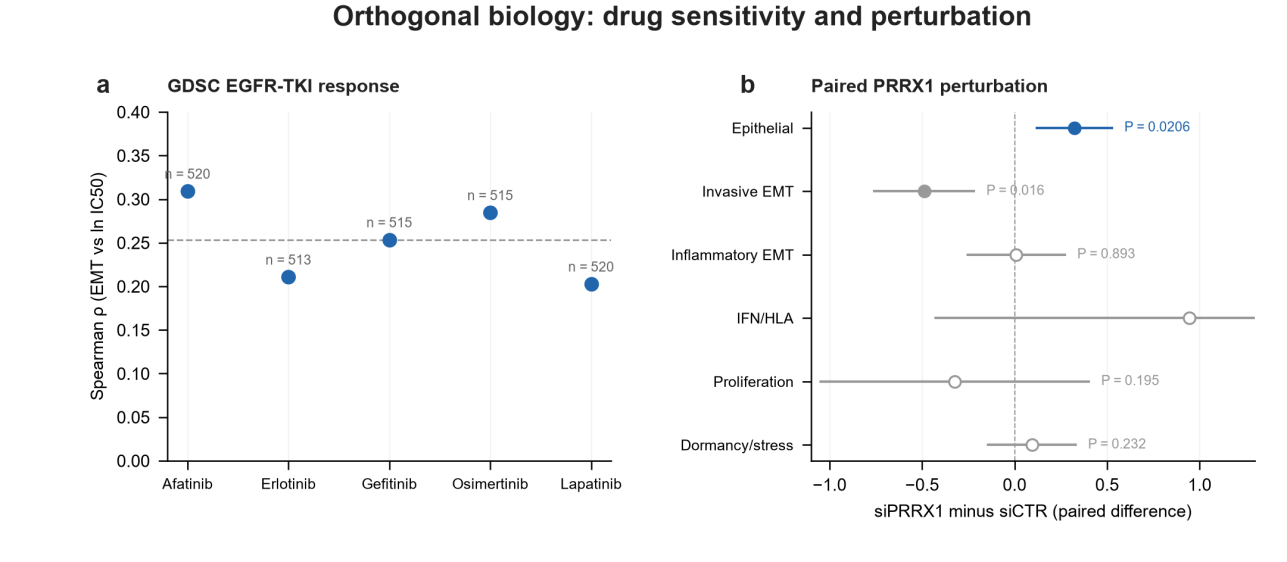

Extended Data Fig. 3 | Orthogonal biology: drug sensitivity and perturbation.

a, Spearman correlation between EMT score and ln(IC50) for five EGFR tyrosine kinase inhibitors in GDSC (points; dashed line shows the median, 0.2535; labels give n = 513–520 cell lines per drug). b, Paired siPRRX1 − siCTR differences across six branches in MDCK cells with t-based 95% intervals; filled symbols indicate paired P < 0.05 (epithelial P = 0.0206, invasive EMT P = 0.0160; n = 3 biological replicate pairs).

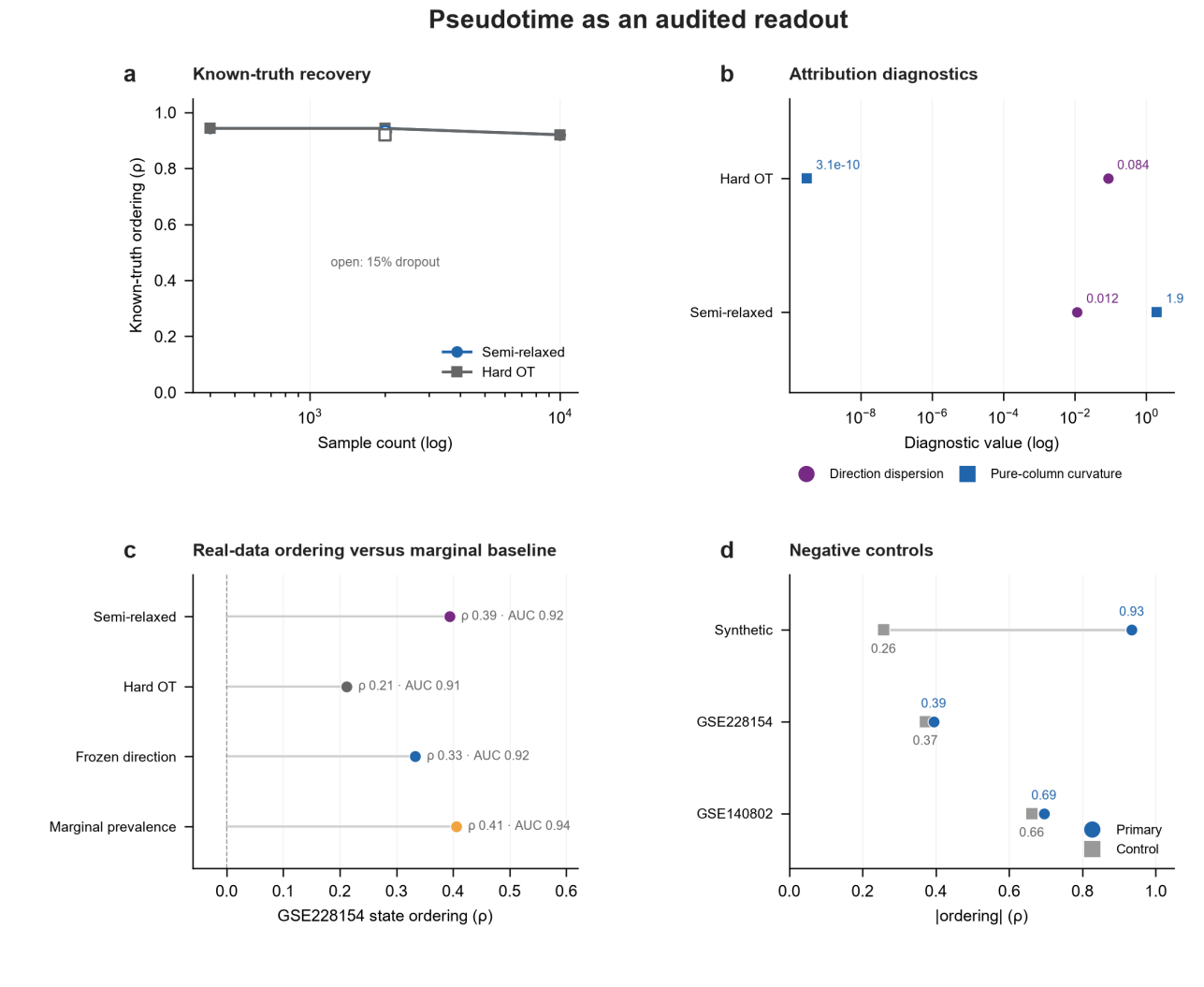

Extended Data Fig. 4 | Pseudotime as an auditable readout.

a, Ground-truth ordering of the hitting-time readout under semi-relaxed and hard-marginal fits in the synthetic chain across sample sizes; open symbols indicate 15% observation dropout. b, Directional dispersion and pure-column curvature for hard-marginal and semi-relaxed fits; ordering is stable under both, but programme attribution is identifiable only in the semi-relaxed fit. c, GSE228154 state ordering for semi-relaxed, hard-marginal, frozen-direction and marginal-prevalence pseudotime, with time-separation AUC annotated; the marginal-prevalence baseline is comparable to the fitted operator (ρ = 0.39 versus 0.41; AUC 0.92 versus 0.94). d, Negative controls (permuted ground truth, shuffled target labels and shuffled coupling) compared with the corresponding main results; the shuffled-coupling control remains correlated (|ρ| = 0.66). These controls constitute the auditing standard proposed in the main text.

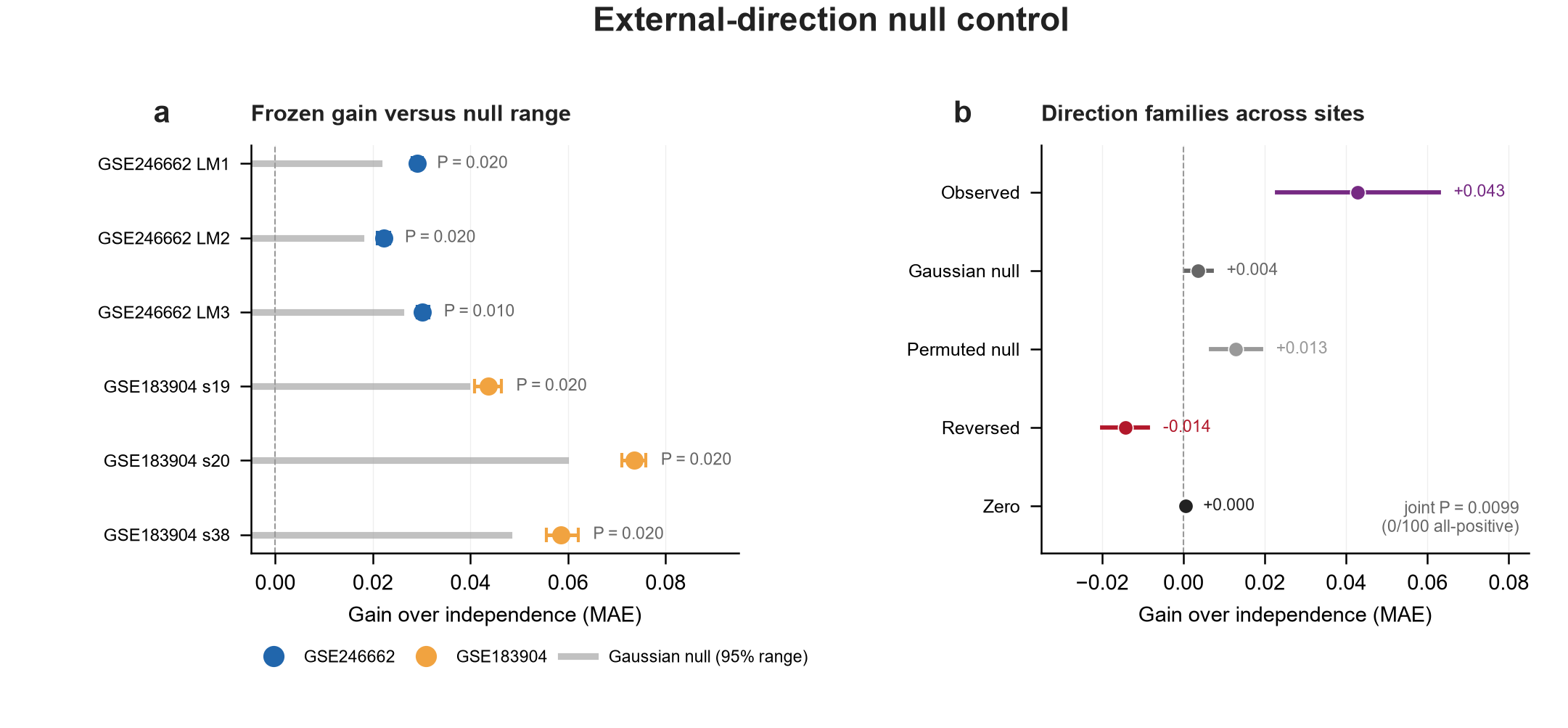

Extended Data Fig. 5 | Negative controls for external direction.

a, Gain of the frozen GSE163558 direction relative to the independent baseline (points with 95% intervals from 200 paired bootstrap resamples per site) together with the 95% range for matched-norm Gaussian directions (grey bars), with three sites in each validation cohort; annotated P values are empirical two-sided estimates over 100 Gaussian directions. b, Mean gain across six sites for the observed direction, Gaussian null directions, coefficient-permutation null directions, the sign-reversed direction and the zero direction (bars show mean ± s.d. across sites). Because of information redundancy among cost features, permutation directions retain a small positive gain and are reported as a weaker control.

Supplementary Table 1 | Future-state composition and persistence performance.

| Metric | Partition | Cases | PERSIST-IOT | Baseline | Difference |
| --- | --- | --- | --- | --- | --- |
| State cross-entropy ↓ | Dev. Outer OOF | 4,867 | 1.0370 | 6.9556ᵃ | −85.1% (relative) |
| State cross-entropy ↓ | External E1 | 7,267 | 0.9859 | 6.5497ᵃ | −84.9% (relative) |
| Dominant-state agreement ↑ | Dev. Outer OOF | 4,867 | 64.43% | 49.70%ᵃ | +14.73 percentage points |
| Dominant-state agreement ↑ | External E1 | 7,267 | 68.20% | 47.50%ᵃ | +20.70 percentage points |
| Sinkhorn divergence ↓ | Dev. Outer OOF | 4,867 | 0.1024 | 0.1784ᵃ | −42.6% (relative) |
| Sinkhorn divergence ↓ | External E1 | 7,267 | 0.1110 | 0.1965ᵃ | −43.5% (relative) |
| Detection Brier ↓ | Dev. Outer OOF | 38,010 | 0.050092 | 0.045277ᵇ | +0.004815 [0.002691, 0.007193] |
| Detection Brier ↓ | External E1 | 109,704 | 0.029756 | 0.031034ᵈ | −0.001279 [−0.002188, −0.000381] |
| Detection AUPRC ↑ | Dev. Outer OOF | 38,010 | 0.78065 | 0.80982ᶜ | −0.02917 |
| Detection AUPRC ↑ | External E1 | 109,704 | 0.71560 | 0.73275ᵈ | −0.01715 |

Comparators: ᵃ transport-ablation control (PERSIST-no-IOT); ᵇ Extra Trees; ᶜ Random Forest; ᵈ HistGradientBoosting. Cross-entropy uses a probability floor of 10⁻¹². Development is the outer fivefold OOF partition grouped by source clone family; E1 is independent experiment 2. Brackets for Brier differences give 95% intervals from 10,000 paired clone bootstrap resamples.

Supplementary Table 2 | Biological comparisons and statistical units.

| Data | Branch | Statistical unit and n | Mean difference [95% CI] | P or q |
| --- | --- | --- | --- | --- |
| TENA, GSE171940 | Invasive EMT | 5 mice, paired | −0.5078 [−0.8589, −0.1567] | P = 0.0159 |
| Cdh, GSE171940 | Invasive EMT | 3 mice, paired | +1.4965 [0.5004, 2.4926] | P = 0.0231 |
| ARTEMIS | IFN/HLA | 79 patients (45 pCR, 34 RD) | +0.2604 [0.1035, 0.4037] | q = 0.0122 |
| ARTEMIS | Invasive EMT | 79 patients (45 pCR, 34 RD) | +0.0030 [−0.0425, 0.0455] | q = 0.8208 |
| MDCK-NBL2, GSE164488 | Epithelial | 3 biological replicate pairs | +0.3234 [0.1204, 0.5264] | P = 0.0206 |
| MDCK-NBL2, GSE164488 | Invasive EMT | 3 biological replicate pairs | −0.4891 [−0.7586, −0.2196] | P = 0.0160 |

Contrasts are ever-EMT minus epithelial lineage for TENA and Cdh, pCR minus RD for ARTEMIS, and siPRRX1 minus siCTR at day 4 under TGFβ for MDCK. Each cohort retains its original branch-score scale. Mouse and MDCK analyses use two-sided paired t tests and raw P values; ARTEMIS intervals use 10,000 patient bootstrap resamples, and q values retain the prespecified Benjamini–Hochberg testing family.

**Extended Data Table 1 | Dataset contracts and information boundaries.**

Source: ED_Table_1_dataset_contract.csv. Field names and values retain the source terminology; blank fields indicate values not supplied in the source. Record numbers identify rows in the source file.

**Analysis group 1**

Fields blank in this group: locked_role.

Fields 1–5; records matched by Record.

| **Record** | **dataset** | **local_path** | **local_status** | **biological_unit** | **modes** |
| --- | --- | --- | --- | --- | --- |
| 1 | GSE140802 | data/processed/GSE140802 | packaged_preprocessed | recipient_mouse | ['retrospective_reconstruction', 'prospective_prediction'] |
| 2 | GSE239651 | data/processed/GSE239651/gse239651_lineage_transition_records.csv | packaged_preprocessed | source_clone_family | ['retrospective_reconstruction'] |
| 5 | macsGESTALT | data/processed/GSE239651/macsgestalt_locked_transition_records.csv | packaged_preprocessed | clone_or_mouse | ['retrospective_reconstruction'] |

Fields 6–7; records matched by Record.

| **Record** | **benchmarks** | **velocity_layers_expected** |
| --- | --- | --- |
| 1 | ['B2_lineage_transition', 'B3_prospective_composition'] | False |
| 2 | ['B2_lineage_transition'] | False |
| 5 | ['B2_lineage_transition'] | False |

**Analysis group 2**

Fields blank in this group: velocity_layers_expected.

Fields 1–5; records matched by Record.

| **Record** | **dataset** | **local_path** | **local_status** | **biological_unit** | **modes** |
| --- | --- | --- | --- | --- | --- |
| 3 | GSE239651_expt1 | data/processed/GSE239651/gse239651_expt1_lineage_panel.parquet | packaged_preprocessed | source_clone_family | ['prospective_prediction'] |
| 4 | GSE239651_expt2 | data/processed/GSE239651/gse239651_expt2_lineage_panel.parquet | packaged_preprocessed | source_clone_family | ['prospective_prediction'] |

Fields 6–7; records matched by Record.

| **Record** | **benchmarks** | **locked_role** |
| --- | --- | --- |
| 3 | ['B3_prospective_composition'] | development_oof |
| 4 | ['B3_prospective_composition'] | external_e1 |

**Extended Data Table 2 | External method versions and repository commits.**

Source: ED_Table_2_method_versions.csv. Field names and values retain the source terminology; blank fields indicate values not supplied in the source. Record numbers identify rows in the source file.

Fields 1–5; records matched by Record.

| **Record** | **method** | **repository** | **commit** | **commit_date** | **local_checkout** |
| --- | --- | --- | --- | --- | --- |
| 1 | Waddington-OT | https://github.com/broadinstitute/wot.git | ca5e94f05699997b01cf5ae13383f9810f0613f6 | 2021-03-31 11:27:28-04:00 | sources/wot |
| 2 | moscot | https://github.com/theislab/moscot.git | 440093ccbb8e70de209157d91da839c55b897821 | 2026-08-25 17:57:58+02:00 | sources/moscot |
| 3 | LineageOT | https://github.com/aforr/LineageOT.git | 6081b402074f7e5934e729e81669aef430219da8 | 2023-07-28 16:25:43+01:00 | sources/lineageot |
| 4 | TIGON | https://github.com/yutongo/TIGON.git | 1ed92cfcc250415fc01b4d344a308b0680cc9635 | 2025-03-22 21:13:35-04:00 | sources/tigon |
| 5 | MIOFlow | https://github.com/KrishnaswamyLab/MIOFlow.git | 36365403d0f23cc3ad1065781c7331bf81debf4e | 2026-06-15 12:51:32-04:00 | sources/mioflow |
| 6 | CellRank_2 | https://github.com/scverse/cellrank.git | d7191d75684c86b58adbb317c8aae7d06f2682f3 | 2026-08-31 11:27:45-04:00 | sources/cellrank2 |
| 7 | PRESCIENT | https://github.com/gifford-lab/prescient.git | 50971c7d495e8763eaa60f83af91f51555ed7ece | 2025-01-20 16:08:40-05:00 | sources/prescient |

Fields 6–7; records matched by Record.

| **Record** | **license_file** | **environment_group** |
| --- | --- | --- |
| 1 | LICENSE | legacy_ot |
| 2 | LICENSE | modern_scverse |
| 3 | LICENSE | legacy_ot |
| 4 | LICENSE | neural_legacy |
| 5 | LICENSE.md | neural_modern |
| 6 | LICENSE | modern_scverse |
| 7 | LICENSE | neural_legacy |

**Extended Data Table 3 | Complete run-level benchmark results.**

Source: ED_Table_3_all_runs_long.csv. Field names and values retain the source terminology; blank fields indicate values not supplied in the source. Record numbers identify rows in the source file.

All 2,616 records have status = success. The fields peak_memory_gb, failure_reason and artifact are blank throughout. Shared run metadata are given above each group; repeat, seed, metric, value and runtime_seconds are listed for every record.

**dataset = identifiable_soft_marginal; benchmark = B1_known_truth; task = known_truth_parameter_and_coupling_recovery; split = synthetic; method = soft_iot; method_version = iot-benchmark-0.1.0**

| **Record** | **repeat** | **seed** | **metric** | **value** | **runtime_seconds** |
| --- | --- | --- | --- | --- | --- |
| 1 | 400 | 20260916.0 | heldout_coupling_relative_frobenius | 0.0207699812570886 | 0.963803000000098 |
| 3 | 400 | 20260917.0 | heldout_coupling_relative_frobenius | 0.0483479491346527 | 1.132093400000031 |
| 5 | 400 | 20260918.0 | heldout_coupling_relative_frobenius | 0.0692560207677088 | 1.0148684000000685 |
| 7 | 400 | 20260919.0 | heldout_coupling_relative_frobenius | 0.0390708011924771 | 0.96355850000009 |
| 9 | 400 | 20260920.0 | heldout_coupling_relative_frobenius | 0.0294202931746749 | 0.8907831000001352 |
| 11 | 2000 | 20260916.0 | heldout_coupling_relative_frobenius | 0.0245554274172476 | 1.0275094999997236 |
| 13 | 2000 | 20260917.0 | heldout_coupling_relative_frobenius | 0.0141101646633622 | 1.0609501000003547 |
| 15 | 2000 | 20260918.0 | heldout_coupling_relative_frobenius | 0.0167051991257651 | 0.9338363000001664 |
| 17 | 2000 | 20260919.0 | heldout_coupling_relative_frobenius | 0.0181263318148039 | 1.0084128000003147 |
| 19 | 2000 | 20260920.0 | heldout_coupling_relative_frobenius | 0.0205997198178066 | 0.8803312000000005 |
| 21 | 10000 | 20260916.0 | heldout_coupling_relative_frobenius | 0.0102713758320923 | 0.945737800000188 |
| 23 | 10000 | 20260917.0 | heldout_coupling_relative_frobenius | 0.0081133524744136 | 0.9946180000001732 |
| 25 | 10000 | 20260918.0 | heldout_coupling_relative_frobenius | 0.0057823675716678 | 0.9884000000001834 |
| 27 | 10000 | 20260919.0 | heldout_coupling_relative_frobenius | 0.0059571819291357 | 1.016133500000251 |
| 29 | 10000 | 20260920.0 | heldout_coupling_relative_frobenius | 0.0112907843262237 | 0.9297907999998642 |
| 181 | 400 | 20260916.0 | coefficient_cosine | 0.9995515514095292 | 0.963803000000098 |
| 183 | 400 | 20260917.0 | coefficient_cosine | 0.9870719458365788 | 1.132093400000031 |
| 185 | 400 | 20260918.0 | coefficient_cosine | 0.9916753028340471 | 1.0148684000000685 |
| 187 | 400 | 20260919.0 | coefficient_cosine | 0.9980348207238112 | 0.96355850000009 |
| 189 | 400 | 20260920.0 | coefficient_cosine | 0.996836900109119 | 0.8907831000001352 |
| 191 | 2000 | 20260916.0 | coefficient_cosine | 0.9960828474529428 | 1.0275094999997236 |
| 193 | 2000 | 20260917.0 | coefficient_cosine | 0.9993891962380566 | 1.0609501000003547 |
| 195 | 2000 | 20260918.0 | coefficient_cosine | 0.9995368749643722 | 0.9338363000001664 |
| 197 | 2000 | 20260919.0 | coefficient_cosine | 0.9984738178844944 | 1.0084128000003147 |
| 199 | 2000 | 20260920.0 | coefficient_cosine | 0.9995603201034752 | 0.8803312000000005 |
| 201 | 10000 | 20260916.0 | coefficient_cosine | 0.9998579564004916 | 0.945737800000188 |
| 203 | 10000 | 20260917.0 | coefficient_cosine | 0.9997504170476496 | 0.9946180000001732 |
| 205 | 10000 | 20260918.0 | coefficient_cosine | 0.9999396659191608 | 0.9884000000001834 |
| 207 | 10000 | 20260919.0 | coefficient_cosine | 0.9998448976822863 | 1.016133500000251 |
| 209 | 10000 | 20260920.0 | coefficient_cosine | 0.999849790398248 | 0.9297907999998642 |
| 361 | 400 | 20260916.0 | coefficient_sign_accuracy | 1.0 | 0.963803000000098 |
| 363 | 400 | 20260917.0 | coefficient_sign_accuracy | 1.0 | 1.132093400000031 |
| 365 | 400 | 20260918.0 | coefficient_sign_accuracy | 0.8333333333333334 | 1.0148684000000685 |
| 367 | 400 | 20260919.0 | coefficient_sign_accuracy | 1.0 | 0.96355850000009 |
| 369 | 400 | 20260920.0 | coefficient_sign_accuracy | 1.0 | 0.8907831000001352 |
| 371 | 2000 | 20260916.0 | coefficient_sign_accuracy | 1.0 | 1.0275094999997236 |
| 373 | 2000 | 20260917.0 | coefficient_sign_accuracy | 1.0 | 1.0609501000003547 |
| 375 | 2000 | 20260918.0 | coefficient_sign_accuracy | 1.0 | 0.9338363000001664 |
| 377 | 2000 | 20260919.0 | coefficient_sign_accuracy | 1.0 | 1.0084128000003147 |
| 379 | 2000 | 20260920.0 | coefficient_sign_accuracy | 1.0 | 0.8803312000000005 |
| 381 | 10000 | 20260916.0 | coefficient_sign_accuracy | 1.0 | 0.945737800000188 |
| 383 | 10000 | 20260917.0 | coefficient_sign_accuracy | 1.0 | 0.9946180000001732 |
| 385 | 10000 | 20260918.0 | coefficient_sign_accuracy | 1.0 | 0.9884000000001834 |
| 387 | 10000 | 20260919.0 | coefficient_sign_accuracy | 1.0 | 1.016133500000251 |
| 389 | 10000 | 20260920.0 | coefficient_sign_accuracy | 1.0 | 0.9297907999998642 |
| 541 | 400 | 20260916.0 | pure_column_minimum_curvature | 0.237363408928104 | 0.963803000000098 |
| 543 | 400 | 20260917.0 | pure_column_minimum_curvature | 0.283165268566421 | 1.132093400000031 |
| 545 | 400 | 20260918.0 | pure_column_minimum_curvature | 0.3292938188792504 | 1.0148684000000685 |
| 547 | 400 | 20260919.0 | pure_column_minimum_curvature | 0.1107706522063267 | 0.96355850000009 |
| 549 | 400 | 20260920.0 | pure_column_minimum_curvature | 0.295903165525857 | 0.8907831000001352 |
| 551 | 2000 | 20260916.0 | pure_column_minimum_curvature | 0.246458566888208 | 1.0275094999997236 |
| 553 | 2000 | 20260917.0 | pure_column_minimum_curvature | 0.2543337140892276 | 1.0609501000003547 |
| 555 | 2000 | 20260918.0 | pure_column_minimum_curvature | 0.3291534254046269 | 0.9338363000001664 |
| 557 | 2000 | 20260919.0 | pure_column_minimum_curvature | 0.1249358936306066 | 1.0084128000003147 |
| 559 | 2000 | 20260920.0 | pure_column_minimum_curvature | 0.2849015992012482 | 0.8803312000000005 |
| 561 | 10000 | 20260916.0 | pure_column_minimum_curvature | 0.2294812202485729 | 0.945737800000188 |
| 563 | 10000 | 20260917.0 | pure_column_minimum_curvature | 0.258035866451678 | 0.9946180000001732 |
| 565 | 10000 | 20260918.0 | pure_column_minimum_curvature | 0.3427808876743654 | 0.9884000000001834 |
| 567 | 10000 | 20260919.0 | pure_column_minimum_curvature | 0.1206422153288144 | 1.016133500000251 |
| 569 | 10000 | 20260920.0 | pure_column_minimum_curvature | 0.2952847126813651 | 0.9297907999998642 |
| 721 | 400 | 20260916.0 | restart_direction_variance | 3.798305901454861e-13 | 0.963803000000098 |
| 723 | 400 | 20260917.0 | restart_direction_variance | 2.788812695977547e-13 | 1.132093400000031 |
| 725 | 400 | 20260918.0 | restart_direction_variance | 3.0719083879883484e-14 | 1.0148684000000685 |
| 727 | 400 | 20260919.0 | restart_direction_variance | 1.396304965294252e-13 | 0.96355850000009 |
| 729 | 400 | 20260920.0 | restart_direction_variance | 2.4505312890274622e-14 | 0.8907831000001352 |
| 731 | 2000 | 20260916.0 | restart_direction_variance | 2.1266280606610045e-13 | 1.0275094999997236 |
| 733 | 2000 | 20260917.0 | restart_direction_variance | 1.916238902879262e-14 | 1.0609501000003547 |
| 735 | 2000 | 20260918.0 | restart_direction_variance | 7.618648496683025e-15 | 0.9338363000001664 |
| 737 | 2000 | 20260919.0 | restart_direction_variance | 3.5994748127953885e-13 | 1.0084128000003147 |
| 739 | 2000 | 20260920.0 | restart_direction_variance | 1.513421867376697e-13 | 0.8803312000000005 |
| 741 | 10000 | 20260916.0 | restart_direction_variance | 1.3659974755404336e-13 | 0.945737800000188 |
| 743 | 10000 | 20260917.0 | restart_direction_variance | 1.451860039091756e-13 | 0.9946180000001732 |
| 745 | 10000 | 20260918.0 | restart_direction_variance | 5.398738512916399e-14 | 0.9884000000001834 |
| 747 | 10000 | 20260919.0 | restart_direction_variance | 1.6498947518012144e-13 | 1.016133500000251 |
| 749 | 10000 | 20260920.0 | restart_direction_variance | 1.0357353217041693e-13 | 0.9297907999998642 |

**dataset = identifiable_soft_marginal; benchmark = B1_known_truth; task = known_truth_parameter_and_coupling_recovery; split = synthetic; method = hard_ot; method_version = iot-benchmark-0.1.0**

| **Record** | **repeat** | **seed** | **metric** | **value** | **runtime_seconds** |
| --- | --- | --- | --- | --- | --- |
| 2 | 400 | 20260916.0 | heldout_coupling_relative_frobenius | 0.08139394538595 | 0.1392136999998001 |
| 4 | 400 | 20260917.0 | heldout_coupling_relative_frobenius | 0.103067401950881 | 0.1504667000003792 |
| 6 | 400 | 20260918.0 | heldout_coupling_relative_frobenius | 0.1101998461388334 | 0.1824176000000079 |
| 8 | 400 | 20260919.0 | heldout_coupling_relative_frobenius | 0.0938627494564115 | 0.1000410000001466 |
| 10 | 400 | 20260920.0 | heldout_coupling_relative_frobenius | 0.0908772555966051 | 0.1080259000000296 |
| 12 | 2000 | 20260916.0 | heldout_coupling_relative_frobenius | 0.038107670109363 | 0.1905112999997982 |
| 14 | 2000 | 20260917.0 | heldout_coupling_relative_frobenius | 0.0339287372982511 | 0.1462578999999095 |
| 16 | 2000 | 20260918.0 | heldout_coupling_relative_frobenius | 0.0464478470486312 | 0.1552415999999539 |
| 18 | 2000 | 20260919.0 | heldout_coupling_relative_frobenius | 0.0443618739585832 | 0.0984231999996154 |
| 20 | 2000 | 20260920.0 | heldout_coupling_relative_frobenius | 0.0297066091114038 | 0.0869949999996606 |
| 22 | 10000 | 20260916.0 | heldout_coupling_relative_frobenius | 0.0132257697783976 | 0.1422238999998626 |
| 24 | 10000 | 20260917.0 | heldout_coupling_relative_frobenius | 0.0221702102194982 | 0.1672464999996918 |
| 26 | 10000 | 20260918.0 | heldout_coupling_relative_frobenius | 0.0317945331356629 | 0.1465640000001258 |
| 28 | 10000 | 20260919.0 | heldout_coupling_relative_frobenius | 0.0201150755156233 | 0.1013747000001785 |
| 30 | 10000 | 20260920.0 | heldout_coupling_relative_frobenius | 0.0167491157153561 | 0.0945558999997047 |
| 182 | 400 | 20260916.0 | coefficient_cosine | 0.7712668930877951 | 0.1392136999998001 |
| 184 | 400 | 20260917.0 | coefficient_cosine | 0.484518532715283 | 0.1504667000003792 |
| 186 | 400 | 20260918.0 | coefficient_cosine | -0.0632766628133919 | 0.1824176000000079 |
| 188 | 400 | 20260919.0 | coefficient_cosine | 0.8666165560203433 | 0.1000410000001466 |
| 190 | 400 | 20260920.0 | coefficient_cosine | 0.5702679681928934 | 0.1080259000000296 |
| 192 | 2000 | 20260916.0 | coefficient_cosine | 0.3190450724977434 | 0.1905112999997982 |
| 194 | 2000 | 20260917.0 | coefficient_cosine | 0.3295995061156788 | 0.1462578999999095 |
| 196 | 2000 | 20260918.0 | coefficient_cosine | 0.4521441527576409 | 0.1552415999999539 |
| 198 | 2000 | 20260919.0 | coefficient_cosine | -0.3488767786058773 | 0.0984231999996154 |
| 200 | 2000 | 20260920.0 | coefficient_cosine | -0.0758335875606693 | 0.0869949999996606 |
| 202 | 10000 | 20260916.0 | coefficient_cosine | 0.316375537876772 | 0.1422238999998626 |
| 204 | 10000 | 20260917.0 | coefficient_cosine | 0.0264409597643551 | 0.1672464999996918 |
| 206 | 10000 | 20260918.0 | coefficient_cosine | 0.4514524370205405 | 0.1465640000001258 |
| 208 | 10000 | 20260919.0 | coefficient_cosine | 0.1699418969750258 | 0.1013747000001785 |
| 210 | 10000 | 20260920.0 | coefficient_cosine | 0.5713784211796334 | 0.0945558999997047 |
| 362 | 400 | 20260916.0 | coefficient_sign_accuracy | 0.8333333333333334 | 0.1392136999998001 |
| 364 | 400 | 20260917.0 | coefficient_sign_accuracy | 0.6666666666666666 | 0.1504667000003792 |
| 366 | 400 | 20260918.0 | coefficient_sign_accuracy | 0.5 | 0.1824176000000079 |
| 368 | 400 | 20260919.0 | coefficient_sign_accuracy | 1.0 | 0.1000410000001466 |
| 370 | 400 | 20260920.0 | coefficient_sign_accuracy | 1.0 | 0.1080259000000296 |
| 372 | 2000 | 20260916.0 | coefficient_sign_accuracy | 0.6666666666666666 | 0.1905112999997982 |
| 374 | 2000 | 20260917.0 | coefficient_sign_accuracy | 0.6666666666666666 | 0.1462578999999095 |
| 376 | 2000 | 20260918.0 | coefficient_sign_accuracy | 0.5 | 0.1552415999999539 |
| 378 | 2000 | 20260919.0 | coefficient_sign_accuracy | 0.3333333333333333 | 0.0984231999996154 |
| 380 | 2000 | 20260920.0 | coefficient_sign_accuracy | 0.6666666666666666 | 0.0869949999996606 |
| 382 | 10000 | 20260916.0 | coefficient_sign_accuracy | 0.6666666666666666 | 0.1422238999998626 |
| 384 | 10000 | 20260917.0 | coefficient_sign_accuracy | 0.5 | 0.1672464999996918 |
| 386 | 10000 | 20260918.0 | coefficient_sign_accuracy | 0.5 | 0.1465640000001258 |
| 388 | 10000 | 20260919.0 | coefficient_sign_accuracy | 0.6666666666666666 | 0.1013747000001785 |
| 390 | 10000 | 20260920.0 | coefficient_sign_accuracy | 1.0 | 0.0945558999997047 |
| 542 | 400 | 20260916.0 | pure_column_minimum_curvature | 0.0 | 0.1392136999998001 |
| 544 | 400 | 20260917.0 | pure_column_minimum_curvature | 0.0 | 0.1504667000003792 |
| 546 | 400 | 20260918.0 | pure_column_minimum_curvature | -8.88178419700125e-10 | 0.1824176000000079 |
| 548 | 400 | 20260919.0 | pure_column_minimum_curvature | -4.4408920985006256e-10 | 0.1000410000001466 |
| 550 | 400 | 20260920.0 | pure_column_minimum_curvature | 0.0 | 0.1080259000000296 |
| 552 | 2000 | 20260916.0 | pure_column_minimum_curvature | 0.0 | 0.1905112999997982 |
| 554 | 2000 | 20260917.0 | pure_column_minimum_curvature | 0.0 | 0.1462578999999095 |
| 556 | 2000 | 20260918.0 | pure_column_minimum_curvature | 0.0 | 0.1552415999999539 |
| 558 | 2000 | 20260919.0 | pure_column_minimum_curvature | 0.0 | 0.0984231999996154 |
| 560 | 2000 | 20260920.0 | pure_column_minimum_curvature | -8.88178419700125e-10 | 0.0869949999996606 |
| 562 | 10000 | 20260916.0 | pure_column_minimum_curvature | -1.7763568394002503e-09 | 0.1422238999998626 |
| 564 | 10000 | 20260917.0 | pure_column_minimum_curvature | 0.0 | 0.1672464999996918 |
| 566 | 10000 | 20260918.0 | pure_column_minimum_curvature | 0.0 | 0.1465640000001258 |
| 568 | 10000 | 20260919.0 | pure_column_minimum_curvature | 0.0 | 0.1013747000001785 |
| 570 | 10000 | 20260920.0 | pure_column_minimum_curvature | 0.0 | 0.0945558999997047 |
| 722 | 400 | 20260916.0 | restart_direction_variance | 0.0647746140579003 | 0.1392136999998001 |
| 724 | 400 | 20260917.0 | restart_direction_variance | 0.0986833835986874 | 0.1504667000003792 |
| 726 | 400 | 20260918.0 | restart_direction_variance | 0.0775051751089891 | 0.1824176000000079 |
| 728 | 400 | 20260919.0 | restart_direction_variance | 0.1286257574375557 | 0.1000410000001466 |
| 730 | 400 | 20260920.0 | restart_direction_variance | 0.1436545875896556 | 0.1080259000000296 |
| 732 | 2000 | 20260916.0 | restart_direction_variance | 0.060894247222499 | 0.1905112999997982 |
| 734 | 2000 | 20260917.0 | restart_direction_variance | 0.1010814282895829 | 0.1462578999999095 |
| 736 | 2000 | 20260918.0 | restart_direction_variance | 0.0793811458598071 | 0.1552415999999539 |
| 738 | 2000 | 20260919.0 | restart_direction_variance | 0.1422373496909778 | 0.0984231999996154 |
| 740 | 2000 | 20260920.0 | restart_direction_variance | 0.1460841871730005 | 0.0869949999996606 |
| 742 | 10000 | 20260916.0 | restart_direction_variance | 0.0616026736463164 | 0.1422238999998626 |
| 744 | 10000 | 20260917.0 | restart_direction_variance | 0.1008099981187067 | 0.1672464999996918 |
| 746 | 10000 | 20260918.0 | restart_direction_variance | 0.0798545869259997 | 0.1465640000001258 |
| 748 | 10000 | 20260919.0 | restart_direction_variance | 0.144255192016548 | 0.1013747000001785 |
| 750 | 10000 | 20260920.0 | restart_direction_variance | 0.1367698752419833 | 0.0945558999997047 |

**dataset = hard_marginal_flat_direction; benchmark = B1_known_truth; task = known_truth_parameter_and_coupling_recovery; split = synthetic; method = soft_iot; method_version = iot-benchmark-0.1.0**

| **Record** | **repeat** | **seed** | **metric** | **value** | **runtime_seconds** |
| --- | --- | --- | --- | --- | --- |
| 31 | 400 | 20260916.0 | heldout_coupling_relative_frobenius | 0.0207699812570886 | 1.5232843999997383 |
| 33 | 400 | 20260917.0 | heldout_coupling_relative_frobenius | 0.0483479491346527 | 1.5501485000004325 |
| 35 | 400 | 20260918.0 | heldout_coupling_relative_frobenius | 0.0692560207677088 | 1.3613313000000744 |
| 37 | 400 | 20260919.0 | heldout_coupling_relative_frobenius | 0.0390708011924771 | 1.1765565000000606 |
| 39 | 400 | 20260920.0 | heldout_coupling_relative_frobenius | 0.0294202931746749 | 1.1393797000000632 |
| 41 | 2000 | 20260916.0 | heldout_coupling_relative_frobenius | 0.0245554274172476 | 1.1556378999998742 |
| 43 | 2000 | 20260917.0 | heldout_coupling_relative_frobenius | 0.0141101646633622 | 1.0604648999997153 |
| 45 | 2000 | 20260918.0 | heldout_coupling_relative_frobenius | 0.0167051991257651 | 1.0904113000001416 |
| 47 | 2000 | 20260919.0 | heldout_coupling_relative_frobenius | 0.0181263318148039 | 0.972682700000405 |
| 49 | 2000 | 20260920.0 | heldout_coupling_relative_frobenius | 0.0205997198178066 | 1.0363537000002907 |
| 51 | 10000 | 20260916.0 | heldout_coupling_relative_frobenius | 0.0102713758320923 | 0.9835258000002796 |
| 53 | 10000 | 20260917.0 | heldout_coupling_relative_frobenius | 0.0081133524744136 | 1.0423848000000362 |
| 55 | 10000 | 20260918.0 | heldout_coupling_relative_frobenius | 0.0057823675716678 | 1.0388536000000386 |
| 57 | 10000 | 20260919.0 | heldout_coupling_relative_frobenius | 0.0059571819291357 | 1.0565594999998211 |
| 59 | 10000 | 20260920.0 | heldout_coupling_relative_frobenius | 0.0112907843262237 | 0.9896129000003384 |
| 211 | 400 | 20260916.0 | coefficient_cosine | 0.9995515514095292 | 1.5232843999997383 |
| 213 | 400 | 20260917.0 | coefficient_cosine | 0.9870719458365788 | 1.5501485000004325 |
| 215 | 400 | 20260918.0 | coefficient_cosine | 0.9916753028340471 | 1.3613313000000744 |
| 217 | 400 | 20260919.0 | coefficient_cosine | 0.9980348207238112 | 1.1765565000000606 |
| 219 | 400 | 20260920.0 | coefficient_cosine | 0.996836900109119 | 1.1393797000000632 |
| 221 | 2000 | 20260916.0 | coefficient_cosine | 0.9960828474529428 | 1.1556378999998742 |
| 223 | 2000 | 20260917.0 | coefficient_cosine | 0.9993891962380566 | 1.0604648999997153 |
| 225 | 2000 | 20260918.0 | coefficient_cosine | 0.9995368749643722 | 1.0904113000001416 |
| 227 | 2000 | 20260919.0 | coefficient_cosine | 0.9984738178844944 | 0.972682700000405 |
| 229 | 2000 | 20260920.0 | coefficient_cosine | 0.9995603201034752 | 1.0363537000002907 |
| 231 | 10000 | 20260916.0 | coefficient_cosine | 0.9998579564004916 | 0.9835258000002796 |
| 233 | 10000 | 20260917.0 | coefficient_cosine | 0.9997504170476496 | 1.0423848000000362 |
| 235 | 10000 | 20260918.0 | coefficient_cosine | 0.9999396659191608 | 1.0388536000000386 |
| 237 | 10000 | 20260919.0 | coefficient_cosine | 0.9998448976822863 | 1.0565594999998211 |
| 239 | 10000 | 20260920.0 | coefficient_cosine | 0.999849790398248 | 0.9896129000003384 |
| 391 | 400 | 20260916.0 | coefficient_sign_accuracy | 1.0 | 1.5232843999997383 |
| 393 | 400 | 20260917.0 | coefficient_sign_accuracy | 1.0 | 1.5501485000004325 |
| 395 | 400 | 20260918.0 | coefficient_sign_accuracy | 0.8333333333333334 | 1.3613313000000744 |
| 397 | 400 | 20260919.0 | coefficient_sign_accuracy | 1.0 | 1.1765565000000606 |
| 399 | 400 | 20260920.0 | coefficient_sign_accuracy | 1.0 | 1.1393797000000632 |
| 401 | 2000 | 20260916.0 | coefficient_sign_accuracy | 1.0 | 1.1556378999998742 |
| 403 | 2000 | 20260917.0 | coefficient_sign_accuracy | 1.0 | 1.0604648999997153 |
| 405 | 2000 | 20260918.0 | coefficient_sign_accuracy | 1.0 | 1.0904113000001416 |
| 407 | 2000 | 20260919.0 | coefficient_sign_accuracy | 1.0 | 0.972682700000405 |
| 409 | 2000 | 20260920.0 | coefficient_sign_accuracy | 1.0 | 1.0363537000002907 |
| 411 | 10000 | 20260916.0 | coefficient_sign_accuracy | 1.0 | 0.9835258000002796 |
| 413 | 10000 | 20260917.0 | coefficient_sign_accuracy | 1.0 | 1.0423848000000362 |
| 415 | 10000 | 20260918.0 | coefficient_sign_accuracy | 1.0 | 1.0388536000000386 |
| 417 | 10000 | 20260919.0 | coefficient_sign_accuracy | 1.0 | 1.0565594999998211 |
| 419 | 10000 | 20260920.0 | coefficient_sign_accuracy | 1.0 | 0.9896129000003384 |
| 571 | 400 | 20260916.0 | pure_column_minimum_curvature | 0.237363408928104 | 1.5232843999997383 |
| 573 | 400 | 20260917.0 | pure_column_minimum_curvature | 0.283165268566421 | 1.5501485000004325 |
| 575 | 400 | 20260918.0 | pure_column_minimum_curvature | 0.3292938188792504 | 1.3613313000000744 |
| 577 | 400 | 20260919.0 | pure_column_minimum_curvature | 0.1107706522063267 | 1.1765565000000606 |
| 579 | 400 | 20260920.0 | pure_column_minimum_curvature | 0.295903165525857 | 1.1393797000000632 |
| 581 | 2000 | 20260916.0 | pure_column_minimum_curvature | 0.246458566888208 | 1.1556378999998742 |
| 583 | 2000 | 20260917.0 | pure_column_minimum_curvature | 0.2543337140892276 | 1.0604648999997153 |
| 585 | 2000 | 20260918.0 | pure_column_minimum_curvature | 0.3291534254046269 | 1.0904113000001416 |
| 587 | 2000 | 20260919.0 | pure_column_minimum_curvature | 0.1249358936306066 | 0.972682700000405 |
| 589 | 2000 | 20260920.0 | pure_column_minimum_curvature | 0.2849015992012482 | 1.0363537000002907 |
| 591 | 10000 | 20260916.0 | pure_column_minimum_curvature | 0.2294812202485729 | 0.9835258000002796 |
| 593 | 10000 | 20260917.0 | pure_column_minimum_curvature | 0.258035866451678 | 1.0423848000000362 |
| 595 | 10000 | 20260918.0 | pure_column_minimum_curvature | 0.3427808876743654 | 1.0388536000000386 |
| 597 | 10000 | 20260919.0 | pure_column_minimum_curvature | 0.1206422153288144 | 1.0565594999998211 |
| 599 | 10000 | 20260920.0 | pure_column_minimum_curvature | 0.2952847126813651 | 0.9896129000003384 |
| 751 | 400 | 20260916.0 | restart_direction_variance | 3.798305901454861e-13 | 1.5232843999997383 |
| 753 | 400 | 20260917.0 | restart_direction_variance | 2.788812695977547e-13 | 1.5501485000004325 |
| 755 | 400 | 20260918.0 | restart_direction_variance | 3.0719083879883484e-14 | 1.3613313000000744 |
| 757 | 400 | 20260919.0 | restart_direction_variance | 1.396304965294252e-13 | 1.1765565000000606 |
| 759 | 400 | 20260920.0 | restart_direction_variance | 2.4505312890274622e-14 | 1.1393797000000632 |
| 761 | 2000 | 20260916.0 | restart_direction_variance | 2.1266280606610045e-13 | 1.1556378999998742 |
| 763 | 2000 | 20260917.0 | restart_direction_variance | 1.916238902879262e-14 | 1.0604648999997153 |
| 765 | 2000 | 20260918.0 | restart_direction_variance | 7.618648496683025e-15 | 1.0904113000001416 |
| 767 | 2000 | 20260919.0 | restart_direction_variance | 3.5994748127953885e-13 | 0.972682700000405 |
| 769 | 2000 | 20260920.0 | restart_direction_variance | 1.513421867376697e-13 | 1.0363537000002907 |
| 771 | 10000 | 20260916.0 | restart_direction_variance | 1.3659974755404336e-13 | 0.9835258000002796 |
| 773 | 10000 | 20260917.0 | restart_direction_variance | 1.451860039091756e-13 | 1.0423848000000362 |
| 775 | 10000 | 20260918.0 | restart_direction_variance | 5.398738512916399e-14 | 1.0388536000000386 |
| 777 | 10000 | 20260919.0 | restart_direction_variance | 1.6498947518012144e-13 | 1.0565594999998211 |
| 779 | 10000 | 20260920.0 | restart_direction_variance | 1.0357353217041693e-13 | 0.9896129000003384 |

**dataset = hard_marginal_flat_direction; benchmark = B1_known_truth; task = known_truth_parameter_and_coupling_recovery; split = synthetic; method = hard_ot; method_version = iot-benchmark-0.1.0**

| **Record** | **repeat** | **seed** | **metric** | **value** | **runtime_seconds** |
| --- | --- | --- | --- | --- | --- |
| 32 | 400 | 20260916.0 | heldout_coupling_relative_frobenius | 0.08139394538595 | 0.2748335000001134 |
| 34 | 400 | 20260917.0 | heldout_coupling_relative_frobenius | 0.103067401950881 | 0.2171367000000827 |
| 36 | 400 | 20260918.0 | heldout_coupling_relative_frobenius | 0.1101998461388334 | 0.2224190000001726 |
| 38 | 400 | 20260919.0 | heldout_coupling_relative_frobenius | 0.0938627494564115 | 0.1316170000000056 |
| 40 | 400 | 20260920.0 | heldout_coupling_relative_frobenius | 0.0908772555966051 | 0.1126588000001902 |
| 42 | 2000 | 20260916.0 | heldout_coupling_relative_frobenius | 0.038107670109363 | 0.1678867000000536 |
| 44 | 2000 | 20260917.0 | heldout_coupling_relative_frobenius | 0.0339287372982511 | 0.1387133999996876 |
| 46 | 2000 | 20260918.0 | heldout_coupling_relative_frobenius | 0.0464478470486312 | 0.160927299999912 |
| 48 | 2000 | 20260919.0 | heldout_coupling_relative_frobenius | 0.0443618739585832 | 0.1193326000002343 |
| 50 | 2000 | 20260920.0 | heldout_coupling_relative_frobenius | 0.0297066091114038 | 0.0963798000002498 |
| 52 | 10000 | 20260916.0 | heldout_coupling_relative_frobenius | 0.0132257697783976 | 0.1285948999998254 |
| 54 | 10000 | 20260917.0 | heldout_coupling_relative_frobenius | 0.0221702102194982 | 0.1711081999997077 |
| 56 | 10000 | 20260918.0 | heldout_coupling_relative_frobenius | 0.0317945331356629 | 0.159653299999718 |
| 58 | 10000 | 20260919.0 | heldout_coupling_relative_frobenius | 0.0201150755156233 | 0.1007915999998658 |
| 60 | 10000 | 20260920.0 | heldout_coupling_relative_frobenius | 0.0167491157153561 | 0.1319333000001279 |
| 212 | 400 | 20260916.0 | coefficient_cosine | 0.7712668930877951 | 0.2748335000001134 |
| 214 | 400 | 20260917.0 | coefficient_cosine | 0.484518532715283 | 0.2171367000000827 |
| 216 | 400 | 20260918.0 | coefficient_cosine | -0.0632766628133919 | 0.2224190000001726 |
| 218 | 400 | 20260919.0 | coefficient_cosine | 0.8666165560203433 | 0.1316170000000056 |
| 220 | 400 | 20260920.0 | coefficient_cosine | 0.5702679681928934 | 0.1126588000001902 |
| 222 | 2000 | 20260916.0 | coefficient_cosine | 0.3190450724977434 | 0.1678867000000536 |
| 224 | 2000 | 20260917.0 | coefficient_cosine | 0.3295995061156788 | 0.1387133999996876 |
| 226 | 2000 | 20260918.0 | coefficient_cosine | 0.4521441527576409 | 0.160927299999912 |
| 228 | 2000 | 20260919.0 | coefficient_cosine | -0.3488767786058773 | 0.1193326000002343 |
| 230 | 2000 | 20260920.0 | coefficient_cosine | -0.0758335875606693 | 0.0963798000002498 |
| 232 | 10000 | 20260916.0 | coefficient_cosine | 0.316375537876772 | 0.1285948999998254 |
| 234 | 10000 | 20260917.0 | coefficient_cosine | 0.0264409597643551 | 0.1711081999997077 |
| 236 | 10000 | 20260918.0 | coefficient_cosine | 0.4514524370205405 | 0.159653299999718 |
| 238 | 10000 | 20260919.0 | coefficient_cosine | 0.1699418969750258 | 0.1007915999998658 |
| 240 | 10000 | 20260920.0 | coefficient_cosine | 0.5713784211796334 | 0.1319333000001279 |
| 392 | 400 | 20260916.0 | coefficient_sign_accuracy | 0.8333333333333334 | 0.2748335000001134 |
| 394 | 400 | 20260917.0 | coefficient_sign_accuracy | 0.6666666666666666 | 0.2171367000000827 |
| 396 | 400 | 20260918.0 | coefficient_sign_accuracy | 0.5 | 0.2224190000001726 |
| 398 | 400 | 20260919.0 | coefficient_sign_accuracy | 1.0 | 0.1316170000000056 |
| 400 | 400 | 20260920.0 | coefficient_sign_accuracy | 1.0 | 0.1126588000001902 |
| 402 | 2000 | 20260916.0 | coefficient_sign_accuracy | 0.6666666666666666 | 0.1678867000000536 |
| 404 | 2000 | 20260917.0 | coefficient_sign_accuracy | 0.6666666666666666 | 0.1387133999996876 |
| 406 | 2000 | 20260918.0 | coefficient_sign_accuracy | 0.5 | 0.160927299999912 |
| 408 | 2000 | 20260919.0 | coefficient_sign_accuracy | 0.3333333333333333 | 0.1193326000002343 |
| 410 | 2000 | 20260920.0 | coefficient_sign_accuracy | 0.6666666666666666 | 0.0963798000002498 |
| 412 | 10000 | 20260916.0 | coefficient_sign_accuracy | 0.6666666666666666 | 0.1285948999998254 |
| 414 | 10000 | 20260917.0 | coefficient_sign_accuracy | 0.5 | 0.1711081999997077 |
| 416 | 10000 | 20260918.0 | coefficient_sign_accuracy | 0.5 | 0.159653299999718 |
| 418 | 10000 | 20260919.0 | coefficient_sign_accuracy | 0.6666666666666666 | 0.1007915999998658 |
| 420 | 10000 | 20260920.0 | coefficient_sign_accuracy | 1.0 | 0.1319333000001279 |
| 572 | 400 | 20260916.0 | pure_column_minimum_curvature | 0.0 | 0.2748335000001134 |
| 574 | 400 | 20260917.0 | pure_column_minimum_curvature | 0.0 | 0.2171367000000827 |
| 576 | 400 | 20260918.0 | pure_column_minimum_curvature | -8.88178419700125e-10 | 0.2224190000001726 |
| 578 | 400 | 20260919.0 | pure_column_minimum_curvature | -4.4408920985006256e-10 | 0.1316170000000056 |
| 580 | 400 | 20260920.0 | pure_column_minimum_curvature | 0.0 | 0.1126588000001902 |
| 582 | 2000 | 20260916.0 | pure_column_minimum_curvature | 0.0 | 0.1678867000000536 |
| 584 | 2000 | 20260917.0 | pure_column_minimum_curvature | 0.0 | 0.1387133999996876 |
| 586 | 2000 | 20260918.0 | pure_column_minimum_curvature | 0.0 | 0.160927299999912 |
| 588 | 2000 | 20260919.0 | pure_column_minimum_curvature | 0.0 | 0.1193326000002343 |
| 590 | 2000 | 20260920.0 | pure_column_minimum_curvature | -8.88178419700125e-10 | 0.0963798000002498 |
| 592 | 10000 | 20260916.0 | pure_column_minimum_curvature | -1.7763568394002503e-09 | 0.1285948999998254 |
| 594 | 10000 | 20260917.0 | pure_column_minimum_curvature | 0.0 | 0.1711081999997077 |
| 596 | 10000 | 20260918.0 | pure_column_minimum_curvature | 0.0 | 0.159653299999718 |
| 598 | 10000 | 20260919.0 | pure_column_minimum_curvature | 0.0 | 0.1007915999998658 |
| 600 | 10000 | 20260920.0 | pure_column_minimum_curvature | 0.0 | 0.1319333000001279 |
| 752 | 400 | 20260916.0 | restart_direction_variance | 0.0647746140579003 | 0.2748335000001134 |
| 754 | 400 | 20260917.0 | restart_direction_variance | 0.0986833835986874 | 0.2171367000000827 |
| 756 | 400 | 20260918.0 | restart_direction_variance | 0.0775051751089891 | 0.2224190000001726 |
| 758 | 400 | 20260919.0 | restart_direction_variance | 0.1286257574375557 | 0.1316170000000056 |
| 760 | 400 | 20260920.0 | restart_direction_variance | 0.1436545875896556 | 0.1126588000001902 |
| 762 | 2000 | 20260916.0 | restart_direction_variance | 0.060894247222499 | 0.1678867000000536 |
| 764 | 2000 | 20260917.0 | restart_direction_variance | 0.1010814282895829 | 0.1387133999996876 |
| 766 | 2000 | 20260918.0 | restart_direction_variance | 0.0793811458598071 | 0.160927299999912 |
| 768 | 2000 | 20260919.0 | restart_direction_variance | 0.1422373496909778 | 0.1193326000002343 |
| 770 | 2000 | 20260920.0 | restart_direction_variance | 0.1460841871730005 | 0.0963798000002498 |
| 772 | 10000 | 20260916.0 | restart_direction_variance | 0.0616026736463164 | 0.1285948999998254 |
| 774 | 10000 | 20260917.0 | restart_direction_variance | 0.1008099981187067 | 0.1711081999997077 |
| 776 | 10000 | 20260918.0 | restart_direction_variance | 0.0798545869259997 | 0.159653299999718 |
| 778 | 10000 | 20260919.0 | restart_direction_variance | 0.144255192016548 | 0.1007915999998658 |
| 780 | 10000 | 20260920.0 | restart_direction_variance | 0.1367698752419833 | 0.1319333000001279 |

**dataset = equivalent_fit_multiple_parameters; benchmark = B1_known_truth; task = known_truth_parameter_and_coupling_recovery; split = synthetic; method = soft_iot; method_version = iot-benchmark-0.1.0**

| **Record** | **repeat** | **seed** | **metric** | **value** | **runtime_seconds** |
| --- | --- | --- | --- | --- | --- |
| 61 | 400 | 20260916.0 | heldout_coupling_relative_frobenius | 0.0427323550456116 | 1.07515420000027 |
| 63 | 400 | 20260917.0 | heldout_coupling_relative_frobenius | 0.0214033771293042 | 0.9713815000000068 |
| 65 | 400 | 20260918.0 | heldout_coupling_relative_frobenius | 0.0434081974442864 | 0.8395845000000008 |
| 67 | 400 | 20260919.0 | heldout_coupling_relative_frobenius | 0.0364069158179338 | 1.1241503999999622 |
| 69 | 400 | 20260920.0 | heldout_coupling_relative_frobenius | 0.0296928592835332 | 1.0202758999998878 |
| 71 | 2000 | 20260916.0 | heldout_coupling_relative_frobenius | 0.0163441461380098 | 1.0049973999998656 |
| 73 | 2000 | 20260917.0 | heldout_coupling_relative_frobenius | 0.0348825800328826 | 1.0348675000000185 |
| 75 | 2000 | 20260918.0 | heldout_coupling_relative_frobenius | 0.0271331947292079 | 0.920255100000304 |
| 77 | 2000 | 20260919.0 | heldout_coupling_relative_frobenius | 0.0119854824622326 | 1.1910577000003286 |
| 79 | 2000 | 20260920.0 | heldout_coupling_relative_frobenius | 0.0150904462852003 | 0.9830117999999856 |
| 81 | 10000 | 20260916.0 | heldout_coupling_relative_frobenius | 0.0111491684792322 | 0.9429312000002028 |
| 83 | 10000 | 20260917.0 | heldout_coupling_relative_frobenius | 0.0083046641255133 | 0.9786874000001262 |
| 85 | 10000 | 20260918.0 | heldout_coupling_relative_frobenius | 0.0047919530726179 | 0.8395958000000974 |
| 87 | 10000 | 20260919.0 | heldout_coupling_relative_frobenius | 0.0099398650088525 | 1.0072248999999829 |
| 89 | 10000 | 20260920.0 | heldout_coupling_relative_frobenius | 0.0062984941245662 | 1.184292199999618 |
| 241 | 400 | 20260916.0 | coefficient_cosine | 0.9933973250804814 | 1.07515420000027 |
| 243 | 400 | 20260917.0 | coefficient_cosine | 0.9962197522673718 | 0.9713815000000068 |
| 245 | 400 | 20260918.0 | coefficient_cosine | 0.9970543774757166 | 0.8395845000000008 |
| 247 | 400 | 20260919.0 | coefficient_cosine | 0.9882838544868005 | 1.1241503999999622 |
| 249 | 400 | 20260920.0 | coefficient_cosine | 0.9961502116092884 | 1.0202758999998878 |
| 251 | 2000 | 20260916.0 | coefficient_cosine | 0.9991766712991011 | 1.0049973999998656 |
| 253 | 2000 | 20260917.0 | coefficient_cosine | 0.9976616705972364 | 1.0348675000000185 |
| 255 | 2000 | 20260918.0 | coefficient_cosine | 0.9972327813034714 | 0.920255100000304 |
| 257 | 2000 | 20260919.0 | coefficient_cosine | 0.9987712549925823 | 1.1910577000003286 |
| 259 | 2000 | 20260920.0 | coefficient_cosine | 0.998865194550723 | 0.9830117999999856 |
| 261 | 10000 | 20260916.0 | coefficient_cosine | 0.9997310842464964 | 0.9429312000002028 |
| 263 | 10000 | 20260917.0 | coefficient_cosine | 0.9999423232429754 | 0.9786874000001262 |
| 265 | 10000 | 20260918.0 | coefficient_cosine | 0.9999084203638527 | 0.8395958000000974 |
| 267 | 10000 | 20260919.0 | coefficient_cosine | 0.9992575602603914 | 1.0072248999999829 |
| 269 | 10000 | 20260920.0 | coefficient_cosine | 0.9995831487565456 | 1.184292199999618 |
| 421 | 400 | 20260916.0 | coefficient_sign_accuracy | 1.0 | 1.07515420000027 |
| 423 | 400 | 20260917.0 | coefficient_sign_accuracy | 1.0 | 0.9713815000000068 |
| 425 | 400 | 20260918.0 | coefficient_sign_accuracy | 0.75 | 0.8395845000000008 |
| 427 | 400 | 20260919.0 | coefficient_sign_accuracy | 1.0 | 1.1241503999999622 |
| 429 | 400 | 20260920.0 | coefficient_sign_accuracy | 1.0 | 1.0202758999998878 |
| 431 | 2000 | 20260916.0 | coefficient_sign_accuracy | 1.0 | 1.0049973999998656 |
| 433 | 2000 | 20260917.0 | coefficient_sign_accuracy | 1.0 | 1.0348675000000185 |
| 435 | 2000 | 20260918.0 | coefficient_sign_accuracy | 1.0 | 0.920255100000304 |
| 437 | 2000 | 20260919.0 | coefficient_sign_accuracy | 1.0 | 1.1910577000003286 |
| 439 | 2000 | 20260920.0 | coefficient_sign_accuracy | 1.0 | 0.9830117999999856 |
| 441 | 10000 | 20260916.0 | coefficient_sign_accuracy | 1.0 | 0.9429312000002028 |
| 443 | 10000 | 20260917.0 | coefficient_sign_accuracy | 1.0 | 0.9786874000001262 |
| 445 | 10000 | 20260918.0 | coefficient_sign_accuracy | 1.0 | 0.8395958000000974 |
| 447 | 10000 | 20260919.0 | coefficient_sign_accuracy | 1.0 | 1.0072248999999829 |
| 449 | 10000 | 20260920.0 | coefficient_sign_accuracy | 1.0 | 1.184292199999618 |
| 601 | 400 | 20260916.0 | pure_column_minimum_curvature | 0.1199783619121319 | 1.07515420000027 |
| 603 | 400 | 20260917.0 | pure_column_minimum_curvature | 0.2637516596237788 | 0.9713815000000068 |
| 605 | 400 | 20260918.0 | pure_column_minimum_curvature | 0.3244111845823738 | 0.8395845000000008 |
| 607 | 400 | 20260919.0 | pure_column_minimum_curvature | 0.1272069249935725 | 1.1241503999999622 |
| 609 | 400 | 20260920.0 | pure_column_minimum_curvature | 0.2829631009504396 | 1.0202758999998878 |
| 611 | 2000 | 20260916.0 | pure_column_minimum_curvature | 0.1437502330503548 | 1.0049973999998656 |
| 613 | 2000 | 20260917.0 | pure_column_minimum_curvature | 0.2399015306409069 | 1.0348675000000185 |
| 615 | 2000 | 20260918.0 | pure_column_minimum_curvature | 0.324619757741118 | 0.920255100000304 |
| 617 | 2000 | 20260919.0 | pure_column_minimum_curvature | 0.1059039700379571 | 1.1910577000003286 |
| 619 | 2000 | 20260920.0 | pure_column_minimum_curvature | 0.2641526317681553 | 0.9830117999999856 |
| 621 | 10000 | 20260916.0 | pure_column_minimum_curvature | 0.1565104570389053 | 0.9429312000002028 |
| 623 | 10000 | 20260917.0 | pure_column_minimum_curvature | 0.2716886933207263 | 0.9786874000001262 |
| 625 | 10000 | 20260918.0 | pure_column_minimum_curvature | 0.3175619514017569 | 0.8395958000000974 |
| 627 | 10000 | 20260919.0 | pure_column_minimum_curvature | 0.1005256939912158 | 1.0072248999999829 |
| 629 | 10000 | 20260920.0 | pure_column_minimum_curvature | 0.2568372825528797 | 1.184292199999618 |
| 781 | 400 | 20260916.0 | restart_direction_variance | 1.3898888218933465e-13 | 1.07515420000027 |
| 783 | 400 | 20260917.0 | restart_direction_variance | 1.670706390901926e-13 | 0.9713815000000068 |
| 785 | 400 | 20260918.0 | restart_direction_variance | 2.4826804479267067e-14 | 0.8395845000000008 |
| 787 | 400 | 20260919.0 | restart_direction_variance | 1.0844987060222321e-13 | 1.1241503999999622 |
| 789 | 400 | 20260920.0 | restart_direction_variance | 2.123970478964794e-13 | 1.0202758999998878 |
| 791 | 2000 | 20260916.0 | restart_direction_variance | 2.3914126694025045e-13 | 1.0049973999998656 |
| 793 | 2000 | 20260917.0 | restart_direction_variance | 1.850604689345744e-12 | 1.0348675000000185 |
| 795 | 2000 | 20260918.0 | restart_direction_variance | 5.3878038306976196e-14 | 0.920255100000304 |
| 797 | 2000 | 20260919.0 | restart_direction_variance | 1.568481989198529e-13 | 1.1910577000003286 |
| 799 | 2000 | 20260920.0 | restart_direction_variance | 2.468956335972468e-13 | 0.9830117999999856 |
| 801 | 10000 | 20260916.0 | restart_direction_variance | 9.279494683521136e-14 | 0.9429312000002028 |
| 803 | 10000 | 20260917.0 | restart_direction_variance | 4.073176032824917e-14 | 0.9786874000001262 |
| 805 | 10000 | 20260918.0 | restart_direction_variance | 9.729260281579984e-14 | 0.8395958000000974 |
| 807 | 10000 | 20260919.0 | restart_direction_variance | 2.220326488191084e-13 | 1.0072248999999829 |
| 809 | 10000 | 20260920.0 | restart_direction_variance | 1.5358519320555148e-13 | 1.184292199999618 |

**dataset = equivalent_fit_multiple_parameters; benchmark = B1_known_truth; task = known_truth_parameter_and_coupling_recovery; split = synthetic; method = hard_ot; method_version = iot-benchmark-0.1.0**

| **Record** | **repeat** | **seed** | **metric** | **value** | **runtime_seconds** |
| --- | --- | --- | --- | --- | --- |
| 62 | 400 | 20260916.0 | heldout_coupling_relative_frobenius | 0.0875453932936106 | 0.0828166999999666 |
| 64 | 400 | 20260917.0 | heldout_coupling_relative_frobenius | 0.0605575656889887 | 0.1086900999998761 |
| 66 | 400 | 20260918.0 | heldout_coupling_relative_frobenius | 0.1311706133258216 | 0.0948812999999972 |
| 68 | 400 | 20260919.0 | heldout_coupling_relative_frobenius | 0.0705487777894709 | 0.1172025000000758 |
| 70 | 400 | 20260920.0 | heldout_coupling_relative_frobenius | 0.1192635665109429 | 0.1098799000001236 |
| 72 | 2000 | 20260916.0 | heldout_coupling_relative_frobenius | 0.0319856428657565 | 0.1029505999999855 |
| 74 | 2000 | 20260917.0 | heldout_coupling_relative_frobenius | 0.0424517505675177 | 0.1132460000003448 |
| 76 | 2000 | 20260918.0 | heldout_coupling_relative_frobenius | 0.0482162871811523 | 0.0879520999997112 |
| 78 | 2000 | 20260919.0 | heldout_coupling_relative_frobenius | 0.0410544587745597 | 0.122143799999776 |
| 80 | 2000 | 20260920.0 | heldout_coupling_relative_frobenius | 0.0355185889801211 | 0.0942646000003151 |
| 82 | 10000 | 20260916.0 | heldout_coupling_relative_frobenius | 0.0169125921828075 | 0.0865029000001413 |
| 84 | 10000 | 20260917.0 | heldout_coupling_relative_frobenius | 0.0147721964442347 | 0.1116302999998879 |
| 86 | 10000 | 20260918.0 | heldout_coupling_relative_frobenius | 0.0215679664373367 | 0.0872058999998444 |
| 88 | 10000 | 20260919.0 | heldout_coupling_relative_frobenius | 0.0208767249517476 | 0.1041626000001088 |
| 90 | 10000 | 20260920.0 | heldout_coupling_relative_frobenius | 0.0100125374149177 | 0.1108207000002039 |
| 242 | 400 | 20260916.0 | coefficient_cosine | -0.086935827326141 | 0.0828166999999666 |
| 244 | 400 | 20260917.0 | coefficient_cosine | 0.5015138814101904 | 0.1086900999998761 |
| 246 | 400 | 20260918.0 | coefficient_cosine | 0.9229497464064876 | 0.0948812999999972 |
| 248 | 400 | 20260919.0 | coefficient_cosine | 0.8935480796886097 | 0.1172025000000758 |
| 250 | 400 | 20260920.0 | coefficient_cosine | 0.5024514328571291 | 0.1098799000001236 |
| 252 | 2000 | 20260916.0 | coefficient_cosine | 0.4765973327281566 | 0.1029505999999855 |
| 254 | 2000 | 20260917.0 | coefficient_cosine | 0.0708728143413903 | 0.1132460000003448 |
| 256 | 2000 | 20260918.0 | coefficient_cosine | -0.3772857049065223 | 0.0879520999997112 |
| 258 | 2000 | 20260919.0 | coefficient_cosine | 0.0246986205077229 | 0.122143799999776 |
| 260 | 2000 | 20260920.0 | coefficient_cosine | -0.4434178673618426 | 0.0942646000003151 |
| 262 | 10000 | 20260916.0 | coefficient_cosine | 0.4775169754532443 | 0.0865029000001413 |
| 264 | 10000 | 20260917.0 | coefficient_cosine | 0.0709185158105209 | 0.1116302999998879 |
| 266 | 10000 | 20260918.0 | coefficient_cosine | -0.7660071869150487 | 0.0872058999998444 |
| 268 | 10000 | 20260919.0 | coefficient_cosine | -0.6109088938530205 | 0.1041626000001088 |
| 270 | 10000 | 20260920.0 | coefficient_cosine | -0.44357625490468 | 0.1108207000002039 |
| 422 | 400 | 20260916.0 | coefficient_sign_accuracy | 0.5 | 0.0828166999999666 |
| 424 | 400 | 20260917.0 | coefficient_sign_accuracy | 0.5 | 0.1086900999998761 |
| 426 | 400 | 20260918.0 | coefficient_sign_accuracy | 1.0 | 0.0948812999999972 |
| 428 | 400 | 20260919.0 | coefficient_sign_accuracy | 1.0 | 0.1172025000000758 |
| 430 | 400 | 20260920.0 | coefficient_sign_accuracy | 1.0 | 0.1098799000001236 |
| 432 | 2000 | 20260916.0 | coefficient_sign_accuracy | 0.75 | 0.1029505999999855 |
| 434 | 2000 | 20260917.0 | coefficient_sign_accuracy | 0.5 | 0.1132460000003448 |
| 436 | 2000 | 20260918.0 | coefficient_sign_accuracy | 0.25 | 0.0879520999997112 |
| 438 | 2000 | 20260919.0 | coefficient_sign_accuracy | 0.5 | 0.122143799999776 |
| 440 | 2000 | 20260920.0 | coefficient_sign_accuracy | 0.25 | 0.0942646000003151 |
| 442 | 10000 | 20260916.0 | coefficient_sign_accuracy | 0.75 | 0.0865029000001413 |
| 444 | 10000 | 20260917.0 | coefficient_sign_accuracy | 0.5 | 0.1116302999998879 |
| 446 | 10000 | 20260918.0 | coefficient_sign_accuracy | 0.25 | 0.0872058999998444 |
| 448 | 10000 | 20260919.0 | coefficient_sign_accuracy | 0.0 | 0.1041626000001088 |
| 450 | 10000 | 20260920.0 | coefficient_sign_accuracy | 0.25 | 0.1108207000002039 |
| 602 | 400 | 20260916.0 | pure_column_minimum_curvature | 0.0 | 0.0828166999999666 |
| 604 | 400 | 20260917.0 | pure_column_minimum_curvature | 0.0 | 0.1086900999998761 |
| 606 | 400 | 20260918.0 | pure_column_minimum_curvature | 0.0 | 0.0948812999999972 |
| 608 | 400 | 20260919.0 | pure_column_minimum_curvature | 0.0 | 0.1172025000000758 |
| 610 | 400 | 20260920.0 | pure_column_minimum_curvature | 0.0 | 0.1098799000001236 |
| 612 | 2000 | 20260916.0 | pure_column_minimum_curvature | 0.0 | 0.1029505999999855 |
| 614 | 2000 | 20260917.0 | pure_column_minimum_curvature | 0.0 | 0.1132460000003448 |
| 616 | 2000 | 20260918.0 | pure_column_minimum_curvature | 0.0 | 0.0879520999997112 |
| 618 | 2000 | 20260919.0 | pure_column_minimum_curvature | 0.0 | 0.122143799999776 |
| 620 | 2000 | 20260920.0 | pure_column_minimum_curvature | 0.0 | 0.0942646000003151 |
| 622 | 10000 | 20260916.0 | pure_column_minimum_curvature | -1.332267629550187e-09 | 0.0865029000001413 |
| 624 | 10000 | 20260917.0 | pure_column_minimum_curvature | 0.0 | 0.1116302999998879 |
| 626 | 10000 | 20260918.0 | pure_column_minimum_curvature | -8.88178419700125e-10 | 0.0872058999998444 |
| 628 | 10000 | 20260919.0 | pure_column_minimum_curvature | 0.0 | 0.1041626000001088 |
| 630 | 10000 | 20260920.0 | pure_column_minimum_curvature | 0.0 | 0.1108207000002039 |
| 782 | 400 | 20260916.0 | restart_direction_variance | 0.2061074051763044 | 0.0828166999999666 |
| 784 | 400 | 20260917.0 | restart_direction_variance | 0.2026443328940148 | 0.1086900999998761 |
| 786 | 400 | 20260918.0 | restart_direction_variance | 0.1939920563809734 | 0.0948812999999972 |
| 788 | 400 | 20260919.0 | restart_direction_variance | 0.1846038746057955 | 0.1172025000000758 |
| 790 | 400 | 20260920.0 | restart_direction_variance | 0.2056493387178563 | 0.1098799000001236 |
| 792 | 2000 | 20260916.0 | restart_direction_variance | 0.2116761136109295 | 0.1029505999999855 |
| 794 | 2000 | 20260917.0 | restart_direction_variance | 0.2045185254456162 | 0.1132460000003448 |
| 796 | 2000 | 20260918.0 | restart_direction_variance | 0.1952584819233156 | 0.0879520999997112 |
| 798 | 2000 | 20260919.0 | restart_direction_variance | 0.184731685252915 | 0.122143799999776 |
| 800 | 2000 | 20260920.0 | restart_direction_variance | 0.2054239093660288 | 0.0942646000003151 |
| 802 | 10000 | 20260916.0 | restart_direction_variance | 0.2125319125819989 | 0.0865029000001413 |
| 804 | 10000 | 20260917.0 | restart_direction_variance | 0.2047884180450052 | 0.1116302999998879 |
| 806 | 10000 | 20260918.0 | restart_direction_variance | 0.1953284558688947 | 0.0872058999998444 |
| 808 | 10000 | 20260919.0 | restart_direction_variance | 0.1847363409882331 | 0.1041626000001088 |
| 810 | 10000 | 20260920.0 | restart_direction_variance | 0.2054664590039889 | 0.1108207000002039 |

**dataset = shared_plus_context_offset; benchmark = B1_known_truth; task = known_truth_parameter_and_coupling_recovery; split = synthetic; method = soft_iot; method_version = iot-benchmark-0.1.0**

| **Record** | **repeat** | **seed** | **metric** | **value** | **runtime_seconds** |
| --- | --- | --- | --- | --- | --- |
| 91 | 400 | 20260916.0 | heldout_coupling_relative_frobenius | 0.176349899588655 | 1.1123382999999194 |
| 93 | 400 | 20260917.0 | heldout_coupling_relative_frobenius | 0.1329219102447165 | 1.1023595999999998 |
| 95 | 400 | 20260918.0 | heldout_coupling_relative_frobenius | 0.0637899611819555 | 1.051800800000365 |
| 97 | 400 | 20260919.0 | heldout_coupling_relative_frobenius | 0.144652293603667 | 1.0626346999997622 |
| 99 | 400 | 20260920.0 | heldout_coupling_relative_frobenius | 0.1813906615021428 | 1.0025394999997843 |
| 101 | 2000 | 20260916.0 | heldout_coupling_relative_frobenius | 0.1615041974620064 | 0.9982817000000068 |
| 103 | 2000 | 20260917.0 | heldout_coupling_relative_frobenius | 0.1418453657176784 | 1.02074599999969 |
| 105 | 2000 | 20260918.0 | heldout_coupling_relative_frobenius | 0.0527372985266395 | 0.8786196999999447 |
| 107 | 2000 | 20260919.0 | heldout_coupling_relative_frobenius | 0.1399247210575211 | 1.0219849000000067 |
| 109 | 2000 | 20260920.0 | heldout_coupling_relative_frobenius | 0.1553583689571544 | 0.9431463000000804 |
| 111 | 10000 | 20260916.0 | heldout_coupling_relative_frobenius | 0.1508604888432931 | 0.9935421999998651 |
| 113 | 10000 | 20260917.0 | heldout_coupling_relative_frobenius | 0.1186783539640445 | 0.963337700000011 |
| 115 | 10000 | 20260918.0 | heldout_coupling_relative_frobenius | 0.0585936962044973 | 0.9668118999998114 |
| 117 | 10000 | 20260919.0 | heldout_coupling_relative_frobenius | 0.1356045644629172 | 1.0804274999995869 |
| 119 | 10000 | 20260920.0 | heldout_coupling_relative_frobenius | 0.1676694214899196 | 1.0419104000002335 |
| 271 | 400 | 20260916.0 | coefficient_cosine | 0.983878363311385 | 1.1123382999999194 |
| 273 | 400 | 20260917.0 | coefficient_cosine | 0.96230632166459 | 1.1023595999999998 |
| 275 | 400 | 20260918.0 | coefficient_cosine | 0.9669640280409444 | 1.051800800000365 |
| 277 | 400 | 20260919.0 | coefficient_cosine | 0.9446925475013228 | 1.0626346999997622 |
| 279 | 400 | 20260920.0 | coefficient_cosine | 0.9948654215672548 | 1.0025394999997843 |
| 281 | 2000 | 20260916.0 | coefficient_cosine | 0.9947159533334876 | 0.9982817000000068 |
| 283 | 2000 | 20260917.0 | coefficient_cosine | 0.9940173153758532 | 1.02074599999969 |
| 285 | 2000 | 20260918.0 | coefficient_cosine | 0.9869180104974438 | 0.8786196999999447 |
| 287 | 2000 | 20260919.0 | coefficient_cosine | 0.991196724280534 | 1.0219849000000067 |
| 289 | 2000 | 20260920.0 | coefficient_cosine | 0.9956360081988342 | 0.9431463000000804 |
| 291 | 10000 | 20260916.0 | coefficient_cosine | 0.9886918705817698 | 0.9935421999998651 |
| 293 | 10000 | 20260917.0 | coefficient_cosine | 0.9912723253419724 | 0.963337700000011 |
| 295 | 10000 | 20260918.0 | coefficient_cosine | 0.9867369147137156 | 0.9668118999998114 |
| 297 | 10000 | 20260919.0 | coefficient_cosine | 0.9936065125452672 | 1.0804274999995869 |
| 299 | 10000 | 20260920.0 | coefficient_cosine | 0.9950878219702546 | 1.0419104000002335 |
| 451 | 400 | 20260916.0 | coefficient_sign_accuracy | 1.0 | 1.1123382999999194 |
| 453 | 400 | 20260917.0 | coefficient_sign_accuracy | 1.0 | 1.1023595999999998 |
| 455 | 400 | 20260918.0 | coefficient_sign_accuracy | 0.8333333333333334 | 1.051800800000365 |
| 457 | 400 | 20260919.0 | coefficient_sign_accuracy | 1.0 | 1.0626346999997622 |
| 459 | 400 | 20260920.0 | coefficient_sign_accuracy | 1.0 | 1.0025394999997843 |
| 461 | 2000 | 20260916.0 | coefficient_sign_accuracy | 1.0 | 0.9982817000000068 |
| 463 | 2000 | 20260917.0 | coefficient_sign_accuracy | 1.0 | 1.02074599999969 |
| 465 | 2000 | 20260918.0 | coefficient_sign_accuracy | 1.0 | 0.8786196999999447 |
| 467 | 2000 | 20260919.0 | coefficient_sign_accuracy | 1.0 | 1.0219849000000067 |
| 469 | 2000 | 20260920.0 | coefficient_sign_accuracy | 1.0 | 0.9431463000000804 |
| 471 | 10000 | 20260916.0 | coefficient_sign_accuracy | 1.0 | 0.9935421999998651 |
| 473 | 10000 | 20260917.0 | coefficient_sign_accuracy | 1.0 | 0.963337700000011 |
| 475 | 10000 | 20260918.0 | coefficient_sign_accuracy | 1.0 | 0.9668118999998114 |
| 477 | 10000 | 20260919.0 | coefficient_sign_accuracy | 1.0 | 1.0804274999995869 |
| 479 | 10000 | 20260920.0 | coefficient_sign_accuracy | 1.0 | 1.0419104000002335 |
| 631 | 400 | 20260916.0 | pure_column_minimum_curvature | 0.1806251819225224 | 1.1123382999999194 |
| 633 | 400 | 20260917.0 | pure_column_minimum_curvature | 0.2427997798370995 | 1.1023595999999998 |
| 635 | 400 | 20260918.0 | pure_column_minimum_curvature | 0.2876988536293368 | 1.051800800000365 |
| 637 | 400 | 20260919.0 | pure_column_minimum_curvature | 0.1217018432697614 | 1.0626346999997622 |
| 639 | 400 | 20260920.0 | pure_column_minimum_curvature | 0.2268075411926418 | 1.0025394999997843 |
| 641 | 2000 | 20260916.0 | pure_column_minimum_curvature | 0.2064917388189258 | 0.9982817000000068 |
| 643 | 2000 | 20260917.0 | pure_column_minimum_curvature | 0.2419019109467513 | 1.02074599999969 |
| 645 | 2000 | 20260918.0 | pure_column_minimum_curvature | 0.311526185292621 | 0.8786196999999447 |
| 647 | 2000 | 20260919.0 | pure_column_minimum_curvature | 0.0918527791604617 | 1.0219849000000067 |
| 649 | 2000 | 20260920.0 | pure_column_minimum_curvature | 0.2594209096429267 | 0.9431463000000804 |
| 651 | 10000 | 20260916.0 | pure_column_minimum_curvature | 0.206235566402313 | 0.9935421999998651 |
| 653 | 10000 | 20260917.0 | pure_column_minimum_curvature | 0.2205297464641376 | 0.963337700000011 |
| 655 | 10000 | 20260918.0 | pure_column_minimum_curvature | 0.3250866451587342 | 0.9668118999998114 |
| 657 | 10000 | 20260919.0 | pure_column_minimum_curvature | 0.1004778722446531 | 1.0804274999995869 |
| 659 | 10000 | 20260920.0 | pure_column_minimum_curvature | 0.2255028102027267 | 1.0419104000002335 |
| 811 | 400 | 20260916.0 | restart_direction_variance | 3.481958222410208e-14 | 1.1123382999999194 |
| 813 | 400 | 20260917.0 | restart_direction_variance | 1.1442553017775506e-13 | 1.1023595999999998 |
| 815 | 400 | 20260918.0 | restart_direction_variance | 5.044204415017759e-14 | 1.051800800000365 |
| 817 | 400 | 20260919.0 | restart_direction_variance | 5.150487815465089e-13 | 1.0626346999997622 |
| 819 | 400 | 20260920.0 | restart_direction_variance | 1.6987860407480638e-13 | 1.0025394999997843 |
| 821 | 2000 | 20260916.0 | restart_direction_variance | 2.605918452145289e-13 | 0.9982817000000068 |
| 823 | 2000 | 20260917.0 | restart_direction_variance | 1.3564681915947158e-13 | 1.02074599999969 |
| 825 | 2000 | 20260918.0 | restart_direction_variance | 2.3223849593205262e-14 | 0.8786196999999447 |
| 827 | 2000 | 20260919.0 | restart_direction_variance | 2.8976006908050733e-13 | 1.0219849000000067 |
| 829 | 2000 | 20260920.0 | restart_direction_variance | 1.488084099766755e-13 | 0.9431463000000804 |
| 831 | 10000 | 20260916.0 | restart_direction_variance | 4.9821214714759734e-14 | 0.9935421999998651 |
| 833 | 10000 | 20260917.0 | restart_direction_variance | 1.2006057206632446e-13 | 0.963337700000011 |
| 835 | 10000 | 20260918.0 | restart_direction_variance | 3.7721664267680624e-14 | 0.9668118999998114 |
| 837 | 10000 | 20260919.0 | restart_direction_variance | 3.6646187386754997e-13 | 1.0804274999995869 |
| 839 | 10000 | 20260920.0 | restart_direction_variance | 8.67179744625099e-14 | 1.0419104000002335 |

**dataset = shared_plus_context_offset; benchmark = B1_known_truth; task = known_truth_parameter_and_coupling_recovery; split = synthetic; method = hard_ot; method_version = iot-benchmark-0.1.0**

| **Record** | **repeat** | **seed** | **metric** | **value** | **runtime_seconds** |
| --- | --- | --- | --- | --- | --- |
| 92 | 400 | 20260916.0 | heldout_coupling_relative_frobenius | 0.0929479307921709 | 0.1654218999997283 |
| 94 | 400 | 20260917.0 | heldout_coupling_relative_frobenius | 0.1037580666377482 | 0.2124383000000307 |
| 96 | 400 | 20260918.0 | heldout_coupling_relative_frobenius | 0.0926607502841339 | 0.1991322000003492 |
| 98 | 400 | 20260919.0 | heldout_coupling_relative_frobenius | 0.0864115351641546 | 0.1177881999997225 |
| 100 | 400 | 20260920.0 | heldout_coupling_relative_frobenius | 0.0836533663402642 | 0.1008627999999589 |
| 102 | 2000 | 20260916.0 | heldout_coupling_relative_frobenius | 0.0224646084401439 | 0.1455492000000049 |
| 104 | 2000 | 20260917.0 | heldout_coupling_relative_frobenius | 0.0315953102262114 | 0.1568135000002257 |
| 106 | 2000 | 20260918.0 | heldout_coupling_relative_frobenius | 0.0630869160203272 | 0.1371281000001545 |
| 108 | 2000 | 20260919.0 | heldout_coupling_relative_frobenius | 0.0359209204658064 | 0.1076626999997643 |
| 110 | 2000 | 20260920.0 | heldout_coupling_relative_frobenius | 0.0398039337845239 | 0.085291600000346 |
| 112 | 10000 | 20260916.0 | heldout_coupling_relative_frobenius | 0.0118957364246071 | 0.1351744000003236 |
| 114 | 10000 | 20260917.0 | heldout_coupling_relative_frobenius | 0.0225459509500254 | 0.1563703000001624 |
| 116 | 10000 | 20260918.0 | heldout_coupling_relative_frobenius | 0.0174803305575769 | 0.1507541000000856 |
| 118 | 10000 | 20260919.0 | heldout_coupling_relative_frobenius | 0.0119123228528906 | 0.0913902999996025 |
| 120 | 10000 | 20260920.0 | heldout_coupling_relative_frobenius | 0.0146843564988675 | 0.0908484999999927 |
| 272 | 400 | 20260916.0 | coefficient_cosine | 0.774355873752146 | 0.1654218999997283 |
| 274 | 400 | 20260917.0 | coefficient_cosine | 0.6050969771079389 | 0.2124383000000307 |
| 276 | 400 | 20260918.0 | coefficient_cosine | 0.4530605726931598 | 0.1991322000003492 |
| 278 | 400 | 20260919.0 | coefficient_cosine | -0.3092309272785552 | 0.1177881999997225 |
| 280 | 400 | 20260920.0 | coefficient_cosine | 0.2557131357030778 | 0.1008627999999589 |
| 282 | 2000 | 20260916.0 | coefficient_cosine | 0.663379793073201 | 0.1455492000000049 |
| 284 | 2000 | 20260917.0 | coefficient_cosine | 0.0340950528890312 | 0.1568135000002257 |
| 286 | 2000 | 20260918.0 | coefficient_cosine | -0.0701477080561162 | 0.1371281000001545 |
| 288 | 2000 | 20260919.0 | coefficient_cosine | -0.3528420535419301 | 0.1076626999997643 |
| 290 | 2000 | 20260920.0 | coefficient_cosine | -0.0498502101016429 | 0.085291600000346 |
| 292 | 10000 | 20260916.0 | coefficient_cosine | 0.3143560047441472 | 0.1351744000003236 |
| 294 | 10000 | 20260917.0 | coefficient_cosine | 0.0229061740865776 | 0.1563703000001624 |
| 296 | 10000 | 20260918.0 | coefficient_cosine | 0.5671697964903925 | 0.1507541000000856 |
| 298 | 10000 | 20260919.0 | coefficient_cosine | 0.8854736399930558 | 0.0913902999996025 |
| 300 | 10000 | 20260920.0 | coefficient_cosine | 0.5708800265730167 | 0.0908484999999927 |
| 452 | 400 | 20260916.0 | coefficient_sign_accuracy | 0.8333333333333334 | 0.1654218999997283 |
| 454 | 400 | 20260917.0 | coefficient_sign_accuracy | 0.6666666666666666 | 0.2124383000000307 |
| 456 | 400 | 20260918.0 | coefficient_sign_accuracy | 0.5 | 0.1991322000003492 |
| 458 | 400 | 20260919.0 | coefficient_sign_accuracy | 0.3333333333333333 | 0.1177881999997225 |
| 460 | 400 | 20260920.0 | coefficient_sign_accuracy | 0.8333333333333334 | 0.1008627999999589 |
| 462 | 2000 | 20260916.0 | coefficient_sign_accuracy | 0.6666666666666666 | 0.1455492000000049 |
| 464 | 2000 | 20260917.0 | coefficient_sign_accuracy | 0.5 | 0.1568135000002257 |
| 466 | 2000 | 20260918.0 | coefficient_sign_accuracy | 0.5 | 0.1371281000001545 |
| 468 | 2000 | 20260919.0 | coefficient_sign_accuracy | 0.3333333333333333 | 0.1076626999997643 |
| 470 | 2000 | 20260920.0 | coefficient_sign_accuracy | 0.6666666666666666 | 0.085291600000346 |
| 472 | 10000 | 20260916.0 | coefficient_sign_accuracy | 0.6666666666666666 | 0.1351744000003236 |
| 474 | 10000 | 20260917.0 | coefficient_sign_accuracy | 0.5 | 0.1563703000001624 |
| 476 | 10000 | 20260918.0 | coefficient_sign_accuracy | 0.6666666666666666 | 0.1507541000000856 |
| 478 | 10000 | 20260919.0 | coefficient_sign_accuracy | 1.0 | 0.0913902999996025 |
| 480 | 10000 | 20260920.0 | coefficient_sign_accuracy | 1.0 | 0.0908484999999927 |
| 632 | 400 | 20260916.0 | pure_column_minimum_curvature | 0.0 | 0.1654218999997283 |
| 634 | 400 | 20260917.0 | pure_column_minimum_curvature | 0.0 | 0.2124383000000307 |
| 636 | 400 | 20260918.0 | pure_column_minimum_curvature | 0.0 | 0.1991322000003492 |
| 638 | 400 | 20260919.0 | pure_column_minimum_curvature | 0.0 | 0.1177881999997225 |
| 640 | 400 | 20260920.0 | pure_column_minimum_curvature | 0.0 | 0.1008627999999589 |
| 642 | 2000 | 20260916.0 | pure_column_minimum_curvature | 0.0 | 0.1455492000000049 |
| 644 | 2000 | 20260917.0 | pure_column_minimum_curvature | 0.0 | 0.1568135000002257 |
| 646 | 2000 | 20260918.0 | pure_column_minimum_curvature | 0.0 | 0.1371281000001545 |
| 648 | 2000 | 20260919.0 | pure_column_minimum_curvature | 0.0 | 0.1076626999997643 |
| 650 | 2000 | 20260920.0 | pure_column_minimum_curvature | 0.0 | 0.085291600000346 |
| 652 | 10000 | 20260916.0 | pure_column_minimum_curvature | 0.0 | 0.1351744000003236 |
| 654 | 10000 | 20260917.0 | pure_column_minimum_curvature | 0.0 | 0.1563703000001624 |
| 656 | 10000 | 20260918.0 | pure_column_minimum_curvature | 0.0 | 0.1507541000000856 |
| 658 | 10000 | 20260919.0 | pure_column_minimum_curvature | 4.4408920985006256e-10 | 0.0913902999996025 |
| 660 | 10000 | 20260920.0 | pure_column_minimum_curvature | 0.0 | 0.0908484999999927 |
| 812 | 400 | 20260916.0 | restart_direction_variance | 0.0594842194835263 | 0.1654218999997283 |
| 814 | 400 | 20260917.0 | restart_direction_variance | 0.1065244548845614 | 0.2124383000000307 |
| 816 | 400 | 20260918.0 | restart_direction_variance | 0.0788389175958989 | 0.1991322000003492 |
| 818 | 400 | 20260919.0 | restart_direction_variance | 0.1311391460060481 | 0.1177881999997225 |
| 820 | 400 | 20260920.0 | restart_direction_variance | 0.1527567481458919 | 0.1008627999999589 |
| 822 | 2000 | 20260916.0 | restart_direction_variance | 0.0602277907308195 | 0.1455492000000049 |
| 824 | 2000 | 20260917.0 | restart_direction_variance | 0.0976984095855669 | 0.1568135000002257 |
| 826 | 2000 | 20260918.0 | restart_direction_variance | 0.0790772311629539 | 0.1371281000001545 |
| 828 | 2000 | 20260919.0 | restart_direction_variance | 0.1433100589614536 | 0.1076626999997643 |
| 830 | 2000 | 20260920.0 | restart_direction_variance | 0.1316227060368269 | 0.085291600000346 |
| 832 | 10000 | 20260916.0 | restart_direction_variance | 0.0620757969709021 | 0.1351744000003236 |
| 834 | 10000 | 20260917.0 | restart_direction_variance | 0.1022596931349284 | 0.1563703000001624 |
| 836 | 10000 | 20260918.0 | restart_direction_variance | 0.0803319214467597 | 0.1507541000000856 |
| 838 | 10000 | 20260919.0 | restart_direction_variance | 0.1415485967961971 | 0.0913902999996025 |
| 840 | 10000 | 20260920.0 | restart_direction_variance | 0.1409021078976285 | 0.0908484999999927 |

**dataset = observation_noise_and_dropout; benchmark = B1_known_truth; task = known_truth_parameter_and_coupling_recovery; split = synthetic; method = soft_iot; method_version = iot-benchmark-0.1.0**

| **Record** | **repeat** | **seed** | **metric** | **value** | **runtime_seconds** |
| --- | --- | --- | --- | --- | --- |
| 121 | 400 | 20260916.0 | heldout_coupling_relative_frobenius | 0.0785321662065585 | 1.0358581000000413 |
| 123 | 400 | 20260917.0 | heldout_coupling_relative_frobenius | 0.0676958818428152 | 0.9919863999998596 |
| 125 | 400 | 20260918.0 | heldout_coupling_relative_frobenius | 0.062622277152564 | 1.0540828999996847 |
| 127 | 400 | 20260919.0 | heldout_coupling_relative_frobenius | 0.1202146354463767 | 1.068597899999986 |
| 129 | 400 | 20260920.0 | heldout_coupling_relative_frobenius | 0.0513226043632217 | 1.0296806000001195 |
| 131 | 2000 | 20260916.0 | heldout_coupling_relative_frobenius | 0.1489549607851971 | 1.0135543000001237 |
| 133 | 2000 | 20260917.0 | heldout_coupling_relative_frobenius | 0.0471815021424743 | 1.0352265000001353 |
| 135 | 2000 | 20260918.0 | heldout_coupling_relative_frobenius | 0.1433219112868007 | 1.039399499999945 |
| 137 | 2000 | 20260919.0 | heldout_coupling_relative_frobenius | 0.0758910386508216 | 0.999287699999968 |
| 139 | 2000 | 20260920.0 | heldout_coupling_relative_frobenius | 0.0576482462669196 | 1.0551024999999754 |
| 141 | 10000 | 20260916.0 | heldout_coupling_relative_frobenius | 0.0774264174877618 | 1.0065113999999085 |
| 143 | 10000 | 20260917.0 | heldout_coupling_relative_frobenius | 0.0818523159965736 | 1.1623053000002983 |
| 145 | 10000 | 20260918.0 | heldout_coupling_relative_frobenius | 0.0911218942030977 | 0.9902772000000368 |
| 147 | 10000 | 20260919.0 | heldout_coupling_relative_frobenius | 0.0672437305964824 | 1.043357399999877 |
| 149 | 10000 | 20260920.0 | heldout_coupling_relative_frobenius | 0.0593827681687093 | 1.035367099999803 |
| 301 | 400 | 20260916.0 | coefficient_cosine | 0.9967472535164869 | 1.0358581000000413 |
| 303 | 400 | 20260917.0 | coefficient_cosine | 0.9767322798458258 | 0.9919863999998596 |
| 305 | 400 | 20260918.0 | coefficient_cosine | 0.9951018292662792 | 1.0540828999996847 |
| 307 | 400 | 20260919.0 | coefficient_cosine | 0.9759492051878528 | 1.068597899999986 |
| 309 | 400 | 20260920.0 | coefficient_cosine | 0.9891256859744548 | 1.0296806000001195 |
| 311 | 2000 | 20260916.0 | coefficient_cosine | 0.9539626078210164 | 1.0135543000001237 |
| 313 | 2000 | 20260917.0 | coefficient_cosine | 0.9829699891816968 | 1.0352265000001353 |
| 315 | 2000 | 20260918.0 | coefficient_cosine | 0.9570087202260006 | 1.039399499999945 |
| 317 | 2000 | 20260919.0 | coefficient_cosine | 0.9819286242784604 | 0.999287699999968 |
| 319 | 2000 | 20260920.0 | coefficient_cosine | 0.982696784288408 | 1.0551024999999754 |
| 321 | 10000 | 20260916.0 | coefficient_cosine | 0.9481737487714266 | 1.0065113999999085 |
| 323 | 10000 | 20260917.0 | coefficient_cosine | 0.9855488641140872 | 1.1623053000002983 |
| 325 | 10000 | 20260918.0 | coefficient_cosine | 0.9904971490355312 | 0.9902772000000368 |
| 327 | 10000 | 20260919.0 | coefficient_cosine | 0.980333690010186 | 1.043357399999877 |
| 329 | 10000 | 20260920.0 | coefficient_cosine | 0.9776783571805892 | 1.035367099999803 |
| 481 | 400 | 20260916.0 | coefficient_sign_accuracy | 1.0 | 1.0358581000000413 |
| 483 | 400 | 20260917.0 | coefficient_sign_accuracy | 0.8333333333333334 | 0.9919863999998596 |
| 485 | 400 | 20260918.0 | coefficient_sign_accuracy | 0.8333333333333334 | 1.0540828999996847 |
| 487 | 400 | 20260919.0 | coefficient_sign_accuracy | 1.0 | 1.068597899999986 |
| 489 | 400 | 20260920.0 | coefficient_sign_accuracy | 0.8333333333333334 | 1.0296806000001195 |
| 491 | 2000 | 20260916.0 | coefficient_sign_accuracy | 1.0 | 1.0135543000001237 |
| 493 | 2000 | 20260917.0 | coefficient_sign_accuracy | 1.0 | 1.0352265000001353 |
| 495 | 2000 | 20260918.0 | coefficient_sign_accuracy | 0.8333333333333334 | 1.039399499999945 |
| 497 | 2000 | 20260919.0 | coefficient_sign_accuracy | 1.0 | 0.999287699999968 |
| 499 | 2000 | 20260920.0 | coefficient_sign_accuracy | 0.8333333333333334 | 1.0551024999999754 |
| 501 | 10000 | 20260916.0 | coefficient_sign_accuracy | 1.0 | 1.0065113999999085 |
| 503 | 10000 | 20260917.0 | coefficient_sign_accuracy | 1.0 | 1.1623053000002983 |
| 505 | 10000 | 20260918.0 | coefficient_sign_accuracy | 1.0 | 0.9902772000000368 |
| 507 | 10000 | 20260919.0 | coefficient_sign_accuracy | 1.0 | 1.043357399999877 |
| 509 | 10000 | 20260920.0 | coefficient_sign_accuracy | 1.0 | 1.035367099999803 |
| 661 | 400 | 20260916.0 | pure_column_minimum_curvature | 0.2300501353857953 | 1.0358581000000413 |
| 663 | 400 | 20260917.0 | pure_column_minimum_curvature | 0.2561734717687614 | 0.9919863999998596 |
| 665 | 400 | 20260918.0 | pure_column_minimum_curvature | 0.3082016704070156 | 1.0540828999996847 |
| 667 | 400 | 20260919.0 | pure_column_minimum_curvature | 0.1348554334867913 | 1.068597899999986 |
| 669 | 400 | 20260920.0 | pure_column_minimum_curvature | 0.2890735504301744 | 1.0296806000001195 |
| 671 | 2000 | 20260916.0 | pure_column_minimum_curvature | 0.2689885496565125 | 1.0135543000001237 |
| 673 | 2000 | 20260917.0 | pure_column_minimum_curvature | 0.2714303373174687 | 1.0352265000001353 |
| 675 | 2000 | 20260918.0 | pure_column_minimum_curvature | 0.3183389041261364 | 1.039399499999945 |
| 677 | 2000 | 20260919.0 | pure_column_minimum_curvature | 0.1017459156926747 | 0.999287699999968 |
| 679 | 2000 | 20260920.0 | pure_column_minimum_curvature | 0.2851953828653109 | 1.0551024999999754 |
| 681 | 10000 | 20260916.0 | pure_column_minimum_curvature | 0.2984883269441241 | 1.0065113999999085 |
| 683 | 10000 | 20260917.0 | pure_column_minimum_curvature | 0.2692835407991367 | 1.1623053000002983 |
| 685 | 10000 | 20260918.0 | pure_column_minimum_curvature | 0.3291365713309346 | 0.9902772000000368 |
| 687 | 10000 | 20260919.0 | pure_column_minimum_curvature | 0.118524962289257 | 1.043357399999877 |
| 689 | 10000 | 20260920.0 | pure_column_minimum_curvature | 0.2751476753104498 | 1.035367099999803 |
| 841 | 400 | 20260916.0 | restart_direction_variance | 5.490838664010235e-13 | 1.0358581000000413 |
| 843 | 400 | 20260917.0 | restart_direction_variance | 1.4711729933985094e-13 | 0.9919863999998596 |
| 845 | 400 | 20260918.0 | restart_direction_variance | 4.4670983196119657e-14 | 1.0540828999996847 |
| 847 | 400 | 20260919.0 | restart_direction_variance | 1.9162461935828538e-13 | 1.068597899999986 |
| 849 | 400 | 20260920.0 | restart_direction_variance | 9.213805659586929e-14 | 1.0296806000001195 |
| 851 | 2000 | 20260916.0 | restart_direction_variance | 2.6465583575920605e-13 | 1.0135543000001237 |
| 853 | 2000 | 20260917.0 | restart_direction_variance | 8.33415011356123e-13 | 1.0352265000001353 |
| 855 | 2000 | 20260918.0 | restart_direction_variance | 6.639936386425447e-14 | 1.039399499999945 |
| 857 | 2000 | 20260919.0 | restart_direction_variance | 1.281811857268472e-13 | 0.999287699999968 |
| 859 | 2000 | 20260920.0 | restart_direction_variance | 1.501260467186616e-13 | 1.0551024999999754 |
| 861 | 10000 | 20260916.0 | restart_direction_variance | 7.107029824757708e-13 | 1.0065113999999085 |
| 863 | 10000 | 20260917.0 | restart_direction_variance | 3.00659890596888e-13 | 1.1623053000002983 |
| 865 | 10000 | 20260918.0 | restart_direction_variance | 5.23780746450211e-14 | 0.9902772000000368 |
| 867 | 10000 | 20260919.0 | restart_direction_variance | 3.184327574699849e-13 | 1.043357399999877 |
| 869 | 10000 | 20260920.0 | restart_direction_variance | 2.206880739515217e-13 | 1.035367099999803 |

**dataset = observation_noise_and_dropout; benchmark = B1_known_truth; task = known_truth_parameter_and_coupling_recovery; split = synthetic; method = hard_ot; method_version = iot-benchmark-0.1.0**

| **Record** | **repeat** | **seed** | **metric** | **value** | **runtime_seconds** |
| --- | --- | --- | --- | --- | --- |
| 122 | 400 | 20260916.0 | heldout_coupling_relative_frobenius | 0.3385310251087775 | 0.1563836000000265 |
| 124 | 400 | 20260917.0 | heldout_coupling_relative_frobenius | 0.1987935033679807 | 0.133145599999807 |
| 126 | 400 | 20260918.0 | heldout_coupling_relative_frobenius | 0.1847956895034365 | 0.1892931000002136 |
| 128 | 400 | 20260919.0 | heldout_coupling_relative_frobenius | 0.2524986818699166 | 0.1380479999997987 |
| 130 | 400 | 20260920.0 | heldout_coupling_relative_frobenius | 0.1897818159811424 | 0.1098606999998992 |
| 132 | 2000 | 20260916.0 | heldout_coupling_relative_frobenius | 0.16915864639523 | 0.1671473999999762 |
| 134 | 2000 | 20260917.0 | heldout_coupling_relative_frobenius | 0.1653341247527186 | 0.1598726000001988 |
| 136 | 2000 | 20260918.0 | heldout_coupling_relative_frobenius | 0.2673372363766526 | 0.1720463000001473 |
| 138 | 2000 | 20260919.0 | heldout_coupling_relative_frobenius | 0.1199773030186652 | 0.1218624000002819 |
| 140 | 2000 | 20260920.0 | heldout_coupling_relative_frobenius | 0.1140306187711368 | 0.1008802999999716 |
| 142 | 10000 | 20260916.0 | heldout_coupling_relative_frobenius | 0.2087395408241687 | 0.1825636000003214 |
| 144 | 10000 | 20260917.0 | heldout_coupling_relative_frobenius | 0.3166819352147227 | 0.1776282999999239 |
| 146 | 10000 | 20260918.0 | heldout_coupling_relative_frobenius | 0.3205768007302343 | 0.1657964999999421 |
| 148 | 10000 | 20260919.0 | heldout_coupling_relative_frobenius | 0.1001306753922329 | 0.1143796999999722 |
| 150 | 10000 | 20260920.0 | heldout_coupling_relative_frobenius | 0.1462393856668217 | 0.1046404999997321 |
| 302 | 400 | 20260916.0 | coefficient_cosine | 0.788320671521866 | 0.1563836000000265 |
| 304 | 400 | 20260917.0 | coefficient_cosine | 0.3009814902726925 | 0.133145599999807 |
| 306 | 400 | 20260918.0 | coefficient_cosine | 0.5522973752933453 | 0.1892931000002136 |
| 308 | 400 | 20260919.0 | coefficient_cosine | 0.2125765308847584 | 0.1380479999997987 |
| 310 | 400 | 20260920.0 | coefficient_cosine | 0.2468557591401806 | 0.1098606999998992 |
| 312 | 2000 | 20260916.0 | coefficient_cosine | 0.2898920609117478 | 0.1671473999999762 |
| 314 | 2000 | 20260917.0 | coefficient_cosine | 0.0563999151773294 | 0.1598726000001988 |
| 316 | 2000 | 20260918.0 | coefficient_cosine | 0.919302640081237 | 0.1720463000001473 |
| 318 | 2000 | 20260919.0 | coefficient_cosine | 0.5695723813291148 | 0.1218624000002819 |
| 320 | 2000 | 20260920.0 | coefficient_cosine | -0.1019750431180877 | 0.1008802999999716 |
| 322 | 10000 | 20260916.0 | coefficient_cosine | 0.318531697740089 | 0.1825636000003214 |
| 324 | 10000 | 20260917.0 | coefficient_cosine | 0.4851799547327994 | 0.1776282999999239 |
| 326 | 10000 | 20260918.0 | coefficient_cosine | 0.5578115224001341 | 0.1657964999999421 |
| 328 | 10000 | 20260919.0 | coefficient_cosine | 0.5659673141968419 | 0.1143796999999722 |
| 330 | 10000 | 20260920.0 | coefficient_cosine | -0.0078934015798952 | 0.1046404999997321 |
| 482 | 400 | 20260916.0 | coefficient_sign_accuracy | 0.8333333333333334 | 0.1563836000000265 |
| 484 | 400 | 20260917.0 | coefficient_sign_accuracy | 0.6666666666666666 | 0.133145599999807 |
| 486 | 400 | 20260918.0 | coefficient_sign_accuracy | 0.6666666666666666 | 0.1892931000002136 |
| 488 | 400 | 20260919.0 | coefficient_sign_accuracy | 0.6666666666666666 | 0.1380479999997987 |
| 490 | 400 | 20260920.0 | coefficient_sign_accuracy | 0.8333333333333334 | 0.1098606999998992 |
| 492 | 2000 | 20260916.0 | coefficient_sign_accuracy | 0.6666666666666666 | 0.1671473999999762 |
| 494 | 2000 | 20260917.0 | coefficient_sign_accuracy | 0.5 | 0.1598726000001988 |
| 496 | 2000 | 20260918.0 | coefficient_sign_accuracy | 1.0 | 0.1720463000001473 |
| 498 | 2000 | 20260919.0 | coefficient_sign_accuracy | 0.8333333333333334 | 0.1218624000002819 |
| 500 | 2000 | 20260920.0 | coefficient_sign_accuracy | 0.6666666666666666 | 0.1008802999999716 |
| 502 | 10000 | 20260916.0 | coefficient_sign_accuracy | 0.6666666666666666 | 0.1825636000003214 |
| 504 | 10000 | 20260917.0 | coefficient_sign_accuracy | 0.6666666666666666 | 0.1776282999999239 |
| 506 | 10000 | 20260918.0 | coefficient_sign_accuracy | 0.6666666666666666 | 0.1657964999999421 |
| 508 | 10000 | 20260919.0 | coefficient_sign_accuracy | 0.8333333333333334 | 0.1143796999999722 |
| 510 | 10000 | 20260920.0 | coefficient_sign_accuracy | 0.6666666666666666 | 0.1046404999997321 |
| 662 | 400 | 20260916.0 | pure_column_minimum_curvature | 0.0 | 0.1563836000000265 |
| 664 | 400 | 20260917.0 | pure_column_minimum_curvature | 0.0 | 0.133145599999807 |
| 666 | 400 | 20260918.0 | pure_column_minimum_curvature | 0.0 | 0.1892931000002136 |
| 668 | 400 | 20260919.0 | pure_column_minimum_curvature | -8.88178419700125e-10 | 0.1380479999997987 |
| 670 | 400 | 20260920.0 | pure_column_minimum_curvature | 0.0 | 0.1098606999998992 |
| 672 | 2000 | 20260916.0 | pure_column_minimum_curvature | 0.0 | 0.1671473999999762 |
| 674 | 2000 | 20260917.0 | pure_column_minimum_curvature | 0.0 | 0.1598726000001988 |
| 676 | 2000 | 20260918.0 | pure_column_minimum_curvature | 0.0 | 0.1720463000001473 |
| 678 | 2000 | 20260919.0 | pure_column_minimum_curvature | 0.0 | 0.1218624000002819 |
| 680 | 2000 | 20260920.0 | pure_column_minimum_curvature | 0.0 | 0.1008802999999716 |
| 682 | 10000 | 20260916.0 | pure_column_minimum_curvature | 0.0 | 0.1825636000003214 |
| 684 | 10000 | 20260917.0 | pure_column_minimum_curvature | 0.0 | 0.1776282999999239 |
| 686 | 10000 | 20260918.0 | pure_column_minimum_curvature | 0.0 | 0.1657964999999421 |
| 688 | 10000 | 20260919.0 | pure_column_minimum_curvature | 0.0 | 0.1143796999999722 |
| 690 | 10000 | 20260920.0 | pure_column_minimum_curvature | 0.0 | 0.1046404999997321 |
| 842 | 400 | 20260916.0 | restart_direction_variance | 0.0482169221165358 | 0.1563836000000265 |
| 844 | 400 | 20260917.0 | restart_direction_variance | 0.1235159775896361 | 0.133145599999807 |
| 846 | 400 | 20260918.0 | restart_direction_variance | 0.0885887351607962 | 0.1892931000002136 |
| 848 | 400 | 20260919.0 | restart_direction_variance | 0.1159147915853171 | 0.1380479999997987 |
| 850 | 400 | 20260920.0 | restart_direction_variance | 0.1600869362558061 | 0.1098606999998992 |
| 852 | 2000 | 20260916.0 | restart_direction_variance | 0.0677332360700882 | 0.1671473999999762 |
| 854 | 2000 | 20260917.0 | restart_direction_variance | 0.0887472868150955 | 0.1598726000001988 |
| 856 | 2000 | 20260918.0 | restart_direction_variance | 0.0696930312573184 | 0.1720463000001473 |
| 858 | 2000 | 20260919.0 | restart_direction_variance | 0.1319561367184711 | 0.1218624000002819 |
| 860 | 2000 | 20260920.0 | restart_direction_variance | 0.1604264088921128 | 0.1008802999999716 |
| 862 | 10000 | 20260916.0 | restart_direction_variance | 0.0611113117739782 | 0.1825636000003214 |
| 864 | 10000 | 20260917.0 | restart_direction_variance | 0.0980965264671948 | 0.1776282999999239 |
| 866 | 10000 | 20260918.0 | restart_direction_variance | 0.0858742272328067 | 0.1657964999999421 |
| 868 | 10000 | 20260919.0 | restart_direction_variance | 0.1355386830150776 | 0.1143796999999722 |
| 870 | 10000 | 20260920.0 | restart_direction_variance | 0.109685851807493 | 0.1046404999997321 |

**dataset = nonlinear_or_hidden_confounding_misspecification; benchmark = B1_known_truth; task = known_truth_parameter_and_coupling_recovery; split = synthetic; method = soft_iot; method_version = iot-benchmark-0.1.0**

| **Record** | **repeat** | **seed** | **metric** | **value** | **runtime_seconds** |
| --- | --- | --- | --- | --- | --- |
| 151 | 400 | 20260916.0 | heldout_coupling_relative_frobenius | 0.5530488137550039 | 1.336510399999952 |
| 153 | 400 | 20260917.0 | heldout_coupling_relative_frobenius | 0.1875729995386292 | 1.113893200000348 |
| 155 | 400 | 20260918.0 | heldout_coupling_relative_frobenius | 0.4613756560568629 | 1.1119886000001316 |
| 157 | 400 | 20260919.0 | heldout_coupling_relative_frobenius | 0.2128855635957705 | 1.0350179000001845 |
| 159 | 400 | 20260920.0 | heldout_coupling_relative_frobenius | 0.200337403918906 | 1.0244719000002078 |
| 161 | 2000 | 20260916.0 | heldout_coupling_relative_frobenius | 0.491584882887873 | 1.476230199999918 |
| 163 | 2000 | 20260917.0 | heldout_coupling_relative_frobenius | 0.2407665401154692 | 1.1467106999998578 |
| 165 | 2000 | 20260918.0 | heldout_coupling_relative_frobenius | 0.5197733071256009 | 1.1300897999999506 |
| 167 | 2000 | 20260919.0 | heldout_coupling_relative_frobenius | 0.2077759461548372 | 1.08436329999995 |
| 169 | 2000 | 20260920.0 | heldout_coupling_relative_frobenius | 0.2562666787257949 | 0.9798384000000624 |
| 171 | 10000 | 20260916.0 | heldout_coupling_relative_frobenius | 0.5323834661893728 | 1.4587419000004047 |
| 173 | 10000 | 20260917.0 | heldout_coupling_relative_frobenius | 0.2017044555456117 | 1.1241569000003435 |
| 175 | 10000 | 20260918.0 | heldout_coupling_relative_frobenius | 0.5198508568221593 | 1.1670874999999796 |
| 177 | 10000 | 20260919.0 | heldout_coupling_relative_frobenius | 0.251756246015258 | 1.1234731999998075 |
| 179 | 10000 | 20260920.0 | heldout_coupling_relative_frobenius | 0.2534860217428559 | 1.0709382000000005 |
| 331 | 400 | 20260916.0 | coefficient_cosine | 0.8966584726140575 | 1.336510399999952 |
| 333 | 400 | 20260917.0 | coefficient_cosine | 0.8289730472803564 | 1.113893200000348 |
| 335 | 400 | 20260918.0 | coefficient_cosine | 0.9544044316458592 | 1.1119886000001316 |
| 337 | 400 | 20260919.0 | coefficient_cosine | 0.9554574627723031 | 1.0350179000001845 |
| 339 | 400 | 20260920.0 | coefficient_cosine | 0.9154458439019598 | 1.0244719000002078 |
| 341 | 2000 | 20260916.0 | coefficient_cosine | 0.902726064130205 | 1.476230199999918 |
| 343 | 2000 | 20260917.0 | coefficient_cosine | 0.860660210840053 | 1.1467106999998578 |
| 345 | 2000 | 20260918.0 | coefficient_cosine | 0.9585627830935328 | 1.1300897999999506 |
| 347 | 2000 | 20260919.0 | coefficient_cosine | 0.9604787581213314 | 1.08436329999995 |
| 349 | 2000 | 20260920.0 | coefficient_cosine | 0.9233885887604308 | 0.9798384000000624 |
| 351 | 10000 | 20260916.0 | coefficient_cosine | 0.8845401639712659 | 1.4587419000004047 |
| 353 | 10000 | 20260917.0 | coefficient_cosine | 0.860745098013487 | 1.1241569000003435 |
| 355 | 10000 | 20260918.0 | coefficient_cosine | 0.9670472557916912 | 1.1670874999999796 |
| 357 | 10000 | 20260919.0 | coefficient_cosine | 0.9631074454434106 | 1.1234731999998075 |
| 359 | 10000 | 20260920.0 | coefficient_cosine | 0.9334519539857776 | 1.0709382000000005 |
| 511 | 400 | 20260916.0 | coefficient_sign_accuracy | 0.8333333333333334 | 1.336510399999952 |
| 513 | 400 | 20260917.0 | coefficient_sign_accuracy | 0.6666666666666666 | 1.113893200000348 |
| 515 | 400 | 20260918.0 | coefficient_sign_accuracy | 0.8333333333333334 | 1.1119886000001316 |
| 517 | 400 | 20260919.0 | coefficient_sign_accuracy | 1.0 | 1.0350179000001845 |
| 519 | 400 | 20260920.0 | coefficient_sign_accuracy | 0.8333333333333334 | 1.0244719000002078 |
| 521 | 2000 | 20260916.0 | coefficient_sign_accuracy | 0.8333333333333334 | 1.476230199999918 |
| 523 | 2000 | 20260917.0 | coefficient_sign_accuracy | 0.6666666666666666 | 1.1467106999998578 |
| 525 | 2000 | 20260918.0 | coefficient_sign_accuracy | 0.8333333333333334 | 1.1300897999999506 |
| 527 | 2000 | 20260919.0 | coefficient_sign_accuracy | 1.0 | 1.08436329999995 |
| 529 | 2000 | 20260920.0 | coefficient_sign_accuracy | 0.8333333333333334 | 0.9798384000000624 |
| 531 | 10000 | 20260916.0 | coefficient_sign_accuracy | 0.8333333333333334 | 1.4587419000004047 |
| 533 | 10000 | 20260917.0 | coefficient_sign_accuracy | 0.6666666666666666 | 1.1241569000003435 |
| 535 | 10000 | 20260918.0 | coefficient_sign_accuracy | 0.8333333333333334 | 1.1670874999999796 |
| 537 | 10000 | 20260919.0 | coefficient_sign_accuracy | 1.0 | 1.1234731999998075 |
| 539 | 10000 | 20260920.0 | coefficient_sign_accuracy | 0.8333333333333334 | 1.0709382000000005 |
| 691 | 400 | 20260916.0 | pure_column_minimum_curvature | 0.2528508473709223 | 1.336510399999952 |
| 693 | 400 | 20260917.0 | pure_column_minimum_curvature | 0.2090664694698318 | 1.113893200000348 |
| 695 | 400 | 20260918.0 | pure_column_minimum_curvature | 0.3039915452696107 | 1.1119886000001316 |
| 697 | 400 | 20260919.0 | pure_column_minimum_curvature | 0.1596318677243857 | 1.0350179000001845 |
| 699 | 400 | 20260920.0 | pure_column_minimum_curvature | 0.2868084658658176 | 1.0244719000002078 |
| 701 | 2000 | 20260916.0 | pure_column_minimum_curvature | 0.2275319093136829 | 1.476230199999918 |
| 703 | 2000 | 20260917.0 | pure_column_minimum_curvature | 0.2040247149004415 | 1.1467106999998578 |
| 705 | 2000 | 20260918.0 | pure_column_minimum_curvature | 0.3354325874838082 | 1.1300897999999506 |
| 707 | 2000 | 20260919.0 | pure_column_minimum_curvature | 0.1409240706351511 | 1.08436329999995 |
| 709 | 2000 | 20260920.0 | pure_column_minimum_curvature | 0.26413883125187 | 0.9798384000000624 |
| 711 | 10000 | 20260916.0 | pure_column_minimum_curvature | 0.2489873067723635 | 1.4587419000004047 |
| 713 | 10000 | 20260917.0 | pure_column_minimum_curvature | 0.2118321718036497 | 1.1241569000003435 |
| 715 | 10000 | 20260918.0 | pure_column_minimum_curvature | 0.3107082746645062 | 1.1670874999999796 |
| 717 | 10000 | 20260919.0 | pure_column_minimum_curvature | 0.1446007931171777 | 1.1234731999998075 |
| 719 | 10000 | 20260920.0 | pure_column_minimum_curvature | 0.2859014975875595 | 1.0709382000000005 |
| 871 | 400 | 20260916.0 | restart_direction_variance | 9.139396645984977e-14 | 1.336510399999952 |
| 873 | 400 | 20260917.0 | restart_direction_variance | 3.510776888534104e-14 | 1.113893200000348 |
| 875 | 400 | 20260918.0 | restart_direction_variance | 4.5856379208771983e-14 | 1.1119886000001316 |
| 877 | 400 | 20260919.0 | restart_direction_variance | 4.001448349241048e-13 | 1.0350179000001845 |
| 879 | 400 | 20260920.0 | restart_direction_variance | 5.266317200382306e-14 | 1.0244719000002078 |
| 881 | 2000 | 20260916.0 | restart_direction_variance | 3.336933208238208e-13 | 1.476230199999918 |
| 883 | 2000 | 20260917.0 | restart_direction_variance | 1.0535411599425149e-13 | 1.1467106999998578 |
| 885 | 2000 | 20260918.0 | restart_direction_variance | 1.1634697621398428e-13 | 1.1300897999999506 |
| 887 | 2000 | 20260919.0 | restart_direction_variance | 5.617280852634681e-13 | 1.08436329999995 |
| 889 | 2000 | 20260920.0 | restart_direction_variance | 2.020776108889045e-14 | 0.9798384000000624 |
| 891 | 10000 | 20260916.0 | restart_direction_variance | 1.1549070369894005e-13 | 1.4587419000004047 |
| 893 | 10000 | 20260917.0 | restart_direction_variance | 1.7243642922941556e-14 | 1.1241569000003435 |
| 895 | 10000 | 20260918.0 | restart_direction_variance | 1.252943046735326e-14 | 1.1670874999999796 |
| 897 | 10000 | 20260919.0 | restart_direction_variance | 5.898265903820428e-14 | 1.1234731999998075 |
| 899 | 10000 | 20260920.0 | restart_direction_variance | 1.0130538200248088e-13 | 1.0709382000000005 |

**dataset = nonlinear_or_hidden_confounding_misspecification; benchmark = B1_known_truth; task = known_truth_parameter_and_coupling_recovery; split = synthetic; method = hard_ot; method_version = iot-benchmark-0.1.0**

| **Record** | **repeat** | **seed** | **metric** | **value** | **runtime_seconds** |
| --- | --- | --- | --- | --- | --- |
| 152 | 400 | 20260916.0 | heldout_coupling_relative_frobenius | 0.555298431766314 | 0.2024116000002322 |
| 154 | 400 | 20260917.0 | heldout_coupling_relative_frobenius | 0.1951288909076906 | 0.2007628999999724 |
| 156 | 400 | 20260918.0 | heldout_coupling_relative_frobenius | 0.447226059780135 | 0.2208121999997274 |
| 158 | 400 | 20260919.0 | heldout_coupling_relative_frobenius | 0.2079947501315176 | 0.12569749999966 |
| 160 | 400 | 20260920.0 | heldout_coupling_relative_frobenius | 0.1883585556446613 | 0.1201066999997237 |
| 162 | 2000 | 20260916.0 | heldout_coupling_relative_frobenius | 0.5091021717136246 | 0.2441287999999986 |
| 164 | 2000 | 20260917.0 | heldout_coupling_relative_frobenius | 0.2359576598099584 | 0.2061431000001903 |
| 166 | 2000 | 20260918.0 | heldout_coupling_relative_frobenius | 0.5163193138019204 | 0.1995476000001872 |
| 168 | 2000 | 20260919.0 | heldout_coupling_relative_frobenius | 0.1969283934338379 | 0.1443789999998443 |
| 170 | 2000 | 20260920.0 | heldout_coupling_relative_frobenius | 0.2530387104095217 | 0.1209948999999142 |
| 172 | 10000 | 20260916.0 | heldout_coupling_relative_frobenius | 0.534234276688569 | 0.2454908999998224 |
| 174 | 10000 | 20260917.0 | heldout_coupling_relative_frobenius | 0.1979790952318598 | 0.204494699999941 |
| 176 | 10000 | 20260918.0 | heldout_coupling_relative_frobenius | 0.4987486497025511 | 0.2254143000000112 |
| 178 | 10000 | 20260919.0 | heldout_coupling_relative_frobenius | 0.2379669061125247 | 0.1464865000002646 |
| 180 | 10000 | 20260920.0 | heldout_coupling_relative_frobenius | 0.2462986375210409 | 0.1322454999999536 |
| 332 | 400 | 20260916.0 | coefficient_cosine | 0.6720125262978333 | 0.2024116000002322 |
| 334 | 400 | 20260917.0 | coefficient_cosine | 0.5798326502004336 | 0.2007628999999724 |
| 336 | 400 | 20260918.0 | coefficient_cosine | 0.0170810173016286 | 0.2208121999997274 |
| 338 | 400 | 20260919.0 | coefficient_cosine | 0.5699760842478638 | 0.12569749999966 |
| 340 | 400 | 20260920.0 | coefficient_cosine | 0.3087393497014509 | 0.1201066999997237 |
| 342 | 2000 | 20260916.0 | coefficient_cosine | 0.7787957164230688 | 0.2441287999999986 |
| 344 | 2000 | 20260917.0 | coefficient_cosine | 0.3136889313382608 | 0.2061431000001903 |
| 346 | 2000 | 20260918.0 | coefficient_cosine | 0.9015184231185902 | 0.1995476000001872 |
| 348 | 2000 | 20260919.0 | coefficient_cosine | -0.2823761639212537 | 0.1443789999998443 |
| 350 | 2000 | 20260920.0 | coefficient_cosine | 0.5562364773440662 | 0.1209948999999142 |
| 352 | 10000 | 20260916.0 | coefficient_cosine | 0.7761515004235421 | 0.2454908999998224 |
| 354 | 10000 | 20260917.0 | coefficient_cosine | 0.4691785073556372 | 0.204494699999941 |
| 356 | 10000 | 20260918.0 | coefficient_cosine | 0.4742066411384317 | 0.2254143000000112 |
| 358 | 10000 | 20260919.0 | coefficient_cosine | 0.5707651963493476 | 0.1464865000002646 |
| 360 | 10000 | 20260920.0 | coefficient_cosine | 0.1617063351267477 | 0.1322454999999536 |
| 512 | 400 | 20260916.0 | coefficient_sign_accuracy | 0.6666666666666666 | 0.2024116000002322 |
| 514 | 400 | 20260917.0 | coefficient_sign_accuracy | 0.5 | 0.2007628999999724 |
| 516 | 400 | 20260918.0 | coefficient_sign_accuracy | 0.5 | 0.2208121999997274 |
| 518 | 400 | 20260919.0 | coefficient_sign_accuracy | 0.8333333333333334 | 0.12569749999966 |
| 520 | 400 | 20260920.0 | coefficient_sign_accuracy | 0.8333333333333334 | 0.1201066999997237 |
| 522 | 2000 | 20260916.0 | coefficient_sign_accuracy | 0.8333333333333334 | 0.2441287999999986 |
| 524 | 2000 | 20260917.0 | coefficient_sign_accuracy | 0.5 | 0.2061431000001903 |
| 526 | 2000 | 20260918.0 | coefficient_sign_accuracy | 1.0 | 0.1995476000001872 |
| 528 | 2000 | 20260919.0 | coefficient_sign_accuracy | 0.3333333333333333 | 0.1443789999998443 |
| 530 | 2000 | 20260920.0 | coefficient_sign_accuracy | 1.0 | 0.1209948999999142 |
| 532 | 10000 | 20260916.0 | coefficient_sign_accuracy | 0.8333333333333334 | 0.2454908999998224 |
| 534 | 10000 | 20260917.0 | coefficient_sign_accuracy | 0.5 | 0.204494699999941 |
| 536 | 10000 | 20260918.0 | coefficient_sign_accuracy | 0.5 | 0.2254143000000112 |
| 538 | 10000 | 20260919.0 | coefficient_sign_accuracy | 0.8333333333333334 | 0.1464865000002646 |
| 540 | 10000 | 20260920.0 | coefficient_sign_accuracy | 0.5 | 0.1322454999999536 |
| 692 | 400 | 20260916.0 | pure_column_minimum_curvature | 4.4408920985006256e-10 | 0.2024116000002322 |
| 694 | 400 | 20260917.0 | pure_column_minimum_curvature | 0.0 | 0.2007628999999724 |
| 696 | 400 | 20260918.0 | pure_column_minimum_curvature | -4.4408920985006256e-10 | 0.2208121999997274 |
| 698 | 400 | 20260919.0 | pure_column_minimum_curvature | 0.0 | 0.12569749999966 |
| 700 | 400 | 20260920.0 | pure_column_minimum_curvature | -8.88178419700125e-10 | 0.1201066999997237 |
| 702 | 2000 | 20260916.0 | pure_column_minimum_curvature | 4.4408920985006256e-10 | 0.2441287999999986 |
| 704 | 2000 | 20260917.0 | pure_column_minimum_curvature | 0.0 | 0.2061431000001903 |
| 706 | 2000 | 20260918.0 | pure_column_minimum_curvature | 0.0 | 0.1995476000001872 |
| 708 | 2000 | 20260919.0 | pure_column_minimum_curvature | 0.0 | 0.1443789999998443 |
| 710 | 2000 | 20260920.0 | pure_column_minimum_curvature | 0.0 | 0.1209948999999142 |
| 712 | 10000 | 20260916.0 | pure_column_minimum_curvature | -8.88178419700125e-10 | 0.2454908999998224 |
| 714 | 10000 | 20260917.0 | pure_column_minimum_curvature | 0.0 | 0.204494699999941 |
| 716 | 10000 | 20260918.0 | pure_column_minimum_curvature | 0.0 | 0.2254143000000112 |
| 718 | 10000 | 20260919.0 | pure_column_minimum_curvature | 0.0 | 0.1464865000002646 |
| 720 | 10000 | 20260920.0 | pure_column_minimum_curvature | -4.4408920985006256e-10 | 0.1322454999999536 |
| 872 | 400 | 20260916.0 | restart_direction_variance | 0.0504520767972573 | 0.2024116000002322 |
| 874 | 400 | 20260917.0 | restart_direction_variance | 0.0879365062436688 | 0.2007628999999724 |
| 876 | 400 | 20260918.0 | restart_direction_variance | 0.0609491429114784 | 0.2208121999997274 |
| 878 | 400 | 20260919.0 | restart_direction_variance | 0.1197389700415968 | 0.12569749999966 |
| 880 | 400 | 20260920.0 | restart_direction_variance | 0.0991152344944836 | 0.1201066999997237 |
| 882 | 2000 | 20260916.0 | restart_direction_variance | 0.0520624070412131 | 0.2441287999999986 |
| 884 | 2000 | 20260917.0 | restart_direction_variance | 0.0980267593513495 | 0.2061431000001903 |
| 886 | 2000 | 20260918.0 | restart_direction_variance | 0.0600716149570704 | 0.1995476000001872 |
| 888 | 2000 | 20260919.0 | restart_direction_variance | 0.1235593532246011 | 0.1443789999998443 |
| 890 | 2000 | 20260920.0 | restart_direction_variance | 0.0953128164306985 | 0.1209948999999142 |
| 892 | 10000 | 20260916.0 | restart_direction_variance | 0.0532632793768051 | 0.2454908999998224 |
| 894 | 10000 | 20260917.0 | restart_direction_variance | 0.09175277978845 | 0.204494699999941 |
| 896 | 10000 | 20260918.0 | restart_direction_variance | 0.0637191027825734 | 0.2254143000000112 |
| 898 | 10000 | 20260919.0 | restart_direction_variance | 0.1225961669094714 | 0.1464865000002646 |
| 900 | 10000 | 20260920.0 | restart_direction_variance | 0.0873372969602336 | 0.1322454999999536 |

**dataset = gse239651_transition_panel; benchmark = B2_lineage_transition; task = transition_recovery; split = retrospective_reconstruction; method = UOT-IOT; method_version = iot-benchmark-0.1.0**

| **Record** | **repeat** | **seed** | **metric** | **value** | **runtime_seconds** |
| --- | --- | --- | --- | --- | --- |
| 901 |  | 20260916.0 | coupling_relative_frobenius | 0.9981903108580128 | 1.2096544999999423 |
| 902 |  | 20260917.0 | coupling_relative_frobenius | 0.998190310911833 | 1.1872864000001755 |
| 903 |  | 20260918.0 | coupling_relative_frobenius | 0.9981903108895868 | 1.2096050999998624 |
| 904 |  | 20260919.0 | coupling_relative_frobenius | 0.998190310870727 | 1.2151920999999677 |
| 905 |  | 20260920.0 | coupling_relative_frobenius | 0.9981903108864354 | 1.195388499999808 |
| 921 |  | 20260916.0 | transition_weighted_l1 | 0.0951153419028303 | 1.2096544999999423 |
| 922 |  | 20260917.0 | transition_weighted_l1 | 0.0951156677659977 | 1.1872864000001755 |
| 923 |  | 20260918.0 | transition_weighted_l1 | 0.0951155151281439 | 1.2096050999998624 |
| 924 |  | 20260919.0 | transition_weighted_l1 | 0.0951153868050402 | 1.2151920999999677 |
| 925 |  | 20260920.0 | transition_weighted_l1 | 0.0951155082735286 | 1.195388499999808 |
| 941 |  | 20260916.0 | fate_pearson | 0.4787361542456028 | 1.2096544999999423 |
| 942 |  | 20260917.0 | fate_pearson | 0.4787360817298395 | 1.1872864000001755 |
| 943 |  | 20260918.0 | fate_pearson | 0.478736115647362 | 1.2096050999998624 |
| 944 |  | 20260919.0 | fate_pearson | 0.4787361441121345 | 1.2151920999999677 |
| 945 |  | 20260920.0 | fate_pearson | 0.4787361171020095 | 1.195388499999808 |
| 961 |  | 20260916.0 | fate_spearman | 0.5112475264202239 | 1.2096544999999423 |
| 962 |  | 20260917.0 | fate_spearman | 0.5108023198928917 | 1.1872864000001755 |
| 963 |  | 20260918.0 | fate_spearman | 0.5111729914765657 | 1.2096050999998624 |
| 964 |  | 20260919.0 | fate_spearman | 0.5113212497037449 | 1.2151920999999677 |
| 965 |  | 20260920.0 | fate_spearman | 0.512172803610128 | 1.195388499999808 |
| 981 |  | 20260916.0 | fate_top1_accuracy | 0.3962703962703963 | 1.2096544999999423 |
| 982 |  | 20260917.0 | fate_top1_accuracy | 0.3962703962703963 | 1.1872864000001755 |
| 983 |  | 20260918.0 | fate_top1_accuracy | 0.3962703962703963 | 1.2096050999998624 |
| 984 |  | 20260919.0 | fate_top1_accuracy | 0.3962703962703963 | 1.2151920999999677 |
| 985 |  | 20260920.0 | fate_top1_accuracy | 0.3962703962703963 | 1.195388499999808 |

**dataset = macsgestalt_transition_panel; benchmark = B2_lineage_transition; task = transition_recovery; split = retrospective_reconstruction; method = UOT-IOT; method_version = iot-benchmark-0.1.0**

| **Record** | **repeat** | **seed** | **metric** | **value** | **runtime_seconds** |
| --- | --- | --- | --- | --- | --- |
| 906 |  | 20260916.0 | coupling_relative_frobenius | 0.8021728510836154 | 0.3401880999999775 |
| 907 |  | 20260917.0 | coupling_relative_frobenius | 0.8021728506771723 | 0.3242322999999487 |
| 908 |  | 20260918.0 | coupling_relative_frobenius | 0.8021728491033728 | 0.366518399999677 |
| 909 |  | 20260919.0 | coupling_relative_frobenius | 0.8021728499429494 | 0.3456639999999424 |
| 910 |  | 20260920.0 | coupling_relative_frobenius | 0.8021728485972119 | 0.3324896999997691 |
| 926 |  | 20260916.0 | transition_weighted_l1 | 0.3322085350486563 | 0.3401880999999775 |
| 927 |  | 20260917.0 | transition_weighted_l1 | 0.3322085362872843 | 0.3242322999999487 |
| 928 |  | 20260918.0 | transition_weighted_l1 | 0.3322085367587918 | 0.366518399999677 |
| 929 |  | 20260919.0 | transition_weighted_l1 | 0.3322085380403349 | 0.3456639999999424 |
| 930 |  | 20260920.0 | transition_weighted_l1 | 0.3322085363145715 | 0.3324896999997691 |
| 946 |  | 20260916.0 | fate_pearson | 0.4037133931804976 | 0.3401880999999775 |
| 947 |  | 20260917.0 | fate_pearson | 0.4037134004495707 | 0.3242322999999487 |
| 948 |  | 20260918.0 | fate_pearson | 0.4037134011405447 | 0.366518399999677 |
| 949 |  | 20260919.0 | fate_pearson | 0.4037134092397098 | 0.3456639999999424 |
| 950 |  | 20260920.0 | fate_pearson | 0.4037134023415135 | 0.3324896999997691 |
| 966 |  | 20260916.0 | fate_spearman | 0.409393221225753 | 0.3401880999999775 |
| 967 |  | 20260917.0 | fate_spearman | 0.4093784245381994 | 0.3242322999999487 |
| 968 |  | 20260918.0 | fate_spearman | 0.4090126247617452 | 0.366518399999677 |
| 969 |  | 20260919.0 | fate_spearman | 0.4085913456423439 | 0.3456639999999424 |
| 970 |  | 20260920.0 | fate_spearman | 0.4087984790783026 | 0.3324896999997691 |
| 986 |  | 20260916.0 | fate_top1_accuracy | 0.5476190476190477 | 0.3401880999999775 |
| 987 |  | 20260917.0 | fate_top1_accuracy | 0.5476190476190477 | 0.3242322999999487 |
| 988 |  | 20260918.0 | fate_top1_accuracy | 0.5476190476190477 | 0.366518399999677 |
| 989 |  | 20260919.0 | fate_top1_accuracy | 0.5476190476190477 | 0.3456639999999424 |
| 990 |  | 20260920.0 | fate_top1_accuracy | 0.5476190476190477 | 0.3324896999997691 |

**dataset = gse140802_t2_t16; benchmark = B2_lineage_transition; task = transition_recovery; split = retrospective_reconstruction; method = UOT-IOT; method_version = iot-benchmark-0.1.0**

| **Record** | **repeat** | **seed** | **metric** | **value** | **runtime_seconds** |
| --- | --- | --- | --- | --- | --- |
| 911 |  | 20260916.0 | coupling_relative_frobenius | 0.9874996653580814 | 0.9998890000001666 |
| 912 |  | 20260917.0 | coupling_relative_frobenius | 0.9874996653580814 | 0.9571274999998424 |
| 913 |  | 20260918.0 | coupling_relative_frobenius | 0.9874996653580814 | 0.8777365000000827 |
| 914 |  | 20260919.0 | coupling_relative_frobenius | 0.9874996653580814 | 0.9703205999999228 |
| 915 |  | 20260920.0 | coupling_relative_frobenius | 0.9874996653580814 | 0.8830874999998741 |
| 931 |  | 20260916.0 | transition_weighted_l1 | 0.0107461842228854 | 0.9998890000001666 |
| 932 |  | 20260917.0 | transition_weighted_l1 | 0.0107461842228854 | 0.9571274999998424 |
| 933 |  | 20260918.0 | transition_weighted_l1 | 0.0107461842228854 | 0.8777365000000827 |
| 934 |  | 20260919.0 | transition_weighted_l1 | 0.0107461842228854 | 0.9703205999999228 |
| 935 |  | 20260920.0 | transition_weighted_l1 | 0.0107461842228854 | 0.8830874999998741 |
| 951 |  | 20260916.0 | fate_pearson | 0.6855422778990485 | 0.9998890000001666 |
| 952 |  | 20260917.0 | fate_pearson | 0.6855422778990484 | 0.9571274999998424 |
| 953 |  | 20260918.0 | fate_pearson | 0.6855422778990484 | 0.8777365000000827 |
| 954 |  | 20260919.0 | fate_pearson | 0.6855422778990485 | 0.9703205999999228 |
| 955 |  | 20260920.0 | fate_pearson | 0.6855422778990484 | 0.8830874999998741 |
| 971 |  | 20260916.0 | fate_spearman | 0.616007979755324 | 0.9998890000001666 |
| 972 |  | 20260917.0 | fate_spearman | 0.6110366997961014 | 0.9571274999998424 |
| 973 |  | 20260918.0 | fate_spearman | 0.6080058943575364 | 0.8777365000000827 |
| 974 |  | 20260919.0 | fate_spearman | 0.6169133579581954 | 0.9703205999999228 |
| 975 |  | 20260920.0 | fate_spearman | 0.6096720439090709 | 0.8830874999998741 |
| 991 |  | 20260916.0 | fate_top1_accuracy | 0.55 | 0.9998890000001666 |
| 992 |  | 20260917.0 | fate_top1_accuracy | 0.55 | 0.9571274999998424 |
| 993 |  | 20260918.0 | fate_top1_accuracy | 0.55 | 0.8777365000000827 |
| 994 |  | 20260919.0 | fate_top1_accuracy | 0.55 | 0.9703205999999228 |
| 995 |  | 20260920.0 | fate_top1_accuracy | 0.55 | 0.8830874999998741 |

**dataset = gse140802_t2_t9; benchmark = B2_lineage_transition; task = transition_recovery; split = retrospective_reconstruction; method = UOT-IOT; method_version = iot-benchmark-0.1.0**

| **Record** | **repeat** | **seed** | **metric** | **value** | **runtime_seconds** |
| --- | --- | --- | --- | --- | --- |
| 916 |  | 20260916.0 | coupling_relative_frobenius | 0.995808566288555 | 1.4134618000002774 |
| 917 |  | 20260917.0 | coupling_relative_frobenius | 0.995808566288555 | 1.3819898000001558 |
| 918 |  | 20260918.0 | coupling_relative_frobenius | 0.995808566288555 | 1.3514521000001878 |
| 919 |  | 20260919.0 | coupling_relative_frobenius | 0.995808566288555 | 1.4133861000000252 |
| 920 |  | 20260920.0 | coupling_relative_frobenius | 0.995808566288555 | 1.3885298000000148 |
| 936 |  | 20260916.0 | transition_weighted_l1 | 0.0150171352657198 | 1.4134618000002774 |
| 937 |  | 20260917.0 | transition_weighted_l1 | 0.0150171352657198 | 1.3819898000001558 |
| 938 |  | 20260918.0 | transition_weighted_l1 | 0.0150171352657198 | 1.3514521000001878 |
| 939 |  | 20260919.0 | transition_weighted_l1 | 0.0150171352657198 | 1.4133861000000252 |
| 940 |  | 20260920.0 | transition_weighted_l1 | 0.0150171352657198 | 1.3885298000000148 |
| 956 |  | 20260916.0 | fate_pearson | 0.5449431551874835 | 1.4134618000002774 |
| 957 |  | 20260917.0 | fate_pearson | 0.5449431551874835 | 1.3819898000001558 |
| 958 |  | 20260918.0 | fate_pearson | 0.5449431551874835 | 1.3514521000001878 |
| 959 |  | 20260919.0 | fate_pearson | 0.5449431551874835 | 1.4133861000000252 |
| 960 |  | 20260920.0 | fate_pearson | 0.5449431551874835 | 1.3885298000000148 |
| 976 |  | 20260916.0 | fate_spearman | 0.5359660819469549 | 1.4134618000002774 |
| 977 |  | 20260917.0 | fate_spearman | 0.534325798976456 | 1.3819898000001558 |
| 978 |  | 20260918.0 | fate_spearman | 0.5358707695380804 | 1.3514521000001878 |
| 979 |  | 20260919.0 | fate_spearman | 0.53586267451377 | 1.4133861000000252 |
| 980 |  | 20260920.0 | fate_spearman | 0.5366456855786991 | 1.3885298000000148 |
| 996 |  | 20260916.0 | fate_top1_accuracy | 0.5526315789473685 | 1.4134618000002774 |
| 997 |  | 20260917.0 | fate_top1_accuracy | 0.5526315789473685 | 1.3819898000001558 |
| 998 |  | 20260918.0 | fate_top1_accuracy | 0.5526315789473685 | 1.3514521000001878 |
| 999 |  | 20260919.0 | fate_top1_accuracy | 0.5526315789473685 | 1.4133861000000252 |
| 1000 |  | 20260920.0 | fate_top1_accuracy | 0.5526315789473685 | 1.3885298000000148 |

**dataset = gse239651_transition_panel; benchmark = B2_lineage_transition; task = transition_recovery; split = retrospective_reconstruction; method = wot; method_version = see_method_status**

| **Record** | **repeat** | **seed** | **metric** | **value** | **runtime_seconds** |
| --- | --- | --- | --- | --- | --- |
| 1001 |  |  | coupling_relative_frobenius | 1.0054741329829409 | 3.874419000000216 |
| 1002 |  |  | transition_weighted_l1 | 0.5622235992510631 | 3.874419000000216 |
| 1003 |  |  | fate_pearson | 0.2829156431995213 | 3.874419000000216 |
| 1004 |  |  | fate_spearman | 0.38966389484292985 | 3.874419000000216 |
| 1005 |  |  | fate_top1_accuracy | 0.2727272727272727 | 3.874419000000216 |

**dataset = gse239651_transition_panel; benchmark = B2_lineage_transition; task = transition_recovery; split = retrospective_reconstruction; method = moscot; method_version = see_method_status**

| **Record** | **repeat** | **seed** | **metric** | **value** | **runtime_seconds** |
| --- | --- | --- | --- | --- | --- |
| 1006 |  |  | coupling_relative_frobenius | 1.0022205766976557 | 10.35936730000003 |
| 1007 |  |  | transition_weighted_l1 | 0.5249415568089765 | 10.35936730000003 |
| 1008 |  |  | fate_pearson | 0.291872157616353 | 10.35936730000003 |
| 1009 |  |  | fate_spearman | 0.38784344981661156 | 10.35936730000003 |
| 1010 |  |  | fate_top1_accuracy | 0.28205128205128205 | 10.35936730000003 |

**dataset = gse239651_transition_panel; benchmark = B2_lineage_transition; task = transition_recovery; split = retrospective_reconstruction; method = lineageot; method_version = see_method_status**

| **Record** | **repeat** | **seed** | **metric** | **value** | **runtime_seconds** |
| --- | --- | --- | --- | --- | --- |
| 1011 |  |  | coupling_relative_frobenius | 1.0026475896743297 | 4.060406099999909 |
| 1012 |  |  | transition_weighted_l1 | 0.5289717795563629 | 4.060406099999909 |
| 1013 |  |  | fate_pearson | 0.28934723571195303 | 4.060406099999909 |
| 1014 |  |  | fate_spearman | 0.37619952262593126 | 4.060406099999909 |
| 1015 |  |  | fate_top1_accuracy | 0.28205128205128205 | 4.060406099999909 |

**dataset = gse239651_transition_panel; benchmark = B2_lineage_transition; task = transition_recovery; split = retrospective_reconstruction; method = cellrank2; method_version = see_method_status**

| **Record** | **repeat** | **seed** | **metric** | **value** | **runtime_seconds** |
| --- | --- | --- | --- | --- | --- |
| 1016 |  |  | fate_pearson | 0.2924215843399057 | 18.17139989999987 |
| 1017 |  |  | fate_spearman | 0.357601455695145 | 18.17139989999987 |
| 1018 |  |  | fate_top1_accuracy | 0.351981351981352 | 18.17139989999987 |

**dataset = macsgestalt_transition_panel; benchmark = B2_lineage_transition; task = transition_recovery; split = retrospective_reconstruction; method = wot; method_version = see_method_status**

| **Record** | **repeat** | **seed** | **metric** | **value** | **runtime_seconds** |
| --- | --- | --- | --- | --- | --- |
| 1019 |  |  | coupling_relative_frobenius | 0.9845342777272762 | 3.2780704000001606 |
| 1020 |  |  | transition_weighted_l1 | 0.6528142047500569 | 3.2780704000001606 |
| 1021 |  |  | fate_pearson | 0.6996693958746467 | 3.2780704000001606 |
| 1022 |  |  | fate_spearman | 0.5498813965385022 | 3.2780704000001606 |
| 1023 |  |  | fate_top1_accuracy | 0.7619047619047619 | 3.2780704000001606 |

**dataset = macsgestalt_transition_panel; benchmark = B2_lineage_transition; task = transition_recovery; split = retrospective_reconstruction; method = moscot; method_version = see_method_status**

| **Record** | **repeat** | **seed** | **metric** | **value** | **runtime_seconds** |
| --- | --- | --- | --- | --- | --- |
| 1024 |  |  | coupling_relative_frobenius | 0.9839620929046672 | 8.9895247999998 |
| 1025 |  |  | transition_weighted_l1 | 0.7191109716690901 | 8.9895247999998 |
| 1026 |  |  | fate_pearson | 0.678623130018003 | 8.9895247999998 |
| 1027 |  |  | fate_spearman | 0.565228977424694 | 8.9895247999998 |
| 1028 |  |  | fate_top1_accuracy | 0.7619047619047619 | 8.9895247999998 |

**dataset = macsgestalt_transition_panel; benchmark = B2_lineage_transition; task = transition_recovery; split = retrospective_reconstruction; method = lineageot; method_version = see_method_status**

| **Record** | **repeat** | **seed** | **metric** | **value** | **runtime_seconds** |
| --- | --- | --- | --- | --- | --- |
| 1029 |  |  | coupling_relative_frobenius | 0.9901266067096133 | 3.2980133000000933 |
| 1030 |  |  | transition_weighted_l1 | 0.7130570022441368 | 3.2980133000000933 |
| 1031 |  |  | fate_pearson | 0.6789880726002673 | 3.2980133000000933 |
| 1032 |  |  | fate_spearman | 0.5499124508501396 | 3.2980133000000933 |
| 1033 |  |  | fate_top1_accuracy | 0.7619047619047619 | 3.2980133000000933 |

**dataset = macsgestalt_transition_panel; benchmark = B2_lineage_transition; task = transition_recovery; split = retrospective_reconstruction; method = cellrank2; method_version = see_method_status**

| **Record** | **repeat** | **seed** | **metric** | **value** | **runtime_seconds** |
| --- | --- | --- | --- | --- | --- |
| 1034 |  |  | fate_pearson | 0.6633623991223192 | 16.732700900000054 |
| 1035 |  |  | fate_spearman | 0.47240906047354825 | 16.732700900000054 |
| 1036 |  |  | fate_top1_accuracy | 0.7380952380952381 | 16.732700900000054 |

**dataset = gse140802_t2_t16; benchmark = B2_lineage_transition; task = transition_recovery; split = retrospective_reconstruction; method = wot; method_version = see_method_status**

| **Record** | **repeat** | **seed** | **metric** | **value** | **runtime_seconds** |
| --- | --- | --- | --- | --- | --- |
| 1037 |  |  | coupling_relative_frobenius | 0.993316708844632 | 3.3637782999999217 |
| 1038 |  |  | transition_weighted_l1 | 0.0634264230787114 | 3.3637782999999217 |
| 1039 |  |  | fate_pearson | 0.6877673287265326 | 3.3637782999999217 |
| 1040 |  |  | fate_spearman | 0.6139979206853131 | 3.3637782999999217 |
| 1041 |  |  | fate_top1_accuracy | 0.55 | 3.3637782999999217 |

**dataset = gse140802_t2_t16; benchmark = B2_lineage_transition; task = transition_recovery; split = retrospective_reconstruction; method = moscot; method_version = see_method_status**

| **Record** | **repeat** | **seed** | **metric** | **value** | **runtime_seconds** |
| --- | --- | --- | --- | --- | --- |
| 1042 |  |  | coupling_relative_frobenius | 0.9933254747328308 | 8.850090099999761 |
| 1043 |  |  | transition_weighted_l1 | 0.06398321642589498 | 8.850090099999761 |
| 1044 |  |  | fate_pearson | 0.6877696582668232 | 8.850090099999761 |
| 1045 |  |  | fate_spearman | 0.6139979206853131 | 8.850090099999761 |
| 1046 |  |  | fate_top1_accuracy | 0.55 | 8.850090099999761 |

**dataset = gse140802_t2_t16; benchmark = B2_lineage_transition; task = transition_recovery; split = retrospective_reconstruction; method = lineageot; method_version = see_method_status**

| **Record** | **repeat** | **seed** | **metric** | **value** | **runtime_seconds** |
| --- | --- | --- | --- | --- | --- |
| 1047 |  |  | coupling_relative_frobenius | 0.9933254747328308 | 4.625729200000023 |
| 1048 |  |  | transition_weighted_l1 | 0.06398321642589481 | 4.625729200000023 |
| 1049 |  |  | fate_pearson | 0.687769658266823 | 4.625729200000023 |
| 1050 |  |  | fate_spearman | 0.6139979206853131 | 4.625729200000023 |
| 1051 |  |  | fate_top1_accuracy | 0.55 | 4.625729200000023 |

**dataset = gse140802_t2_t16; benchmark = B2_lineage_transition; task = transition_recovery; split = retrospective_reconstruction; method = cellrank2; method_version = see_method_status**

| **Record** | **repeat** | **seed** | **metric** | **value** | **runtime_seconds** |
| --- | --- | --- | --- | --- | --- |
| 1052 |  |  | fate_pearson | 0.67655602763955 | 19.560309099999813 |
| 1053 |  |  | fate_spearman | 0.5966825914669532 | 19.560309099999813 |
| 1054 |  |  | fate_top1_accuracy | 0.55 | 19.560309099999813 |

**dataset = gse140802_t2_t9; benchmark = B2_lineage_transition; task = transition_recovery; split = retrospective_reconstruction; method = wot; method_version = see_method_status**

| **Record** | **repeat** | **seed** | **metric** | **value** | **runtime_seconds** |
| --- | --- | --- | --- | --- | --- |
| 1055 |  |  | coupling_relative_frobenius | 0.9985001033741996 | 3.410049699999945 |
| 1056 |  |  | transition_weighted_l1 | 0.08270910860975397 | 3.410049699999945 |
| 1057 |  |  | fate_pearson | 0.5474001244314879 | 3.410049699999945 |
| 1058 |  |  | fate_spearman | 0.5319530890326584 | 3.410049699999945 |
| 1059 |  |  | fate_top1_accuracy | 0.5526315789473685 | 3.410049699999945 |

**dataset = gse140802_t2_t9; benchmark = B2_lineage_transition; task = transition_recovery; split = retrospective_reconstruction; method = moscot; method_version = see_method_status**

| **Record** | **repeat** | **seed** | **metric** | **value** | **runtime_seconds** |
| --- | --- | --- | --- | --- | --- |
| 1060 |  |  | coupling_relative_frobenius | 0.9985046076760864 | 9.358690800000204 |
| 1061 |  |  | transition_weighted_l1 | 0.08193419577073678 | 9.358690800000204 |
| 1062 |  |  | fate_pearson | 0.5474099056098471 | 9.358690800000204 |
| 1063 |  |  | fate_spearman | 0.535114767297792 | 9.358690800000204 |
| 1064 |  |  | fate_top1_accuracy | 0.5526315789473685 | 9.358690800000204 |

**dataset = gse140802_t2_t9; benchmark = B2_lineage_transition; task = transition_recovery; split = retrospective_reconstruction; method = lineageot; method_version = see_method_status**

| **Record** | **repeat** | **seed** | **metric** | **value** | **runtime_seconds** |
| --- | --- | --- | --- | --- | --- |
| 1065 |  |  | coupling_relative_frobenius | 0.9985046076760864 | 3.2721207999998114 |
| 1066 |  |  | transition_weighted_l1 | 0.08193419577073693 | 3.2721207999998114 |
| 1067 |  |  | fate_pearson | 0.5474099056098471 | 3.2721207999998114 |
| 1068 |  |  | fate_spearman | 0.535114767297792 | 3.2721207999998114 |
| 1069 |  |  | fate_top1_accuracy | 0.5526315789473685 | 3.2721207999998114 |

**dataset = gse140802_t2_t9; benchmark = B2_lineage_transition; task = transition_recovery; split = retrospective_reconstruction; method = cellrank2; method_version = see_method_status**

| **Record** | **repeat** | **seed** | **metric** | **value** | **runtime_seconds** |
| --- | --- | --- | --- | --- | --- |
| 1070 |  |  | fate_pearson | 0.49974074260933526 | 16.78689940000004 |
| 1071 |  |  | fate_spearman | 0.49382603672138053 | 16.78689940000004 |
| 1072 |  |  | fate_top1_accuracy | 0.5526315789473685 | 16.78689940000004 |

**dataset = gse239651_transition_panel; benchmark = B2_lineage_transition; task = transition_recovery; split = retrospective_reconstruction; method = mioflow; method_version = see_method_status**

| **Record** | **repeat** | **seed** | **metric** | **value** | **runtime_seconds** |
| --- | --- | --- | --- | --- | --- |
| 1073 |  | 20260916.0 | coupling_relative_frobenius | 0.9990306434785268 | 20.785635199999888 |
| 1074 |  | 20260916.0 | transition_weighted_l1 | 0.15124430580487555 | 20.785635199999888 |
| 1075 |  | 20260916.0 | fate_pearson | 0.4457723560156528 | 20.785635199999888 |
| 1076 |  | 20260916.0 | fate_spearman | 0.4971807678112818 | 20.785635199999888 |
| 1077 |  | 20260916.0 | fate_top1_accuracy | 0.3146853146853147 | 20.785635199999888 |
| 1088 |  | 20260917.0 | coupling_relative_frobenius | 0.9991305893250799 | 15.186479299999974 |
| 1089 |  | 20260917.0 | transition_weighted_l1 | 0.08961296702415208 | 15.186479299999974 |
| 1090 |  | 20260917.0 | fate_pearson | 0.46262135276915517 | 15.186479299999974 |
| 1091 |  | 20260917.0 | fate_spearman | 0.5005027085190424 | 15.186479299999974 |
| 1092 |  | 20260917.0 | fate_top1_accuracy | 0.30536130536130535 | 15.186479299999974 |
| 1103 |  | 20260918.0 | coupling_relative_frobenius | 0.9991713809927019 | 15.013544199999616 |
| 1104 |  | 20260918.0 | transition_weighted_l1 | 0.15078987712976266 | 15.013544199999616 |
| 1105 |  | 20260918.0 | fate_pearson | 0.4238849624211725 | 15.013544199999616 |
| 1106 |  | 20260918.0 | fate_spearman | 0.4832492718476891 | 15.013544199999616 |
| 1107 |  | 20260918.0 | fate_top1_accuracy | 0.289044289044289 | 15.013544199999616 |
| 1118 |  | 20260919.0 | coupling_relative_frobenius | 0.9990234889032085 | 16.295011199999863 |
| 1119 |  | 20260919.0 | transition_weighted_l1 | 0.1090210624912312 | 16.295011199999863 |
| 1120 |  | 20260919.0 | fate_pearson | 0.4623677605801095 | 16.295011199999863 |
| 1121 |  | 20260919.0 | fate_spearman | 0.5082522542819 | 16.295011199999863 |
| 1122 |  | 20260919.0 | fate_top1_accuracy | 0.36596736596736595 | 16.295011199999863 |
| 1133 |  | 20260920.0 | coupling_relative_frobenius | 0.9991693450866036 | 14.431510399999752 |
| 1134 |  | 20260920.0 | transition_weighted_l1 | 0.1761695755833731 | 14.431510399999752 |
| 1135 |  | 20260920.0 | fate_pearson | 0.40010244646046966 | 14.431510399999752 |
| 1136 |  | 20260920.0 | fate_spearman | 0.4787012339391481 | 14.431510399999752 |
| 1137 |  | 20260920.0 | fate_top1_accuracy | 0.2634032634032634 | 14.431510399999752 |

**dataset = gse239651_transition_panel; benchmark = B2_lineage_transition; task = transition_recovery; split = retrospective_reconstruction; method = prescient; method_version = see_method_status**

| **Record** | **repeat** | **seed** | **metric** | **value** | **runtime_seconds** |
| --- | --- | --- | --- | --- | --- |
| 1078 |  | 20260916.0 | coupling_relative_frobenius | 0.9989931339374124 | 12.2056977000002 |
| 1079 |  | 20260916.0 | transition_weighted_l1 | 0.2589233773640866 | 12.2056977000002 |
| 1080 |  | 20260916.0 | fate_pearson | 0.42489057252961243 | 12.2056977000002 |
| 1081 |  | 20260916.0 | fate_spearman | 0.5015536355339979 | 12.2056977000002 |
| 1082 |  | 20260916.0 | fate_top1_accuracy | 0.29836829836829837 | 12.2056977000002 |
| 1093 |  | 20260917.0 | coupling_relative_frobenius | 0.998988910745598 | 12.208909500000118 |
| 1094 |  | 20260917.0 | transition_weighted_l1 | 0.256290709271331 | 12.208909500000118 |
| 1095 |  | 20260917.0 | fate_pearson | 0.42477072574786057 | 12.208909500000118 |
| 1096 |  | 20260917.0 | fate_spearman | 0.49879474172118893 | 12.208909500000118 |
| 1097 |  | 20260917.0 | fate_top1_accuracy | 0.289044289044289 | 12.208909500000118 |
| 1108 |  | 20260918.0 | coupling_relative_frobenius | 0.9989803645534394 | 12.23315360000015 |
| 1109 |  | 20260918.0 | transition_weighted_l1 | 0.25938942190811265 | 12.23315360000015 |
| 1110 |  | 20260918.0 | fate_pearson | 0.4294377567185322 | 12.23315360000015 |
| 1111 |  | 20260918.0 | fate_spearman | 0.5042407915791847 | 12.23315360000015 |
| 1112 |  | 20260918.0 | fate_top1_accuracy | 0.289044289044289 | 12.23315360000015 |
| 1123 |  | 20260919.0 | coupling_relative_frobenius | 0.9989827293994876 | 12.084383700000217 |
| 1124 |  | 20260919.0 | transition_weighted_l1 | 0.2573956860911987 | 12.084383700000217 |
| 1125 |  | 20260919.0 | fate_pearson | 0.4261876335511731 | 12.084383700000217 |
| 1126 |  | 20260919.0 | fate_spearman | 0.5026588018070919 | 12.084383700000217 |
| 1127 |  | 20260919.0 | fate_top1_accuracy | 0.29603729603729606 | 12.084383700000217 |
| 1138 |  | 20260920.0 | coupling_relative_frobenius | 0.99902630185711 | 11.99105150000014 |
| 1139 |  | 20260920.0 | transition_weighted_l1 | 0.26115910638959683 | 11.99105150000014 |
| 1140 |  | 20260920.0 | fate_pearson | 0.42499226661117656 | 11.99105150000014 |
| 1141 |  | 20260920.0 | fate_spearman | 0.5004066267460906 | 11.99105150000014 |
| 1142 |  | 20260920.0 | fate_top1_accuracy | 0.27972027972027974 | 11.99105150000014 |

**dataset = gse239651_transition_panel; benchmark = B2_lineage_transition; task = transition_recovery; split = retrospective_reconstruction; method = tigon; method_version = see_method_status**

| **Record** | **repeat** | **seed** | **metric** | **value** | **runtime_seconds** |
| --- | --- | --- | --- | --- | --- |
| 1083 |  | 20260916.0 | coupling_relative_frobenius | 0.9991391301135109 | 83.49555340000006 |
| 1084 |  | 20260916.0 | transition_weighted_l1 | 0.29741171788925036 | 83.49555340000006 |
| 1085 |  | 20260916.0 | fate_pearson | 0.41484693294079034 | 83.49555340000006 |
| 1086 |  | 20260916.0 | fate_spearman | 0.5015412113083324 | 83.49555340000006 |
| 1087 |  | 20260916.0 | fate_top1_accuracy | 0.289044289044289 | 83.49555340000006 |
| 1098 |  | 20260917.0 | coupling_relative_frobenius | 0.9991422789470741 | 88.4980099999998 |
| 1099 |  | 20260917.0 | transition_weighted_l1 | 0.29768360586486936 | 88.4980099999998 |
| 1100 |  | 20260917.0 | fate_pearson | 0.412946624984967 | 88.4980099999998 |
| 1101 |  | 20260917.0 | fate_spearman | 0.5015799138379464 | 88.4980099999998 |
| 1102 |  | 20260917.0 | fate_top1_accuracy | 0.289044289044289 | 88.4980099999998 |
| 1113 |  | 20260918.0 | coupling_relative_frobenius | 0.9991617292599619 | 89.59955020000007 |
| 1114 |  | 20260918.0 | transition_weighted_l1 | 0.30154038778735015 | 89.59955020000007 |
| 1115 |  | 20260918.0 | fate_pearson | 0.41191261312826627 | 89.59955020000007 |
| 1116 |  | 20260918.0 | fate_spearman | 0.501820187865442 | 89.59955020000007 |
| 1117 |  | 20260918.0 | fate_top1_accuracy | 0.27505827505827507 | 89.59955020000007 |
| 1128 |  | 20260919.0 | coupling_relative_frobenius | 0.99914683448574 | 90.81693210000049 |
| 1129 |  | 20260919.0 | transition_weighted_l1 | 0.2988756784191658 | 90.81693210000049 |
| 1130 |  | 20260919.0 | fate_pearson | 0.4124209980019498 | 90.81693210000049 |
| 1131 |  | 20260919.0 | fate_spearman | 0.5014812988914691 | 90.81693210000049 |
| 1132 |  | 20260919.0 | fate_top1_accuracy | 0.289044289044289 | 90.81693210000049 |
| 1143 |  | 20260920.0 | coupling_relative_frobenius | 0.9991584221037534 | 87.81660160000047 |
| 1144 |  | 20260920.0 | transition_weighted_l1 | 0.29869847968356433 | 87.81660160000047 |
| 1145 |  | 20260920.0 | fate_pearson | 0.41282626195714295 | 87.81660160000047 |
| 1146 |  | 20260920.0 | fate_spearman | 0.5017362252219898 | 87.81660160000047 |
| 1147 |  | 20260920.0 | fate_top1_accuracy | 0.289044289044289 | 87.81660160000047 |

**dataset = macsgestalt_transition_panel; benchmark = B2_lineage_transition; task = transition_recovery; split = retrospective_reconstruction; method = mioflow; method_version = see_method_status**

| **Record** | **repeat** | **seed** | **metric** | **value** | **runtime_seconds** |
| --- | --- | --- | --- | --- | --- |
| 1148 |  | 20260916.0 | coupling_relative_frobenius | 1.003561646595841 | 11.044323500000246 |
| 1149 |  | 20260916.0 | transition_weighted_l1 | 1.3202806211689606 | 11.044323500000246 |
| 1150 |  | 20260916.0 | fate_pearson | 0.3605240131826958 | 11.044323500000246 |
| 1151 |  | 20260916.0 | fate_spearman | 0.38192696254805125 | 11.044323500000246 |
| 1152 |  | 20260916.0 | fate_top1_accuracy | 0.4523809523809524 | 11.044323500000246 |
| 1163 |  | 20260917.0 | coupling_relative_frobenius | 1.00300696197363 | 11.484083600000304 |
| 1164 |  | 20260917.0 | transition_weighted_l1 | 1.335157605047449 | 11.484083600000304 |
| 1165 |  | 20260917.0 | fate_pearson | 0.42690216489193766 | 11.484083600000304 |
| 1166 |  | 20260917.0 | fate_spearman | 0.4082066544270583 | 11.484083600000304 |
| 1167 |  | 20260917.0 | fate_top1_accuracy | 0.5476190476190477 | 11.484083600000304 |
| 1178 |  | 20260918.0 | coupling_relative_frobenius | 1.0020868131043903 | 11.78794199999993 |
| 1179 |  | 20260918.0 | transition_weighted_l1 | 1.33810838528661 | 11.78794199999993 |
| 1180 |  | 20260918.0 | fate_pearson | 0.407414803511065 | 11.78794199999993 |
| 1181 |  | 20260918.0 | fate_spearman | 0.41991117438915326 | 11.78794199999993 |
| 1182 |  | 20260918.0 | fate_top1_accuracy | 0.47619047619047616 | 11.78794199999993 |
| 1193 |  | 20260919.0 | coupling_relative_frobenius | 0.9913139871393861 | 11.830186999999569 |
| 1194 |  | 20260919.0 | transition_weighted_l1 | 1.2174254614207702 | 11.830186999999569 |
| 1195 |  | 20260919.0 | fate_pearson | 0.48569311533340637 | 11.830186999999569 |
| 1196 |  | 20260919.0 | fate_spearman | 0.44278581561224356 | 11.830186999999569 |
| 1197 |  | 20260919.0 | fate_top1_accuracy | 0.5476190476190477 | 11.830186999999569 |
| 1208 |  | 20260920.0 | coupling_relative_frobenius | 1.001329332116878 | 12.103159299999788 |
| 1209 |  | 20260920.0 | transition_weighted_l1 | 1.2143644353824117 | 12.103159299999788 |
| 1210 |  | 20260920.0 | fate_pearson | 0.44898998880477314 | 12.103159299999788 |
| 1211 |  | 20260920.0 | fate_spearman | 0.41080833999840616 | 12.103159299999788 |
| 1212 |  | 20260920.0 | fate_top1_accuracy | 0.5 | 12.103159299999788 |

**dataset = macsgestalt_transition_panel; benchmark = B2_lineage_transition; task = transition_recovery; split = retrospective_reconstruction; method = prescient; method_version = see_method_status**

| **Record** | **repeat** | **seed** | **metric** | **value** | **runtime_seconds** |
| --- | --- | --- | --- | --- | --- |
| 1153 |  | 20260916.0 | coupling_relative_frobenius | 0.91914396672175 | 4.970087499999863 |
| 1154 |  | 20260916.0 | transition_weighted_l1 | 0.6758110035515296 | 4.970087499999863 |
| 1155 |  | 20260916.0 | fate_pearson | 0.7472279095775682 | 4.970087499999863 |
| 1156 |  | 20260916.0 | fate_spearman | 0.621023741518435 | 4.970087499999863 |
| 1157 |  | 20260916.0 | fate_top1_accuracy | 0.7857142857142857 | 4.970087499999863 |
| 1168 |  | 20260917.0 | coupling_relative_frobenius | 0.9273143508480319 | 4.637583499999892 |
| 1169 |  | 20260917.0 | transition_weighted_l1 | 0.7075988608674194 | 4.637583499999892 |
| 1170 |  | 20260917.0 | fate_pearson | 0.7463828770088128 | 4.637583499999892 |
| 1171 |  | 20260917.0 | fate_spearman | 0.6092115493554686 | 4.637583499999892 |
| 1172 |  | 20260917.0 | fate_top1_accuracy | 0.7857142857142857 | 4.637583499999892 |
| 1183 |  | 20260918.0 | coupling_relative_frobenius | 0.9280846625566778 | 5.187965800000711 |
| 1184 |  | 20260918.0 | transition_weighted_l1 | 0.7301966681847867 | 5.187965800000711 |
| 1185 |  | 20260918.0 | fate_pearson | 0.7424762694139533 | 5.187965800000711 |
| 1186 |  | 20260918.0 | fate_spearman | 0.6152303803833008 | 5.187965800000711 |
| 1187 |  | 20260918.0 | fate_top1_accuracy | 0.7619047619047619 | 5.187965800000711 |
| 1198 |  | 20260919.0 | coupling_relative_frobenius | 0.9232862449047136 | 4.753263999999945 |
| 1199 |  | 20260919.0 | transition_weighted_l1 | 0.7041725058485173 | 4.753263999999945 |
| 1200 |  | 20260919.0 | fate_pearson | 0.7482161474805122 | 4.753263999999945 |
| 1201 |  | 20260919.0 | fate_spearman | 0.6109473235889521 | 4.753263999999945 |
| 1202 |  | 20260919.0 | fate_top1_accuracy | 0.7857142857142857 | 4.753263999999945 |
| 1213 |  | 20260920.0 | coupling_relative_frobenius | 0.934123566609378 | 4.769642400000521 |
| 1214 |  | 20260920.0 | transition_weighted_l1 | 0.762888689901581 | 4.769642400000521 |
| 1215 |  | 20260920.0 | fate_pearson | 0.7375832873134232 | 4.769642400000521 |
| 1216 |  | 20260920.0 | fate_spearman | 0.6031986769551924 | 4.769642400000521 |
| 1217 |  | 20260920.0 | fate_top1_accuracy | 0.7857142857142857 | 4.769642400000521 |

**dataset = macsgestalt_transition_panel; benchmark = B2_lineage_transition; task = transition_recovery; split = retrospective_reconstruction; method = tigon; method_version = see_method_status**

| **Record** | **repeat** | **seed** | **metric** | **value** | **runtime_seconds** |
| --- | --- | --- | --- | --- | --- |
| 1158 |  | 20260916.0 | coupling_relative_frobenius | 0.9299757145560769 | 19.89577809999992 |
| 1159 |  | 20260916.0 | transition_weighted_l1 | 0.7174029970849015 | 19.89577809999992 |
| 1160 |  | 20260916.0 | fate_pearson | 0.7512864617151734 | 19.89577809999992 |
| 1161 |  | 20260916.0 | fate_spearman | 0.6165217856723526 | 19.89577809999992 |
| 1162 |  | 20260916.0 | fate_top1_accuracy | 0.8095238095238095 | 19.89577809999992 |
| 1173 |  | 20260917.0 | coupling_relative_frobenius | 0.9354347742344629 | 19.72212709999985 |
| 1174 |  | 20260917.0 | transition_weighted_l1 | 0.7526388148578879 | 19.72212709999985 |
| 1175 |  | 20260917.0 | fate_pearson | 0.7409543006229233 | 19.72212709999985 |
| 1176 |  | 20260917.0 | fate_spearman | 0.6115787545899247 | 19.72212709999985 |
| 1177 |  | 20260917.0 | fate_top1_accuracy | 0.7857142857142857 | 19.72212709999985 |
| 1188 |  | 20260918.0 | coupling_relative_frobenius | 0.9541013749938375 | 19.38227749999988 |
| 1189 |  | 20260918.0 | transition_weighted_l1 | 0.9283452308512878 | 19.38227749999988 |
| 1190 |  | 20260918.0 | fate_pearson | 0.7211851023313107 | 19.38227749999988 |
| 1191 |  | 20260918.0 | fate_spearman | 0.6107621708556491 | 19.38227749999988 |
| 1192 |  | 20260918.0 | fate_top1_accuracy | 0.7619047619047619 | 19.38227749999988 |
| 1203 |  | 20260919.0 | coupling_relative_frobenius | 0.9536765217854136 | 20.06999889999952 |
| 1204 |  | 20260919.0 | transition_weighted_l1 | 0.8469570562504978 | 20.06999889999952 |
| 1205 |  | 20260919.0 | fate_pearson | 0.7351985165769845 | 20.06999889999952 |
| 1206 |  | 20260919.0 | fate_spearman | 0.6168417691672761 | 20.06999889999952 |
| 1207 |  | 20260919.0 | fate_top1_accuracy | 0.7619047619047619 | 20.06999889999952 |
| 1218 |  | 20260920.0 | coupling_relative_frobenius | 0.9263516686389875 | 20.34762430000046 |
| 1219 |  | 20260920.0 | transition_weighted_l1 | 0.7061222389859736 | 20.34762430000046 |
| 1220 |  | 20260920.0 | fate_pearson | 0.7483483145378048 | 20.34762430000046 |
| 1221 |  | 20260920.0 | fate_spearman | 0.6170738402927092 | 20.34762430000046 |
| 1222 |  | 20260920.0 | fate_top1_accuracy | 0.8095238095238095 | 20.34762430000046 |

**dataset = gse140802_t2_t16; benchmark = B2_lineage_transition; task = transition_recovery; split = retrospective_reconstruction; method = mioflow; method_version = see_method_status**

| **Record** | **repeat** | **seed** | **metric** | **value** | **runtime_seconds** |
| --- | --- | --- | --- | --- | --- |
| 1223 |  | 20260916.0 | coupling_relative_frobenius | 0.9931673643287151 | 12.013453700000357 |
| 1224 |  | 20260916.0 | transition_weighted_l1 | 0.12310121097551577 | 12.013453700000357 |
| 1225 |  | 20260916.0 | fate_pearson | 0.6810018026138667 | 12.013453700000357 |
| 1226 |  | 20260916.0 | fate_spearman | 0.6065945304599655 | 12.013453700000357 |
| 1227 |  | 20260916.0 | fate_top1_accuracy | 0.55 | 12.013453700000357 |
| 1238 |  | 20260917.0 | coupling_relative_frobenius | 0.9935871894927563 | 11.470924399999603 |
| 1239 |  | 20260917.0 | transition_weighted_l1 | 0.14834119800970394 | 11.470924399999603 |
| 1240 |  | 20260917.0 | fate_pearson | 0.6685203639977878 | 11.470924399999603 |
| 1241 |  | 20260917.0 | fate_spearman | 0.6186308852868236 | 11.470924399999603 |
| 1242 |  | 20260917.0 | fate_top1_accuracy | 0.55 | 11.470924399999603 |
| 1253 |  | 20260918.0 | coupling_relative_frobenius | 0.9933128210872902 | 12.363045500000226 |
| 1254 |  | 20260918.0 | transition_weighted_l1 | 0.14061398827279162 | 12.363045500000226 |
| 1255 |  | 20260918.0 | fate_pearson | 0.6730154071381761 | 12.363045500000226 |
| 1256 |  | 20260918.0 | fate_spearman | 0.6107744208463829 | 12.363045500000226 |
| 1257 |  | 20260918.0 | fate_top1_accuracy | 0.55 | 12.363045500000226 |
| 1268 |  | 20260919.0 | coupling_relative_frobenius | 0.9931021585305273 | 11.767347699999846 |
| 1269 |  | 20260919.0 | transition_weighted_l1 | 0.12814822404067627 | 11.767347699999846 |
| 1270 |  | 20260919.0 | fate_pearson | 0.6776899817881051 | 11.767347699999846 |
| 1271 |  | 20260919.0 | fate_spearman | 0.6070230070884197 | 11.767347699999846 |
| 1272 |  | 20260919.0 | fate_top1_accuracy | 0.55 | 11.767347699999846 |
| 1283 |  | 20260920.0 | coupling_relative_frobenius | 0.9934705059604081 | 11.402560200000153 |
| 1284 |  | 20260920.0 | transition_weighted_l1 | 0.14199395317972316 | 11.402560200000153 |
| 1285 |  | 20260920.0 | fate_pearson | 0.6718380460006822 | 11.402560200000153 |
| 1286 |  | 20260920.0 | fate_spearman | 0.610372550112441 | 11.402560200000153 |
| 1287 |  | 20260920.0 | fate_top1_accuracy | 0.55 | 11.402560200000153 |

**dataset = gse140802_t2_t16; benchmark = B2_lineage_transition; task = transition_recovery; split = retrospective_reconstruction; method = prescient; method_version = see_method_status**

| **Record** | **repeat** | **seed** | **metric** | **value** | **runtime_seconds** |
| --- | --- | --- | --- | --- | --- |
| 1228 |  | 20260916.0 | coupling_relative_frobenius | 0.9898361198925616 | 4.76325420000012 |
| 1229 |  | 20260916.0 | transition_weighted_l1 | 0.02752394374287959 | 4.76325420000012 |
| 1230 |  | 20260916.0 | fate_pearson | 0.6857582828457882 | 4.76325420000012 |
| 1231 |  | 20260916.0 | fate_spearman | 0.6021522172747454 | 4.76325420000012 |
| 1232 |  | 20260916.0 | fate_top1_accuracy | 0.55 | 4.76325420000012 |
| 1243 |  | 20260917.0 | coupling_relative_frobenius | 0.9895187892038269 | 4.930187599999954 |
| 1244 |  | 20260917.0 | transition_weighted_l1 | 0.04041397518849717 | 4.930187599999954 |
| 1245 |  | 20260917.0 | fate_pearson | 0.6897292478617645 | 4.930187599999954 |
| 1246 |  | 20260917.0 | fate_spearman | 0.6141203973215643 | 4.930187599999954 |
| 1247 |  | 20260917.0 | fate_top1_accuracy | 0.55 | 4.930187599999954 |
| 1258 |  | 20260918.0 | coupling_relative_frobenius | 0.9895855511195137 | 4.707105800000136 |
| 1259 |  | 20260918.0 | transition_weighted_l1 | 0.025040151718579224 | 4.707105800000136 |
| 1260 |  | 20260918.0 | fate_pearson | 0.6812914512747107 | 4.707105800000136 |
| 1261 |  | 20260918.0 | fate_spearman | 0.6045381468293745 | 4.707105800000136 |
| 1262 |  | 20260918.0 | fate_top1_accuracy | 0.55 | 4.707105800000136 |
| 1273 |  | 20260919.0 | coupling_relative_frobenius | 0.988352785860451 | 4.738925100000415 |
| 1274 |  | 20260919.0 | transition_weighted_l1 | 0.02667510125642086 | 4.738925100000415 |
| 1275 |  | 20260919.0 | fate_pearson | 0.6898767443702183 | 4.738925100000415 |
| 1276 |  | 20260919.0 | fate_spearman | 0.600838769337358 | 4.738925100000415 |
| 1277 |  | 20260919.0 | fate_top1_accuracy | 0.55 | 4.738925100000415 |
| 1288 |  | 20260920.0 | coupling_relative_frobenius | 0.9893671615277244 | 4.724510500000179 |
| 1289 |  | 20260920.0 | transition_weighted_l1 | 0.02573092121783031 | 4.724510500000179 |
| 1290 |  | 20260920.0 | fate_pearson | 0.6739427338800289 | 4.724510500000179 |
| 1291 |  | 20260920.0 | fate_spearman | 0.5960219559088142 | 4.724510500000179 |
| 1292 |  | 20260920.0 | fate_top1_accuracy | 0.55 | 4.724510500000179 |

**dataset = gse140802_t2_t16; benchmark = B2_lineage_transition; task = transition_recovery; split = retrospective_reconstruction; method = tigon; method_version = see_method_status**

| **Record** | **repeat** | **seed** | **metric** | **value** | **runtime_seconds** |
| --- | --- | --- | --- | --- | --- |
| 1233 |  | 20260916.0 | coupling_relative_frobenius | 0.9932713686217347 | 24.13769130000037 |
| 1234 |  | 20260916.0 | transition_weighted_l1 | 0.10975185802219731 | 24.13769130000037 |
| 1235 |  | 20260916.0 | fate_pearson | 0.6819716979552833 | 24.13769130000037 |
| 1236 |  | 20260916.0 | fate_spearman | 0.6052923729585108 | 24.13769130000037 |
| 1237 |  | 20260916.0 | fate_top1_accuracy | 0.55 | 24.13769130000037 |
| 1248 |  | 20260917.0 | coupling_relative_frobenius | 0.9932723625317816 | 23.325883700000304 |
| 1249 |  | 20260917.0 | transition_weighted_l1 | 0.10975811833751334 | 23.325883700000304 |
| 1250 |  | 20260917.0 | fate_pearson | 0.6820383698968957 | 23.325883700000304 |
| 1251 |  | 20260917.0 | fate_spearman | 0.6076782372425027 | 23.325883700000304 |
| 1252 |  | 20260917.0 | fate_top1_accuracy | 0.55 | 23.325883700000304 |
| 1263 |  | 20260918.0 | coupling_relative_frobenius | 0.9932709221334548 | 21.92691970000033 |
| 1264 |  | 20260918.0 | transition_weighted_l1 | 0.10971196373946275 | 21.92691970000033 |
| 1265 |  | 20260918.0 | fate_pearson | 0.6819729290361615 | 21.92691970000033 |
| 1266 |  | 20260918.0 | fate_spearman | 0.6098168197392374 | 21.92691970000033 |
| 1267 |  | 20260918.0 | fate_top1_accuracy | 0.55 | 21.92691970000033 |
| 1278 |  | 20260919.0 | coupling_relative_frobenius | 0.9932696642305019 | 22.086581900000056 |
| 1279 |  | 20260919.0 | transition_weighted_l1 | 0.10953817767962025 | 22.086581900000056 |
| 1280 |  | 20260919.0 | fate_pearson | 0.6820279187780509 | 22.086581900000056 |
| 1281 |  | 20260919.0 | fate_spearman | 0.6117600201283903 | 22.086581900000056 |
| 1282 |  | 20260919.0 | fate_top1_accuracy | 0.55 | 22.086581900000056 |
| 1293 |  | 20260920.0 | coupling_relative_frobenius | 0.9932726606554857 | 23.77054660000067 |
| 1294 |  | 20260920.0 | transition_weighted_l1 | 0.10981740927327081 | 23.77054660000067 |
| 1295 |  | 20260920.0 | fate_pearson | 0.681993818914351 | 23.77054660000067 |
| 1296 |  | 20260920.0 | fate_spearman | 0.6089527806652585 | 23.77054660000067 |
| 1297 |  | 20260920.0 | fate_top1_accuracy | 0.55 | 23.77054660000067 |

**dataset = gse140802_t2_t9; benchmark = B2_lineage_transition; task = transition_recovery; split = retrospective_reconstruction; method = mioflow; method_version = see_method_status**

| **Record** | **repeat** | **seed** | **metric** | **value** | **runtime_seconds** |
| --- | --- | --- | --- | --- | --- |
| 1298 |  | 20260916.0 | coupling_relative_frobenius | 0.9975218930432536 | 13.74652270000024 |
| 1299 |  | 20260916.0 | transition_weighted_l1 | 0.1161616005967642 | 13.74652270000024 |
| 1300 |  | 20260916.0 | fate_pearson | 0.5320828005890784 | 13.74652270000024 |
| 1301 |  | 20260916.0 | fate_spearman | 0.5338474510666846 | 13.74652270000024 |
| 1302 |  | 20260916.0 | fate_top1_accuracy | 0.5526315789473685 | 13.74652270000024 |
| 1313 |  | 20260917.0 | coupling_relative_frobenius | 0.998089853442605 | 13.662903800000096 |
| 1314 |  | 20260917.0 | transition_weighted_l1 | 0.17415058978497658 | 13.662903800000096 |
| 1315 |  | 20260917.0 | fate_pearson | 0.5128638259823372 | 13.662903800000096 |
| 1316 |  | 20260917.0 | fate_spearman | 0.5275253677125754 | 13.662903800000096 |
| 1317 |  | 20260917.0 | fate_top1_accuracy | 0.5526315789473685 | 13.662903800000096 |
| 1328 |  | 20260918.0 | coupling_relative_frobenius | 0.9979621503877042 | 13.745216499999517 |
| 1329 |  | 20260918.0 | transition_weighted_l1 | 0.15875943908411158 | 13.745216499999517 |
| 1330 |  | 20260918.0 | fate_pearson | 0.5184957074281172 | 13.745216499999517 |
| 1331 |  | 20260918.0 | fate_spearman | 0.5305098878431908 | 13.745216499999517 |
| 1332 |  | 20260918.0 | fate_top1_accuracy | 0.5526315789473685 | 13.745216499999517 |
| 1343 |  | 20260919.0 | coupling_relative_frobenius | 0.9978430034987719 | 14.100329599999895 |
| 1344 |  | 20260919.0 | transition_weighted_l1 | 0.14472116391286802 | 14.100329599999895 |
| 1345 |  | 20260919.0 | fate_pearson | 0.5230953087251339 | 14.100329599999895 |
| 1346 |  | 20260919.0 | fate_spearman | 0.5297222697884598 | 14.100329599999895 |
| 1347 |  | 20260919.0 | fate_top1_accuracy | 0.5526315789473685 | 14.100329599999895 |
| 1358 |  | 20260920.0 | coupling_relative_frobenius | 0.9981916210019995 | 12.533916000000318 |
| 1359 |  | 20260920.0 | transition_weighted_l1 | 0.17486659452716632 | 12.533916000000318 |
| 1360 |  | 20260920.0 | fate_pearson | 0.511146297378903 | 12.533916000000318 |
| 1361 |  | 20260920.0 | fate_spearman | 0.5044788677468828 | 12.533916000000318 |
| 1362 |  | 20260920.0 | fate_top1_accuracy | 0.24342105263157895 | 12.533916000000318 |

**dataset = gse140802_t2_t9; benchmark = B2_lineage_transition; task = transition_recovery; split = retrospective_reconstruction; method = prescient; method_version = see_method_status**

| **Record** | **repeat** | **seed** | **metric** | **value** | **runtime_seconds** |
| --- | --- | --- | --- | --- | --- |
| 1303 |  | 20260916.0 | coupling_relative_frobenius | 0.997119789970787 | 6.1162162000000535 |
| 1304 |  | 20260916.0 | transition_weighted_l1 | 0.02638442834552323 | 6.1162162000000535 |
| 1305 |  | 20260916.0 | fate_pearson | 0.5415687988006731 | 6.1162162000000535 |
| 1306 |  | 20260916.0 | fate_spearman | 0.5308736356354434 | 6.1162162000000535 |
| 1307 |  | 20260916.0 | fate_top1_accuracy | 0.5526315789473685 | 6.1162162000000535 |
| 1318 |  | 20260917.0 | coupling_relative_frobenius | 0.9968915287909355 | 5.9572438000004695 |
| 1319 |  | 20260917.0 | transition_weighted_l1 | 0.01837516633798943 | 5.9572438000004695 |
| 1320 |  | 20260917.0 | fate_pearson | 0.5462248525924126 | 5.9572438000004695 |
| 1321 |  | 20260917.0 | fate_spearman | 0.5324479246967051 | 5.9572438000004695 |
| 1322 |  | 20260917.0 | fate_top1_accuracy | 0.5526315789473685 | 5.9572438000004695 |
| 1333 |  | 20260918.0 | coupling_relative_frobenius | 0.9970729972402402 | 6.154530699999668 |
| 1334 |  | 20260918.0 | transition_weighted_l1 | 0.026131759999358052 | 6.154530699999668 |
| 1335 |  | 20260918.0 | fate_pearson | 0.5392771694323856 | 6.154530699999668 |
| 1336 |  | 20260918.0 | fate_spearman | 0.5289425066240131 | 6.154530699999668 |
| 1337 |  | 20260918.0 | fate_top1_accuracy | 0.5526315789473685 | 6.154530699999668 |
| 1348 |  | 20260919.0 | coupling_relative_frobenius | 0.9971257777717669 | 6.35136059999968 |
| 1349 |  | 20260919.0 | transition_weighted_l1 | 0.03113853323446794 | 6.35136059999968 |
| 1350 |  | 20260919.0 | fate_pearson | 0.5443327544981922 | 6.35136059999968 |
| 1351 |  | 20260919.0 | fate_spearman | 0.530476594663815 | 6.35136059999968 |
| 1352 |  | 20260919.0 | fate_top1_accuracy | 0.5526315789473685 | 6.35136059999968 |
| 1363 |  | 20260920.0 | coupling_relative_frobenius | 0.9972446330235923 | 6.240835099999458 |
| 1364 |  | 20260920.0 | transition_weighted_l1 | 0.02575313877475246 | 6.240835099999458 |
| 1365 |  | 20260920.0 | fate_pearson | 0.5500566576323485 | 6.240835099999458 |
| 1366 |  | 20260920.0 | fate_spearman | 0.5395875911639605 | 6.240835099999458 |
| 1367 |  | 20260920.0 | fate_top1_accuracy | 0.5526315789473685 | 6.240835099999458 |

**dataset = gse140802_t2_t9; benchmark = B2_lineage_transition; task = transition_recovery; split = retrospective_reconstruction; method = tigon; method_version = see_method_status**

| **Record** | **repeat** | **seed** | **metric** | **value** | **runtime_seconds** |
| --- | --- | --- | --- | --- | --- |
| 1308 |  | 20260916.0 | coupling_relative_frobenius | 0.9982439942852676 | 41.948921600000176 |
| 1309 |  | 20260916.0 | transition_weighted_l1 | 0.152639589286058 | 41.948921600000176 |
| 1310 |  | 20260916.0 | fate_pearson | 0.5244065451815121 | 41.948921600000176 |
| 1311 |  | 20260916.0 | fate_spearman | 0.5301091764604083 | 41.948921600000176 |
| 1312 |  | 20260916.0 | fate_top1_accuracy | 0.5526315789473685 | 41.948921600000176 |
| 1323 |  | 20260917.0 | coupling_relative_frobenius | 0.9982413821611922 | 42.25566600000002 |
| 1324 |  | 20260917.0 | transition_weighted_l1 | 0.15248766675439096 | 42.25566600000002 |
| 1325 |  | 20260917.0 | fate_pearson | 0.5244607500097886 | 42.25566600000002 |
| 1326 |  | 20260917.0 | fate_spearman | 0.5319742370616225 | 42.25566600000002 |
| 1327 |  | 20260917.0 | fate_top1_accuracy | 0.5526315789473685 | 42.25566600000002 |
| 1338 |  | 20260918.0 | coupling_relative_frobenius | 0.9982429368571443 | 42.34471649999978 |
| 1339 |  | 20260918.0 | transition_weighted_l1 | 0.15247571488102915 | 42.34471649999978 |
| 1340 |  | 20260918.0 | fate_pearson | 0.5244415169912201 | 42.34471649999978 |
| 1341 |  | 20260918.0 | fate_spearman | 0.5324268658293035 | 42.34471649999978 |
| 1342 |  | 20260918.0 | fate_top1_accuracy | 0.5526315789473685 | 42.34471649999978 |
| 1353 |  | 20260919.0 | coupling_relative_frobenius | 0.9982425935671609 | 42.208458599999176 |
| 1354 |  | 20260919.0 | transition_weighted_l1 | 0.1523226551068403 | 42.208458599999176 |
| 1355 |  | 20260919.0 | fate_pearson | 0.5245765493452326 | 42.208458599999176 |
| 1356 |  | 20260919.0 | fate_spearman | 0.5316546308176608 | 42.208458599999176 |
| 1357 |  | 20260919.0 | fate_top1_accuracy | 0.5526315789473685 | 42.208458599999176 |
| 1368 |  | 20260920.0 | coupling_relative_frobenius | 0.998243736860358 | 42.86661980000008 |
| 1369 |  | 20260920.0 | transition_weighted_l1 | 0.152423661141033 | 42.86661980000008 |
| 1370 |  | 20260920.0 | fate_pearson | 0.5245675066070653 | 42.86661980000008 |
| 1371 |  | 20260920.0 | fate_spearman | 0.5326905306015322 | 42.86661980000008 |
| 1372 |  | 20260920.0 | fate_top1_accuracy | 0.5526315789473685 | 42.86661980000008 |

**dataset = GSE239651_expt1; benchmark = B3_prospective_composition; task = clone_level_future_state_composition; split = locked_existing; method = PERSIST-IOT; method_version = locked_primary_rerun**

| **Record** | **repeat** | **seed** | **metric** | **value** | **runtime_seconds** |
| --- | --- | --- | --- | --- | --- |
| 1373 |  | 20260916.0 | cross_entropy | 1.0370164120742835 |  |
| 1377 |  | 20260916.0 | sinkhorn_divergence | 0.1024034576297865 |  |
| 1381 |  | 20260916.0 | top1_accuracy | 0.6443394288062462 |  |
| 1385 |  | 20260916.0 | brier_score | 0.2851258100055829 |  |
| 1389 |  | 20260916.0 | expected_calibration_error | 0.0887882896405101 |  |

**dataset = GSE239651_expt1; benchmark = B3_prospective_composition; task = clone_level_future_state_composition; split = locked_existing; method = PERSIST-no-IOT; method_version = locked_primary_rerun**

| **Record** | **repeat** | **seed** | **metric** | **value** | **runtime_seconds** |
| --- | --- | --- | --- | --- | --- |
| 1374 |  | 20260916.0 | cross_entropy | 6.955561833086343 |  |
| 1378 |  | 20260916.0 | sinkhorn_divergence | 0.1784467192593128 |  |
| 1382 |  | 20260916.0 | top1_accuracy | 0.4970207520032874 |  |
| 1386 |  | 20260916.0 | brier_score | 0.5164761222815634 |  |
| 1390 |  | 20260916.0 | expected_calibration_error | 0.2392807832146085 |  |

**dataset = GSE239651_expt2; benchmark = B3_prospective_composition; task = clone_level_future_state_composition; split = locked_existing; method = PERSIST-IOT; method_version = locked_primary_rerun**

| **Record** | **repeat** | **seed** | **metric** | **value** | **runtime_seconds** |
| --- | --- | --- | --- | --- | --- |
| 1375 |  | 20260916.0 | cross_entropy | 0.9859026620053872 |  |
| 1379 |  | 20260916.0 | sinkhorn_divergence | 0.1110045964504361 |  |
| 1383 |  | 20260916.0 | top1_accuracy | 0.6819870648135407 |  |
| 1387 |  | 20260916.0 | brier_score | 0.3592909630817728 |  |
| 1391 |  | 20260916.0 | expected_calibration_error | 0.0915102460671916 |  |

**dataset = GSE239651_expt2; benchmark = B3_prospective_composition; task = clone_level_future_state_composition; split = locked_existing; method = PERSIST-no-IOT; method_version = locked_primary_rerun**

| **Record** | **repeat** | **seed** | **metric** | **value** | **runtime_seconds** |
| --- | --- | --- | --- | --- | --- |
| 1376 |  | 20260916.0 | cross_entropy | 6.549742700206335 |  |
| 1380 |  | 20260916.0 | sinkhorn_divergence | 0.1965103520808689 |  |
| 1384 |  | 20260916.0 | top1_accuracy | 0.475024081464153 |  |
| 1388 |  | 20260916.0 | brier_score | 0.597281963260918 |  |
| 1392 |  | 20260916.0 | expected_calibration_error | 0.2342588619523692 |  |

**dataset = GSE239651_expt2; benchmark = B3_prospective_composition; task = population_future_state_composition; split = expt1_to_locked_expt2; method = source-carry-forward; method_version = see_method_status**

| **Record** | **repeat** | **seed** | **metric** | **value** | **runtime_seconds** |
| --- | --- | --- | --- | --- | --- |
| 1393 | gse239651_dmso_day3::DMSO | 20260916.0 | cross_entropy | 0.9453034735830608 | 0.1561231999994561 |
| 1410 | gse239651_dmso_day6::DMSO | 20260916.0 | cross_entropy | 1.0432801252777673 | 0.1849554000000353 |
| 1427 | gse239651_dmso_day69::DMSO | 20260916.0 | cross_entropy | 0.905026595273441 | 0.2196296000001893 |
| 1444 | gse239651_dmso_day98::DMSO | 20260916.0 | cross_entropy | 0.994094655963954 | 0.2158336000002236 |
| 1461 | gse239651_ispinesib_day3::ispinesib | 20260916.0 | cross_entropy | 0.8354442804489903 | 0.2572759000004225 |
| 1478 | gse239651_ispinesib_day6::ispinesib | 20260916.0 | cross_entropy | 1.0450354764491314 | 0.2187656999994942 |
| 1495 | gse239651_ispinesib_day69::ispinesib | 20260916.0 | cross_entropy | 0.9997951524140428 | 0.2187080000003334 |
| 1512 | gse239651_ispinesib_day98::ispinesib | 20260916.0 | cross_entropy | 0.9712333308951135 | 0.2183641999999963 |
| 1699 | gse239651_dmso_day3::DMSO | 20260916.0 | sinkhorn_divergence | 0.0057580065874268 | 0.1561231999994561 |
| 1716 | gse239651_dmso_day6::DMSO | 20260916.0 | sinkhorn_divergence | 0.0422153065329133 | 0.1849554000000353 |
| 1733 | gse239651_dmso_day69::DMSO | 20260916.0 | sinkhorn_divergence | 0.0720306082192917 | 0.2196296000001893 |
| 1750 | gse239651_dmso_day98::DMSO | 20260916.0 | sinkhorn_divergence | 0.033227232366992 | 0.2158336000002236 |
| 1767 | gse239651_ispinesib_day3::ispinesib | 20260916.0 | sinkhorn_divergence | 0.0694115623419238 | 0.2572759000004225 |
| 1784 | gse239651_ispinesib_day6::ispinesib | 20260916.0 | sinkhorn_divergence | 0.0738155793871123 | 0.2187656999994942 |
| 1801 | gse239651_ispinesib_day69::ispinesib | 20260916.0 | sinkhorn_divergence | 0.0661734460421103 | 0.2187080000003334 |
| 1818 | gse239651_ispinesib_day98::ispinesib | 20260916.0 | sinkhorn_divergence | 0.066458495815634 | 0.2183641999999963 |
| 2005 | gse239651_dmso_day3::DMSO | 20260916.0 | top1_accuracy | 1.0 | 0.1561231999994561 |
| 2022 | gse239651_dmso_day6::DMSO | 20260916.0 | top1_accuracy | 1.0 | 0.1849554000000353 |
| 2039 | gse239651_dmso_day69::DMSO | 20260916.0 | top1_accuracy | 1.0 | 0.2196296000001893 |
| 2056 | gse239651_dmso_day98::DMSO | 20260916.0 | top1_accuracy | 1.0 | 0.2158336000002236 |
| 2073 | gse239651_ispinesib_day3::ispinesib | 20260916.0 | top1_accuracy | 1.0 | 0.2572759000004225 |
| 2090 | gse239651_ispinesib_day6::ispinesib | 20260916.0 | top1_accuracy | 1.0 | 0.2187656999994942 |
| 2107 | gse239651_ispinesib_day69::ispinesib | 20260916.0 | top1_accuracy | 1.0 | 0.2187080000003334 |
| 2124 | gse239651_ispinesib_day98::ispinesib | 20260916.0 | top1_accuracy | 1.0 | 0.2183641999999963 |
| 2311 | gse239651_dmso_day3::DMSO | 20260916.0 | multiclass_brier | 0.0011126460665103 | 0.1561231999994561 |
| 2328 | gse239651_dmso_day6::DMSO | 20260916.0 | multiclass_brier | 0.0541439647402735 | 0.1849554000000353 |
| 2345 | gse239651_dmso_day69::DMSO | 20260916.0 | multiclass_brier | 0.204327930688437 | 0.2196296000001893 |
| 2362 | gse239651_dmso_day98::DMSO | 20260916.0 | multiclass_brier | 0.0368181356973505 | 0.2158336000002236 |
| 2379 | gse239651_ispinesib_day3::ispinesib | 20260916.0 | multiclass_brier | 0.2276792778685996 | 0.2572759000004225 |
| 2396 | gse239651_ispinesib_day6::ispinesib | 20260916.0 | multiclass_brier | 0.1832575668179687 | 0.2187656999994942 |
| 2413 | gse239651_ispinesib_day69::ispinesib | 20260916.0 | multiclass_brier | 0.1658227600582185 | 0.2187080000003334 |
| 2430 | gse239651_ispinesib_day98::ispinesib | 20260916.0 | multiclass_brier | 0.1450375981627853 | 0.2183641999999963 |

**dataset = GSE239651_expt2; benchmark = B3_prospective_composition; task = population_future_state_composition; split = expt1_to_locked_expt2; method = development-target-transfer; method_version = see_method_status**

| **Record** | **repeat** | **seed** | **metric** | **value** | **runtime_seconds** |
| --- | --- | --- | --- | --- | --- |
| 1394 | gse239651_dmso_day3::DMSO | 20260916.0 | cross_entropy | 0.9513839598786596 | 0.1614518000005773 |
| 1411 | gse239651_dmso_day6::DMSO | 20260916.0 | cross_entropy | 0.964639344856504 | 0.1975183000004108 |
| 1428 | gse239651_dmso_day69::DMSO | 20260916.0 | cross_entropy | 0.7632222554906667 | 0.1586776000003738 |
| 1445 | gse239651_dmso_day98::DMSO | 20260916.0 | cross_entropy | 1.0183799358048429 | 0.1661822000005486 |
| 1462 | gse239651_ispinesib_day3::ispinesib | 20260916.0 | cross_entropy | 0.636907795064926 | 0.1867031000001588 |
| 1479 | gse239651_ispinesib_day6::ispinesib | 20260916.0 | cross_entropy | 0.8110292202747693 | 0.1451276000007055 |
| 1496 | gse239651_ispinesib_day69::ispinesib | 20260916.0 | cross_entropy | 0.818458460421515 | 0.167641399999411 |
| 1513 | gse239651_ispinesib_day98::ispinesib | 20260916.0 | cross_entropy | 0.9127610849878 | 0.1554246000014245 |
| 1700 | gse239651_dmso_day3::DMSO | 20260916.0 | sinkhorn_divergence | 0.0147489786815011 | 0.1614518000005773 |
| 1717 | gse239651_dmso_day6::DMSO | 20260916.0 | sinkhorn_divergence | 0.0132550046481408 | 0.1975183000004108 |
| 1734 | gse239651_dmso_day69::DMSO | 20260916.0 | sinkhorn_divergence | 0.0693790297893027 | 0.1586776000003738 |
| 1751 | gse239651_dmso_day98::DMSO | 20260916.0 | sinkhorn_divergence | 0.0626060625227409 | 0.1661822000005486 |
| 1768 | gse239651_ispinesib_day3::ispinesib | 20260916.0 | sinkhorn_divergence | 0.0292859573852665 | 0.1867031000001588 |
| 1785 | gse239651_ispinesib_day6::ispinesib | 20260916.0 | sinkhorn_divergence | 0.0208356901415962 | 0.1451276000007055 |
| 1802 | gse239651_ispinesib_day69::ispinesib | 20260916.0 | sinkhorn_divergence | 0.0321664019287899 | 0.167641399999411 |
| 1819 | gse239651_ispinesib_day98::ispinesib | 20260916.0 | sinkhorn_divergence | 0.039882955974131 | 0.1554246000014245 |
| 2006 | gse239651_dmso_day3::DMSO | 20260916.0 | top1_accuracy | 0.0 | 0.1614518000005773 |
| 2023 | gse239651_dmso_day6::DMSO | 20260916.0 | top1_accuracy | 1.0 | 0.1975183000004108 |
| 2040 | gse239651_dmso_day69::DMSO | 20260916.0 | top1_accuracy | 1.0 | 0.1586776000003738 |
| 2057 | gse239651_dmso_day98::DMSO | 20260916.0 | top1_accuracy | 1.0 | 0.1661822000005486 |
| 2074 | gse239651_ispinesib_day3::ispinesib | 20260916.0 | top1_accuracy | 1.0 | 0.1867031000001588 |
| 2091 | gse239651_ispinesib_day6::ispinesib | 20260916.0 | top1_accuracy | 1.0 | 0.1451276000007055 |
| 2108 | gse239651_ispinesib_day69::ispinesib | 20260916.0 | top1_accuracy | 1.0 | 0.167641399999411 |
| 2125 | gse239651_ispinesib_day98::ispinesib | 20260916.0 | top1_accuracy | 1.0 | 0.1554246000014245 |
| 2312 | gse239651_dmso_day3::DMSO | 20260916.0 | multiclass_brier | 0.0054947247353515 | 0.1614518000005773 |
| 2329 | gse239651_dmso_day6::DMSO | 20260916.0 | multiclass_brier | 0.0054901443327234 | 0.1975183000004108 |
| 2346 | gse239651_dmso_day69::DMSO | 20260916.0 | multiclass_brier | 0.0511871607876314 | 0.1586776000003738 |
| 2363 | gse239651_dmso_day98::DMSO | 20260916.0 | multiclass_brier | 0.0171207989199834 | 0.1661822000005486 |
| 2380 | gse239651_ispinesib_day3::ispinesib | 20260916.0 | multiclass_brier | 0.0119143028967951 | 0.1867031000001588 |
| 2397 | gse239651_ispinesib_day6::ispinesib | 20260916.0 | multiclass_brier | 0.0121673024362299 | 0.1451276000007055 |
| 2414 | gse239651_ispinesib_day69::ispinesib | 20260916.0 | multiclass_brier | 0.0312449371789714 | 0.167641399999411 |
| 2431 | gse239651_ispinesib_day98::ispinesib | 20260916.0 | multiclass_brier | 0.0241215259705728 | 0.1554246000014245 |

**dataset = GSE239651_expt2; benchmark = B3_prospective_composition; task = population_future_state_composition; split = expt1_to_locked_expt2; method = prescient; method_version = see_method_status**

| **Record** | **repeat** | **seed** | **metric** | **value** | **runtime_seconds** |
| --- | --- | --- | --- | --- | --- |
| 1395 | gse239651_dmso_day3::DMSO | 20260916.0 | cross_entropy | 0.988116830378835 | 16.968943099999706 |
| 1396 | gse239651_dmso_day3::DMSO | 20260917.0 | cross_entropy | 0.989816494885139 | 16.857099100000596 |
| 1397 | gse239651_dmso_day3::DMSO | 20260918.0 | cross_entropy | 0.9870363226600358 | 17.230403200000183 |
| 1398 | gse239651_dmso_day3::DMSO | 20260919.0 | cross_entropy | 0.9881711142543196 | 17.66758740000023 |
| 1399 | gse239651_dmso_day3::DMSO | 20260920.0 | cross_entropy | 0.9892621542786588 | 18.999660499999663 |
| 1412 | gse239651_dmso_day6::DMSO | 20260916.0 | cross_entropy | 1.021477522156026 | 18.23375170000054 |
| 1413 | gse239651_dmso_day6::DMSO | 20260917.0 | cross_entropy | 1.0223202622410534 | 17.320406599999842 |
| 1414 | gse239651_dmso_day6::DMSO | 20260918.0 | cross_entropy | 1.0223459302433011 | 17.21284259999993 |
| 1415 | gse239651_dmso_day6::DMSO | 20260919.0 | cross_entropy | 1.019392034281461 | 17.633608499999355 |
| 1416 | gse239651_dmso_day6::DMSO | 20260920.0 | cross_entropy | 1.0209650708717106 | 17.563787300000513 |
| 1429 | gse239651_dmso_day69::DMSO | 20260916.0 | cross_entropy | 0.9067866721940832 | 17.763366000000133 |
| 1430 | gse239651_dmso_day69::DMSO | 20260917.0 | cross_entropy | 0.9035112164362826 | 17.281015699999443 |
| 1431 | gse239651_dmso_day69::DMSO | 20260918.0 | cross_entropy | 0.9040541804832004 | 16.85154969999985 |
| 1432 | gse239651_dmso_day69::DMSO | 20260919.0 | cross_entropy | 0.901981319722351 | 17.224731799999972 |
| 1433 | gse239651_dmso_day69::DMSO | 20260920.0 | cross_entropy | 0.9022455285621018 | 16.87041489999956 |
| 1446 | gse239651_dmso_day98::DMSO | 20260916.0 | cross_entropy | 1.0361537987184122 | 18.396034000000327 |
| 1447 | gse239651_dmso_day98::DMSO | 20260917.0 | cross_entropy | 1.0366309020954825 | 17.86212599999999 |
| 1448 | gse239651_dmso_day98::DMSO | 20260918.0 | cross_entropy | 1.0355081935653916 | 17.750324900000123 |
| 1449 | gse239651_dmso_day98::DMSO | 20260919.0 | cross_entropy | 1.0345819205947353 | 17.46751789999962 |
| 1450 | gse239651_dmso_day98::DMSO | 20260920.0 | cross_entropy | 1.0367795670619897 | 17.304312700000082 |
| 1463 | gse239651_ispinesib_day3::ispinesib | 20260916.0 | cross_entropy | 0.7472219863409492 | 22.08064430000013 |
| 1464 | gse239651_ispinesib_day3::ispinesib | 20260917.0 | cross_entropy | 0.7504327558235545 | 20.677488299999823 |
| 1465 | gse239651_ispinesib_day3::ispinesib | 20260918.0 | cross_entropy | 0.7559797446459916 | 20.723336900000504 |
| 1466 | gse239651_ispinesib_day3::ispinesib | 20260919.0 | cross_entropy | 0.751555563905548 | 21.28973709999991 |
| 1467 | gse239651_ispinesib_day3::ispinesib | 20260920.0 | cross_entropy | 0.7524161703598998 | 21.415869599999496 |
| 1480 | gse239651_ispinesib_day6::ispinesib | 20260916.0 | cross_entropy | 0.8697304394507896 | 17.542980799998986 |
| 1481 | gse239651_ispinesib_day6::ispinesib | 20260917.0 | cross_entropy | 0.8722063018856927 | 16.59724190000088 |
| 1482 | gse239651_ispinesib_day6::ispinesib | 20260918.0 | cross_entropy | 0.8785745348663916 | 17.076539499999853 |
| 1483 | gse239651_ispinesib_day6::ispinesib | 20260919.0 | cross_entropy | 0.8750376200076058 | 16.597240299999612 |
| 1484 | gse239651_ispinesib_day6::ispinesib | 20260920.0 | cross_entropy | 0.8674423409671306 | 17.31988039999851 |
| 1497 | gse239651_ispinesib_day69::ispinesib | 20260916.0 | cross_entropy | 0.9546607390302708 | 18.12783660000059 |
| 1498 | gse239651_ispinesib_day69::ispinesib | 20260917.0 | cross_entropy | 0.9554981033373392 | 17.08631940000123 |
| 1499 | gse239651_ispinesib_day69::ispinesib | 20260918.0 | cross_entropy | 0.9565598731468228 | 17.45098550000148 |
| 1500 | gse239651_ispinesib_day69::ispinesib | 20260919.0 | cross_entropy | 0.9553990554379824 | 17.094880399999965 |
| 1501 | gse239651_ispinesib_day69::ispinesib | 20260920.0 | cross_entropy | 0.9531725951236174 | 17.22556960000111 |
| 1514 | gse239651_ispinesib_day98::ispinesib | 20260916.0 | cross_entropy | 0.8740512551101787 | 17.698182500000257 |
| 1515 | gse239651_ispinesib_day98::ispinesib | 20260917.0 | cross_entropy | 0.8749556806784632 | 16.309630499999912 |
| 1516 | gse239651_ispinesib_day98::ispinesib | 20260918.0 | cross_entropy | 0.8809506503156996 | 16.448863300000085 |
| 1517 | gse239651_ispinesib_day98::ispinesib | 20260919.0 | cross_entropy | 0.8752058461695048 | 16.350034600000072 |
| 1518 | gse239651_ispinesib_day98::ispinesib | 20260920.0 | cross_entropy | 0.8773505330478346 | 16.955900299999485 |
| 1701 | gse239651_dmso_day3::DMSO | 20260916.0 | sinkhorn_divergence | 0.0484359552541962 | 16.968943099999706 |
| 1702 | gse239651_dmso_day3::DMSO | 20260917.0 | sinkhorn_divergence | 0.0498232219979004 | 16.857099100000596 |
| 1703 | gse239651_dmso_day3::DMSO | 20260918.0 | sinkhorn_divergence | 0.0473019104429932 | 17.230403200000183 |
| 1704 | gse239651_dmso_day3::DMSO | 20260919.0 | sinkhorn_divergence | 0.0494100277006611 | 17.66758740000023 |
| 1705 | gse239651_dmso_day3::DMSO | 20260920.0 | sinkhorn_divergence | 0.0506210759415023 | 18.999660499999663 |
| 1718 | gse239651_dmso_day6::DMSO | 20260916.0 | sinkhorn_divergence | 0.0600155910432542 | 18.23375170000054 |
| 1719 | gse239651_dmso_day6::DMSO | 20260917.0 | sinkhorn_divergence | 0.0605280658686593 | 17.320406599999842 |
| 1720 | gse239651_dmso_day6::DMSO | 20260918.0 | sinkhorn_divergence | 0.0599071013628854 | 17.21284259999993 |
| 1721 | gse239651_dmso_day6::DMSO | 20260919.0 | sinkhorn_divergence | 0.0585772074998889 | 17.633608499999355 |
| 1722 | gse239651_dmso_day6::DMSO | 20260920.0 | sinkhorn_divergence | 0.060448827634423 | 17.563787300000513 |
| 1735 | gse239651_dmso_day69::DMSO | 20260916.0 | sinkhorn_divergence | 0.1155175354670177 | 17.763366000000133 |
| 1736 | gse239651_dmso_day69::DMSO | 20260917.0 | sinkhorn_divergence | 0.1148593617033118 | 17.281015699999443 |
| 1737 | gse239651_dmso_day69::DMSO | 20260918.0 | sinkhorn_divergence | 0.113137478804419 | 16.85154969999985 |
| 1738 | gse239651_dmso_day69::DMSO | 20260919.0 | sinkhorn_divergence | 0.1138840999224217 | 17.224731799999972 |
| 1739 | gse239651_dmso_day69::DMSO | 20260920.0 | sinkhorn_divergence | 0.1148544099874131 | 16.87041489999956 |
| 1752 | gse239651_dmso_day98::DMSO | 20260916.0 | sinkhorn_divergence | 0.0833812886536702 | 18.396034000000327 |
| 1753 | gse239651_dmso_day98::DMSO | 20260917.0 | sinkhorn_divergence | 0.0838291069229871 | 17.86212599999999 |
| 1754 | gse239651_dmso_day98::DMSO | 20260918.0 | sinkhorn_divergence | 0.0832024685785667 | 17.750324900000123 |
| 1755 | gse239651_dmso_day98::DMSO | 20260919.0 | sinkhorn_divergence | 0.0839688516455 | 17.46751789999962 |
| 1756 | gse239651_dmso_day98::DMSO | 20260920.0 | sinkhorn_divergence | 0.0856431908998516 | 17.304312700000082 |
| 1769 | gse239651_ispinesib_day3::ispinesib | 20260916.0 | sinkhorn_divergence | 0.0892693436432935 | 22.08064430000013 |
| 1770 | gse239651_ispinesib_day3::ispinesib | 20260917.0 | sinkhorn_divergence | 0.0920792760515613 | 20.677488299999823 |
| 1771 | gse239651_ispinesib_day3::ispinesib | 20260918.0 | sinkhorn_divergence | 0.0918704183403168 | 20.723336900000504 |
| 1772 | gse239651_ispinesib_day3::ispinesib | 20260919.0 | sinkhorn_divergence | 0.0910900945278862 | 21.28973709999991 |
| 1773 | gse239651_ispinesib_day3::ispinesib | 20260920.0 | sinkhorn_divergence | 0.0926773926224235 | 21.415869599999496 |
| 1786 | gse239651_ispinesib_day6::ispinesib | 20260916.0 | sinkhorn_divergence | 0.058186910530651 | 17.542980799998986 |
| 1787 | gse239651_ispinesib_day6::ispinesib | 20260917.0 | sinkhorn_divergence | 0.0606091250093585 | 16.59724190000088 |
| 1788 | gse239651_ispinesib_day6::ispinesib | 20260918.0 | sinkhorn_divergence | 0.0620267674229096 | 17.076539499999853 |
| 1789 | gse239651_ispinesib_day6::ispinesib | 20260919.0 | sinkhorn_divergence | 0.0609685109908412 | 16.597240299999612 |
| 1790 | gse239651_ispinesib_day6::ispinesib | 20260920.0 | sinkhorn_divergence | 0.0595712318732289 | 17.31988039999851 |
| 1803 | gse239651_ispinesib_day69::ispinesib | 20260916.0 | sinkhorn_divergence | 0.0947951486549829 | 18.12783660000059 |
| 1804 | gse239651_ispinesib_day69::ispinesib | 20260917.0 | sinkhorn_divergence | 0.095459924641461 | 17.08631940000123 |
| 1805 | gse239651_ispinesib_day69::ispinesib | 20260918.0 | sinkhorn_divergence | 0.0945470135483168 | 17.45098550000148 |
| 1806 | gse239651_ispinesib_day69::ispinesib | 20260919.0 | sinkhorn_divergence | 0.0949192565135455 | 17.094880399999965 |
| 1807 | gse239651_ispinesib_day69::ispinesib | 20260920.0 | sinkhorn_divergence | 0.0955417858721151 | 17.22556960000111 |
| 1820 | gse239651_ispinesib_day98::ispinesib | 20260916.0 | sinkhorn_divergence | 0.0635666522975798 | 17.698182500000257 |
| 1821 | gse239651_ispinesib_day98::ispinesib | 20260917.0 | sinkhorn_divergence | 0.0657766401830957 | 16.309630499999912 |
| 1822 | gse239651_ispinesib_day98::ispinesib | 20260918.0 | sinkhorn_divergence | 0.0668271868668956 | 16.448863300000085 |
| 1823 | gse239651_ispinesib_day98::ispinesib | 20260919.0 | sinkhorn_divergence | 0.0643794898128316 | 16.350034600000072 |
| 1824 | gse239651_ispinesib_day98::ispinesib | 20260920.0 | sinkhorn_divergence | 0.0665345336532973 | 16.955900299999485 |
| 2007 | gse239651_dmso_day3::DMSO | 20260916.0 | top1_accuracy | 1.0 | 16.968943099999706 |
| 2008 | gse239651_dmso_day3::DMSO | 20260917.0 | top1_accuracy | 1.0 | 16.857099100000596 |
| 2009 | gse239651_dmso_day3::DMSO | 20260918.0 | top1_accuracy | 1.0 | 17.230403200000183 |
| 2010 | gse239651_dmso_day3::DMSO | 20260919.0 | top1_accuracy | 1.0 | 17.66758740000023 |
| 2011 | gse239651_dmso_day3::DMSO | 20260920.0 | top1_accuracy | 1.0 | 18.999660499999663 |
| 2024 | gse239651_dmso_day6::DMSO | 20260916.0 | top1_accuracy | 1.0 | 18.23375170000054 |
| 2025 | gse239651_dmso_day6::DMSO | 20260917.0 | top1_accuracy | 1.0 | 17.320406599999842 |
| 2026 | gse239651_dmso_day6::DMSO | 20260918.0 | top1_accuracy | 1.0 | 17.21284259999993 |
| 2027 | gse239651_dmso_day6::DMSO | 20260919.0 | top1_accuracy | 1.0 | 17.633608499999355 |
| 2028 | gse239651_dmso_day6::DMSO | 20260920.0 | top1_accuracy | 1.0 | 17.563787300000513 |
| 2041 | gse239651_dmso_day69::DMSO | 20260916.0 | top1_accuracy | 1.0 | 17.763366000000133 |
| 2042 | gse239651_dmso_day69::DMSO | 20260917.0 | top1_accuracy | 1.0 | 17.281015699999443 |
| 2043 | gse239651_dmso_day69::DMSO | 20260918.0 | top1_accuracy | 1.0 | 16.85154969999985 |
| 2044 | gse239651_dmso_day69::DMSO | 20260919.0 | top1_accuracy | 1.0 | 17.224731799999972 |
| 2045 | gse239651_dmso_day69::DMSO | 20260920.0 | top1_accuracy | 1.0 | 16.87041489999956 |
| 2058 | gse239651_dmso_day98::DMSO | 20260916.0 | top1_accuracy | 1.0 | 18.396034000000327 |
| 2059 | gse239651_dmso_day98::DMSO | 20260917.0 | top1_accuracy | 1.0 | 17.86212599999999 |
| 2060 | gse239651_dmso_day98::DMSO | 20260918.0 | top1_accuracy | 1.0 | 17.750324900000123 |
| 2061 | gse239651_dmso_day98::DMSO | 20260919.0 | top1_accuracy | 1.0 | 17.46751789999962 |
| 2062 | gse239651_dmso_day98::DMSO | 20260920.0 | top1_accuracy | 1.0 | 17.304312700000082 |
| 2075 | gse239651_ispinesib_day3::ispinesib | 20260916.0 | top1_accuracy | 1.0 | 22.08064430000013 |
| 2076 | gse239651_ispinesib_day3::ispinesib | 20260917.0 | top1_accuracy | 1.0 | 20.677488299999823 |
| 2077 | gse239651_ispinesib_day3::ispinesib | 20260918.0 | top1_accuracy | 1.0 | 20.723336900000504 |
| 2078 | gse239651_ispinesib_day3::ispinesib | 20260919.0 | top1_accuracy | 1.0 | 21.28973709999991 |
| 2079 | gse239651_ispinesib_day3::ispinesib | 20260920.0 | top1_accuracy | 1.0 | 21.415869599999496 |
| 2092 | gse239651_ispinesib_day6::ispinesib | 20260916.0 | top1_accuracy | 1.0 | 17.542980799998986 |
| 2093 | gse239651_ispinesib_day6::ispinesib | 20260917.0 | top1_accuracy | 1.0 | 16.59724190000088 |
| 2094 | gse239651_ispinesib_day6::ispinesib | 20260918.0 | top1_accuracy | 1.0 | 17.076539499999853 |
| 2095 | gse239651_ispinesib_day6::ispinesib | 20260919.0 | top1_accuracy | 1.0 | 16.597240299999612 |
| 2096 | gse239651_ispinesib_day6::ispinesib | 20260920.0 | top1_accuracy | 1.0 | 17.31988039999851 |
| 2109 | gse239651_ispinesib_day69::ispinesib | 20260916.0 | top1_accuracy | 1.0 | 18.12783660000059 |
| 2110 | gse239651_ispinesib_day69::ispinesib | 20260917.0 | top1_accuracy | 1.0 | 17.08631940000123 |
| 2111 | gse239651_ispinesib_day69::ispinesib | 20260918.0 | top1_accuracy | 1.0 | 17.45098550000148 |
| 2112 | gse239651_ispinesib_day69::ispinesib | 20260919.0 | top1_accuracy | 1.0 | 17.094880399999965 |
| 2113 | gse239651_ispinesib_day69::ispinesib | 20260920.0 | top1_accuracy | 1.0 | 17.22556960000111 |
| 2126 | gse239651_ispinesib_day98::ispinesib | 20260916.0 | top1_accuracy | 1.0 | 17.698182500000257 |
| 2127 | gse239651_ispinesib_day98::ispinesib | 20260917.0 | top1_accuracy | 1.0 | 16.309630499999912 |
| 2128 | gse239651_ispinesib_day98::ispinesib | 20260918.0 | top1_accuracy | 1.0 | 16.448863300000085 |
| 2129 | gse239651_ispinesib_day98::ispinesib | 20260919.0 | top1_accuracy | 1.0 | 16.350034600000072 |
| 2130 | gse239651_ispinesib_day98::ispinesib | 20260920.0 | top1_accuracy | 1.0 | 16.955900299999485 |
| 2313 | gse239651_dmso_day3::DMSO | 20260916.0 | multiclass_brier | 0.002259532277463 | 16.968943099999706 |
| 2314 | gse239651_dmso_day3::DMSO | 20260917.0 | multiclass_brier | 0.0025637870877154 | 16.857099100000596 |
| 2315 | gse239651_dmso_day3::DMSO | 20260918.0 | multiclass_brier | 0.0023029091613764 | 17.230403200000183 |
| 2316 | gse239651_dmso_day3::DMSO | 20260919.0 | multiclass_brier | 0.0025739836546477 | 17.66758740000023 |
| 2317 | gse239651_dmso_day3::DMSO | 20260920.0 | multiclass_brier | 0.0025929635205282 | 18.999660499999663 |
| 2330 | gse239651_dmso_day6::DMSO | 20260916.0 | multiclass_brier | 0.0236788267797141 | 18.23375170000054 |
| 2331 | gse239651_dmso_day6::DMSO | 20260917.0 | multiclass_brier | 0.0234920286573976 | 17.320406599999842 |
| 2332 | gse239651_dmso_day6::DMSO | 20260918.0 | multiclass_brier | 0.0239262161997716 | 17.21284259999993 |
| 2333 | gse239651_dmso_day6::DMSO | 20260919.0 | multiclass_brier | 0.0225947070426399 | 17.633608499999355 |
| 2334 | gse239651_dmso_day6::DMSO | 20260920.0 | multiclass_brier | 0.0232885959366488 | 17.563787300000513 |
| 2347 | gse239651_dmso_day69::DMSO | 20260916.0 | multiclass_brier | 0.1581866743059591 | 17.763366000000133 |
| 2348 | gse239651_dmso_day69::DMSO | 20260917.0 | multiclass_brier | 0.1563726637811303 | 17.281015699999443 |
| 2349 | gse239651_dmso_day69::DMSO | 20260918.0 | multiclass_brier | 0.158810373192059 | 16.85154969999985 |
| 2350 | gse239651_dmso_day69::DMSO | 20260919.0 | multiclass_brier | 0.1561260197453837 | 17.224731799999972 |
| 2351 | gse239651_dmso_day69::DMSO | 20260920.0 | multiclass_brier | 0.1551669661087373 | 16.87041489999956 |
| 2364 | gse239651_dmso_day98::DMSO | 20260916.0 | multiclass_brier | 0.0315977588730373 | 18.396034000000327 |
| 2365 | gse239651_dmso_day98::DMSO | 20260917.0 | multiclass_brier | 0.0312851102595558 | 17.86212599999999 |
| 2366 | gse239651_dmso_day98::DMSO | 20260918.0 | multiclass_brier | 0.0314164566454223 | 17.750324900000123 |
| 2367 | gse239651_dmso_day98::DMSO | 20260919.0 | multiclass_brier | 0.03004070313915 | 17.46751789999962 |
| 2368 | gse239651_dmso_day98::DMSO | 20260920.0 | multiclass_brier | 0.0304758743485865 | 17.304312700000082 |
| 2381 | gse239651_ispinesib_day3::ispinesib | 20260916.0 | multiclass_brier | 0.1142115091429974 | 22.08064430000013 |
| 2382 | gse239651_ispinesib_day3::ispinesib | 20260917.0 | multiclass_brier | 0.1153732353946578 | 20.677488299999823 |
| 2383 | gse239651_ispinesib_day3::ispinesib | 20260918.0 | multiclass_brier | 0.1207365137813238 | 20.723336900000504 |
| 2384 | gse239651_ispinesib_day3::ispinesib | 20260919.0 | multiclass_brier | 0.1177723787320185 | 21.28973709999991 |
| 2385 | gse239651_ispinesib_day3::ispinesib | 20260920.0 | multiclass_brier | 0.1161076948481532 | 21.415869599999496 |
| 2398 | gse239651_ispinesib_day6::ispinesib | 20260916.0 | multiclass_brier | 0.0322746577400538 | 17.542980799998986 |
| 2399 | gse239651_ispinesib_day6::ispinesib | 20260917.0 | multiclass_brier | 0.0330876395291671 | 16.59724190000088 |
| 2400 | gse239651_ispinesib_day6::ispinesib | 20260918.0 | multiclass_brier | 0.0372840890555846 | 17.076539499999853 |
| 2401 | gse239651_ispinesib_day6::ispinesib | 20260919.0 | multiclass_brier | 0.0345374304861293 | 16.597240299999612 |
| 2402 | gse239651_ispinesib_day6::ispinesib | 20260920.0 | multiclass_brier | 0.0311386536751697 | 17.31988039999851 |
| 2415 | gse239651_ispinesib_day69::ispinesib | 20260916.0 | multiclass_brier | 0.1026108735479904 | 18.12783660000059 |
| 2416 | gse239651_ispinesib_day69::ispinesib | 20260917.0 | multiclass_brier | 0.1027363287824218 | 17.08631940000123 |
| 2417 | gse239651_ispinesib_day69::ispinesib | 20260918.0 | multiclass_brier | 0.1040987694361634 | 17.45098550000148 |
| 2418 | gse239651_ispinesib_day69::ispinesib | 20260919.0 | multiclass_brier | 0.1036513048308584 | 17.094880399999965 |
| 2419 | gse239651_ispinesib_day69::ispinesib | 20260920.0 | multiclass_brier | 0.1020899895367715 | 17.22556960000111 |
| 2432 | gse239651_ispinesib_day98::ispinesib | 20260916.0 | multiclass_brier | 0.0434028132899428 | 17.698182500000257 |
| 2433 | gse239651_ispinesib_day98::ispinesib | 20260917.0 | multiclass_brier | 0.0431312357660619 | 16.309630499999912 |
| 2434 | gse239651_ispinesib_day98::ispinesib | 20260918.0 | multiclass_brier | 0.047951070502577 | 16.448863300000085 |
| 2435 | gse239651_ispinesib_day98::ispinesib | 20260919.0 | multiclass_brier | 0.0452041200981334 | 16.350034600000072 |
| 2436 | gse239651_ispinesib_day98::ispinesib | 20260920.0 | multiclass_brier | 0.043781827979546 | 16.955900299999485 |

**dataset = GSE239651_expt2; benchmark = B3_prospective_composition; task = population_future_state_composition; split = expt1_to_locked_expt2; method = tigon; method_version = see_method_status**

| **Record** | **repeat** | **seed** | **metric** | **value** | **runtime_seconds** |
| --- | --- | --- | --- | --- | --- |
| 1400 | gse239651_dmso_day3::DMSO | 20260916.0 | cross_entropy | 0.9521334769425784 | 115.63007089999974 |
| 1401 | gse239651_dmso_day3::DMSO | 20260917.0 | cross_entropy | 0.9474000160733492 | 115.80890920000002 |
| 1402 | gse239651_dmso_day3::DMSO | 20260918.0 | cross_entropy | 0.9451291075523968 | 117.5844432999993 |
| 1403 | gse239651_dmso_day3::DMSO | 20260919.0 | cross_entropy | 0.9530187402299086 | 120.40731559999948 |
| 1404 | gse239651_dmso_day3::DMSO | 20260920.0 | cross_entropy | 0.9507245634793974 | 117.94565039999998 |
| 1417 | gse239651_dmso_day6::DMSO | 20260916.0 | cross_entropy | 1.0457180002094923 | 121.35117600000012 |
| 1418 | gse239651_dmso_day6::DMSO | 20260917.0 | cross_entropy | 1.0443281132316218 | 116.50757129999964 |
| 1419 | gse239651_dmso_day6::DMSO | 20260918.0 | cross_entropy | 1.0400927214989828 | 107.0824107999997 |
| 1420 | gse239651_dmso_day6::DMSO | 20260919.0 | cross_entropy | 1.0421299204079324 | 106.08582360000035 |
| 1421 | gse239651_dmso_day6::DMSO | 20260920.0 | cross_entropy | 1.0455438105838946 | 110.6642585999998 |
| 1434 | gse239651_dmso_day69::DMSO | 20260916.0 | cross_entropy | 0.9096970106701734 | 105.98453539999991 |
| 1435 | gse239651_dmso_day69::DMSO | 20260917.0 | cross_entropy | 0.9060121931522171 | 103.9811552000001 |
| 1436 | gse239651_dmso_day69::DMSO | 20260918.0 | cross_entropy | 0.9028858670419416 | 105.9265863999999 |
| 1437 | gse239651_dmso_day69::DMSO | 20260919.0 | cross_entropy | 0.9042311439382232 | 101.50770660000036 |
| 1438 | gse239651_dmso_day69::DMSO | 20260920.0 | cross_entropy | 0.9085035893131652 | 104.91881490000014 |
| 1451 | gse239651_dmso_day98::DMSO | 20260916.0 | cross_entropy | 0.9961311493271858 | 122.23007440000038 |
| 1452 | gse239651_dmso_day98::DMSO | 20260917.0 | cross_entropy | 0.9942793669235644 | 133.0048740000002 |
| 1453 | gse239651_dmso_day98::DMSO | 20260918.0 | cross_entropy | 0.9916083647675102 | 133.1269975999994 |
| 1454 | gse239651_dmso_day98::DMSO | 20260919.0 | cross_entropy | 0.9986215270093676 | 133.16438820000076 |
| 1455 | gse239651_dmso_day98::DMSO | 20260920.0 | cross_entropy | 0.9955591024797272 | 134.2977045000007 |
| 1468 | gse239651_ispinesib_day3::ispinesib | 20260916.0 | cross_entropy | 0.8427814256732556 | 132.31578749999971 |
| 1469 | gse239651_ispinesib_day3::ispinesib | 20260917.0 | cross_entropy | 0.8379110997507561 | 130.98438579999856 |
| 1470 | gse239651_ispinesib_day3::ispinesib | 20260918.0 | cross_entropy | 0.8356117987830278 | 118.62514700000064 |
| 1471 | gse239651_ispinesib_day3::ispinesib | 20260919.0 | cross_entropy | 0.8383913317752819 | 111.87683459999973 |
| 1472 | gse239651_ispinesib_day3::ispinesib | 20260920.0 | cross_entropy | 0.8375538378855529 | 112.4667773000001 |
| 1485 | gse239651_ispinesib_day6::ispinesib | 20260916.0 | cross_entropy | 1.0498956096662082 | 115.93260999999984 |
| 1486 | gse239651_ispinesib_day6::ispinesib | 20260917.0 | cross_entropy | 1.045320488718252 | 112.2782141000007 |
| 1487 | gse239651_ispinesib_day6::ispinesib | 20260918.0 | cross_entropy | 1.039842537217211 | 110.78718130000016 |
| 1488 | gse239651_ispinesib_day6::ispinesib | 20260919.0 | cross_entropy | 1.044393810896549 | 112.84757480000007 |
| 1489 | gse239651_ispinesib_day6::ispinesib | 20260920.0 | cross_entropy | 1.045580012201369 | 113.0381097999998 |
| 1502 | gse239651_ispinesib_day69::ispinesib | 20260916.0 | cross_entropy | 1.0020514767410678 | 113.43000900000152 |
| 1503 | gse239651_ispinesib_day69::ispinesib | 20260917.0 | cross_entropy | 1.000600370242782 | 114.5526747999993 |
| 1504 | gse239651_ispinesib_day69::ispinesib | 20260918.0 | cross_entropy | 0.9966490280047926 | 116.72172610000052 |
| 1505 | gse239651_ispinesib_day69::ispinesib | 20260919.0 | cross_entropy | 0.9970005690914938 | 112.28214730000036 |
| 1506 | gse239651_ispinesib_day69::ispinesib | 20260920.0 | cross_entropy | 1.0025436591175714 | 112.40618619999988 |
| 1519 | gse239651_ispinesib_day98::ispinesib | 20260916.0 | cross_entropy | 0.9775043339666536 | 111.89460319999851 |
| 1520 | gse239651_ispinesib_day98::ispinesib | 20260917.0 | cross_entropy | 0.97242346967758 | 109.15929640000104 |
| 1521 | gse239651_ispinesib_day98::ispinesib | 20260918.0 | cross_entropy | 0.968583240548352 | 109.93124539999916 |
| 1522 | gse239651_ispinesib_day98::ispinesib | 20260919.0 | cross_entropy | 0.9771744646011356 | 112.23595419999948 |
| 1523 | gse239651_ispinesib_day98::ispinesib | 20260920.0 | cross_entropy | 0.976948216949525 | 113.44597980000071 |
| 1706 | gse239651_dmso_day3::DMSO | 20260916.0 | sinkhorn_divergence | 0.0133843641894747 | 115.63007089999974 |
| 1707 | gse239651_dmso_day3::DMSO | 20260917.0 | sinkhorn_divergence | 0.0073169499488512 | 115.80890920000002 |
| 1708 | gse239651_dmso_day3::DMSO | 20260918.0 | sinkhorn_divergence | 0.0055338298552435 | 117.5844432999993 |
| 1709 | gse239651_dmso_day3::DMSO | 20260919.0 | sinkhorn_divergence | 0.0125342684506611 | 120.40731559999948 |
| 1710 | gse239651_dmso_day3::DMSO | 20260920.0 | sinkhorn_divergence | 0.0109625506650411 | 117.94565039999998 |
| 1723 | gse239651_dmso_day6::DMSO | 20260916.0 | sinkhorn_divergence | 0.0480520131433918 | 121.35117600000012 |
| 1724 | gse239651_dmso_day6::DMSO | 20260917.0 | sinkhorn_divergence | 0.0437239352823244 | 116.50757129999964 |
| 1725 | gse239651_dmso_day6::DMSO | 20260918.0 | sinkhorn_divergence | 0.0415113746514258 | 107.0824107999997 |
| 1726 | gse239651_dmso_day6::DMSO | 20260919.0 | sinkhorn_divergence | 0.0434164596500704 | 106.08582360000035 |
| 1727 | gse239651_dmso_day6::DMSO | 20260920.0 | sinkhorn_divergence | 0.0455950233058628 | 110.6642585999998 |
| 1740 | gse239651_dmso_day69::DMSO | 20260916.0 | sinkhorn_divergence | 0.0794703314092915 | 105.98453539999991 |
| 1741 | gse239651_dmso_day69::DMSO | 20260917.0 | sinkhorn_divergence | 0.0749670902847424 | 103.9811552000001 |
| 1742 | gse239651_dmso_day69::DMSO | 20260918.0 | sinkhorn_divergence | 0.0723898902514691 | 105.9265863999999 |
| 1743 | gse239651_dmso_day69::DMSO | 20260919.0 | sinkhorn_divergence | 0.074638090652598 | 101.50770660000036 |
| 1744 | gse239651_dmso_day69::DMSO | 20260920.0 | sinkhorn_divergence | 0.0766434869472568 | 104.91881490000014 |
| 1757 | gse239651_dmso_day98::DMSO | 20260916.0 | sinkhorn_divergence | 0.0395211441503247 | 122.23007440000038 |
| 1758 | gse239651_dmso_day98::DMSO | 20260917.0 | sinkhorn_divergence | 0.0343718009671043 | 133.0048740000002 |
| 1759 | gse239651_dmso_day98::DMSO | 20260918.0 | sinkhorn_divergence | 0.0340663667713531 | 133.1269975999994 |
| 1760 | gse239651_dmso_day98::DMSO | 20260919.0 | sinkhorn_divergence | 0.0393284965205327 | 133.16438820000076 |
| 1761 | gse239651_dmso_day98::DMSO | 20260920.0 | sinkhorn_divergence | 0.0365010158598626 | 134.2977045000007 |
| 1774 | gse239651_ispinesib_day3::ispinesib | 20260916.0 | sinkhorn_divergence | 0.0765983546210553 | 132.31578749999971 |
| 1775 | gse239651_ispinesib_day3::ispinesib | 20260917.0 | sinkhorn_divergence | 0.0727085874338132 | 130.98438579999856 |
| 1776 | gse239651_ispinesib_day3::ispinesib | 20260918.0 | sinkhorn_divergence | 0.0705535387627701 | 118.62514700000064 |
| 1777 | gse239651_ispinesib_day3::ispinesib | 20260919.0 | sinkhorn_divergence | 0.0745785059447318 | 111.87683459999973 |
| 1778 | gse239651_ispinesib_day3::ispinesib | 20260920.0 | sinkhorn_divergence | 0.0717339419772254 | 112.4667773000001 |
| 1791 | gse239651_ispinesib_day6::ispinesib | 20260916.0 | sinkhorn_divergence | 0.0797668048590401 | 115.93260999999984 |
| 1792 | gse239651_ispinesib_day6::ispinesib | 20260917.0 | sinkhorn_divergence | 0.0755305479914991 | 112.2782141000007 |
| 1793 | gse239651_ispinesib_day6::ispinesib | 20260918.0 | sinkhorn_divergence | 0.0731410218030623 | 110.78718130000016 |
| 1794 | gse239651_ispinesib_day6::ispinesib | 20260919.0 | sinkhorn_divergence | 0.075131142795574 | 112.84757480000007 |
| 1795 | gse239651_ispinesib_day6::ispinesib | 20260920.0 | sinkhorn_divergence | 0.0767511460326792 | 113.0381097999998 |
| 1808 | gse239651_ispinesib_day69::ispinesib | 20260916.0 | sinkhorn_divergence | 0.0737489034307971 | 113.43000900000152 |
| 1809 | gse239651_ispinesib_day69::ispinesib | 20260917.0 | sinkhorn_divergence | 0.0693213787322065 | 114.5526747999993 |
| 1810 | gse239651_ispinesib_day69::ispinesib | 20260918.0 | sinkhorn_divergence | 0.0668244705015984 | 116.72172610000052 |
| 1811 | gse239651_ispinesib_day69::ispinesib | 20260919.0 | sinkhorn_divergence | 0.068172468337015 | 112.28214730000036 |
| 1812 | gse239651_ispinesib_day69::ispinesib | 20260920.0 | sinkhorn_divergence | 0.0711422741318041 | 112.40618619999988 |
| 1825 | gse239651_ispinesib_day98::ispinesib | 20260916.0 | sinkhorn_divergence | 0.072231080085733 | 111.89460319999851 |
| 1826 | gse239651_ispinesib_day98::ispinesib | 20260917.0 | sinkhorn_divergence | 0.0678937898953271 | 109.15929640000104 |
| 1827 | gse239651_ispinesib_day98::ispinesib | 20260918.0 | sinkhorn_divergence | 0.0659319265777596 | 109.93124539999916 |
| 1828 | gse239651_ispinesib_day98::ispinesib | 20260919.0 | sinkhorn_divergence | 0.0720296256200508 | 112.23595419999948 |
| 1829 | gse239651_ispinesib_day98::ispinesib | 20260920.0 | sinkhorn_divergence | 0.0703773458988668 | 113.44597980000071 |
| 2012 | gse239651_dmso_day3::DMSO | 20260916.0 | top1_accuracy | 1.0 | 115.63007089999974 |
| 2013 | gse239651_dmso_day3::DMSO | 20260917.0 | top1_accuracy | 1.0 | 115.80890920000002 |
| 2014 | gse239651_dmso_day3::DMSO | 20260918.0 | top1_accuracy | 1.0 | 117.5844432999993 |
| 2015 | gse239651_dmso_day3::DMSO | 20260919.0 | top1_accuracy | 1.0 | 120.40731559999948 |
| 2016 | gse239651_dmso_day3::DMSO | 20260920.0 | top1_accuracy | 1.0 | 117.94565039999998 |
| 2029 | gse239651_dmso_day6::DMSO | 20260916.0 | top1_accuracy | 1.0 | 121.35117600000012 |
| 2030 | gse239651_dmso_day6::DMSO | 20260917.0 | top1_accuracy | 1.0 | 116.50757129999964 |
| 2031 | gse239651_dmso_day6::DMSO | 20260918.0 | top1_accuracy | 1.0 | 107.0824107999997 |
| 2032 | gse239651_dmso_day6::DMSO | 20260919.0 | top1_accuracy | 1.0 | 106.08582360000035 |
| 2033 | gse239651_dmso_day6::DMSO | 20260920.0 | top1_accuracy | 1.0 | 110.6642585999998 |
| 2046 | gse239651_dmso_day69::DMSO | 20260916.0 | top1_accuracy | 1.0 | 105.98453539999991 |
| 2047 | gse239651_dmso_day69::DMSO | 20260917.0 | top1_accuracy | 1.0 | 103.9811552000001 |
| 2048 | gse239651_dmso_day69::DMSO | 20260918.0 | top1_accuracy | 1.0 | 105.9265863999999 |
| 2049 | gse239651_dmso_day69::DMSO | 20260919.0 | top1_accuracy | 1.0 | 101.50770660000036 |
| 2050 | gse239651_dmso_day69::DMSO | 20260920.0 | top1_accuracy | 1.0 | 104.91881490000014 |
| 2063 | gse239651_dmso_day98::DMSO | 20260916.0 | top1_accuracy | 1.0 | 122.23007440000038 |
| 2064 | gse239651_dmso_day98::DMSO | 20260917.0 | top1_accuracy | 1.0 | 133.0048740000002 |
| 2065 | gse239651_dmso_day98::DMSO | 20260918.0 | top1_accuracy | 1.0 | 133.1269975999994 |
| 2066 | gse239651_dmso_day98::DMSO | 20260919.0 | top1_accuracy | 1.0 | 133.16438820000076 |
| 2067 | gse239651_dmso_day98::DMSO | 20260920.0 | top1_accuracy | 1.0 | 134.2977045000007 |
| 2080 | gse239651_ispinesib_day3::ispinesib | 20260916.0 | top1_accuracy | 1.0 | 132.31578749999971 |
| 2081 | gse239651_ispinesib_day3::ispinesib | 20260917.0 | top1_accuracy | 1.0 | 130.98438579999856 |
| 2082 | gse239651_ispinesib_day3::ispinesib | 20260918.0 | top1_accuracy | 1.0 | 118.62514700000064 |
| 2083 | gse239651_ispinesib_day3::ispinesib | 20260919.0 | top1_accuracy | 1.0 | 111.87683459999973 |
| 2084 | gse239651_ispinesib_day3::ispinesib | 20260920.0 | top1_accuracy | 1.0 | 112.4667773000001 |
| 2097 | gse239651_ispinesib_day6::ispinesib | 20260916.0 | top1_accuracy | 1.0 | 115.93260999999984 |
| 2098 | gse239651_ispinesib_day6::ispinesib | 20260917.0 | top1_accuracy | 1.0 | 112.2782141000007 |
| 2099 | gse239651_ispinesib_day6::ispinesib | 20260918.0 | top1_accuracy | 1.0 | 110.78718130000016 |
| 2100 | gse239651_ispinesib_day6::ispinesib | 20260919.0 | top1_accuracy | 1.0 | 112.84757480000007 |
| 2101 | gse239651_ispinesib_day6::ispinesib | 20260920.0 | top1_accuracy | 1.0 | 113.0381097999998 |
| 2114 | gse239651_ispinesib_day69::ispinesib | 20260916.0 | top1_accuracy | 1.0 | 113.43000900000152 |
| 2115 | gse239651_ispinesib_day69::ispinesib | 20260917.0 | top1_accuracy | 1.0 | 114.5526747999993 |
| 2116 | gse239651_ispinesib_day69::ispinesib | 20260918.0 | top1_accuracy | 1.0 | 116.72172610000052 |
| 2117 | gse239651_ispinesib_day69::ispinesib | 20260919.0 | top1_accuracy | 1.0 | 112.28214730000036 |
| 2118 | gse239651_ispinesib_day69::ispinesib | 20260920.0 | top1_accuracy | 1.0 | 112.40618619999988 |
| 2131 | gse239651_ispinesib_day98::ispinesib | 20260916.0 | top1_accuracy | 1.0 | 111.89460319999851 |
| 2132 | gse239651_ispinesib_day98::ispinesib | 20260917.0 | top1_accuracy | 1.0 | 109.15929640000104 |
| 2133 | gse239651_ispinesib_day98::ispinesib | 20260918.0 | top1_accuracy | 1.0 | 109.93124539999916 |
| 2134 | gse239651_ispinesib_day98::ispinesib | 20260919.0 | top1_accuracy | 1.0 | 112.23595419999948 |
| 2135 | gse239651_ispinesib_day98::ispinesib | 20260920.0 | top1_accuracy | 1.0 | 113.44597980000071 |
| 2318 | gse239651_dmso_day3::DMSO | 20260916.0 | multiclass_brier | 0.0009111063496398 | 115.63007089999974 |
| 2319 | gse239651_dmso_day3::DMSO | 20260917.0 | multiclass_brier | 0.0009260749834626 | 115.80890920000002 |
| 2320 | gse239651_dmso_day3::DMSO | 20260918.0 | multiclass_brier | 0.0008980462270904 | 117.5844432999993 |
| 2321 | gse239651_dmso_day3::DMSO | 20260919.0 | multiclass_brier | 0.000844339914709 | 120.40731559999948 |
| 2322 | gse239651_dmso_day3::DMSO | 20260920.0 | multiclass_brier | 0.0009432315189906 | 117.94565039999998 |
| 2335 | gse239651_dmso_day6::DMSO | 20260916.0 | multiclass_brier | 0.0531086120818408 | 121.35117600000012 |
| 2336 | gse239651_dmso_day6::DMSO | 20260917.0 | multiclass_brier | 0.0529785973105019 | 116.50757129999964 |
| 2337 | gse239651_dmso_day6::DMSO | 20260918.0 | multiclass_brier | 0.0529438574794169 | 107.0824107999997 |
| 2338 | gse239651_dmso_day6::DMSO | 20260919.0 | multiclass_brier | 0.0522938501118152 | 106.08582360000035 |
| 2339 | gse239651_dmso_day6::DMSO | 20260920.0 | multiclass_brier | 0.0535257944262386 | 110.6642585999998 |
| 2352 | gse239651_dmso_day69::DMSO | 20260916.0 | multiclass_brier | 0.2037178663332163 | 105.98453539999991 |
| 2353 | gse239651_dmso_day69::DMSO | 20260917.0 | multiclass_brier | 0.2022733186015859 | 103.9811552000001 |
| 2354 | gse239651_dmso_day69::DMSO | 20260918.0 | multiclass_brier | 0.2023411864973956 | 105.9265863999999 |
| 2355 | gse239651_dmso_day69::DMSO | 20260919.0 | multiclass_brier | 0.2012983182587824 | 101.50770660000036 |
| 2356 | gse239651_dmso_day69::DMSO | 20260920.0 | multiclass_brier | 0.2038049525123659 | 104.91881490000014 |
| 2369 | gse239651_dmso_day98::DMSO | 20260916.0 | multiclass_brier | 0.0353309034699373 | 122.23007440000038 |
| 2370 | gse239651_dmso_day98::DMSO | 20260917.0 | multiclass_brier | 0.0357923499122602 | 133.0048740000002 |
| 2371 | gse239651_dmso_day98::DMSO | 20260918.0 | multiclass_brier | 0.0360235961008311 | 133.1269975999994 |
| 2372 | gse239651_dmso_day98::DMSO | 20260919.0 | multiclass_brier | 0.0358262459225888 | 133.16438820000076 |
| 2373 | gse239651_dmso_day98::DMSO | 20260920.0 | multiclass_brier | 0.0362147925663268 | 134.2977045000007 |
| 2386 | gse239651_ispinesib_day3::ispinesib | 20260916.0 | multiclass_brier | 0.2283620194775748 | 132.31578749999971 |
| 2387 | gse239651_ispinesib_day3::ispinesib | 20260917.0 | multiclass_brier | 0.2273348367522623 | 130.98438579999856 |
| 2388 | gse239651_ispinesib_day3::ispinesib | 20260918.0 | multiclass_brier | 0.226674613483055 | 118.62514700000064 |
| 2389 | gse239651_ispinesib_day3::ispinesib | 20260919.0 | multiclass_brier | 0.2255436710835624 | 111.87683459999973 |
| 2390 | gse239651_ispinesib_day3::ispinesib | 20260920.0 | multiclass_brier | 0.2267839093546517 | 112.4667773000001 |
| 2403 | gse239651_ispinesib_day6::ispinesib | 20260916.0 | multiclass_brier | 0.182396281366485 | 115.93260999999984 |
| 2404 | gse239651_ispinesib_day6::ispinesib | 20260917.0 | multiclass_brier | 0.1809077940640159 | 112.2782141000007 |
| 2405 | gse239651_ispinesib_day6::ispinesib | 20260918.0 | multiclass_brier | 0.1813708677561772 | 110.78718130000016 |
| 2406 | gse239651_ispinesib_day6::ispinesib | 20260919.0 | multiclass_brier | 0.1807549237304095 | 112.84757480000007 |
| 2407 | gse239651_ispinesib_day6::ispinesib | 20260920.0 | multiclass_brier | 0.181845240028914 | 113.0381097999998 |
| 2420 | gse239651_ispinesib_day69::ispinesib | 20260916.0 | multiclass_brier | 0.1645149216598599 | 113.43000900000152 |
| 2421 | gse239651_ispinesib_day69::ispinesib | 20260917.0 | multiclass_brier | 0.1638850120326528 | 114.5526747999993 |
| 2422 | gse239651_ispinesib_day69::ispinesib | 20260918.0 | multiclass_brier | 0.1640311230983581 | 116.72172610000052 |
| 2423 | gse239651_ispinesib_day69::ispinesib | 20260919.0 | multiclass_brier | 0.1625920562310538 | 112.28214730000036 |
| 2424 | gse239651_ispinesib_day69::ispinesib | 20260920.0 | multiclass_brier | 0.1652601018868799 | 112.40618619999988 |
| 2437 | gse239651_ispinesib_day98::ispinesib | 20260916.0 | multiclass_brier | 0.1444986425106006 | 111.89460319999851 |
| 2438 | gse239651_ispinesib_day98::ispinesib | 20260917.0 | multiclass_brier | 0.1443123823243783 | 109.15929640000104 |
| 2439 | gse239651_ispinesib_day98::ispinesib | 20260918.0 | multiclass_brier | 0.1437487858262217 | 109.93124539999916 |
| 2440 | gse239651_ispinesib_day98::ispinesib | 20260919.0 | multiclass_brier | 0.1432918214418461 | 112.23595419999948 |
| 2441 | gse239651_ispinesib_day98::ispinesib | 20260920.0 | multiclass_brier | 0.1441465188108963 | 113.44597980000071 |

**dataset = GSE239651_expt2; benchmark = B3_prospective_composition; task = population_future_state_composition; split = expt1_to_locked_expt2; method = mioflow; method_version = see_method_status**

| **Record** | **repeat** | **seed** | **metric** | **value** | **runtime_seconds** |
| --- | --- | --- | --- | --- | --- |
| 1405 | gse239651_dmso_day3::DMSO | 20260916.0 | cross_entropy | 0.9962334672608402 | 14.598558699999558 |
| 1406 | gse239651_dmso_day3::DMSO | 20260917.0 | cross_entropy | 1.0050423152604069 | 13.951207300000531 |
| 1407 | gse239651_dmso_day3::DMSO | 20260918.0 | cross_entropy | 0.9906237654754012 | 12.65088199999991 |
| 1408 | gse239651_dmso_day3::DMSO | 20260919.0 | cross_entropy | 0.9550429379275278 | 13.814521600000262 |
| 1409 | gse239651_dmso_day3::DMSO | 20260920.0 | cross_entropy | 0.989576019497666 | 12.666965100000198 |
| 1422 | gse239651_dmso_day6::DMSO | 20260916.0 | cross_entropy | 0.9867712479737538 | 14.926148600000488 |
| 1423 | gse239651_dmso_day6::DMSO | 20260917.0 | cross_entropy | 0.9761577795254678 | 14.69445739999992 |
| 1424 | gse239651_dmso_day6::DMSO | 20260918.0 | cross_entropy | 0.9924476010080994 | 13.91017599999941 |
| 1425 | gse239651_dmso_day6::DMSO | 20260919.0 | cross_entropy | 0.968150597485614 | 14.022097099999884 |
| 1426 | gse239651_dmso_day6::DMSO | 20260920.0 | cross_entropy | 0.9999722640493784 | 13.102705799999969 |
| 1439 | gse239651_dmso_day69::DMSO | 20260916.0 | cross_entropy | 0.7872586211634485 | 14.52330400000028 |
| 1440 | gse239651_dmso_day69::DMSO | 20260917.0 | cross_entropy | 0.8394467595871631 | 14.84278539999923 |
| 1441 | gse239651_dmso_day69::DMSO | 20260918.0 | cross_entropy | 0.7918003016581237 | 13.503484500000011 |
| 1442 | gse239651_dmso_day69::DMSO | 20260919.0 | cross_entropy | 0.7843198586891725 | 14.792220300000736 |
| 1443 | gse239651_dmso_day69::DMSO | 20260920.0 | cross_entropy | 0.8123924618593176 | 13.586769199999251 |
| 1456 | gse239651_dmso_day98::DMSO | 20260916.0 | cross_entropy | 1.0224475921410483 | 13.158562699999491 |
| 1457 | gse239651_dmso_day98::DMSO | 20260917.0 | cross_entropy | 1.0295030076015743 | 14.319500599999628 |
| 1458 | gse239651_dmso_day98::DMSO | 20260918.0 | cross_entropy | 1.0223521663431203 | 13.963670500000262 |
| 1459 | gse239651_dmso_day98::DMSO | 20260919.0 | cross_entropy | 0.998135221842274 | 14.330297299999984 |
| 1460 | gse239651_dmso_day98::DMSO | 20260920.0 | cross_entropy | 1.0043230942894543 | 13.118254199999685 |
| 1473 | gse239651_ispinesib_day3::ispinesib | 20260916.0 | cross_entropy | 0.6135368584097628 | 12.298813100000189 |
| 1474 | gse239651_ispinesib_day3::ispinesib | 20260917.0 | cross_entropy | 0.6760299544141531 | 13.309447600000569 |
| 1475 | gse239651_ispinesib_day3::ispinesib | 20260918.0 | cross_entropy | 0.59428046462675 | 10.8384786000006 |
| 1476 | gse239651_ispinesib_day3::ispinesib | 20260919.0 | cross_entropy | 0.6356671528615768 | 12.781641300000048 |
| 1477 | gse239651_ispinesib_day3::ispinesib | 20260920.0 | cross_entropy | 0.7098456439461713 | 13.663610400000834 |
| 1490 | gse239651_ispinesib_day6::ispinesib | 20260916.0 | cross_entropy | 0.9127383824819372 | 11.874877599999309 |
| 1491 | gse239651_ispinesib_day6::ispinesib | 20260917.0 | cross_entropy | 0.8986029602813538 | 12.621181499998784 |
| 1492 | gse239651_ispinesib_day6::ispinesib | 20260918.0 | cross_entropy | 0.9141133660625064 | 12.361971999998786 |
| 1493 | gse239651_ispinesib_day6::ispinesib | 20260919.0 | cross_entropy | 0.8676519747342687 | 13.0319528 |
| 1494 | gse239651_ispinesib_day6::ispinesib | 20260920.0 | cross_entropy | 0.929472761588232 | 12.743641600000046 |
| 1507 | gse239651_ispinesib_day69::ispinesib | 20260916.0 | cross_entropy | 0.8197224200014208 | 13.13460490000034 |
| 1508 | gse239651_ispinesib_day69::ispinesib | 20260917.0 | cross_entropy | 0.8631950699125395 | 13.319859400000496 |
| 1509 | gse239651_ispinesib_day69::ispinesib | 20260918.0 | cross_entropy | 0.8473716259338324 | 13.1839473 |
| 1510 | gse239651_ispinesib_day69::ispinesib | 20260919.0 | cross_entropy | 0.8586908016994187 | 12.159537500001534 |
| 1511 | gse239651_ispinesib_day69::ispinesib | 20260920.0 | cross_entropy | 0.8993052003934544 | 12.749141899999811 |
| 1524 | gse239651_ispinesib_day98::ispinesib | 20260916.0 | cross_entropy | 0.8560450287219907 | 12.829244099999414 |
| 1525 | gse239651_ispinesib_day98::ispinesib | 20260917.0 | cross_entropy | 0.8557644344027052 | 11.73789880000004 |
| 1526 | gse239651_ispinesib_day98::ispinesib | 20260918.0 | cross_entropy | 0.8539252257084766 | 11.17984780000006 |
| 1527 | gse239651_ispinesib_day98::ispinesib | 20260919.0 | cross_entropy | 0.8156429348520877 | 12.00023929999952 |
| 1528 | gse239651_ispinesib_day98::ispinesib | 20260920.0 | cross_entropy | 0.8916816750049608 | 10.802229799999624 |
| 1711 | gse239651_dmso_day3::DMSO | 20260916.0 | sinkhorn_divergence | 0.0221656330970374 | 14.598558699999558 |
| 1712 | gse239651_dmso_day3::DMSO | 20260917.0 | sinkhorn_divergence | 0.0307861149677218 | 13.951207300000531 |
| 1713 | gse239651_dmso_day3::DMSO | 20260918.0 | sinkhorn_divergence | 0.0221783146710876 | 12.65088199999991 |
| 1714 | gse239651_dmso_day3::DMSO | 20260919.0 | sinkhorn_divergence | 0.0119957933934749 | 13.814521600000262 |
| 1715 | gse239651_dmso_day3::DMSO | 20260920.0 | sinkhorn_divergence | 0.0371476887226505 | 12.666965100000198 |
| 1728 | gse239651_dmso_day6::DMSO | 20260916.0 | sinkhorn_divergence | 0.0294825697651997 | 14.926148600000488 |
| 1729 | gse239651_dmso_day6::DMSO | 20260917.0 | sinkhorn_divergence | 0.0231087803502442 | 14.69445739999992 |
| 1730 | gse239651_dmso_day6::DMSO | 20260918.0 | sinkhorn_divergence | 0.0278741835829767 | 13.91017599999941 |
| 1731 | gse239651_dmso_day6::DMSO | 20260919.0 | sinkhorn_divergence | 0.0125342033566239 | 14.022097099999884 |
| 1732 | gse239651_dmso_day6::DMSO | 20260920.0 | sinkhorn_divergence | 0.0470771822545301 | 13.102705799999969 |
| 1745 | gse239651_dmso_day69::DMSO | 20260916.0 | sinkhorn_divergence | 0.0907233147857463 | 14.52330400000028 |
| 1746 | gse239651_dmso_day69::DMSO | 20260917.0 | sinkhorn_divergence | 0.0967490948339089 | 14.84278539999923 |
| 1747 | gse239651_dmso_day69::DMSO | 20260918.0 | sinkhorn_divergence | 0.0911644217815995 | 13.503484500000011 |
| 1748 | gse239651_dmso_day69::DMSO | 20260919.0 | sinkhorn_divergence | 0.0678850865180176 | 14.792220300000736 |
| 1749 | gse239651_dmso_day69::DMSO | 20260920.0 | sinkhorn_divergence | 0.084435427565002 | 13.586769199999251 |
| 1762 | gse239651_dmso_day98::DMSO | 20260916.0 | sinkhorn_divergence | 0.065388434698078 | 13.158562699999491 |
| 1763 | gse239651_dmso_day98::DMSO | 20260917.0 | sinkhorn_divergence | 0.0638156009877749 | 14.319500599999628 |
| 1764 | gse239651_dmso_day98::DMSO | 20260918.0 | sinkhorn_divergence | 0.0507339761611228 | 13.963670500000262 |
| 1765 | gse239651_dmso_day98::DMSO | 20260919.0 | sinkhorn_divergence | 0.0453966291815568 | 14.330297299999984 |
| 1766 | gse239651_dmso_day98::DMSO | 20260920.0 | sinkhorn_divergence | 0.057461030769125 | 13.118254199999685 |
| 1779 | gse239651_ispinesib_day3::ispinesib | 20260916.0 | sinkhorn_divergence | 0.0577872033080676 | 12.298813100000189 |
| 1780 | gse239651_ispinesib_day3::ispinesib | 20260917.0 | sinkhorn_divergence | 0.0580318663088776 | 13.309447600000569 |
| 1781 | gse239651_ispinesib_day3::ispinesib | 20260918.0 | sinkhorn_divergence | 0.0310650785804466 | 10.8384786000006 |
| 1782 | gse239651_ispinesib_day3::ispinesib | 20260919.0 | sinkhorn_divergence | 0.0340272649837307 | 12.781641300000048 |
| 1783 | gse239651_ispinesib_day3::ispinesib | 20260920.0 | sinkhorn_divergence | 0.0660910687459542 | 13.663610400000834 |
| 1796 | gse239651_ispinesib_day6::ispinesib | 20260916.0 | sinkhorn_divergence | 0.0828642046126163 | 11.874877599999309 |
| 1797 | gse239651_ispinesib_day6::ispinesib | 20260917.0 | sinkhorn_divergence | 0.0534203593088263 | 12.621181499998784 |
| 1798 | gse239651_ispinesib_day6::ispinesib | 20260918.0 | sinkhorn_divergence | 0.0627562619767706 | 12.361971999998786 |
| 1799 | gse239651_ispinesib_day6::ispinesib | 20260919.0 | sinkhorn_divergence | 0.0361933870431288 | 13.0319528 |
| 1800 | gse239651_ispinesib_day6::ispinesib | 20260920.0 | sinkhorn_divergence | 0.0585354150437482 | 12.743641600000046 |
| 1813 | gse239651_ispinesib_day69::ispinesib | 20260916.0 | sinkhorn_divergence | 0.0321955795492395 | 13.13460490000034 |
| 1814 | gse239651_ispinesib_day69::ispinesib | 20260917.0 | sinkhorn_divergence | 0.0533456031221731 | 13.319859400000496 |
| 1815 | gse239651_ispinesib_day69::ispinesib | 20260918.0 | sinkhorn_divergence | 0.0403382038133384 | 13.1839473 |
| 1816 | gse239651_ispinesib_day69::ispinesib | 20260919.0 | sinkhorn_divergence | 0.0396773714203108 | 12.159537500001534 |
| 1817 | gse239651_ispinesib_day69::ispinesib | 20260920.0 | sinkhorn_divergence | 0.0677769474137668 | 12.749141899999811 |
| 1830 | gse239651_ispinesib_day98::ispinesib | 20260916.0 | sinkhorn_divergence | 0.0515636024423813 | 12.829244099999414 |
| 1831 | gse239651_ispinesib_day98::ispinesib | 20260917.0 | sinkhorn_divergence | 0.0487125327841218 | 11.73789880000004 |
| 1832 | gse239651_ispinesib_day98::ispinesib | 20260918.0 | sinkhorn_divergence | 0.0448081893918791 | 11.17984780000006 |
| 1833 | gse239651_ispinesib_day98::ispinesib | 20260919.0 | sinkhorn_divergence | 0.0263658895497084 | 12.00023929999952 |
| 1834 | gse239651_ispinesib_day98::ispinesib | 20260920.0 | sinkhorn_divergence | 0.0628905096442678 | 10.802229799999624 |
| 2017 | gse239651_dmso_day3::DMSO | 20260916.0 | top1_accuracy | 1.0 | 14.598558699999558 |
| 2018 | gse239651_dmso_day3::DMSO | 20260917.0 | top1_accuracy | 1.0 | 13.951207300000531 |
| 2019 | gse239651_dmso_day3::DMSO | 20260918.0 | top1_accuracy | 1.0 | 12.65088199999991 |
| 2020 | gse239651_dmso_day3::DMSO | 20260919.0 | top1_accuracy | 1.0 | 13.814521600000262 |
| 2021 | gse239651_dmso_day3::DMSO | 20260920.0 | top1_accuracy | 1.0 | 12.666965100000198 |
| 2034 | gse239651_dmso_day6::DMSO | 20260916.0 | top1_accuracy | 1.0 | 14.926148600000488 |
| 2035 | gse239651_dmso_day6::DMSO | 20260917.0 | top1_accuracy | 1.0 | 14.69445739999992 |
| 2036 | gse239651_dmso_day6::DMSO | 20260918.0 | top1_accuracy | 1.0 | 13.91017599999941 |
| 2037 | gse239651_dmso_day6::DMSO | 20260919.0 | top1_accuracy | 1.0 | 14.022097099999884 |
| 2038 | gse239651_dmso_day6::DMSO | 20260920.0 | top1_accuracy | 1.0 | 13.102705799999969 |
| 2051 | gse239651_dmso_day69::DMSO | 20260916.0 | top1_accuracy | 1.0 | 14.52330400000028 |
| 2052 | gse239651_dmso_day69::DMSO | 20260917.0 | top1_accuracy | 1.0 | 14.84278539999923 |
| 2053 | gse239651_dmso_day69::DMSO | 20260918.0 | top1_accuracy | 1.0 | 13.503484500000011 |
| 2054 | gse239651_dmso_day69::DMSO | 20260919.0 | top1_accuracy | 1.0 | 14.792220300000736 |
| 2055 | gse239651_dmso_day69::DMSO | 20260920.0 | top1_accuracy | 1.0 | 13.586769199999251 |
| 2068 | gse239651_dmso_day98::DMSO | 20260916.0 | top1_accuracy | 1.0 | 13.158562699999491 |
| 2069 | gse239651_dmso_day98::DMSO | 20260917.0 | top1_accuracy | 1.0 | 14.319500599999628 |
| 2070 | gse239651_dmso_day98::DMSO | 20260918.0 | top1_accuracy | 1.0 | 13.963670500000262 |
| 2071 | gse239651_dmso_day98::DMSO | 20260919.0 | top1_accuracy | 1.0 | 14.330297299999984 |
| 2072 | gse239651_dmso_day98::DMSO | 20260920.0 | top1_accuracy | 1.0 | 13.118254199999685 |
| 2085 | gse239651_ispinesib_day3::ispinesib | 20260916.0 | top1_accuracy | 1.0 | 12.298813100000189 |
| 2086 | gse239651_ispinesib_day3::ispinesib | 20260917.0 | top1_accuracy | 1.0 | 13.309447600000569 |
| 2087 | gse239651_ispinesib_day3::ispinesib | 20260918.0 | top1_accuracy | 1.0 | 10.8384786000006 |
| 2088 | gse239651_ispinesib_day3::ispinesib | 20260919.0 | top1_accuracy | 1.0 | 12.781641300000048 |
| 2089 | gse239651_ispinesib_day3::ispinesib | 20260920.0 | top1_accuracy | 1.0 | 13.663610400000834 |
| 2102 | gse239651_ispinesib_day6::ispinesib | 20260916.0 | top1_accuracy | 1.0 | 11.874877599999309 |
| 2103 | gse239651_ispinesib_day6::ispinesib | 20260917.0 | top1_accuracy | 1.0 | 12.621181499998784 |
| 2104 | gse239651_ispinesib_day6::ispinesib | 20260918.0 | top1_accuracy | 1.0 | 12.361971999998786 |
| 2105 | gse239651_ispinesib_day6::ispinesib | 20260919.0 | top1_accuracy | 1.0 | 13.0319528 |
| 2106 | gse239651_ispinesib_day6::ispinesib | 20260920.0 | top1_accuracy | 1.0 | 12.743641600000046 |
| 2119 | gse239651_ispinesib_day69::ispinesib | 20260916.0 | top1_accuracy | 1.0 | 13.13460490000034 |
| 2120 | gse239651_ispinesib_day69::ispinesib | 20260917.0 | top1_accuracy | 1.0 | 13.319859400000496 |
| 2121 | gse239651_ispinesib_day69::ispinesib | 20260918.0 | top1_accuracy | 1.0 | 13.1839473 |
| 2122 | gse239651_ispinesib_day69::ispinesib | 20260919.0 | top1_accuracy | 1.0 | 12.159537500001534 |
| 2123 | gse239651_ispinesib_day69::ispinesib | 20260920.0 | top1_accuracy | 1.0 | 12.749141899999811 |
| 2136 | gse239651_ispinesib_day98::ispinesib | 20260916.0 | top1_accuracy | 1.0 | 12.829244099999414 |
| 2137 | gse239651_ispinesib_day98::ispinesib | 20260917.0 | top1_accuracy | 1.0 | 11.73789880000004 |
| 2138 | gse239651_ispinesib_day98::ispinesib | 20260918.0 | top1_accuracy | 1.0 | 11.17984780000006 |
| 2139 | gse239651_ispinesib_day98::ispinesib | 20260919.0 | top1_accuracy | 1.0 | 12.00023929999952 |
| 2140 | gse239651_ispinesib_day98::ispinesib | 20260920.0 | top1_accuracy | 1.0 | 10.802229799999624 |
| 2323 | gse239651_dmso_day3::DMSO | 20260916.0 | multiclass_brier | 0.0035618650299663 | 14.598558699999558 |
| 2324 | gse239651_dmso_day3::DMSO | 20260917.0 | multiclass_brier | 0.0036466455617465 | 13.951207300000531 |
| 2325 | gse239651_dmso_day3::DMSO | 20260918.0 | multiclass_brier | 0.0030228590675321 | 12.65088199999991 |
| 2326 | gse239651_dmso_day3::DMSO | 20260919.0 | multiclass_brier | 0.0028224297216845 | 13.814521600000262 |
| 2327 | gse239651_dmso_day3::DMSO | 20260920.0 | multiclass_brier | 0.0026561374181277 | 12.666965100000198 |
| 2340 | gse239651_dmso_day6::DMSO | 20260916.0 | multiclass_brier | 0.0096100424902748 | 14.926148600000488 |
| 2341 | gse239651_dmso_day6::DMSO | 20260917.0 | multiclass_brier | 0.0018909468706494 | 14.69445739999992 |
| 2342 | gse239651_dmso_day6::DMSO | 20260918.0 | multiclass_brier | 0.0070663822641806 | 13.91017599999941 |
| 2343 | gse239651_dmso_day6::DMSO | 20260919.0 | multiclass_brier | 0.0026068460292203 | 14.022097099999884 |
| 2344 | gse239651_dmso_day6::DMSO | 20260920.0 | multiclass_brier | 0.0073075947151736 | 13.102705799999969 |
| 2357 | gse239651_dmso_day69::DMSO | 20260916.0 | multiclass_brier | 0.0524870644686214 | 14.52330400000028 |
| 2358 | gse239651_dmso_day69::DMSO | 20260917.0 | multiclass_brier | 0.1043293802992651 | 14.84278539999923 |
| 2359 | gse239651_dmso_day69::DMSO | 20260918.0 | multiclass_brier | 0.0621905161892407 | 13.503484500000011 |
| 2360 | gse239651_dmso_day69::DMSO | 20260919.0 | multiclass_brier | 0.0783023425354155 | 14.792220300000736 |
| 2361 | gse239651_dmso_day69::DMSO | 20260920.0 | multiclass_brier | 0.0921960856769879 | 13.586769199999251 |
| 2374 | gse239651_dmso_day98::DMSO | 20260916.0 | multiclass_brier | 0.0069152953159737 | 13.158562699999491 |
| 2375 | gse239651_dmso_day98::DMSO | 20260917.0 | multiclass_brier | 0.0117659649590919 | 14.319500599999628 |
| 2376 | gse239651_dmso_day98::DMSO | 20260918.0 | multiclass_brier | 0.0074036866807733 | 13.963670500000262 |
| 2377 | gse239651_dmso_day98::DMSO | 20260919.0 | multiclass_brier | 0.012077768709125 | 14.330297299999984 |
| 2378 | gse239651_dmso_day98::DMSO | 20260920.0 | multiclass_brier | 0.0121024439555332 | 13.118254199999685 |
| 2391 | gse239651_ispinesib_day3::ispinesib | 20260916.0 | multiclass_brier | 0.0075947681013464 | 12.298813100000189 |
| 2392 | gse239651_ispinesib_day3::ispinesib | 20260917.0 | multiclass_brier | 0.0701236964656074 | 13.309447600000569 |
| 2393 | gse239651_ispinesib_day3::ispinesib | 20260918.0 | multiclass_brier | 0.0082475853419296 | 10.8384786000006 |
| 2394 | gse239651_ispinesib_day3::ispinesib | 20260919.0 | multiclass_brier | 0.0476370069849326 | 12.781641300000048 |
| 2395 | gse239651_ispinesib_day3::ispinesib | 20260920.0 | multiclass_brier | 0.096945366375192 | 13.663610400000834 |
| 2408 | gse239651_ispinesib_day6::ispinesib | 20260916.0 | multiclass_brier | 0.0091723678281701 | 11.874877599999309 |
| 2409 | gse239651_ispinesib_day6::ispinesib | 20260917.0 | multiclass_brier | 0.0497540484850227 | 12.621181499998784 |
| 2410 | gse239651_ispinesib_day6::ispinesib | 20260918.0 | multiclass_brier | 0.0090627826090539 | 12.361971999998786 |
| 2411 | gse239651_ispinesib_day6::ispinesib | 20260919.0 | multiclass_brier | 0.0258024275076997 | 13.0319528 |
| 2412 | gse239651_ispinesib_day6::ispinesib | 20260920.0 | multiclass_brier | 0.0730777422116582 | 12.743641600000046 |
| 2425 | gse239651_ispinesib_day69::ispinesib | 20260916.0 | multiclass_brier | 0.0169674009205449 | 13.13460490000034 |
| 2426 | gse239651_ispinesib_day69::ispinesib | 20260917.0 | multiclass_brier | 0.0472351380357399 | 13.319859400000496 |
| 2427 | gse239651_ispinesib_day69::ispinesib | 20260918.0 | multiclass_brier | 0.025625616000506 | 13.1839473 |
| 2428 | gse239651_ispinesib_day69::ispinesib | 20260919.0 | multiclass_brier | 0.0488561383080453 | 12.159537500001534 |
| 2429 | gse239651_ispinesib_day69::ispinesib | 20260920.0 | multiclass_brier | 0.0712923154599093 | 12.749141899999811 |
| 2442 | gse239651_ispinesib_day98::ispinesib | 20260916.0 | multiclass_brier | 0.0030622820592636 | 12.829244099999414 |
| 2443 | gse239651_ispinesib_day98::ispinesib | 20260917.0 | multiclass_brier | 0.0348994289724153 | 11.73789880000004 |
| 2444 | gse239651_ispinesib_day98::ispinesib | 20260918.0 | multiclass_brier | 0.0023426782198596 | 11.17984780000006 |
| 2445 | gse239651_ispinesib_day98::ispinesib | 20260919.0 | multiclass_brier | 0.011081090837405 | 12.00023929999952 |
| 2446 | gse239651_ispinesib_day98::ispinesib | 20260920.0 | multiclass_brier | 0.0517477243466723 | 10.802229799999624 |

**dataset = GSE140802; benchmark = B3_prospective_composition; task = population_future_state_composition; split = held_out_animal; method = source-carry-forward; method_version = see_method_status**

| **Record** | **repeat** | **seed** | **metric** | **value** | **runtime_seconds** |
| --- | --- | --- | --- | --- | --- |
| 1529 | gse140802_t2_t16_holdout_A4::heldout_animal_A4 | 20260916.0 | cross_entropy | 26.2988111692677 | 0.1700598000006721 |
| 1546 | gse140802_t2_t16_holdout_A5::heldout_animal_A5 | 20260916.0 | cross_entropy | 24.15077799827343 | 0.2426227999985712 |
| 1563 | gse140802_t2_t16_holdout_B4::heldout_animal_B4 | 20260916.0 | cross_entropy | 26.05425088920436 | 0.2292116999997233 |
| 1580 | gse140802_t2_t16_holdout_B5::heldout_animal_B5 | 20260916.0 | cross_entropy | 24.476656466031105 | 0.2221160999997664 |
| 1597 | gse140802_t2_t9_holdout_A1::heldout_animal_A1 | 20260916.0 | cross_entropy | 21.58347687168758 | 0.2186065000005328 |
| 1614 | gse140802_t2_t9_holdout_A2::heldout_animal_A2 | 20260916.0 | cross_entropy | 21.74428980784592 | 0.2386432999992394 |
| 1631 | gse140802_t2_t9_holdout_A3::heldout_animal_A3 | 20260916.0 | cross_entropy | 23.75618628983284 | 0.2116293000017322 |
| 1648 | gse140802_t2_t9_holdout_B1::heldout_animal_B1 | 20260916.0 | cross_entropy | 23.51060568636025 | 0.2309928000013315 |
| 1665 | gse140802_t2_t9_holdout_B2::heldout_animal_B2 | 20260916.0 | cross_entropy | 23.961276123969288 | 0.2897888000006787 |
| 1682 | gse140802_t2_t9_holdout_B3::heldout_animal_B3 | 20260916.0 | cross_entropy | 19.62202948812317 | 0.2153218000003107 |
| 1835 | gse140802_t2_t16_holdout_A4::heldout_animal_A4 | 20260916.0 | sinkhorn_divergence | 1.903571428571428 | 0.1700598000006721 |
| 1852 | gse140802_t2_t16_holdout_A5::heldout_animal_A5 | 20260916.0 | sinkhorn_divergence | 1.7480916030534353 | 0.2426227999985712 |
| 1869 | gse140802_t2_t16_holdout_B4::heldout_animal_B4 | 20260916.0 | sinkhorn_divergence | 1.885869565217391 | 0.2292116999997233 |
| 1886 | gse140802_t2_t16_holdout_B5::heldout_animal_B5 | 20260916.0 | sinkhorn_divergence | 1.7716794731064769 | 0.2221160999997664 |
| 1903 | gse140802_t2_t9_holdout_A1::heldout_animal_A1 | 20260916.0 | sinkhorn_divergence | 1.562264150943396 | 0.2186065000005328 |
| 1920 | gse140802_t2_t9_holdout_A2::heldout_animal_A2 | 20260916.0 | sinkhorn_divergence | 1.573904179408767 | 0.2386432999992394 |
| 1937 | gse140802_t2_t9_holdout_A3::heldout_animal_A3 | 20260916.0 | sinkhorn_divergence | 1.719530102790015 | 0.2116293000017322 |
| 1954 | gse140802_t2_t9_holdout_B1::heldout_animal_B1 | 20260916.0 | sinkhorn_divergence | 1.7017543859649122 | 0.2309928000013315 |
| 1971 | gse140802_t2_t9_holdout_B2::heldout_animal_B2 | 20260916.0 | sinkhorn_divergence | 1.734375 | 0.2897888000006787 |
| 1988 | gse140802_t2_t9_holdout_B3::heldout_animal_B3 | 20260916.0 | sinkhorn_divergence | 1.420289855072464 | 0.2153218000003107 |
| 2141 | gse140802_t2_t16_holdout_A4::heldout_animal_A4 | 20260916.0 | top1_accuracy | 0.0 | 0.1700598000006721 |
| 2158 | gse140802_t2_t16_holdout_A5::heldout_animal_A5 | 20260916.0 | top1_accuracy | 0.0 | 0.2426227999985712 |
| 2175 | gse140802_t2_t16_holdout_B4::heldout_animal_B4 | 20260916.0 | top1_accuracy | 0.0 | 0.2292116999997233 |
| 2192 | gse140802_t2_t16_holdout_B5::heldout_animal_B5 | 20260916.0 | top1_accuracy | 0.0 | 0.2221160999997664 |
| 2209 | gse140802_t2_t9_holdout_A1::heldout_animal_A1 | 20260916.0 | top1_accuracy | 0.0 | 0.2186065000005328 |
| 2226 | gse140802_t2_t9_holdout_A2::heldout_animal_A2 | 20260916.0 | top1_accuracy | 0.0 | 0.2386432999992394 |
| 2243 | gse140802_t2_t9_holdout_A3::heldout_animal_A3 | 20260916.0 | top1_accuracy | 0.0 | 0.2116293000017322 |
| 2260 | gse140802_t2_t9_holdout_B1::heldout_animal_B1 | 20260916.0 | top1_accuracy | 0.0 | 0.2309928000013315 |
| 2277 | gse140802_t2_t9_holdout_B2::heldout_animal_B2 | 20260916.0 | top1_accuracy | 0.0 | 0.2897888000006787 |
| 2294 | gse140802_t2_t9_holdout_B3::heldout_animal_B3 | 20260916.0 | top1_accuracy | 1.0 | 0.2153218000003107 |
| 2447 | gse140802_t2_t16_holdout_A4::heldout_animal_A4 | 20260916.0 | multiclass_brier | 1.2991581632653062 | 0.1700598000006721 |
| 2464 | gse140802_t2_t16_holdout_A5::heldout_animal_A5 | 20260916.0 | multiclass_brier | 0.971213798729678 | 0.2426227999985712 |
| 2481 | gse140802_t2_t16_holdout_B4::heldout_animal_B4 | 20260916.0 | multiclass_brier | 1.2567934782608692 | 0.2292116999997233 |
| 2498 | gse140802_t2_t16_holdout_B5::heldout_animal_B5 | 20260916.0 | multiclass_brier | 0.9732902288290092 | 0.2221160999997664 |
| 2515 | gse140802_t2_t9_holdout_A1::heldout_animal_A1 | 20260916.0 | multiclass_brier | 0.7573086507653971 | 0.2186065000005328 |
| 2532 | gse140802_t2_t9_holdout_A2::heldout_animal_A2 | 20260916.0 | multiclass_brier | 0.8012738116638835 | 0.2386432999992394 |
| 2549 | gse140802_t2_t9_holdout_A3::heldout_animal_A3 | 20260916.0 | multiclass_brier | 1.0066273360631877 | 0.2116293000017322 |
| 2566 | gse140802_t2_t9_holdout_B1::heldout_animal_B1 | 20260916.0 | multiclass_brier | 0.9085872576177284 | 0.2309928000013315 |
| 2583 | gse140802_t2_t9_holdout_B2::heldout_animal_B2 | 20260916.0 | multiclass_brier | 0.955283203125 | 0.2897888000006787 |
| 2600 | gse140802_t2_t9_holdout_B3::heldout_animal_B3 | 20260916.0 | multiclass_brier | 0.6145767695862214 | 0.2153218000003107 |

**dataset = GSE140802; benchmark = B3_prospective_composition; task = population_future_state_composition; split = held_out_animal; method = development-target-transfer; method_version = see_method_status**

| **Record** | **repeat** | **seed** | **metric** | **value** | **runtime_seconds** |
| --- | --- | --- | --- | --- | --- |
| 1530 | gse140802_t2_t16_holdout_A4::heldout_animal_A4 | 20260916.0 | cross_entropy | 1.3406626155650965 | 0.1493849999988015 |
| 1547 | gse140802_t2_t16_holdout_A5::heldout_animal_A5 | 20260916.0 | cross_entropy | 1.7254243067129305 | 0.1738934000004519 |
| 1564 | gse140802_t2_t16_holdout_B4::heldout_animal_B4 | 20260916.0 | cross_entropy | 1.3659068866352768 | 0.1878577000006771 |
| 1581 | gse140802_t2_t16_holdout_B5::heldout_animal_B5 | 20260916.0 | cross_entropy | 1.8958968729219468 | 0.1606494000006932 |
| 1598 | gse140802_t2_t9_holdout_A1::heldout_animal_A1 | 20260916.0 | cross_entropy | 1.8187451107316104 | 0.1584171000013157 |
| 1615 | gse140802_t2_t9_holdout_A2::heldout_animal_A2 | 20260916.0 | cross_entropy | 1.673093262434782 | 0.1566253999990294 |
| 1632 | gse140802_t2_t9_holdout_A3::heldout_animal_A3 | 20260916.0 | cross_entropy | 1.5982215943035776 | 0.1445597000001726 |
| 1649 | gse140802_t2_t9_holdout_B1::heldout_animal_B1 | 20260916.0 | cross_entropy | 1.7659453620562098 | 0.1843263000009756 |
| 1666 | gse140802_t2_t9_holdout_B2::heldout_animal_B2 | 20260916.0 | cross_entropy | 1.7238366810966512 | 0.1819457000001421 |
| 1683 | gse140802_t2_t9_holdout_B3::heldout_animal_B3 | 20260916.0 | cross_entropy | 1.8715845149667325 | 0.1918287999997119 |
| 1836 | gse140802_t2_t16_holdout_A4::heldout_animal_A4 | 20260916.0 | sinkhorn_divergence | 0.4709372392694909 | 0.1493849999988015 |
| 1853 | gse140802_t2_t16_holdout_A5::heldout_animal_A5 | 20260916.0 | sinkhorn_divergence | 0.2903516606513632 | 0.1738934000004519 |
| 1870 | gse140802_t2_t16_holdout_B4::heldout_animal_B4 | 20260916.0 | sinkhorn_divergence | 0.3516677538322587 | 0.1878577000006771 |
| 1887 | gse140802_t2_t16_holdout_B5::heldout_animal_B5 | 20260916.0 | sinkhorn_divergence | 0.4671475615493178 | 0.1606494000006932 |
| 1904 | gse140802_t2_t9_holdout_A1::heldout_animal_A1 | 20260916.0 | sinkhorn_divergence | 0.4152358048487207 | 0.1584171000013157 |
| 1921 | gse140802_t2_t9_holdout_A2::heldout_animal_A2 | 20260916.0 | sinkhorn_divergence | 0.2669841273495116 | 0.1566253999990294 |
| 1938 | gse140802_t2_t9_holdout_A3::heldout_animal_A3 | 20260916.0 | sinkhorn_divergence | 0.3293542308974324 | 0.1445597000001726 |
| 1955 | gse140802_t2_t9_holdout_B1::heldout_animal_B1 | 20260916.0 | sinkhorn_divergence | 0.3596459248741613 | 0.1843263000009756 |
| 1972 | gse140802_t2_t9_holdout_B2::heldout_animal_B2 | 20260916.0 | sinkhorn_divergence | 0.1482862903222953 | 0.1819457000001421 |
| 1989 | gse140802_t2_t9_holdout_B3::heldout_animal_B3 | 20260916.0 | sinkhorn_divergence | 0.4361164646827603 | 0.1918287999997119 |
| 2142 | gse140802_t2_t16_holdout_A4::heldout_animal_A4 | 20260916.0 | top1_accuracy | 1.0 | 0.1493849999988015 |
| 2159 | gse140802_t2_t16_holdout_A5::heldout_animal_A5 | 20260916.0 | top1_accuracy | 0.0 | 0.1738934000004519 |
| 2176 | gse140802_t2_t16_holdout_B4::heldout_animal_B4 | 20260916.0 | top1_accuracy | 1.0 | 0.1878577000006771 |
| 2193 | gse140802_t2_t16_holdout_B5::heldout_animal_B5 | 20260916.0 | top1_accuracy | 1.0 | 0.1606494000006932 |
| 2210 | gse140802_t2_t9_holdout_A1::heldout_animal_A1 | 20260916.0 | top1_accuracy | 0.0 | 0.1584171000013157 |
| 2227 | gse140802_t2_t9_holdout_A2::heldout_animal_A2 | 20260916.0 | top1_accuracy | 1.0 | 0.1566253999990294 |
| 2244 | gse140802_t2_t9_holdout_A3::heldout_animal_A3 | 20260916.0 | top1_accuracy | 1.0 | 0.1445597000001726 |
| 2261 | gse140802_t2_t9_holdout_B1::heldout_animal_B1 | 20260916.0 | top1_accuracy | 1.0 | 0.1843263000009756 |
| 2278 | gse140802_t2_t9_holdout_B2::heldout_animal_B2 | 20260916.0 | top1_accuracy | 1.0 | 0.1819457000001421 |
| 2295 | gse140802_t2_t9_holdout_B3::heldout_animal_B3 | 20260916.0 | top1_accuracy | 0.0 | 0.1918287999997119 |
| 2448 | gse140802_t2_t16_holdout_A4::heldout_animal_A4 | 20260916.0 | multiclass_brier | 0.0547309358368039 | 0.1493849999988015 |
| 2465 | gse140802_t2_t16_holdout_A5::heldout_animal_A5 | 20260916.0 | multiclass_brier | 0.0266684416305751 | 0.1738934000004519 |
| 2482 | gse140802_t2_t16_holdout_B4::heldout_animal_B4 | 20260916.0 | multiclass_brier | 0.0373651625912385 | 0.1878577000006771 |
| 2499 | gse140802_t2_t16_holdout_B5::heldout_animal_B5 | 20260916.0 | multiclass_brier | 0.0543303080620969 | 0.1606494000006932 |
| 2516 | gse140802_t2_t9_holdout_A1::heldout_animal_A1 | 20260916.0 | multiclass_brier | 0.0440149099072515 | 0.1584171000013157 |
| 2533 | gse140802_t2_t9_holdout_A2::heldout_animal_A2 | 20260916.0 | multiclass_brier | 0.0141366501090766 | 0.1566253999990294 |
| 2550 | gse140802_t2_t9_holdout_A3::heldout_animal_A3 | 20260916.0 | multiclass_brier | 0.0320561630397554 | 0.1445597000001726 |
| 2567 | gse140802_t2_t9_holdout_B1::heldout_animal_B1 | 20260916.0 | multiclass_brier | 0.0255850698236066 | 0.1843263000009756 |
| 2584 | gse140802_t2_t9_holdout_B2::heldout_animal_B2 | 20260916.0 | multiclass_brier | 0.0054948560256243 | 0.1819457000001421 |
| 2601 | gse140802_t2_t9_holdout_B3::heldout_animal_B3 | 20260916.0 | multiclass_brier | 0.0441248100767381 | 0.1918287999997119 |

**dataset = GSE140802; benchmark = B3_prospective_composition; task = population_future_state_composition; split = held_out_animal; method = prescient; method_version = see_method_status**

| **Record** | **repeat** | **seed** | **metric** | **value** | **runtime_seconds** |
| --- | --- | --- | --- | --- | --- |
| 1531 | gse140802_t2_t16_holdout_A4::heldout_animal_A4 | 20260916.0 | cross_entropy | 1.316015191179251 | 4.053417700000864 |
| 1532 | gse140802_t2_t16_holdout_A4::heldout_animal_A4 | 20260917.0 | cross_entropy | 1.2911359334589805 | 3.204100699998889 |
| 1533 | gse140802_t2_t16_holdout_A4::heldout_animal_A4 | 20260918.0 | cross_entropy | 1.301474474685988 | 3.3533478999997897 |
| 1534 | gse140802_t2_t16_holdout_A4::heldout_animal_A4 | 20260919.0 | cross_entropy | 1.3044740913172468 | 3.1164227000008395 |
| 1535 | gse140802_t2_t16_holdout_A4::heldout_animal_A4 | 20260920.0 | cross_entropy | 1.2653430367385092 | 3.0896375000011176 |
| 1548 | gse140802_t2_t16_holdout_A5::heldout_animal_A5 | 20260916.0 | cross_entropy | 1.705692479022567 | 4.5700616000012815 |
| 1549 | gse140802_t2_t16_holdout_A5::heldout_animal_A5 | 20260917.0 | cross_entropy | 1.7235413237278043 | 3.3212940000012168 |
| 1550 | gse140802_t2_t16_holdout_A5::heldout_animal_A5 | 20260918.0 | cross_entropy | 1.7321177416848992 | 3.447330699998929 |
| 1551 | gse140802_t2_t16_holdout_A5::heldout_animal_A5 | 20260919.0 | cross_entropy | 1.7291872278391411 | 3.5357979999989766 |
| 1552 | gse140802_t2_t16_holdout_A5::heldout_animal_A5 | 20260920.0 | cross_entropy | 1.7189957132729614 | 3.3114573000002565 |
| 1565 | gse140802_t2_t16_holdout_B4::heldout_animal_B4 | 20260916.0 | cross_entropy | 1.347887008379161 | 4.257299099999727 |
| 1566 | gse140802_t2_t16_holdout_B4::heldout_animal_B4 | 20260917.0 | cross_entropy | 1.3307254916387738 | 3.433584399999745 |
| 1567 | gse140802_t2_t16_holdout_B4::heldout_animal_B4 | 20260918.0 | cross_entropy | 1.3794278875664578 | 3.293855899999471 |
| 1568 | gse140802_t2_t16_holdout_B4::heldout_animal_B4 | 20260919.0 | cross_entropy | 1.329578933230586 | 3.4075792000003275 |
| 1569 | gse140802_t2_t16_holdout_B4::heldout_animal_B4 | 20260920.0 | cross_entropy | 1.3122318631964618 | 3.4460927000000083 |
| 1582 | gse140802_t2_t16_holdout_B5::heldout_animal_B5 | 20260916.0 | cross_entropy | 1.8742700782800508 | 3.99507579999954 |
| 1583 | gse140802_t2_t16_holdout_B5::heldout_animal_B5 | 20260917.0 | cross_entropy | 1.920984807900613 | 3.2548543000011705 |
| 1584 | gse140802_t2_t16_holdout_B5::heldout_animal_B5 | 20260918.0 | cross_entropy | 1.872829798618899 | 3.5004533999999694 |
| 1585 | gse140802_t2_t16_holdout_B5::heldout_animal_B5 | 20260919.0 | cross_entropy | 1.900468035317768 | 3.516241100000116 |
| 1586 | gse140802_t2_t16_holdout_B5::heldout_animal_B5 | 20260920.0 | cross_entropy | 1.8927967411024325 | 3.2575082999992446 |
| 1599 | gse140802_t2_t9_holdout_A1::heldout_animal_A1 | 20260916.0 | cross_entropy | 1.7646861949849124 | 6.082265799999732 |
| 1600 | gse140802_t2_t9_holdout_A1::heldout_animal_A1 | 20260917.0 | cross_entropy | 1.7864032129844565 | 4.650442899999689 |
| 1601 | gse140802_t2_t9_holdout_A1::heldout_animal_A1 | 20260918.0 | cross_entropy | 1.806138970880387 | 4.783161699999255 |
| 1602 | gse140802_t2_t9_holdout_A1::heldout_animal_A1 | 20260919.0 | cross_entropy | 1.804183804776984 | 4.6483391999991 |
| 1603 | gse140802_t2_t9_holdout_A1::heldout_animal_A1 | 20260920.0 | cross_entropy | 1.789784886276779 | 4.753589799998736 |
| 1616 | gse140802_t2_t9_holdout_A2::heldout_animal_A2 | 20260916.0 | cross_entropy | 1.6908597370141858 | 4.972166300000026 |
| 1617 | gse140802_t2_t9_holdout_A2::heldout_animal_A2 | 20260917.0 | cross_entropy | 1.6846125551956592 | 3.842328200000338 |
| 1618 | gse140802_t2_t9_holdout_A2::heldout_animal_A2 | 20260918.0 | cross_entropy | 1.7009148161300085 | 4.119202800000494 |
| 1619 | gse140802_t2_t9_holdout_A2::heldout_animal_A2 | 20260919.0 | cross_entropy | 1.7115885765479997 | 3.81281620000118 |
| 1620 | gse140802_t2_t9_holdout_A2::heldout_animal_A2 | 20260920.0 | cross_entropy | 1.7009196199973826 | 3.738774900000863 |
| 1633 | gse140802_t2_t9_holdout_A3::heldout_animal_A3 | 20260916.0 | cross_entropy | 1.6030800787630473 | 5.939865999998801 |
| 1634 | gse140802_t2_t9_holdout_A3::heldout_animal_A3 | 20260917.0 | cross_entropy | 1.6038268292170554 | 4.551500900000974 |
| 1635 | gse140802_t2_t9_holdout_A3::heldout_animal_A3 | 20260918.0 | cross_entropy | 1.6101702585519813 | 4.758961400000771 |
| 1636 | gse140802_t2_t9_holdout_A3::heldout_animal_A3 | 20260919.0 | cross_entropy | 1.602640753276684 | 4.833847699999751 |
| 1637 | gse140802_t2_t9_holdout_A3::heldout_animal_A3 | 20260920.0 | cross_entropy | 1.5801436599609828 | 4.673117500000444 |
| 1650 | gse140802_t2_t9_holdout_B1::heldout_animal_B1 | 20260916.0 | cross_entropy | 1.7703091832974347 | 5.552295299999969 |
| 1651 | gse140802_t2_t9_holdout_B1::heldout_animal_B1 | 20260917.0 | cross_entropy | 1.7812187458044089 | 4.847299600000042 |
| 1652 | gse140802_t2_t9_holdout_B1::heldout_animal_B1 | 20260918.0 | cross_entropy | 1.7673957514599694 | 4.970558700000765 |
| 1653 | gse140802_t2_t9_holdout_B1::heldout_animal_B1 | 20260919.0 | cross_entropy | 1.750893590847897 | 4.4982584999997925 |
| 1654 | gse140802_t2_t9_holdout_B1::heldout_animal_B1 | 20260920.0 | cross_entropy | 1.8028536227123952 | 4.594531699998697 |
| 1667 | gse140802_t2_t9_holdout_B2::heldout_animal_B2 | 20260916.0 | cross_entropy | 1.7374519608887848 | 5.467017599999963 |
| 1668 | gse140802_t2_t9_holdout_B2::heldout_animal_B2 | 20260917.0 | cross_entropy | 1.7103603881452718 | 4.698739200001 |
| 1669 | gse140802_t2_t9_holdout_B2::heldout_animal_B2 | 20260918.0 | cross_entropy | 1.709456553131094 | 4.595043199999054 |
| 1670 | gse140802_t2_t9_holdout_B2::heldout_animal_B2 | 20260919.0 | cross_entropy | 1.7223934369492406 | 4.621019300000626 |
| 1671 | gse140802_t2_t9_holdout_B2::heldout_animal_B2 | 20260920.0 | cross_entropy | 1.7223866905704437 | 4.688922999999704 |
| 1684 | gse140802_t2_t9_holdout_B3::heldout_animal_B3 | 20260916.0 | cross_entropy | 1.8333675822796287 | 5.576064099999712 |
| 1685 | gse140802_t2_t9_holdout_B3::heldout_animal_B3 | 20260917.0 | cross_entropy | 1.8451779289307448 | 4.809393899999122 |
| 1686 | gse140802_t2_t9_holdout_B3::heldout_animal_B3 | 20260918.0 | cross_entropy | 1.830623937495852 | 4.4938695000000735 |
| 1687 | gse140802_t2_t9_holdout_B3::heldout_animal_B3 | 20260919.0 | cross_entropy | 1.828680638209716 | 4.501582599999892 |
| 1688 | gse140802_t2_t9_holdout_B3::heldout_animal_B3 | 20260920.0 | cross_entropy | 1.8271311581089071 | 4.37952100000075 |
| 1837 | gse140802_t2_t16_holdout_A4::heldout_animal_A4 | 20260916.0 | sinkhorn_divergence | 0.4292880408367738 | 4.053417700000864 |
| 1838 | gse140802_t2_t16_holdout_A4::heldout_animal_A4 | 20260917.0 | sinkhorn_divergence | 0.3870266507952893 | 3.204100699998889 |
| 1839 | gse140802_t2_t16_holdout_A4::heldout_animal_A4 | 20260918.0 | sinkhorn_divergence | 0.4063325339791563 | 3.3533478999997897 |
| 1840 | gse140802_t2_t16_holdout_A4::heldout_animal_A4 | 20260919.0 | sinkhorn_divergence | 0.4116172884058466 | 3.1164227000008395 |
| 1841 | gse140802_t2_t16_holdout_A4::heldout_animal_A4 | 20260920.0 | sinkhorn_divergence | 0.2963445626711744 | 3.0896375000011176 |
| 1854 | gse140802_t2_t16_holdout_A5::heldout_animal_A5 | 20260916.0 | sinkhorn_divergence | 0.2416498629086157 | 4.5700616000012815 |
| 1855 | gse140802_t2_t16_holdout_A5::heldout_animal_A5 | 20260917.0 | sinkhorn_divergence | 0.300790082491526 | 3.3212940000012168 |
| 1856 | gse140802_t2_t16_holdout_A5::heldout_animal_A5 | 20260918.0 | sinkhorn_divergence | 0.3088636084451347 | 3.447330699998929 |
| 1857 | gse140802_t2_t16_holdout_A5::heldout_animal_A5 | 20260919.0 | sinkhorn_divergence | 0.313750070028142 | 3.5357979999989766 |
| 1858 | gse140802_t2_t16_holdout_A5::heldout_animal_A5 | 20260920.0 | sinkhorn_divergence | 0.2797978959747897 | 3.3114573000002565 |
| 1871 | gse140802_t2_t16_holdout_B4::heldout_animal_B4 | 20260916.0 | sinkhorn_divergence | 0.2502223258284171 | 4.257299099999727 |
| 1872 | gse140802_t2_t16_holdout_B4::heldout_animal_B4 | 20260917.0 | sinkhorn_divergence | 0.2267077167132517 | 3.433584399999745 |
| 1873 | gse140802_t2_t16_holdout_B4::heldout_animal_B4 | 20260918.0 | sinkhorn_divergence | 0.3427998798527027 | 3.293855899999471 |
| 1874 | gse140802_t2_t16_holdout_B4::heldout_animal_B4 | 20260919.0 | sinkhorn_divergence | 0.2368373810436764 | 3.4075792000003275 |
| 1875 | gse140802_t2_t16_holdout_B4::heldout_animal_B4 | 20260920.0 | sinkhorn_divergence | 0.1770639439048382 | 3.4460927000000083 |
| 1888 | gse140802_t2_t16_holdout_B5::heldout_animal_B5 | 20260916.0 | sinkhorn_divergence | 0.4282986417502202 | 3.99507579999954 |
| 1889 | gse140802_t2_t16_holdout_B5::heldout_animal_B5 | 20260917.0 | sinkhorn_divergence | 0.4906807853848293 | 3.2548543000011705 |
| 1890 | gse140802_t2_t16_holdout_B5::heldout_animal_B5 | 20260918.0 | sinkhorn_divergence | 0.4586071408469527 | 3.5004533999999694 |
| 1891 | gse140802_t2_t16_holdout_B5::heldout_animal_B5 | 20260919.0 | sinkhorn_divergence | 0.4471311858733052 | 3.516241100000116 |
| 1892 | gse140802_t2_t16_holdout_B5::heldout_animal_B5 | 20260920.0 | sinkhorn_divergence | 0.4594800581812567 | 3.2575082999992446 |
| 1905 | gse140802_t2_t9_holdout_A1::heldout_animal_A1 | 20260916.0 | sinkhorn_divergence | 0.2805458223334896 | 6.082265799999732 |
| 1906 | gse140802_t2_t9_holdout_A1::heldout_animal_A1 | 20260917.0 | sinkhorn_divergence | 0.321036727365302 | 4.650442899999689 |
| 1907 | gse140802_t2_t9_holdout_A1::heldout_animal_A1 | 20260918.0 | sinkhorn_divergence | 0.3837208318053245 | 4.783161699999255 |
| 1908 | gse140802_t2_t9_holdout_A1::heldout_animal_A1 | 20260919.0 | sinkhorn_divergence | 0.3447013997629052 | 4.6483391999991 |
| 1909 | gse140802_t2_t9_holdout_A1::heldout_animal_A1 | 20260920.0 | sinkhorn_divergence | 0.3141684495122834 | 4.753589799998736 |
| 1922 | gse140802_t2_t9_holdout_A2::heldout_animal_A2 | 20260916.0 | sinkhorn_divergence | 0.2924583434881501 | 4.972166300000026 |
| 1923 | gse140802_t2_t9_holdout_A2::heldout_animal_A2 | 20260917.0 | sinkhorn_divergence | 0.2905074155109574 | 3.842328200000338 |
| 1924 | gse140802_t2_t9_holdout_A2::heldout_animal_A2 | 20260918.0 | sinkhorn_divergence | 0.3234037816886558 | 4.119202800000494 |
| 1925 | gse140802_t2_t9_holdout_A2::heldout_animal_A2 | 20260919.0 | sinkhorn_divergence | 0.3294348490958023 | 3.81281620000118 |
| 1926 | gse140802_t2_t9_holdout_A2::heldout_animal_A2 | 20260920.0 | sinkhorn_divergence | 0.3032308750318628 | 3.738774900000863 |
| 1939 | gse140802_t2_t9_holdout_A3::heldout_animal_A3 | 20260916.0 | sinkhorn_divergence | 0.3496447387152048 | 5.939865999998801 |
| 1940 | gse140802_t2_t9_holdout_A3::heldout_animal_A3 | 20260917.0 | sinkhorn_divergence | 0.3109018311010788 | 4.551500900000974 |
| 1941 | gse140802_t2_t9_holdout_A3::heldout_animal_A3 | 20260918.0 | sinkhorn_divergence | 0.334839800623607 | 4.758961400000771 |
| 1942 | gse140802_t2_t9_holdout_A3::heldout_animal_A3 | 20260919.0 | sinkhorn_divergence | 0.3353062973871464 | 4.833847699999751 |
| 1943 | gse140802_t2_t9_holdout_A3::heldout_animal_A3 | 20260920.0 | sinkhorn_divergence | 0.2871858013306437 | 4.673117500000444 |
| 1956 | gse140802_t2_t9_holdout_B1::heldout_animal_B1 | 20260916.0 | sinkhorn_divergence | 0.3077360906003555 | 5.552295299999969 |
| 1957 | gse140802_t2_t9_holdout_B1::heldout_animal_B1 | 20260917.0 | sinkhorn_divergence | 0.3391868015434169 | 4.847299600000042 |
| 1958 | gse140802_t2_t9_holdout_B1::heldout_animal_B1 | 20260918.0 | sinkhorn_divergence | 0.3203157149970015 | 4.970558700000765 |
| 1959 | gse140802_t2_t9_holdout_B1::heldout_animal_B1 | 20260919.0 | sinkhorn_divergence | 0.3047890122850536 | 4.4982584999997925 |
| 1960 | gse140802_t2_t9_holdout_B1::heldout_animal_B1 | 20260920.0 | sinkhorn_divergence | 0.3926886158664279 | 4.594531699998697 |
| 1973 | gse140802_t2_t9_holdout_B2::heldout_animal_B2 | 20260916.0 | sinkhorn_divergence | 0.2472196714608149 | 5.467017599999963 |
| 1974 | gse140802_t2_t9_holdout_B2::heldout_animal_B2 | 20260917.0 | sinkhorn_divergence | 0.1638580441172384 | 4.698739200001 |
| 1975 | gse140802_t2_t9_holdout_B2::heldout_animal_B2 | 20260918.0 | sinkhorn_divergence | 0.1357114931438811 | 4.595043199999054 |
| 1976 | gse140802_t2_t9_holdout_B2::heldout_animal_B2 | 20260919.0 | sinkhorn_divergence | 0.200197694207902 | 4.621019300000626 |
| 1977 | gse140802_t2_t9_holdout_B2::heldout_animal_B2 | 20260920.0 | sinkhorn_divergence | 0.1975580247895247 | 4.688922999999704 |
| 1990 | gse140802_t2_t9_holdout_B3::heldout_animal_B3 | 20260916.0 | sinkhorn_divergence | 0.3134136247222192 | 5.576064099999712 |
| 1991 | gse140802_t2_t9_holdout_B3::heldout_animal_B3 | 20260917.0 | sinkhorn_divergence | 0.3716680073707227 | 4.809393899999122 |
| 1992 | gse140802_t2_t9_holdout_B3::heldout_animal_B3 | 20260918.0 | sinkhorn_divergence | 0.3653171146367702 | 4.4938695000000735 |
| 1993 | gse140802_t2_t9_holdout_B3::heldout_animal_B3 | 20260919.0 | sinkhorn_divergence | 0.3210943843717977 | 4.501582599999892 |
| 1994 | gse140802_t2_t9_holdout_B3::heldout_animal_B3 | 20260920.0 | sinkhorn_divergence | 0.333450099665907 | 4.37952100000075 |
| 2143 | gse140802_t2_t16_holdout_A4::heldout_animal_A4 | 20260916.0 | top1_accuracy | 1.0 | 4.053417700000864 |
| 2144 | gse140802_t2_t16_holdout_A4::heldout_animal_A4 | 20260917.0 | top1_accuracy | 1.0 | 3.204100699998889 |
| 2145 | gse140802_t2_t16_holdout_A4::heldout_animal_A4 | 20260918.0 | top1_accuracy | 1.0 | 3.3533478999997897 |
| 2146 | gse140802_t2_t16_holdout_A4::heldout_animal_A4 | 20260919.0 | top1_accuracy | 1.0 | 3.1164227000008395 |
| 2147 | gse140802_t2_t16_holdout_A4::heldout_animal_A4 | 20260920.0 | top1_accuracy | 1.0 | 3.0896375000011176 |
| 2160 | gse140802_t2_t16_holdout_A5::heldout_animal_A5 | 20260916.0 | top1_accuracy | 0.0 | 4.5700616000012815 |
| 2161 | gse140802_t2_t16_holdout_A5::heldout_animal_A5 | 20260917.0 | top1_accuracy | 0.0 | 3.3212940000012168 |
| 2162 | gse140802_t2_t16_holdout_A5::heldout_animal_A5 | 20260918.0 | top1_accuracy | 0.0 | 3.447330699998929 |
| 2163 | gse140802_t2_t16_holdout_A5::heldout_animal_A5 | 20260919.0 | top1_accuracy | 0.0 | 3.5357979999989766 |
| 2164 | gse140802_t2_t16_holdout_A5::heldout_animal_A5 | 20260920.0 | top1_accuracy | 0.0 | 3.3114573000002565 |
| 2177 | gse140802_t2_t16_holdout_B4::heldout_animal_B4 | 20260916.0 | top1_accuracy | 1.0 | 4.257299099999727 |
| 2178 | gse140802_t2_t16_holdout_B4::heldout_animal_B4 | 20260917.0 | top1_accuracy | 1.0 | 3.433584399999745 |
| 2179 | gse140802_t2_t16_holdout_B4::heldout_animal_B4 | 20260918.0 | top1_accuracy | 1.0 | 3.293855899999471 |
| 2180 | gse140802_t2_t16_holdout_B4::heldout_animal_B4 | 20260919.0 | top1_accuracy | 1.0 | 3.4075792000003275 |
| 2181 | gse140802_t2_t16_holdout_B4::heldout_animal_B4 | 20260920.0 | top1_accuracy | 1.0 | 3.4460927000000083 |
| 2194 | gse140802_t2_t16_holdout_B5::heldout_animal_B5 | 20260916.0 | top1_accuracy | 1.0 | 3.99507579999954 |
| 2195 | gse140802_t2_t16_holdout_B5::heldout_animal_B5 | 20260917.0 | top1_accuracy | 1.0 | 3.2548543000011705 |
| 2196 | gse140802_t2_t16_holdout_B5::heldout_animal_B5 | 20260918.0 | top1_accuracy | 1.0 | 3.5004533999999694 |
| 2197 | gse140802_t2_t16_holdout_B5::heldout_animal_B5 | 20260919.0 | top1_accuracy | 1.0 | 3.516241100000116 |
| 2198 | gse140802_t2_t16_holdout_B5::heldout_animal_B5 | 20260920.0 | top1_accuracy | 1.0 | 3.2575082999992446 |
| 2211 | gse140802_t2_t9_holdout_A1::heldout_animal_A1 | 20260916.0 | top1_accuracy | 0.0 | 6.082265799999732 |
| 2212 | gse140802_t2_t9_holdout_A1::heldout_animal_A1 | 20260917.0 | top1_accuracy | 0.0 | 4.650442899999689 |
| 2213 | gse140802_t2_t9_holdout_A1::heldout_animal_A1 | 20260918.0 | top1_accuracy | 0.0 | 4.783161699999255 |
| 2214 | gse140802_t2_t9_holdout_A1::heldout_animal_A1 | 20260919.0 | top1_accuracy | 0.0 | 4.6483391999991 |
| 2215 | gse140802_t2_t9_holdout_A1::heldout_animal_A1 | 20260920.0 | top1_accuracy | 0.0 | 4.753589799998736 |
| 2228 | gse140802_t2_t9_holdout_A2::heldout_animal_A2 | 20260916.0 | top1_accuracy | 1.0 | 4.972166300000026 |
| 2229 | gse140802_t2_t9_holdout_A2::heldout_animal_A2 | 20260917.0 | top1_accuracy | 1.0 | 3.842328200000338 |
| 2230 | gse140802_t2_t9_holdout_A2::heldout_animal_A2 | 20260918.0 | top1_accuracy | 1.0 | 4.119202800000494 |
| 2231 | gse140802_t2_t9_holdout_A2::heldout_animal_A2 | 20260919.0 | top1_accuracy | 1.0 | 3.81281620000118 |
| 2232 | gse140802_t2_t9_holdout_A2::heldout_animal_A2 | 20260920.0 | top1_accuracy | 1.0 | 3.738774900000863 |
| 2245 | gse140802_t2_t9_holdout_A3::heldout_animal_A3 | 20260916.0 | top1_accuracy | 1.0 | 5.939865999998801 |
| 2246 | gse140802_t2_t9_holdout_A3::heldout_animal_A3 | 20260917.0 | top1_accuracy | 1.0 | 4.551500900000974 |
| 2247 | gse140802_t2_t9_holdout_A3::heldout_animal_A3 | 20260918.0 | top1_accuracy | 1.0 | 4.758961400000771 |
| 2248 | gse140802_t2_t9_holdout_A3::heldout_animal_A3 | 20260919.0 | top1_accuracy | 1.0 | 4.833847699999751 |
| 2249 | gse140802_t2_t9_holdout_A3::heldout_animal_A3 | 20260920.0 | top1_accuracy | 1.0 | 4.673117500000444 |
| 2262 | gse140802_t2_t9_holdout_B1::heldout_animal_B1 | 20260916.0 | top1_accuracy | 1.0 | 5.552295299999969 |
| 2263 | gse140802_t2_t9_holdout_B1::heldout_animal_B1 | 20260917.0 | top1_accuracy | 1.0 | 4.847299600000042 |
| 2264 | gse140802_t2_t9_holdout_B1::heldout_animal_B1 | 20260918.0 | top1_accuracy | 1.0 | 4.970558700000765 |
| 2265 | gse140802_t2_t9_holdout_B1::heldout_animal_B1 | 20260919.0 | top1_accuracy | 1.0 | 4.4982584999997925 |
| 2266 | gse140802_t2_t9_holdout_B1::heldout_animal_B1 | 20260920.0 | top1_accuracy | 1.0 | 4.594531699998697 |
| 2279 | gse140802_t2_t9_holdout_B2::heldout_animal_B2 | 20260916.0 | top1_accuracy | 1.0 | 5.467017599999963 |
| 2280 | gse140802_t2_t9_holdout_B2::heldout_animal_B2 | 20260917.0 | top1_accuracy | 1.0 | 4.698739200001 |
| 2281 | gse140802_t2_t9_holdout_B2::heldout_animal_B2 | 20260918.0 | top1_accuracy | 1.0 | 4.595043199999054 |
| 2282 | gse140802_t2_t9_holdout_B2::heldout_animal_B2 | 20260919.0 | top1_accuracy | 1.0 | 4.621019300000626 |
| 2283 | gse140802_t2_t9_holdout_B2::heldout_animal_B2 | 20260920.0 | top1_accuracy | 1.0 | 4.688922999999704 |
| 2296 | gse140802_t2_t9_holdout_B3::heldout_animal_B3 | 20260916.0 | top1_accuracy | 0.0 | 5.576064099999712 |
| 2297 | gse140802_t2_t9_holdout_B3::heldout_animal_B3 | 20260917.0 | top1_accuracy | 0.0 | 4.809393899999122 |
| 2298 | gse140802_t2_t9_holdout_B3::heldout_animal_B3 | 20260918.0 | top1_accuracy | 0.0 | 4.4938695000000735 |
| 2299 | gse140802_t2_t9_holdout_B3::heldout_animal_B3 | 20260919.0 | top1_accuracy | 0.0 | 4.501582599999892 |
| 2300 | gse140802_t2_t9_holdout_B3::heldout_animal_B3 | 20260920.0 | top1_accuracy | 0.0 | 4.37952100000075 |
| 2449 | gse140802_t2_t16_holdout_A4::heldout_animal_A4 | 20260916.0 | multiclass_brier | 0.0383831452381602 | 4.053417700000864 |
| 2450 | gse140802_t2_t16_holdout_A4::heldout_animal_A4 | 20260917.0 | multiclass_brier | 0.0303203135893136 | 3.204100699998889 |
| 2451 | gse140802_t2_t16_holdout_A4::heldout_animal_A4 | 20260918.0 | multiclass_brier | 0.0315470392442751 | 3.3533478999997897 |
| 2452 | gse140802_t2_t16_holdout_A4::heldout_animal_A4 | 20260919.0 | multiclass_brier | 0.0305767092120795 | 3.1164227000008395 |
| 2453 | gse140802_t2_t16_holdout_A4::heldout_animal_A4 | 20260920.0 | multiclass_brier | 0.0167419548518687 | 3.0896375000011176 |
| 2466 | gse140802_t2_t16_holdout_A5::heldout_animal_A5 | 20260916.0 | multiclass_brier | 0.0168559776073227 | 4.5700616000012815 |
| 2467 | gse140802_t2_t16_holdout_A5::heldout_animal_A5 | 20260917.0 | multiclass_brier | 0.0284128069395085 | 3.3212940000012168 |
| 2468 | gse140802_t2_t16_holdout_A5::heldout_animal_A5 | 20260918.0 | multiclass_brier | 0.0286041844955103 | 3.447330699998929 |
| 2469 | gse140802_t2_t16_holdout_A5::heldout_animal_A5 | 20260919.0 | multiclass_brier | 0.0199434764044091 | 3.5357979999989766 |
| 2470 | gse140802_t2_t16_holdout_A5::heldout_animal_A5 | 20260920.0 | multiclass_brier | 0.0229204037625375 | 3.3114573000002565 |
| 2483 | gse140802_t2_t16_holdout_B4::heldout_animal_B4 | 20260916.0 | multiclass_brier | 0.0142118245238723 | 4.257299099999727 |
| 2484 | gse140802_t2_t16_holdout_B4::heldout_animal_B4 | 20260917.0 | multiclass_brier | 0.0127994557202447 | 3.433584399999745 |
| 2485 | gse140802_t2_t16_holdout_B4::heldout_animal_B4 | 20260918.0 | multiclass_brier | 0.0243608953531053 | 3.293855899999471 |
| 2486 | gse140802_t2_t16_holdout_B4::heldout_animal_B4 | 20260919.0 | multiclass_brier | 0.0145924787598298 | 3.4075792000003275 |
| 2487 | gse140802_t2_t16_holdout_B4::heldout_animal_B4 | 20260920.0 | multiclass_brier | 0.0090867969908604 | 3.4460927000000083 |
| 2500 | gse140802_t2_t16_holdout_B5::heldout_animal_B5 | 20260916.0 | multiclass_brier | 0.0529005796399025 | 3.99507579999954 |
| 2501 | gse140802_t2_t16_holdout_B5::heldout_animal_B5 | 20260917.0 | multiclass_brier | 0.0661389827438839 | 3.2548543000011705 |
| 2502 | gse140802_t2_t16_holdout_B5::heldout_animal_B5 | 20260918.0 | multiclass_brier | 0.0527612588276294 | 3.5004533999999694 |
| 2503 | gse140802_t2_t16_holdout_B5::heldout_animal_B5 | 20260919.0 | multiclass_brier | 0.0564866677826343 | 3.516241100000116 |
| 2504 | gse140802_t2_t16_holdout_B5::heldout_animal_B5 | 20260920.0 | multiclass_brier | 0.0658423547340629 | 3.2575082999992446 |
| 2517 | gse140802_t2_t9_holdout_A1::heldout_animal_A1 | 20260916.0 | multiclass_brier | 0.0181420744776314 | 6.082265799999732 |
| 2518 | gse140802_t2_t9_holdout_A1::heldout_animal_A1 | 20260917.0 | multiclass_brier | 0.0234280965180468 | 4.650442899999689 |
| 2519 | gse140802_t2_t9_holdout_A1::heldout_animal_A1 | 20260918.0 | multiclass_brier | 0.0289773102918727 | 4.783161699999255 |
| 2520 | gse140802_t2_t9_holdout_A1::heldout_animal_A1 | 20260919.0 | multiclass_brier | 0.0255679428428665 | 4.6483391999991 |
| 2521 | gse140802_t2_t9_holdout_A1::heldout_animal_A1 | 20260920.0 | multiclass_brier | 0.0241976951293648 | 4.753589799998736 |
| 2534 | gse140802_t2_t9_holdout_A2::heldout_animal_A2 | 20260916.0 | multiclass_brier | 0.0146691840125786 | 4.972166300000026 |
| 2535 | gse140802_t2_t9_holdout_A2::heldout_animal_A2 | 20260917.0 | multiclass_brier | 0.0132680004465489 | 3.842328200000338 |
| 2536 | gse140802_t2_t9_holdout_A2::heldout_animal_A2 | 20260918.0 | multiclass_brier | 0.0160821666200155 | 4.119202800000494 |
| 2537 | gse140802_t2_t9_holdout_A2::heldout_animal_A2 | 20260919.0 | multiclass_brier | 0.0195649161506279 | 3.81281620000118 |
| 2538 | gse140802_t2_t9_holdout_A2::heldout_animal_A2 | 20260920.0 | multiclass_brier | 0.0159677885981451 | 3.738774900000863 |
| 2551 | gse140802_t2_t9_holdout_A3::heldout_animal_A3 | 20260916.0 | multiclass_brier | 0.0371102785068755 | 5.939865999998801 |
| 2552 | gse140802_t2_t9_holdout_A3::heldout_animal_A3 | 20260917.0 | multiclass_brier | 0.0306713186778667 | 4.551500900000974 |
| 2553 | gse140802_t2_t9_holdout_A3::heldout_animal_A3 | 20260918.0 | multiclass_brier | 0.0314997292682441 | 4.758961400000771 |
| 2554 | gse140802_t2_t9_holdout_A3::heldout_animal_A3 | 20260919.0 | multiclass_brier | 0.0335131584067983 | 4.833847699999751 |
| 2555 | gse140802_t2_t9_holdout_A3::heldout_animal_A3 | 20260920.0 | multiclass_brier | 0.025767449765389 | 4.673117500000444 |
| 2568 | gse140802_t2_t9_holdout_B1::heldout_animal_B1 | 20260916.0 | multiclass_brier | 0.0179803783063773 | 5.552295299999969 |
| 2569 | gse140802_t2_t9_holdout_B1::heldout_animal_B1 | 20260917.0 | multiclass_brier | 0.0238244978760392 | 4.847299600000042 |
| 2570 | gse140802_t2_t9_holdout_B1::heldout_animal_B1 | 20260918.0 | multiclass_brier | 0.0218255986291359 | 4.970558700000765 |
| 2571 | gse140802_t2_t9_holdout_B1::heldout_animal_B1 | 20260919.0 | multiclass_brier | 0.0185395188708786 | 4.4982584999997925 |
| 2572 | gse140802_t2_t9_holdout_B1::heldout_animal_B1 | 20260920.0 | multiclass_brier | 0.0313954881791075 | 4.594531699998697 |
| 2585 | gse140802_t2_t9_holdout_B2::heldout_animal_B2 | 20260916.0 | multiclass_brier | 0.0185102440697142 | 5.467017599999963 |
| 2586 | gse140802_t2_t9_holdout_B2::heldout_animal_B2 | 20260917.0 | multiclass_brier | 0.0061100752173676 | 4.698739200001 |
| 2587 | gse140802_t2_t9_holdout_B2::heldout_animal_B2 | 20260918.0 | multiclass_brier | 0.006246940072374 | 4.595043199999054 |
| 2588 | gse140802_t2_t9_holdout_B2::heldout_animal_B2 | 20260919.0 | multiclass_brier | 0.0112688686721183 | 4.621019300000626 |
| 2589 | gse140802_t2_t9_holdout_B2::heldout_animal_B2 | 20260920.0 | multiclass_brier | 0.0102255383097357 | 4.688922999999704 |
| 2602 | gse140802_t2_t9_holdout_B3::heldout_animal_B3 | 20260916.0 | multiclass_brier | 0.0266668847220796 | 5.576064099999712 |
| 2603 | gse140802_t2_t9_holdout_B3::heldout_animal_B3 | 20260917.0 | multiclass_brier | 0.0353758821716126 | 4.809393899999122 |
| 2604 | gse140802_t2_t9_holdout_B3::heldout_animal_B3 | 20260918.0 | multiclass_brier | 0.0285283041939432 | 4.4938695000000735 |
| 2605 | gse140802_t2_t9_holdout_B3::heldout_animal_B3 | 20260919.0 | multiclass_brier | 0.0293544737476736 | 4.501582599999892 |
| 2606 | gse140802_t2_t9_holdout_B3::heldout_animal_B3 | 20260920.0 | multiclass_brier | 0.0319328631228811 | 4.37952100000075 |

**dataset = GSE140802; benchmark = B3_prospective_composition; task = population_future_state_composition; split = held_out_animal; method = tigon; method_version = see_method_status**

| **Record** | **repeat** | **seed** | **metric** | **value** | **runtime_seconds** |
| --- | --- | --- | --- | --- | --- |
| 1536 | gse140802_t2_t16_holdout_A4::heldout_animal_A4 | 20260916.0 | cross_entropy | 6.628638523555659 | 18.7477760999991 |
| 1537 | gse140802_t2_t16_holdout_A4::heldout_animal_A4 | 20260917.0 | cross_entropy | 13.314192209732484 | 18.90845099999933 |
| 1538 | gse140802_t2_t16_holdout_A4::heldout_animal_A4 | 20260918.0 | cross_entropy | 18.38754204812657 | 18.86250409999957 |
| 1539 | gse140802_t2_t16_holdout_A4::heldout_animal_A4 | 20260919.0 | cross_entropy | 6.703480033200709 | 18.556916100000308 |
| 1540 | gse140802_t2_t16_holdout_A4::heldout_animal_A4 | 20260920.0 | cross_entropy | 6.569138974450974 | 19.21572930000002 |
| 1553 | gse140802_t2_t16_holdout_A5::heldout_animal_A5 | 20260916.0 | cross_entropy | 17.816609851391302 | 18.083980899999005 |
| 1554 | gse140802_t2_t16_holdout_A5::heldout_animal_A5 | 20260917.0 | cross_entropy | 15.97052592143594 | 17.243624500000806 |
| 1555 | gse140802_t2_t16_holdout_A5::heldout_animal_A5 | 20260918.0 | cross_entropy | 19.4803984521054 | 16.70138820000102 |
| 1556 | gse140802_t2_t16_holdout_A5::heldout_animal_A5 | 20260919.0 | cross_entropy | 7.55859773873657 | 16.343827699998656 |
| 1557 | gse140802_t2_t16_holdout_A5::heldout_animal_A5 | 20260920.0 | cross_entropy | 18.454191322679023 | 17.04764460000115 |
| 1570 | gse140802_t2_t16_holdout_B4::heldout_animal_B4 | 20260916.0 | cross_entropy | 18.414696814394865 | 18.270538199998555 |
| 1571 | gse140802_t2_t16_holdout_B4::heldout_animal_B4 | 20260917.0 | cross_entropy | 11.888055196667947 | 18.67055160000018 |
| 1572 | gse140802_t2_t16_holdout_B4::heldout_animal_B4 | 20260918.0 | cross_entropy | 13.802903935133584 | 17.15463649999947 |
| 1573 | gse140802_t2_t16_holdout_B4::heldout_animal_B4 | 20260919.0 | cross_entropy | 24.897452148531286 | 18.24013940000077 |
| 1574 | gse140802_t2_t16_holdout_B4::heldout_animal_B4 | 20260920.0 | cross_entropy | 24.672646815244548 | 18.50993170000038 |
| 1587 | gse140802_t2_t16_holdout_B5::heldout_animal_B5 | 20260916.0 | cross_entropy | 17.818659570506174 | 16.593354099999488 |
| 1588 | gse140802_t2_t16_holdout_B5::heldout_animal_B5 | 20260917.0 | cross_entropy | 8.46513681581251 | 17.23571390000143 |
| 1589 | gse140802_t2_t16_holdout_B5::heldout_animal_B5 | 20260918.0 | cross_entropy | 15.232162242468371 | 15.609840099999928 |
| 1590 | gse140802_t2_t16_holdout_B5::heldout_animal_B5 | 20260919.0 | cross_entropy | 13.663079572857528 | 16.929675400000633 |
| 1591 | gse140802_t2_t16_holdout_B5::heldout_animal_B5 | 20260920.0 | cross_entropy | 16.11281820464011 | 16.818953899999542 |
| 1604 | gse140802_t2_t9_holdout_A1::heldout_animal_A1 | 20260916.0 | cross_entropy | 13.387776100135442 | 35.19958609999958 |
| 1605 | gse140802_t2_t9_holdout_A1::heldout_animal_A1 | 20260917.0 | cross_entropy | 15.743356511209411 | 35.73301150000043 |
| 1606 | gse140802_t2_t9_holdout_A1::heldout_animal_A1 | 20260918.0 | cross_entropy | 12.876795674304296 | 34.27448600000025 |
| 1607 | gse140802_t2_t9_holdout_A1::heldout_animal_A1 | 20260919.0 | cross_entropy | 16.98787117820552 | 34.57905799999935 |
| 1608 | gse140802_t2_t9_holdout_A1::heldout_animal_A1 | 20260920.0 | cross_entropy | 8.74539195600556 | 35.60164129999976 |
| 1621 | gse140802_t2_t9_holdout_A2::heldout_animal_A2 | 20260916.0 | cross_entropy | 14.26532379526578 | 28.64675840000018 |
| 1622 | gse140802_t2_t9_holdout_A2::heldout_animal_A2 | 20260917.0 | cross_entropy | 13.35667889732795 | 28.38912030000029 |
| 1623 | gse140802_t2_t9_holdout_A2::heldout_animal_A2 | 20260918.0 | cross_entropy | 14.571201666310063 | 28.875523199998497 |
| 1624 | gse140802_t2_t9_holdout_A2::heldout_animal_A2 | 20260919.0 | cross_entropy | 12.4957714401987 | 28.72528229999989 |
| 1625 | gse140802_t2_t9_holdout_A2::heldout_animal_A2 | 20260920.0 | cross_entropy | 10.96019810387066 | 29.20851380000022 |
| 1638 | gse140802_t2_t9_holdout_A3::heldout_animal_A3 | 20260916.0 | cross_entropy | 18.2670032577458 | 35.041050100000575 |
| 1639 | gse140802_t2_t9_holdout_A3::heldout_animal_A3 | 20260917.0 | cross_entropy | 11.74621536378181 | 35.58157460000075 |
| 1640 | gse140802_t2_t9_holdout_A3::heldout_animal_A3 | 20260918.0 | cross_entropy | 17.631370700460263 | 34.25661670000045 |
| 1641 | gse140802_t2_t9_holdout_A3::heldout_animal_A3 | 20260919.0 | cross_entropy | 13.26265938748563 | 34.176104100000885 |
| 1642 | gse140802_t2_t9_holdout_A3::heldout_animal_A3 | 20260920.0 | cross_entropy | 9.173811489471795 | 35.86829039999975 |
| 1655 | gse140802_t2_t9_holdout_B1::heldout_animal_B1 | 20260916.0 | cross_entropy | 15.487180654857749 | 33.22356929999842 |
| 1656 | gse140802_t2_t9_holdout_B1::heldout_animal_B1 | 20260917.0 | cross_entropy | 15.809833686827051 | 33.27631409999958 |
| 1657 | gse140802_t2_t9_holdout_B1::heldout_animal_B1 | 20260918.0 | cross_entropy | 20.32887356864767 | 33.15298780000012 |
| 1658 | gse140802_t2_t9_holdout_B1::heldout_animal_B1 | 20260919.0 | cross_entropy | 14.644997315676912 | 33.69306250000045 |
| 1659 | gse140802_t2_t9_holdout_B1::heldout_animal_B1 | 20260920.0 | cross_entropy | 9.214911959544612 | 34.06901469999866 |
| 1672 | gse140802_t2_t9_holdout_B2::heldout_animal_B2 | 20260916.0 | cross_entropy | 16.79368238383781 | 35.26367279999977 |
| 1673 | gse140802_t2_t9_holdout_B2::heldout_animal_B2 | 20260917.0 | cross_entropy | 13.749594279281007 | 34.13817350000136 |
| 1674 | gse140802_t2_t9_holdout_B2::heldout_animal_B2 | 20260918.0 | cross_entropy | 19.41127047029803 | 33.405341800000315 |
| 1675 | gse140802_t2_t9_holdout_B2::heldout_animal_B2 | 20260919.0 | cross_entropy | 13.050373470941883 | 33.09283400000095 |
| 1676 | gse140802_t2_t9_holdout_B2::heldout_animal_B2 | 20260920.0 | cross_entropy | 10.21723403423272 | 33.8409458999995 |
| 1689 | gse140802_t2_t9_holdout_B3::heldout_animal_B3 | 20260916.0 | cross_entropy | 12.829095530920872 | 33.34541990000071 |
| 1690 | gse140802_t2_t9_holdout_B3::heldout_animal_B3 | 20260917.0 | cross_entropy | 12.774760558316355 | 34.24396390000038 |
| 1691 | gse140802_t2_t9_holdout_B3::heldout_animal_B3 | 20260918.0 | cross_entropy | 12.370172229920492 | 34.28840390000005 |
| 1692 | gse140802_t2_t9_holdout_B3::heldout_animal_B3 | 20260919.0 | cross_entropy | 12.357865885882116 | 34.8315333999999 |
| 1693 | gse140802_t2_t9_holdout_B3::heldout_animal_B3 | 20260920.0 | cross_entropy | 8.243626318985154 | 34.792659299999286 |
| 1842 | gse140802_t2_t16_holdout_A4::heldout_animal_A4 | 20260916.0 | sinkhorn_divergence | 1.8876476762866727 | 18.7477760999991 |
| 1843 | gse140802_t2_t16_holdout_A4::heldout_animal_A4 | 20260917.0 | sinkhorn_divergence | 1.8906593147187487 | 18.90845099999933 |
| 1844 | gse140802_t2_t16_holdout_A4::heldout_animal_A4 | 20260918.0 | sinkhorn_divergence | 1.8890994994059929 | 18.86250409999957 |
| 1845 | gse140802_t2_t16_holdout_A4::heldout_animal_A4 | 20260919.0 | sinkhorn_divergence | 1.886470957709537 | 18.556916100000308 |
| 1846 | gse140802_t2_t16_holdout_A4::heldout_animal_A4 | 20260920.0 | sinkhorn_divergence | 1.8754559988159087 | 19.21572930000002 |
| 1859 | gse140802_t2_t16_holdout_A5::heldout_animal_A5 | 20260916.0 | sinkhorn_divergence | 1.7477817057986065 | 18.083980899999005 |
| 1860 | gse140802_t2_t16_holdout_A5::heldout_animal_A5 | 20260917.0 | sinkhorn_divergence | 1.7107065672090835 | 17.243624500000806 |
| 1861 | gse140802_t2_t16_holdout_A5::heldout_animal_A5 | 20260918.0 | sinkhorn_divergence | 1.7245590673389035 | 16.70138820000102 |
| 1862 | gse140802_t2_t16_holdout_A5::heldout_animal_A5 | 20260919.0 | sinkhorn_divergence | 1.731308467916521 | 16.343827699998656 |
| 1863 | gse140802_t2_t16_holdout_A5::heldout_animal_A5 | 20260920.0 | sinkhorn_divergence | 1.7431307057084937 | 17.04764460000115 |
| 1876 | gse140802_t2_t16_holdout_B4::heldout_animal_B4 | 20260916.0 | sinkhorn_divergence | 1.878772888532546 | 18.270538199998555 |
| 1877 | gse140802_t2_t16_holdout_B4::heldout_animal_B4 | 20260917.0 | sinkhorn_divergence | 1.861138487188818 | 18.67055160000018 |
| 1878 | gse140802_t2_t16_holdout_B4::heldout_animal_B4 | 20260918.0 | sinkhorn_divergence | 1.8798515691332025 | 17.15463649999947 |
| 1879 | gse140802_t2_t16_holdout_B4::heldout_animal_B4 | 20260919.0 | sinkhorn_divergence | 1.8853628411931016 | 18.24013940000077 |
| 1880 | gse140802_t2_t16_holdout_B4::heldout_animal_B4 | 20260920.0 | sinkhorn_divergence | 1.8795055709689208 | 18.50993170000038 |
| 1893 | gse140802_t2_t16_holdout_B5::heldout_animal_B5 | 20260916.0 | sinkhorn_divergence | 1.7693380504331544 | 16.593354099999488 |
| 1894 | gse140802_t2_t16_holdout_B5::heldout_animal_B5 | 20260917.0 | sinkhorn_divergence | 1.7416326420470285 | 17.23571390000143 |
| 1895 | gse140802_t2_t16_holdout_B5::heldout_animal_B5 | 20260918.0 | sinkhorn_divergence | 1.757053533352916 | 15.609840099999928 |
| 1896 | gse140802_t2_t16_holdout_B5::heldout_animal_B5 | 20260919.0 | sinkhorn_divergence | 1.754418505567023 | 16.929675400000633 |
| 1897 | gse140802_t2_t16_holdout_B5::heldout_animal_B5 | 20260920.0 | sinkhorn_divergence | 1.7593989944519817 | 16.818953899999542 |
| 1910 | gse140802_t2_t9_holdout_A1::heldout_animal_A1 | 20260916.0 | sinkhorn_divergence | 1.550099573958161 | 35.19958609999958 |
| 1911 | gse140802_t2_t9_holdout_A1::heldout_animal_A1 | 20260917.0 | sinkhorn_divergence | 1.5393444639669296 | 35.73301150000043 |
| 1912 | gse140802_t2_t9_holdout_A1::heldout_animal_A1 | 20260918.0 | sinkhorn_divergence | 1.550236307298667 | 34.27448600000025 |
| 1913 | gse140802_t2_t9_holdout_A1::heldout_animal_A1 | 20260919.0 | sinkhorn_divergence | 1.5541903055825042 | 34.57905799999935 |
| 1914 | gse140802_t2_t9_holdout_A1::heldout_animal_A1 | 20260920.0 | sinkhorn_divergence | 1.559301705276708 | 35.60164129999976 |
| 1927 | gse140802_t2_t9_holdout_A2::heldout_animal_A2 | 20260916.0 | sinkhorn_divergence | 1.5616100597658016 | 28.64675840000018 |
| 1928 | gse140802_t2_t9_holdout_A2::heldout_animal_A2 | 20260917.0 | sinkhorn_divergence | 1.5508559088536606 | 28.38912030000029 |
| 1929 | gse140802_t2_t9_holdout_A2::heldout_animal_A2 | 20260918.0 | sinkhorn_divergence | 1.5621030916409893 | 28.875523199998497 |
| 1930 | gse140802_t2_t9_holdout_A2::heldout_animal_A2 | 20260919.0 | sinkhorn_divergence | 1.567243384322748 | 28.72528229999989 |
| 1931 | gse140802_t2_t9_holdout_A2::heldout_animal_A2 | 20260920.0 | sinkhorn_divergence | 1.5708184480094385 | 29.20851380000022 |
| 1944 | gse140802_t2_t9_holdout_A3::heldout_animal_A3 | 20260916.0 | sinkhorn_divergence | 1.7072735399754857 | 35.041050100000575 |
| 1945 | gse140802_t2_t9_holdout_A3::heldout_animal_A3 | 20260917.0 | sinkhorn_divergence | 1.6970379412793668 | 35.58157460000075 |
| 1946 | gse140802_t2_t9_holdout_A3::heldout_animal_A3 | 20260918.0 | sinkhorn_divergence | 1.7067295235812734 | 34.25661670000045 |
| 1947 | gse140802_t2_t9_holdout_A3::heldout_animal_A3 | 20260919.0 | sinkhorn_divergence | 1.7124693535493554 | 34.176104100000885 |
| 1948 | gse140802_t2_t9_holdout_A3::heldout_animal_A3 | 20260920.0 | sinkhorn_divergence | 1.716230958657071 | 35.86829039999975 |
| 1961 | gse140802_t2_t9_holdout_B1::heldout_animal_B1 | 20260916.0 | sinkhorn_divergence | 1.6888813391353643 | 33.22356929999842 |
| 1962 | gse140802_t2_t9_holdout_B1::heldout_animal_B1 | 20260917.0 | sinkhorn_divergence | 1.6791378373856598 | 33.27631409999958 |
| 1963 | gse140802_t2_t9_holdout_B1::heldout_animal_B1 | 20260918.0 | sinkhorn_divergence | 1.690190744401712 | 33.15298780000012 |
| 1964 | gse140802_t2_t9_holdout_B1::heldout_animal_B1 | 20260919.0 | sinkhorn_divergence | 1.6945087240288257 | 33.69306250000045 |
| 1965 | gse140802_t2_t9_holdout_B1::heldout_animal_B1 | 20260920.0 | sinkhorn_divergence | 1.6980926244787788 | 34.06901469999866 |
| 1978 | gse140802_t2_t9_holdout_B2::heldout_animal_B2 | 20260916.0 | sinkhorn_divergence | 1.721558027145386 | 35.26367279999977 |
| 1979 | gse140802_t2_t9_holdout_B2::heldout_animal_B2 | 20260917.0 | sinkhorn_divergence | 1.7115126019597866 | 34.13817350000136 |
| 1980 | gse140802_t2_t9_holdout_B2::heldout_animal_B2 | 20260918.0 | sinkhorn_divergence | 1.7222380461908124 | 33.405341800000315 |
| 1981 | gse140802_t2_t9_holdout_B2::heldout_animal_B2 | 20260919.0 | sinkhorn_divergence | 1.7270898116655808 | 33.09283400000095 |
| 1982 | gse140802_t2_t9_holdout_B2::heldout_animal_B2 | 20260920.0 | sinkhorn_divergence | 1.7307060561284546 | 33.8409458999995 |
| 1995 | gse140802_t2_t9_holdout_B3::heldout_animal_B3 | 20260916.0 | sinkhorn_divergence | 1.4072528280691556 | 33.34541990000071 |
| 1996 | gse140802_t2_t9_holdout_B3::heldout_animal_B3 | 20260917.0 | sinkhorn_divergence | 1.397422486689103 | 34.24396390000038 |
| 1997 | gse140802_t2_t9_holdout_B3::heldout_animal_B3 | 20260918.0 | sinkhorn_divergence | 1.4086917950688838 | 34.28840390000005 |
| 1998 | gse140802_t2_t9_holdout_B3::heldout_animal_B3 | 20260919.0 | sinkhorn_divergence | 1.411134077109765 | 34.8315333999999 |
| 1999 | gse140802_t2_t9_holdout_B3::heldout_animal_B3 | 20260920.0 | sinkhorn_divergence | 1.4169451291822586 | 34.792659299999286 |
| 2148 | gse140802_t2_t16_holdout_A4::heldout_animal_A4 | 20260916.0 | top1_accuracy | 0.0 | 18.7477760999991 |
| 2149 | gse140802_t2_t16_holdout_A4::heldout_animal_A4 | 20260917.0 | top1_accuracy | 0.0 | 18.90845099999933 |
| 2150 | gse140802_t2_t16_holdout_A4::heldout_animal_A4 | 20260918.0 | top1_accuracy | 0.0 | 18.86250409999957 |
| 2151 | gse140802_t2_t16_holdout_A4::heldout_animal_A4 | 20260919.0 | top1_accuracy | 0.0 | 18.556916100000308 |
| 2152 | gse140802_t2_t16_holdout_A4::heldout_animal_A4 | 20260920.0 | top1_accuracy | 0.0 | 19.21572930000002 |
| 2165 | gse140802_t2_t16_holdout_A5::heldout_animal_A5 | 20260916.0 | top1_accuracy | 0.0 | 18.083980899999005 |
| 2166 | gse140802_t2_t16_holdout_A5::heldout_animal_A5 | 20260917.0 | top1_accuracy | 0.0 | 17.243624500000806 |
| 2167 | gse140802_t2_t16_holdout_A5::heldout_animal_A5 | 20260918.0 | top1_accuracy | 0.0 | 16.70138820000102 |
| 2168 | gse140802_t2_t16_holdout_A5::heldout_animal_A5 | 20260919.0 | top1_accuracy | 0.0 | 16.343827699998656 |
| 2169 | gse140802_t2_t16_holdout_A5::heldout_animal_A5 | 20260920.0 | top1_accuracy | 0.0 | 17.04764460000115 |
| 2182 | gse140802_t2_t16_holdout_B4::heldout_animal_B4 | 20260916.0 | top1_accuracy | 0.0 | 18.270538199998555 |
| 2183 | gse140802_t2_t16_holdout_B4::heldout_animal_B4 | 20260917.0 | top1_accuracy | 0.0 | 18.67055160000018 |
| 2184 | gse140802_t2_t16_holdout_B4::heldout_animal_B4 | 20260918.0 | top1_accuracy | 0.0 | 17.15463649999947 |
| 2185 | gse140802_t2_t16_holdout_B4::heldout_animal_B4 | 20260919.0 | top1_accuracy | 0.0 | 18.24013940000077 |
| 2186 | gse140802_t2_t16_holdout_B4::heldout_animal_B4 | 20260920.0 | top1_accuracy | 0.0 | 18.50993170000038 |
| 2199 | gse140802_t2_t16_holdout_B5::heldout_animal_B5 | 20260916.0 | top1_accuracy | 0.0 | 16.593354099999488 |
| 2200 | gse140802_t2_t16_holdout_B5::heldout_animal_B5 | 20260917.0 | top1_accuracy | 0.0 | 17.23571390000143 |
| 2201 | gse140802_t2_t16_holdout_B5::heldout_animal_B5 | 20260918.0 | top1_accuracy | 0.0 | 15.609840099999928 |
| 2202 | gse140802_t2_t16_holdout_B5::heldout_animal_B5 | 20260919.0 | top1_accuracy | 0.0 | 16.929675400000633 |
| 2203 | gse140802_t2_t16_holdout_B5::heldout_animal_B5 | 20260920.0 | top1_accuracy | 0.0 | 16.818953899999542 |
| 2216 | gse140802_t2_t9_holdout_A1::heldout_animal_A1 | 20260916.0 | top1_accuracy | 0.0 | 35.19958609999958 |
| 2217 | gse140802_t2_t9_holdout_A1::heldout_animal_A1 | 20260917.0 | top1_accuracy | 0.0 | 35.73301150000043 |
| 2218 | gse140802_t2_t9_holdout_A1::heldout_animal_A1 | 20260918.0 | top1_accuracy | 0.0 | 34.27448600000025 |
| 2219 | gse140802_t2_t9_holdout_A1::heldout_animal_A1 | 20260919.0 | top1_accuracy | 0.0 | 34.57905799999935 |
| 2220 | gse140802_t2_t9_holdout_A1::heldout_animal_A1 | 20260920.0 | top1_accuracy | 0.0 | 35.60164129999976 |
| 2233 | gse140802_t2_t9_holdout_A2::heldout_animal_A2 | 20260916.0 | top1_accuracy | 0.0 | 28.64675840000018 |
| 2234 | gse140802_t2_t9_holdout_A2::heldout_animal_A2 | 20260917.0 | top1_accuracy | 0.0 | 28.38912030000029 |
| 2235 | gse140802_t2_t9_holdout_A2::heldout_animal_A2 | 20260918.0 | top1_accuracy | 0.0 | 28.875523199998497 |
| 2236 | gse140802_t2_t9_holdout_A2::heldout_animal_A2 | 20260919.0 | top1_accuracy | 0.0 | 28.72528229999989 |
| 2237 | gse140802_t2_t9_holdout_A2::heldout_animal_A2 | 20260920.0 | top1_accuracy | 0.0 | 29.20851380000022 |
| 2250 | gse140802_t2_t9_holdout_A3::heldout_animal_A3 | 20260916.0 | top1_accuracy | 0.0 | 35.041050100000575 |
| 2251 | gse140802_t2_t9_holdout_A3::heldout_animal_A3 | 20260917.0 | top1_accuracy | 0.0 | 35.58157460000075 |
| 2252 | gse140802_t2_t9_holdout_A3::heldout_animal_A3 | 20260918.0 | top1_accuracy | 0.0 | 34.25661670000045 |
| 2253 | gse140802_t2_t9_holdout_A3::heldout_animal_A3 | 20260919.0 | top1_accuracy | 0.0 | 34.176104100000885 |
| 2254 | gse140802_t2_t9_holdout_A3::heldout_animal_A3 | 20260920.0 | top1_accuracy | 0.0 | 35.86829039999975 |
| 2267 | gse140802_t2_t9_holdout_B1::heldout_animal_B1 | 20260916.0 | top1_accuracy | 0.0 | 33.22356929999842 |
| 2268 | gse140802_t2_t9_holdout_B1::heldout_animal_B1 | 20260917.0 | top1_accuracy | 0.0 | 33.27631409999958 |
| 2269 | gse140802_t2_t9_holdout_B1::heldout_animal_B1 | 20260918.0 | top1_accuracy | 0.0 | 33.15298780000012 |
| 2270 | gse140802_t2_t9_holdout_B1::heldout_animal_B1 | 20260919.0 | top1_accuracy | 0.0 | 33.69306250000045 |
| 2271 | gse140802_t2_t9_holdout_B1::heldout_animal_B1 | 20260920.0 | top1_accuracy | 0.0 | 34.06901469999866 |
| 2284 | gse140802_t2_t9_holdout_B2::heldout_animal_B2 | 20260916.0 | top1_accuracy | 0.0 | 35.26367279999977 |
| 2285 | gse140802_t2_t9_holdout_B2::heldout_animal_B2 | 20260917.0 | top1_accuracy | 0.0 | 34.13817350000136 |
| 2286 | gse140802_t2_t9_holdout_B2::heldout_animal_B2 | 20260918.0 | top1_accuracy | 0.0 | 33.405341800000315 |
| 2287 | gse140802_t2_t9_holdout_B2::heldout_animal_B2 | 20260919.0 | top1_accuracy | 0.0 | 33.09283400000095 |
| 2288 | gse140802_t2_t9_holdout_B2::heldout_animal_B2 | 20260920.0 | top1_accuracy | 0.0 | 33.8409458999995 |
| 2301 | gse140802_t2_t9_holdout_B3::heldout_animal_B3 | 20260916.0 | top1_accuracy | 1.0 | 33.34541990000071 |
| 2302 | gse140802_t2_t9_holdout_B3::heldout_animal_B3 | 20260917.0 | top1_accuracy | 1.0 | 34.24396390000038 |
| 2303 | gse140802_t2_t9_holdout_B3::heldout_animal_B3 | 20260918.0 | top1_accuracy | 1.0 | 34.28840390000005 |
| 2304 | gse140802_t2_t9_holdout_B3::heldout_animal_B3 | 20260919.0 | top1_accuracy | 1.0 | 34.8315333999999 |
| 2305 | gse140802_t2_t9_holdout_B3::heldout_animal_B3 | 20260920.0 | top1_accuracy | 1.0 | 34.792659299999286 |
| 2454 | gse140802_t2_t16_holdout_A4::heldout_animal_A4 | 20260916.0 | multiclass_brier | 1.2808333056011716 | 18.7477760999991 |
| 2455 | gse140802_t2_t16_holdout_A4::heldout_animal_A4 | 20260917.0 | multiclass_brier | 1.2838265922052985 | 18.90845099999933 |
| 2456 | gse140802_t2_t16_holdout_A4::heldout_animal_A4 | 20260918.0 | multiclass_brier | 1.2843719309230923 | 18.86250409999957 |
| 2457 | gse140802_t2_t16_holdout_A4::heldout_animal_A4 | 20260919.0 | multiclass_brier | 1.2788856820869523 | 18.556916100000308 |
| 2458 | gse140802_t2_t16_holdout_A4::heldout_animal_A4 | 20260920.0 | multiclass_brier | 1.267319207826361 | 19.21572930000002 |
| 2471 | gse140802_t2_t16_holdout_A5::heldout_animal_A5 | 20260916.0 | multiclass_brier | 0.9708388947257688 | 18.083980899999005 |
| 2472 | gse140802_t2_t16_holdout_A5::heldout_animal_A5 | 20260917.0 | multiclass_brier | 0.93274252495029 | 17.243624500000806 |
| 2473 | gse140802_t2_t16_holdout_A5::heldout_animal_A5 | 20260918.0 | multiclass_brier | 0.948322448250163 | 16.70138820000102 |
| 2474 | gse140802_t2_t16_holdout_A5::heldout_animal_A5 | 20260919.0 | multiclass_brier | 0.9527377778001078 | 16.343827699998656 |
| 2475 | gse140802_t2_t16_holdout_A5::heldout_animal_A5 | 20260920.0 | multiclass_brier | 0.9655835566191078 | 17.04764460000115 |
| 2488 | gse140802_t2_t16_holdout_B4::heldout_animal_B4 | 20260916.0 | multiclass_brier | 1.249304258494365 | 18.270538199998555 |
| 2489 | gse140802_t2_t16_holdout_B4::heldout_animal_B4 | 20260917.0 | multiclass_brier | 1.2260145061521657 | 18.67055160000018 |
| 2490 | gse140802_t2_t16_holdout_B4::heldout_animal_B4 | 20260918.0 | multiclass_brier | 1.25002608434258 | 17.15463649999947 |
| 2491 | gse140802_t2_t16_holdout_B4::heldout_animal_B4 | 20260919.0 | multiclass_brier | 1.2562855056537412 | 18.24013940000077 |
| 2492 | gse140802_t2_t16_holdout_B4::heldout_animal_B4 | 20260920.0 | multiclass_brier | 1.2506003152964111 | 18.50993170000038 |
| 2505 | gse140802_t2_t16_holdout_B5::heldout_animal_B5 | 20260916.0 | multiclass_brier | 0.970754384997074 | 16.593354099999488 |
| 2506 | gse140802_t2_t16_holdout_B5::heldout_animal_B5 | 20260917.0 | multiclass_brier | 0.9408248466968756 | 17.23571390000143 |
| 2507 | gse140802_t2_t16_holdout_B5::heldout_animal_B5 | 20260918.0 | multiclass_brier | 0.9576179951752232 | 15.609840099999928 |
| 2508 | gse140802_t2_t16_holdout_B5::heldout_animal_B5 | 20260919.0 | multiclass_brier | 0.9556078830840904 | 16.929675400000633 |
| 2509 | gse140802_t2_t16_holdout_B5::heldout_animal_B5 | 20260920.0 | multiclass_brier | 0.9604938374128672 | 16.818953899999542 |
| 2522 | gse140802_t2_t9_holdout_A1::heldout_animal_A1 | 20260916.0 | multiclass_brier | 0.746456712876549 | 35.19958609999958 |
| 2523 | gse140802_t2_t9_holdout_A1::heldout_animal_A1 | 20260917.0 | multiclass_brier | 0.7358480844367087 | 35.73301150000043 |
| 2524 | gse140802_t2_t9_holdout_A1::heldout_animal_A1 | 20260918.0 | multiclass_brier | 0.7466538421415297 | 34.27448600000025 |
| 2525 | gse140802_t2_t9_holdout_A1::heldout_animal_A1 | 20260919.0 | multiclass_brier | 0.7498330913413428 | 34.57905799999935 |
| 2526 | gse140802_t2_t9_holdout_A1::heldout_animal_A1 | 20260920.0 | multiclass_brier | 0.7546083785384786 | 35.60164129999976 |
| 2539 | gse140802_t2_t9_holdout_A2::heldout_animal_A2 | 20260916.0 | multiclass_brier | 0.790241679015412 | 28.64675840000018 |
| 2540 | gse140802_t2_t9_holdout_A2::heldout_animal_A2 | 20260917.0 | multiclass_brier | 0.7771646748079996 | 28.38912030000029 |
| 2541 | gse140802_t2_t9_holdout_A2::heldout_animal_A2 | 20260918.0 | multiclass_brier | 0.7900712064577438 | 28.875523199998497 |
| 2542 | gse140802_t2_t9_holdout_A2::heldout_animal_A2 | 20260919.0 | multiclass_brier | 0.7942899040829162 | 28.72528229999989 |
| 2543 | gse140802_t2_t9_holdout_A2::heldout_animal_A2 | 20260920.0 | multiclass_brier | 0.798684640147121 | 29.20851380000022 |
| 2556 | gse140802_t2_t9_holdout_A3::heldout_animal_A3 | 20260916.0 | multiclass_brier | 0.99518638574774 | 35.041050100000575 |
| 2557 | gse140802_t2_t9_holdout_A3::heldout_animal_A3 | 20260917.0 | multiclass_brier | 0.9791218255317292 | 35.58157460000075 |
| 2558 | gse140802_t2_t9_holdout_A3::heldout_animal_A3 | 20260918.0 | multiclass_brier | 0.9946708871670918 | 34.25661670000045 |
| 2559 | gse140802_t2_t9_holdout_A3::heldout_animal_A3 | 20260919.0 | multiclass_brier | 0.9981731760382538 | 34.176104100000885 |
| 2560 | gse140802_t2_t9_holdout_A3::heldout_animal_A3 | 20260920.0 | multiclass_brier | 1.0034453079301957 | 35.86829039999975 |
| 2573 | gse140802_t2_t9_holdout_B1::heldout_animal_B1 | 20260916.0 | multiclass_brier | 0.896323555075049 | 33.22356929999842 |
| 2574 | gse140802_t2_t9_holdout_B1::heldout_animal_B1 | 20260917.0 | multiclass_brier | 0.884405705564945 | 33.27631409999958 |
| 2575 | gse140802_t2_t9_holdout_B1::heldout_animal_B1 | 20260918.0 | multiclass_brier | 0.8977914465472956 | 33.15298780000012 |
| 2576 | gse140802_t2_t9_holdout_B1::heldout_animal_B1 | 20260919.0 | multiclass_brier | 0.901078697325552 | 33.69306250000045 |
| 2577 | gse140802_t2_t9_holdout_B1::heldout_animal_B1 | 20260920.0 | multiclass_brier | 0.90507669731375 | 34.06901469999866 |
| 2590 | gse140802_t2_t9_holdout_B2::heldout_animal_B2 | 20260916.0 | multiclass_brier | 0.9428985584824214 | 35.26367279999977 |
| 2591 | gse140802_t2_t9_holdout_B2::heldout_animal_B2 | 20260917.0 | multiclass_brier | 0.928890678335478 | 34.13817350000136 |
| 2592 | gse140802_t2_t9_holdout_B2::heldout_animal_B2 | 20260918.0 | multiclass_brier | 0.9433644555319626 | 33.405341800000315 |
| 2593 | gse140802_t2_t9_holdout_B2::heldout_animal_B2 | 20260919.0 | multiclass_brier | 0.9469928253180362 | 33.09283400000095 |
| 2594 | gse140802_t2_t9_holdout_B2::heldout_animal_B2 | 20260920.0 | multiclass_brier | 0.95166159700494 | 33.8409458999995 |
| 2607 | gse140802_t2_t9_holdout_B3::heldout_animal_B3 | 20260916.0 | multiclass_brier | 0.604169659373308 | 33.34541990000071 |
| 2608 | gse140802_t2_t9_holdout_B3::heldout_animal_B3 | 20260917.0 | multiclass_brier | 0.5945431471079732 | 34.24396390000038 |
| 2609 | gse140802_t2_t9_holdout_B3::heldout_animal_B3 | 20260918.0 | multiclass_brier | 0.605418994031077 | 34.28840390000005 |
| 2610 | gse140802_t2_t9_holdout_B3::heldout_animal_B3 | 20260919.0 | multiclass_brier | 0.6067287107839512 | 34.8315333999999 |
| 2611 | gse140802_t2_t9_holdout_B3::heldout_animal_B3 | 20260920.0 | multiclass_brier | 0.6118256235940106 | 34.792659299999286 |

**dataset = GSE140802; benchmark = B3_prospective_composition; task = population_future_state_composition; split = held_out_animal; method = mioflow; method_version = see_method_status**

| **Record** | **repeat** | **seed** | **metric** | **value** | **runtime_seconds** |
| --- | --- | --- | --- | --- | --- |
| 1541 | gse140802_t2_t16_holdout_A4::heldout_animal_A4 | 20260916.0 | cross_entropy | 1.8612552698122504 | 11.381618999999771 |
| 1542 | gse140802_t2_t16_holdout_A4::heldout_animal_A4 | 20260917.0 | cross_entropy | 1.9238425283562672 | 10.828887900001064 |
| 1543 | gse140802_t2_t16_holdout_A4::heldout_animal_A4 | 20260918.0 | cross_entropy | 1.7218107118444663 | 9.209258099999715 |
| 1544 | gse140802_t2_t16_holdout_A4::heldout_animal_A4 | 20260919.0 | cross_entropy | 2.108926086745366 | 8.240687600000456 |
| 1545 | gse140802_t2_t16_holdout_A4::heldout_animal_A4 | 20260920.0 | cross_entropy | 1.8945537015130285 | 9.692493900000043 |
| 1558 | gse140802_t2_t16_holdout_A5::heldout_animal_A5 | 20260916.0 | cross_entropy | 2.159897919546067 | 10.093173400000524 |
| 1559 | gse140802_t2_t16_holdout_A5::heldout_animal_A5 | 20260917.0 | cross_entropy | 2.104483410736593 | 10.584232999999584 |
| 1560 | gse140802_t2_t16_holdout_A5::heldout_animal_A5 | 20260918.0 | cross_entropy | 1.988812334727837 | 9.87224210000022 |
| 1561 | gse140802_t2_t16_holdout_A5::heldout_animal_A5 | 20260919.0 | cross_entropy | 3.0689502206314034 | 10.484369800000422 |
| 1562 | gse140802_t2_t16_holdout_A5::heldout_animal_A5 | 20260920.0 | cross_entropy | 2.102423124439561 | 10.327723500000502 |
| 1575 | gse140802_t2_t16_holdout_B4::heldout_animal_B4 | 20260916.0 | cross_entropy | 1.8763606821826544 | 10.841824400000403 |
| 1576 | gse140802_t2_t16_holdout_B4::heldout_animal_B4 | 20260917.0 | cross_entropy | 1.9811944318973824 | 9.889872700001431 |
| 1577 | gse140802_t2_t16_holdout_B4::heldout_animal_B4 | 20260918.0 | cross_entropy | 1.7457279750248849 | 9.274696800001038 |
| 1578 | gse140802_t2_t16_holdout_B4::heldout_animal_B4 | 20260919.0 | cross_entropy | 2.4452239735407537 | 10.27010639999935 |
| 1579 | gse140802_t2_t16_holdout_B4::heldout_animal_B4 | 20260920.0 | cross_entropy | 1.9223560007916936 | 8.525537100000292 |
| 1592 | gse140802_t2_t16_holdout_B5::heldout_animal_B5 | 20260916.0 | cross_entropy | 2.20587444115234 | 12.640654699998777 |
| 1593 | gse140802_t2_t16_holdout_B5::heldout_animal_B5 | 20260917.0 | cross_entropy | 2.1416934873494875 | 10.670966799998496 |
| 1594 | gse140802_t2_t16_holdout_B5::heldout_animal_B5 | 20260918.0 | cross_entropy | 2.096334465129865 | 12.243431499999131 |
| 1595 | gse140802_t2_t16_holdout_B5::heldout_animal_B5 | 20260919.0 | cross_entropy | 3.3490365749674496 | 11.23134349999964 |
| 1596 | gse140802_t2_t16_holdout_B5::heldout_animal_B5 | 20260920.0 | cross_entropy | 2.266219988470884 | 9.15854279999985 |
| 1609 | gse140802_t2_t9_holdout_A1::heldout_animal_A1 | 20260916.0 | cross_entropy | 3.0584053111522658 | 13.377239100000224 |
| 1610 | gse140802_t2_t9_holdout_A1::heldout_animal_A1 | 20260917.0 | cross_entropy | 2.0500414087517784 | 12.320767999999587 |
| 1611 | gse140802_t2_t9_holdout_A1::heldout_animal_A1 | 20260918.0 | cross_entropy | 1.9713803718651923 | 11.287288700001229 |
| 1612 | gse140802_t2_t9_holdout_A1::heldout_animal_A1 | 20260919.0 | cross_entropy | 2.811135312634924 | 12.021956800001137 |
| 1613 | gse140802_t2_t9_holdout_A1::heldout_animal_A1 | 20260920.0 | cross_entropy | 2.108913203925716 | 11.787559000000329 |
| 1626 | gse140802_t2_t9_holdout_A2::heldout_animal_A2 | 20260916.0 | cross_entropy | 1.7911311218762926 | 12.12737290000041 |
| 1627 | gse140802_t2_t9_holdout_A2::heldout_animal_A2 | 20260917.0 | cross_entropy | 1.9610628821197047 | 10.070772400000353 |
| 1628 | gse140802_t2_t9_holdout_A2::heldout_animal_A2 | 20260918.0 | cross_entropy | 1.8791390020852567 | 11.219067100000755 |
| 1629 | gse140802_t2_t9_holdout_A2::heldout_animal_A2 | 20260919.0 | cross_entropy | 2.1176320370032555 | 11.366179100001318 |
| 1630 | gse140802_t2_t9_holdout_A2::heldout_animal_A2 | 20260920.0 | cross_entropy | 2.05062372802716 | 9.137043200000337 |
| 1643 | gse140802_t2_t9_holdout_A3::heldout_animal_A3 | 20260916.0 | cross_entropy | 2.816129533903609 | 13.791628000000856 |
| 1644 | gse140802_t2_t9_holdout_A3::heldout_animal_A3 | 20260917.0 | cross_entropy | 2.022614511917723 | 12.229715900000883 |
| 1645 | gse140802_t2_t9_holdout_A3::heldout_animal_A3 | 20260918.0 | cross_entropy | 1.9560714375066683 | 11.05017670000052 |
| 1646 | gse140802_t2_t9_holdout_A3::heldout_animal_A3 | 20260919.0 | cross_entropy | 2.2484836949811084 | 13.881138999999166 |
| 1647 | gse140802_t2_t9_holdout_A3::heldout_animal_A3 | 20260920.0 | cross_entropy | 2.1865211808881653 | 11.94272550000096 |
| 1660 | gse140802_t2_t9_holdout_B1::heldout_animal_B1 | 20260916.0 | cross_entropy | 3.1972012571364465 | 13.191303499999776 |
| 1661 | gse140802_t2_t9_holdout_B1::heldout_animal_B1 | 20260917.0 | cross_entropy | 2.162483715372566 | 10.363531999999395 |
| 1662 | gse140802_t2_t9_holdout_B1::heldout_animal_B1 | 20260918.0 | cross_entropy | 2.0760517608246127 | 13.302652300000773 |
| 1663 | gse140802_t2_t9_holdout_B1::heldout_animal_B1 | 20260919.0 | cross_entropy | 3.16413949434576 | 12.217151799999556 |
| 1664 | gse140802_t2_t9_holdout_B1::heldout_animal_B1 | 20260920.0 | cross_entropy | 2.248140777998532 | 10.476480999999694 |
| 1677 | gse140802_t2_t9_holdout_B2::heldout_animal_B2 | 20260916.0 | cross_entropy | 3.231657325282693 | 14.824461000000156 |
| 1678 | gse140802_t2_t9_holdout_B2::heldout_animal_B2 | 20260917.0 | cross_entropy | 2.135831008691647 | 12.637680699999692 |
| 1679 | gse140802_t2_t9_holdout_B2::heldout_animal_B2 | 20260918.0 | cross_entropy | 2.095972381828932 | 11.214854900001228 |
| 1680 | gse140802_t2_t9_holdout_B2::heldout_animal_B2 | 20260919.0 | cross_entropy | 2.414663556972569 | 12.83930209999926 |
| 1681 | gse140802_t2_t9_holdout_B2::heldout_animal_B2 | 20260920.0 | cross_entropy | 2.2982243510511484 | 12.024557300001106 |
| 1694 | gse140802_t2_t9_holdout_B3::heldout_animal_B3 | 20260916.0 | cross_entropy | 3.463647814378984 | 12.764826900000116 |
| 1695 | gse140802_t2_t9_holdout_B3::heldout_animal_B3 | 20260917.0 | cross_entropy | 1.9574193923028929 | 10.196368300001268 |
| 1696 | gse140802_t2_t9_holdout_B3::heldout_animal_B3 | 20260918.0 | cross_entropy | 1.9541166117093949 | 12.860726999999317 |
| 1697 | gse140802_t2_t9_holdout_B3::heldout_animal_B3 | 20260919.0 | cross_entropy | 2.583104885696264 | 11.956813599999805 |
| 1698 | gse140802_t2_t9_holdout_B3::heldout_animal_B3 | 20260920.0 | cross_entropy | 2.03850830636923 | 11.306239899999127 |
| 1847 | gse140802_t2_t16_holdout_A4::heldout_animal_A4 | 20260916.0 | sinkhorn_divergence | 0.980810657358308 | 11.381618999999771 |
| 1848 | gse140802_t2_t16_holdout_A4::heldout_animal_A4 | 20260917.0 | sinkhorn_divergence | 1.0899683885903255 | 10.828887900001064 |
| 1849 | gse140802_t2_t16_holdout_A4::heldout_animal_A4 | 20260918.0 | sinkhorn_divergence | 0.9390669766006344 | 9.209258099999715 |
| 1850 | gse140802_t2_t16_holdout_A4::heldout_animal_A4 | 20260919.0 | sinkhorn_divergence | 0.8986607127135735 | 8.240687600000456 |
| 1851 | gse140802_t2_t16_holdout_A4::heldout_animal_A4 | 20260920.0 | sinkhorn_divergence | 1.0766710466680358 | 9.692493900000043 |
| 1864 | gse140802_t2_t16_holdout_A5::heldout_animal_A5 | 20260916.0 | sinkhorn_divergence | 0.7411452218540713 | 10.093173400000524 |
| 1865 | gse140802_t2_t16_holdout_A5::heldout_animal_A5 | 20260917.0 | sinkhorn_divergence | 0.8847402034097184 | 10.584232999999584 |
| 1866 | gse140802_t2_t16_holdout_A5::heldout_animal_A5 | 20260918.0 | sinkhorn_divergence | 0.6834093542161573 | 9.87224210000022 |
| 1867 | gse140802_t2_t16_holdout_A5::heldout_animal_A5 | 20260919.0 | sinkhorn_divergence | 0.6958133822905775 | 10.484369800000422 |
| 1868 | gse140802_t2_t16_holdout_A5::heldout_animal_A5 | 20260920.0 | sinkhorn_divergence | 0.8665980690609001 | 10.327723500000502 |
| 1881 | gse140802_t2_t16_holdout_B4::heldout_animal_B4 | 20260916.0 | sinkhorn_divergence | 0.953870143586528 | 10.841824400000403 |
| 1882 | gse140802_t2_t16_holdout_B4::heldout_animal_B4 | 20260917.0 | sinkhorn_divergence | 1.0212267533420878 | 9.889872700001431 |
| 1883 | gse140802_t2_t16_holdout_B4::heldout_animal_B4 | 20260918.0 | sinkhorn_divergence | 0.8512396492734637 | 9.274696800001038 |
| 1884 | gse140802_t2_t16_holdout_B4::heldout_animal_B4 | 20260919.0 | sinkhorn_divergence | 0.8311092803239272 | 10.27010639999935 |
| 1885 | gse140802_t2_t16_holdout_B4::heldout_animal_B4 | 20260920.0 | sinkhorn_divergence | 0.9878213494915488 | 8.525537100000292 |
| 1898 | gse140802_t2_t16_holdout_B5::heldout_animal_B5 | 20260916.0 | sinkhorn_divergence | 0.8104640670253792 | 12.640654699998777 |
| 1899 | gse140802_t2_t16_holdout_B5::heldout_animal_B5 | 20260917.0 | sinkhorn_divergence | 0.7941436265579783 | 10.670966799998496 |
| 1900 | gse140802_t2_t16_holdout_B5::heldout_animal_B5 | 20260918.0 | sinkhorn_divergence | 0.6811199809270838 | 12.243431499999131 |
| 1901 | gse140802_t2_t16_holdout_B5::heldout_animal_B5 | 20260919.0 | sinkhorn_divergence | 0.7523900352055842 | 11.23134349999964 |
| 1902 | gse140802_t2_t16_holdout_B5::heldout_animal_B5 | 20260920.0 | sinkhorn_divergence | 0.8600864513490297 | 9.15854279999985 |
| 1915 | gse140802_t2_t9_holdout_A1::heldout_animal_A1 | 20260916.0 | sinkhorn_divergence | 0.5781817411866412 | 13.377239100000224 |
| 1916 | gse140802_t2_t9_holdout_A1::heldout_animal_A1 | 20260917.0 | sinkhorn_divergence | 0.715949477376505 | 12.320767999999587 |
| 1917 | gse140802_t2_t9_holdout_A1::heldout_animal_A1 | 20260918.0 | sinkhorn_divergence | 0.6273604772348292 | 11.287288700001229 |
| 1918 | gse140802_t2_t9_holdout_A1::heldout_animal_A1 | 20260919.0 | sinkhorn_divergence | 0.7594412517809926 | 12.021956800001137 |
| 1919 | gse140802_t2_t9_holdout_A1::heldout_animal_A1 | 20260920.0 | sinkhorn_divergence | 0.8278675023018632 | 11.787559000000329 |
| 1932 | gse140802_t2_t9_holdout_A2::heldout_animal_A2 | 20260916.0 | sinkhorn_divergence | 0.4920071583111014 | 12.12737290000041 |
| 1933 | gse140802_t2_t9_holdout_A2::heldout_animal_A2 | 20260917.0 | sinkhorn_divergence | 0.7935444618252103 | 10.070772400000353 |
| 1934 | gse140802_t2_t9_holdout_A2::heldout_animal_A2 | 20260918.0 | sinkhorn_divergence | 0.6258808332734538 | 11.219067100000755 |
| 1935 | gse140802_t2_t9_holdout_A2::heldout_animal_A2 | 20260919.0 | sinkhorn_divergence | 0.6487719691187582 | 11.366179100001318 |
| 1936 | gse140802_t2_t9_holdout_A2::heldout_animal_A2 | 20260920.0 | sinkhorn_divergence | 0.8372427044801463 | 9.137043200000337 |
| 1949 | gse140802_t2_t9_holdout_A3::heldout_animal_A3 | 20260916.0 | sinkhorn_divergence | 0.6315147467140592 | 13.791628000000856 |
| 1950 | gse140802_t2_t9_holdout_A3::heldout_animal_A3 | 20260917.0 | sinkhorn_divergence | 0.8823136219924315 | 12.229715900000883 |
| 1951 | gse140802_t2_t9_holdout_A3::heldout_animal_A3 | 20260918.0 | sinkhorn_divergence | 0.7990194353153154 | 11.05017670000052 |
| 1952 | gse140802_t2_t9_holdout_A3::heldout_animal_A3 | 20260919.0 | sinkhorn_divergence | 0.7956587122760758 | 13.881138999999166 |
| 1953 | gse140802_t2_t9_holdout_A3::heldout_animal_A3 | 20260920.0 | sinkhorn_divergence | 0.9890373850793512 | 11.94272550000096 |
| 1966 | gse140802_t2_t9_holdout_B1::heldout_animal_B1 | 20260916.0 | sinkhorn_divergence | 0.6352953793262024 | 13.191303499999776 |
| 1967 | gse140802_t2_t9_holdout_B1::heldout_animal_B1 | 20260917.0 | sinkhorn_divergence | 0.8731605084663274 | 10.363531999999395 |
| 1968 | gse140802_t2_t9_holdout_B1::heldout_animal_B1 | 20260918.0 | sinkhorn_divergence | 0.7968595422704015 | 13.302652300000773 |
| 1969 | gse140802_t2_t9_holdout_B1::heldout_animal_B1 | 20260919.0 | sinkhorn_divergence | 0.7566500364239579 | 12.217151799999556 |
| 1970 | gse140802_t2_t9_holdout_B1::heldout_animal_B1 | 20260920.0 | sinkhorn_divergence | 0.9827456970208298 | 10.476480999999694 |
| 1983 | gse140802_t2_t9_holdout_B2::heldout_animal_B2 | 20260916.0 | sinkhorn_divergence | 0.6058282440374283 | 14.824461000000156 |
| 1984 | gse140802_t2_t9_holdout_B2::heldout_animal_B2 | 20260917.0 | sinkhorn_divergence | 0.886779434943901 | 12.637680699999692 |
| 1985 | gse140802_t2_t9_holdout_B2::heldout_animal_B2 | 20260918.0 | sinkhorn_divergence | 0.8085329002301228 | 11.214854900001228 |
| 1986 | gse140802_t2_t9_holdout_B2::heldout_animal_B2 | 20260919.0 | sinkhorn_divergence | 0.8020695595077193 | 12.83930209999926 |
| 1987 | gse140802_t2_t9_holdout_B2::heldout_animal_B2 | 20260920.0 | sinkhorn_divergence | 1.0050296129435023 | 12.024557300001106 |
| 2000 | gse140802_t2_t9_holdout_B3::heldout_animal_B3 | 20260916.0 | sinkhorn_divergence | 0.3628710608711901 | 12.764826900000116 |
| 2001 | gse140802_t2_t9_holdout_B3::heldout_animal_B3 | 20260917.0 | sinkhorn_divergence | 0.5725662188439201 | 10.196368300001268 |
| 2002 | gse140802_t2_t9_holdout_B3::heldout_animal_B3 | 20260918.0 | sinkhorn_divergence | 0.4763252957244984 | 12.860726999999317 |
| 2003 | gse140802_t2_t9_holdout_B3::heldout_animal_B3 | 20260919.0 | sinkhorn_divergence | 0.6138426406311196 | 11.956813599999805 |
| 2004 | gse140802_t2_t9_holdout_B3::heldout_animal_B3 | 20260920.0 | sinkhorn_divergence | 0.677216637303347 | 11.306239899999127 |
| 2153 | gse140802_t2_t16_holdout_A4::heldout_animal_A4 | 20260916.0 | top1_accuracy | 0.0 | 11.381618999999771 |
| 2154 | gse140802_t2_t16_holdout_A4::heldout_animal_A4 | 20260917.0 | top1_accuracy | 0.0 | 10.828887900001064 |
| 2155 | gse140802_t2_t16_holdout_A4::heldout_animal_A4 | 20260918.0 | top1_accuracy | 0.0 | 9.209258099999715 |
| 2156 | gse140802_t2_t16_holdout_A4::heldout_animal_A4 | 20260919.0 | top1_accuracy | 0.0 | 8.240687600000456 |
| 2157 | gse140802_t2_t16_holdout_A4::heldout_animal_A4 | 20260920.0 | top1_accuracy | 0.0 | 9.692493900000043 |
| 2170 | gse140802_t2_t16_holdout_A5::heldout_animal_A5 | 20260916.0 | top1_accuracy | 0.0 | 10.093173400000524 |
| 2171 | gse140802_t2_t16_holdout_A5::heldout_animal_A5 | 20260917.0 | top1_accuracy | 0.0 | 10.584232999999584 |
| 2172 | gse140802_t2_t16_holdout_A5::heldout_animal_A5 | 20260918.0 | top1_accuracy | 0.0 | 9.87224210000022 |
| 2173 | gse140802_t2_t16_holdout_A5::heldout_animal_A5 | 20260919.0 | top1_accuracy | 0.0 | 10.484369800000422 |
| 2174 | gse140802_t2_t16_holdout_A5::heldout_animal_A5 | 20260920.0 | top1_accuracy | 0.0 | 10.327723500000502 |
| 2187 | gse140802_t2_t16_holdout_B4::heldout_animal_B4 | 20260916.0 | top1_accuracy | 0.0 | 10.841824400000403 |
| 2188 | gse140802_t2_t16_holdout_B4::heldout_animal_B4 | 20260917.0 | top1_accuracy | 0.0 | 9.889872700001431 |
| 2189 | gse140802_t2_t16_holdout_B4::heldout_animal_B4 | 20260918.0 | top1_accuracy | 0.0 | 9.274696800001038 |
| 2190 | gse140802_t2_t16_holdout_B4::heldout_animal_B4 | 20260919.0 | top1_accuracy | 0.0 | 10.27010639999935 |
| 2191 | gse140802_t2_t16_holdout_B4::heldout_animal_B4 | 20260920.0 | top1_accuracy | 0.0 | 8.525537100000292 |
| 2204 | gse140802_t2_t16_holdout_B5::heldout_animal_B5 | 20260916.0 | top1_accuracy | 0.0 | 12.640654699998777 |
| 2205 | gse140802_t2_t16_holdout_B5::heldout_animal_B5 | 20260917.0 | top1_accuracy | 0.0 | 10.670966799998496 |
| 2206 | gse140802_t2_t16_holdout_B5::heldout_animal_B5 | 20260918.0 | top1_accuracy | 0.0 | 12.243431499999131 |
| 2207 | gse140802_t2_t16_holdout_B5::heldout_animal_B5 | 20260919.0 | top1_accuracy | 0.0 | 11.23134349999964 |
| 2208 | gse140802_t2_t16_holdout_B5::heldout_animal_B5 | 20260920.0 | top1_accuracy | 0.0 | 9.15854279999985 |
| 2221 | gse140802_t2_t9_holdout_A1::heldout_animal_A1 | 20260916.0 | top1_accuracy | 0.0 | 13.377239100000224 |
| 2222 | gse140802_t2_t9_holdout_A1::heldout_animal_A1 | 20260917.0 | top1_accuracy | 0.0 | 12.320767999999587 |
| 2223 | gse140802_t2_t9_holdout_A1::heldout_animal_A1 | 20260918.0 | top1_accuracy | 0.0 | 11.287288700001229 |
| 2224 | gse140802_t2_t9_holdout_A1::heldout_animal_A1 | 20260919.0 | top1_accuracy | 0.0 | 12.021956800001137 |
| 2225 | gse140802_t2_t9_holdout_A1::heldout_animal_A1 | 20260920.0 | top1_accuracy | 0.0 | 11.787559000000329 |
| 2238 | gse140802_t2_t9_holdout_A2::heldout_animal_A2 | 20260916.0 | top1_accuracy | 0.0 | 12.12737290000041 |
| 2239 | gse140802_t2_t9_holdout_A2::heldout_animal_A2 | 20260917.0 | top1_accuracy | 0.0 | 10.070772400000353 |
| 2240 | gse140802_t2_t9_holdout_A2::heldout_animal_A2 | 20260918.0 | top1_accuracy | 0.0 | 11.219067100000755 |
| 2241 | gse140802_t2_t9_holdout_A2::heldout_animal_A2 | 20260919.0 | top1_accuracy | 0.0 | 11.366179100001318 |
| 2242 | gse140802_t2_t9_holdout_A2::heldout_animal_A2 | 20260920.0 | top1_accuracy | 0.0 | 9.137043200000337 |
| 2255 | gse140802_t2_t9_holdout_A3::heldout_animal_A3 | 20260916.0 | top1_accuracy | 0.0 | 13.791628000000856 |
| 2256 | gse140802_t2_t9_holdout_A3::heldout_animal_A3 | 20260917.0 | top1_accuracy | 0.0 | 12.229715900000883 |
| 2257 | gse140802_t2_t9_holdout_A3::heldout_animal_A3 | 20260918.0 | top1_accuracy | 0.0 | 11.05017670000052 |
| 2258 | gse140802_t2_t9_holdout_A3::heldout_animal_A3 | 20260919.0 | top1_accuracy | 0.0 | 13.881138999999166 |
| 2259 | gse140802_t2_t9_holdout_A3::heldout_animal_A3 | 20260920.0 | top1_accuracy | 0.0 | 11.94272550000096 |
| 2272 | gse140802_t2_t9_holdout_B1::heldout_animal_B1 | 20260916.0 | top1_accuracy | 0.0 | 13.191303499999776 |
| 2273 | gse140802_t2_t9_holdout_B1::heldout_animal_B1 | 20260917.0 | top1_accuracy | 0.0 | 10.363531999999395 |
| 2274 | gse140802_t2_t9_holdout_B1::heldout_animal_B1 | 20260918.0 | top1_accuracy | 0.0 | 13.302652300000773 |
| 2275 | gse140802_t2_t9_holdout_B1::heldout_animal_B1 | 20260919.0 | top1_accuracy | 0.0 | 12.217151799999556 |
| 2276 | gse140802_t2_t9_holdout_B1::heldout_animal_B1 | 20260920.0 | top1_accuracy | 0.0 | 10.476480999999694 |
| 2289 | gse140802_t2_t9_holdout_B2::heldout_animal_B2 | 20260916.0 | top1_accuracy | 0.0 | 14.824461000000156 |
| 2290 | gse140802_t2_t9_holdout_B2::heldout_animal_B2 | 20260917.0 | top1_accuracy | 0.0 | 12.637680699999692 |
| 2291 | gse140802_t2_t9_holdout_B2::heldout_animal_B2 | 20260918.0 | top1_accuracy | 0.0 | 11.214854900001228 |
| 2292 | gse140802_t2_t9_holdout_B2::heldout_animal_B2 | 20260919.0 | top1_accuracy | 0.0 | 12.83930209999926 |
| 2293 | gse140802_t2_t9_holdout_B2::heldout_animal_B2 | 20260920.0 | top1_accuracy | 0.0 | 12.024557300001106 |
| 2306 | gse140802_t2_t9_holdout_B3::heldout_animal_B3 | 20260916.0 | top1_accuracy | 1.0 | 12.764826900000116 |
| 2307 | gse140802_t2_t9_holdout_B3::heldout_animal_B3 | 20260917.0 | top1_accuracy | 1.0 | 10.196368300001268 |
| 2308 | gse140802_t2_t9_holdout_B3::heldout_animal_B3 | 20260918.0 | top1_accuracy | 1.0 | 12.860726999999317 |
| 2309 | gse140802_t2_t9_holdout_B3::heldout_animal_B3 | 20260919.0 | top1_accuracy | 1.0 | 11.956813599999805 |
| 2310 | gse140802_t2_t9_holdout_B3::heldout_animal_B3 | 20260920.0 | top1_accuracy | 1.0 | 11.306239899999127 |
| 2459 | gse140802_t2_t16_holdout_A4::heldout_animal_A4 | 20260916.0 | multiclass_brier | 0.2879224097313449 | 11.381618999999771 |
| 2460 | gse140802_t2_t16_holdout_A4::heldout_animal_A4 | 20260917.0 | multiclass_brier | 0.418700539950658 | 10.828887900001064 |
| 2461 | gse140802_t2_t16_holdout_A4::heldout_animal_A4 | 20260918.0 | multiclass_brier | 0.2889041136511321 | 9.209258099999715 |
| 2462 | gse140802_t2_t16_holdout_A4::heldout_animal_A4 | 20260919.0 | multiclass_brier | 0.2480139344874812 | 8.240687600000456 |
| 2463 | gse140802_t2_t16_holdout_A4::heldout_animal_A4 | 20260920.0 | multiclass_brier | 0.4146974029246018 | 9.692493900000043 |
| 2476 | gse140802_t2_t16_holdout_A5::heldout_animal_A5 | 20260916.0 | multiclass_brier | 0.1933379039856612 | 10.093173400000524 |
| 2477 | gse140802_t2_t16_holdout_A5::heldout_animal_A5 | 20260917.0 | multiclass_brier | 0.2517274688682777 | 10.584232999999584 |
| 2478 | gse140802_t2_t16_holdout_A5::heldout_animal_A5 | 20260918.0 | multiclass_brier | 0.159496151037691 | 9.87224210000022 |
| 2479 | gse140802_t2_t16_holdout_A5::heldout_animal_A5 | 20260919.0 | multiclass_brier | 0.1598673649490057 | 10.484369800000422 |
| 2480 | gse140802_t2_t16_holdout_A5::heldout_animal_A5 | 20260920.0 | multiclass_brier | 0.2421074096687304 | 10.327723500000502 |
| 2493 | gse140802_t2_t16_holdout_B4::heldout_animal_B4 | 20260916.0 | multiclass_brier | 0.3002771071957894 | 10.841824400000403 |
| 2494 | gse140802_t2_t16_holdout_B4::heldout_animal_B4 | 20260917.0 | multiclass_brier | 0.4053656393470253 | 9.889872700001431 |
| 2495 | gse140802_t2_t16_holdout_B4::heldout_animal_B4 | 20260918.0 | multiclass_brier | 0.2627598261710732 | 9.274696800001038 |
| 2496 | gse140802_t2_t16_holdout_B4::heldout_animal_B4 | 20260919.0 | multiclass_brier | 0.224802090287262 | 10.27010639999935 |
| 2497 | gse140802_t2_t16_holdout_B4::heldout_animal_B4 | 20260920.0 | multiclass_brier | 0.3839598544616181 | 8.525537100000292 |
| 2510 | gse140802_t2_t16_holdout_B5::heldout_animal_B5 | 20260916.0 | multiclass_brier | 0.1507807039744906 | 12.640654699998777 |
| 2511 | gse140802_t2_t16_holdout_B5::heldout_animal_B5 | 20260917.0 | multiclass_brier | 0.1911512565306557 | 10.670966799998496 |
| 2512 | gse140802_t2_t16_holdout_B5::heldout_animal_B5 | 20260918.0 | multiclass_brier | 0.1298748006324776 | 12.243431499999131 |
| 2513 | gse140802_t2_t16_holdout_B5::heldout_animal_B5 | 20260919.0 | multiclass_brier | 0.1331050105595784 | 11.23134349999964 |
| 2514 | gse140802_t2_t16_holdout_B5::heldout_animal_B5 | 20260920.0 | multiclass_brier | 0.2256915924931253 | 9.15854279999985 |
| 2527 | gse140802_t2_t9_holdout_A1::heldout_animal_A1 | 20260916.0 | multiclass_brier | 0.0688268668945295 | 13.377239100000224 |
| 2528 | gse140802_t2_t9_holdout_A1::heldout_animal_A1 | 20260917.0 | multiclass_brier | 0.1665036268703011 | 12.320767999999587 |
| 2529 | gse140802_t2_t9_holdout_A1::heldout_animal_A1 | 20260918.0 | multiclass_brier | 0.1228750595512927 | 11.287288700001229 |
| 2530 | gse140802_t2_t9_holdout_A1::heldout_animal_A1 | 20260919.0 | multiclass_brier | 0.1696290439873226 | 12.021956800001137 |
| 2531 | gse140802_t2_t9_holdout_A1::heldout_animal_A1 | 20260920.0 | multiclass_brier | 0.2090936369584717 | 11.787559000000329 |
| 2544 | gse140802_t2_t9_holdout_A2::heldout_animal_A2 | 20260916.0 | multiclass_brier | 0.0668366458810835 | 12.12737290000041 |
| 2545 | gse140802_t2_t9_holdout_A2::heldout_animal_A2 | 20260917.0 | multiclass_brier | 0.1803540846286523 | 10.070772400000353 |
| 2546 | gse140802_t2_t9_holdout_A2::heldout_animal_A2 | 20260918.0 | multiclass_brier | 0.1342176464893071 | 11.219067100000755 |
| 2547 | gse140802_t2_t9_holdout_A2::heldout_animal_A2 | 20260919.0 | multiclass_brier | 0.1253790194959132 | 11.366179100001318 |
| 2548 | gse140802_t2_t9_holdout_A2::heldout_animal_A2 | 20260920.0 | multiclass_brier | 0.2326749014521689 | 9.137043200000337 |
| 2561 | gse140802_t2_t9_holdout_A3::heldout_animal_A3 | 20260916.0 | multiclass_brier | 0.1373086371556833 | 13.791628000000856 |
| 2562 | gse140802_t2_t9_holdout_A3::heldout_animal_A3 | 20260917.0 | multiclass_brier | 0.3024411702977623 | 12.229715900000883 |
| 2563 | gse140802_t2_t9_holdout_A3::heldout_animal_A3 | 20260918.0 | multiclass_brier | 0.2519251106629385 | 11.05017670000052 |
| 2564 | gse140802_t2_t9_holdout_A3::heldout_animal_A3 | 20260919.0 | multiclass_brier | 0.217127601733783 | 13.881138999999166 |
| 2565 | gse140802_t2_t9_holdout_A3::heldout_animal_A3 | 20260920.0 | multiclass_brier | 0.3803584171399884 | 11.94272550000096 |
| 2578 | gse140802_t2_t9_holdout_B1::heldout_animal_B1 | 20260916.0 | multiclass_brier | 0.1127191055473708 | 13.191303499999776 |
| 2579 | gse140802_t2_t9_holdout_B1::heldout_animal_B1 | 20260917.0 | multiclass_brier | 0.2450683268637843 | 10.363531999999395 |
| 2580 | gse140802_t2_t9_holdout_B1::heldout_animal_B1 | 20260918.0 | multiclass_brier | 0.1964339811688312 | 13.302652300000773 |
| 2581 | gse140802_t2_t9_holdout_B1::heldout_animal_B1 | 20260919.0 | multiclass_brier | 0.222976676516553 | 12.217151799999556 |
| 2582 | gse140802_t2_t9_holdout_B1::heldout_animal_B1 | 20260920.0 | multiclass_brier | 0.3015318852987078 | 10.476480999999694 |
| 2595 | gse140802_t2_t9_holdout_B2::heldout_animal_B2 | 20260916.0 | multiclass_brier | 0.1117177654304648 | 14.824461000000156 |
| 2596 | gse140802_t2_t9_holdout_B2::heldout_animal_B2 | 20260917.0 | multiclass_brier | 0.2607157934429064 | 12.637680699999692 |
| 2597 | gse140802_t2_t9_holdout_B2::heldout_animal_B2 | 20260918.0 | multiclass_brier | 0.214883962411488 | 11.214854900001228 |
| 2598 | gse140802_t2_t9_holdout_B2::heldout_animal_B2 | 20260919.0 | multiclass_brier | 0.2060874524957289 | 12.83930209999926 |
| 2599 | gse140802_t2_t9_holdout_B2::heldout_animal_B2 | 20260920.0 | multiclass_brier | 0.3372706117235449 | 12.024557300001106 |
| 2612 | gse140802_t2_t9_holdout_B3::heldout_animal_B3 | 20260916.0 | multiclass_brier | 0.0296105225966837 | 12.764826900000116 |
| 2613 | gse140802_t2_t9_holdout_B3::heldout_animal_B3 | 20260917.0 | multiclass_brier | 0.1029298405974077 | 10.196368300001268 |
| 2614 | gse140802_t2_t9_holdout_B3::heldout_animal_B3 | 20260918.0 | multiclass_brier | 0.069298039157558 | 12.860726999999317 |
| 2615 | gse140802_t2_t9_holdout_B3::heldout_animal_B3 | 20260919.0 | multiclass_brier | 0.0996288523604585 | 11.956813599999805 |
| 2616 | gse140802_t2_t9_holdout_B3::heldout_animal_B3 | 20260920.0 | multiclass_brier | 0.1392183242787452 | 11.306239899999127 |

**Extended Data Table 4 | Non-inferiority and paired comparisons.**

Source: ED_Table_4_noninferiority_paired.csv. Field names and values retain the source terminology; blank fields indicate values not supplied in the source. Record numbers identify rows in the source file.

**Analysis group 1**

Fields blank in this group: comparator, margin, score_definition, mean_margin_score, noninferior, comparison, paired_dataset_n.

Fields 1–5; records matched by Record.

| **Record** | **benchmark** | **scenario** | **metric** | **positive_favors** | **paired_n** |
| --- | --- | --- | --- | --- | --- |
| 1 | B1_known_truth | equivalent_fit_multiple_parameters | heldout_coupling_relative_frobenius | soft_iot | 15.0 |
| 2 | B1_known_truth | equivalent_fit_multiple_parameters | coefficient_cosine | soft_iot | 15.0 |
| 3 | B1_known_truth | equivalent_fit_multiple_parameters | coefficient_sign_accuracy | soft_iot | 15.0 |
| 4 | B1_known_truth | equivalent_fit_multiple_parameters | pure_column_minimum_curvature | soft_iot | 15.0 |
| 5 | B1_known_truth | equivalent_fit_multiple_parameters | restart_direction_variance | soft_iot | 15.0 |
| 6 | B1_known_truth | hard_marginal_flat_direction | heldout_coupling_relative_frobenius | soft_iot | 15.0 |
| 7 | B1_known_truth | hard_marginal_flat_direction | coefficient_cosine | soft_iot | 15.0 |
| 8 | B1_known_truth | hard_marginal_flat_direction | coefficient_sign_accuracy | soft_iot | 15.0 |
| 9 | B1_known_truth | hard_marginal_flat_direction | pure_column_minimum_curvature | soft_iot | 15.0 |
| 10 | B1_known_truth | hard_marginal_flat_direction | restart_direction_variance | soft_iot | 15.0 |
| 11 | B1_known_truth | identifiable_soft_marginal | heldout_coupling_relative_frobenius | soft_iot | 15.0 |
| 12 | B1_known_truth | identifiable_soft_marginal | coefficient_cosine | soft_iot | 15.0 |
| 13 | B1_known_truth | identifiable_soft_marginal | coefficient_sign_accuracy | soft_iot | 15.0 |
| 14 | B1_known_truth | identifiable_soft_marginal | pure_column_minimum_curvature | soft_iot | 15.0 |
| 15 | B1_known_truth | identifiable_soft_marginal | restart_direction_variance | soft_iot | 15.0 |
| 16 | B1_known_truth | nonlinear_or_hidden_confounding_misspecification | heldout_coupling_relative_frobenius | soft_iot | 15.0 |
| 17 | B1_known_truth | nonlinear_or_hidden_confounding_misspecification | coefficient_cosine | soft_iot | 15.0 |
| 18 | B1_known_truth | nonlinear_or_hidden_confounding_misspecification | coefficient_sign_accuracy | soft_iot | 15.0 |
| 19 | B1_known_truth | nonlinear_or_hidden_confounding_misspecification | pure_column_minimum_curvature | soft_iot | 15.0 |
| 20 | B1_known_truth | nonlinear_or_hidden_confounding_misspecification | restart_direction_variance | soft_iot | 15.0 |
| 21 | B1_known_truth | observation_noise_and_dropout | heldout_coupling_relative_frobenius | soft_iot | 15.0 |
| 22 | B1_known_truth | observation_noise_and_dropout | coefficient_cosine | soft_iot | 15.0 |
| 23 | B1_known_truth | observation_noise_and_dropout | coefficient_sign_accuracy | soft_iot | 15.0 |
| 24 | B1_known_truth | observation_noise_and_dropout | pure_column_minimum_curvature | soft_iot | 15.0 |
| 25 | B1_known_truth | observation_noise_and_dropout | restart_direction_variance | soft_iot | 15.0 |
| 26 | B1_known_truth | shared_plus_context_offset | heldout_coupling_relative_frobenius | soft_iot | 15.0 |
| 27 | B1_known_truth | shared_plus_context_offset | coefficient_cosine | soft_iot | 15.0 |
| 28 | B1_known_truth | shared_plus_context_offset | coefficient_sign_accuracy | soft_iot | 15.0 |
| 29 | B1_known_truth | shared_plus_context_offset | pure_column_minimum_curvature | soft_iot | 15.0 |
| 30 | B1_known_truth | shared_plus_context_offset | restart_direction_variance | soft_iot | 15.0 |

Fields 6–8; records matched by Record.

| **Record** | **mean_difference** | **ci95_lower** | **ci95_upper** |
| --- | --- | --- | --- |
| 1 | 0.0288593975486667 | 0.0167000624886133 | 0.0430744361132459 |
| 2 | 0.916553331106456 | 0.652352621106563 | 1.1781892314519191 |
| 3 | 0.45 | 0.2833333333333333 | 0.6166666666666667 |
| 4 | 0.2199842290551145 | 0.1792142365294088 | 0.2603262079866238 |
| 5 | 0.199830487323871 | 0.1951443068159652 | 0.2042973395561561 |
| 6 | 0.028908779328022 | 0.0191151808582945 | 0.0390590112133629 |
| 7 | 0.6749596933186999 | 0.5129515338141685 | 0.8414419213574108 |
| 8 | 0.3111111111111111 | 0.2111111111111111 | 0.411111111111111 |
| 9 | 0.249500294646765 | 0.2118359757200908 | 0.2839642311713852 |
| 10 | 0.1044142801317334 | 0.0881728024412423 | 0.1200933727832787 |
| 11 | 0.028908779328022 | 0.0191151808582945 | 0.0390590112133629 |
| 12 | 0.6749596933186999 | 0.5129515338141685 | 0.8414419213574108 |
| 13 | 0.3111111111111111 | 0.2111111111111111 | 0.411111111111111 |
| 14 | 0.249500294646765 | 0.2118359757200908 | 0.2839642311713852 |
| 15 | 0.1044142801317334 | 0.0881728024412423 | 0.1200933727832787 |
| 16 | -0.0046658890356185 | -0.0092000935083892 | 0.0002528778612028 |
| 17 | 0.4598756258613382 | 0.3098769897627247 | 0.6265084739995905 |
| 18 | 0.1555555555555555 | 0.0555555555555555 | 0.2666666666666666 |
| 19 | 0.2390954236671424 | 0.2089216723540528 | 0.2682463760480165 |
| 20 | 0.0843929004872588 | 0.071608779446144 | 0.0970209557038204 |
| 21 | 0.1241463088251642 | 0.090039839477343 | 0.1609383323319466 |
| 22 | 0.5947089279809432 | 0.4542354903754507 | 0.732048299352083 |
| 23 | 0.2222222222222222 | 0.1333333333333334 | 0.3 |
| 24 | 0.2503089619132481 | 0.213495366672145 | 0.2833454308892591 |
| 25 | 0.1030124041835771 | 0.0860665835686519 | 0.1196129225041015 |
| 26 | -0.083403944491157 | -0.1091640855889779 | -0.0550596932477628 |
| 27 | 0.6937446660531419 | 0.5063854725215005 | 0.8846555590470682 |
| 28 | 0.3444444444444444 | 0.2444444444444444 | 0.4444444444444445 |
| 29 | 0.2165772909161016 | 0.1826948299576969 | 0.2493914377092269 |
| 30 | 0.1045198532558373 | 0.0880907373081598 | 0.1206534119893146 |

**Analysis group 2**

Fields blank in this group: scenario, positive_favors, mean_difference, comparison, paired_dataset_n.

Fields 1–5; records matched by Record.

| **Record** | **benchmark** | **metric** | **paired_n** | **ci95_lower** | **ci95_upper** |
| --- | --- | --- | --- | --- | --- |
| 31 | B1_known_truth | coupling_relative_frobenius | 90.0 | 0.0115585769970671 | 0.0421038783493074 |
| 32 | B2_lineage_transition | fate_spearman | 4.0 | 0.0129463602791433 | 0.1692175928162474 |
| 33 | B2_lineage_transition | top1_accuracy | 4.0 | -0.1128571428571428 | 0.0632167832167832 |
| 34 | B2_lineage_transition | coupling_relative_frobenius | 4.0 | 0.0533389746063495 | 0.1917424799418539 |
| 35 | B2_lineage_transition | transition_weighted_l1 | 4.0 | 0.063724981658929 | 0.4384031002127444 |
| 36 | B2_lineage_transition | fate_spearman | 4.0 | -0.055503365147403 | 0.1509402103135678 |
| 37 | B2_lineage_transition | top1_accuracy | 4.0 | -0.1307142857142856 | 0.1156643356643357 |
| 38 | B2_lineage_transition | coupling_relative_frobenius | 4.0 | 0.0514396281759108 | 0.1990772200858603 |
| 39 | B2_lineage_transition | transition_weighted_l1 | 4.0 | 0.0718656726825576 | 0.795962976795176 |
| 40 | B2_lineage_transition | fate_spearman | 4.0 | 0.0489775730253626 | 0.064141782110066 |
| 41 | B2_lineage_transition | top1_accuracy | 4.0 | 0.0454605263157894 | 0.1071478521478521 |
| 42 | B2_lineage_transition | coupling_relative_frobenius | 4.0 | 0.0533389746063495 | 0.1867758344140819 |
| 43 | B2_lineage_transition | transition_weighted_l1 | 4.0 | 0.063724981658929 | 0.439465567218467 |
| 44 | B2_lineage_transition | fate_spearman | 4.0 | -0.0669907600783188 | 0.1422072649205576 |
| 45 | B2_lineage_transition | top1_accuracy | 4.0 | -0.1307142857142856 | 0.1156643356643357 |
| 46 | B2_lineage_transition | coupling_relative_frobenius | 4.0 | 0.0508900237748595 | 0.1406871588574592 |
| 47 | B2_lineage_transition | transition_weighted_l1 | 4.0 | 0.0158008878364086 | 0.3197448836815826 |
| 48 | B2_lineage_transition | fate_spearman | 4.0 | -0.0999674120280321 | 0.0593026782820371 |
| 49 | B2_lineage_transition | top1_accuracy | 4.0 | -0.1449999999999999 | 0.1093706293706294 |
| 50 | B2_lineage_transition | coupling_relative_frobenius | 4.0 | 0.0516316804713852 | 0.1516345483509796 |
| 51 | B2_lineage_transition | transition_weighted_l1 | 4.0 | 0.124765655748946 | 0.4094685996323265 |
| 52 | B2_lineage_transition | fate_spearman | 4.0 | -0.1031498553105635 | 0.0581904953488726 |
| 53 | B2_lineage_transition | top1_accuracy | 4.0 | -0.1485714285714285 | 0.1125174825174825 |
| 54 | B2_lineage_transition | coupling_relative_frobenius | 4.0 | 0.0538517888779582 | 0.1875618260317773 |
| 55 | B2_lineage_transition | transition_weighted_l1 | 4.0 | 0.0638394943921416 | 0.4242328368680701 |
| 56 | B2_lineage_transition | fate_spearman | 4.0 | -0.0546896548473916 | 0.1408419311508189 |
| 57 | B2_lineage_transition | top1_accuracy | 4.0 | -0.1307142857142856 | 0.1226573426573426 |
| 58 | B3_prospective_composition | cross_entropy | 2.0 | 5.891327173211264 | 6.266323512666378 |
| 59 | B3_prospective_composition | sinkhorn_divergence | 2.0 | 0.0849655975924919 | 0.0953312732344762 |
| 60 | B3_prospective_composition | top1_accuracy | 2.0 | 0.1773186768029588 | 0.2369629833493876 |

Fields 6–10; records matched by Record.

| **Record** | **comparator** | **margin** | **score_definition** | **mean_margin_score** | **noninferior** |
| --- | --- | --- | --- | --- | --- |
| 31 | hard_ot | 0.05 | reference*(1+relative_margin)-IOT | 0.0266525060640259 | True |
| 32 | cellrank2 | 0.05 | IOT-reference+absolute_margin | 0.0869801125447475 | True |
| 33 | cellrank2 | 0.03 | IOT-reference+absolute_margin | -0.0065467865467865 | False |
| 34 | lineageot | 0.05 | reference*(1+relative_margin)-IOT | 0.1000407750804208 | True |
| 35 | lineageot | 0.05 | reference*(1+relative_margin)-IOT | 0.2510640409358367 | True |
| 36 | lineageot | 0.05 | IOT-reference+absolute_margin | 0.0483037332692103 | False |
| 37 | lineageot | 0.03 | IOT-reference+absolute_margin | 0.00498334998335 | False |
| 38 | mioflow | 0.05 | reference*(1+relative_margin)-IOT | 0.1016184712455346 | True |
| 39 | mioflow | 0.05 | reference*(1+relative_margin)-IOT | 0.3357623585945941 | True |
| 40 | mioflow | 0.05 | IOT-reference+absolute_margin | 0.0565596775677143 | True |
| 41 | mioflow | 0.03 | IOT-reference+absolute_margin | 0.0783193341745973 | True |
| 42 | moscot | 0.05 | reference*(1+relative_margin)-IOT | 0.0983104993002455 | True |
| 43 | moscot | 0.05 | reference*(1+relative_margin)-IOT | 0.251595274438698 | True |
| 44 | moscot | 0.05 | IOT-reference+absolute_margin | 0.0415636198279016 | False |
| 45 | moscot | 0.03 | IOT-reference+absolute_margin | 0.00498334998335 | False |
| 46 | prescient | 0.05 | reference*(1+relative_margin)-IOT | 0.0809317186403825 | True |
| 47 | prescient | 0.05 | reference*(1+relative_margin)-IOT | 0.1569453053442987 | True |
| 48 | prescient | 0.05 | IOT-reference+absolute_margin | 0.0047465982017694 | False |
| 49 | prescient | 0.03 | IOT-reference+absolute_margin | -0.0018764568764568 | False |
| 50 | tigon | 0.05 | reference*(1+relative_margin)-IOT | 0.0858573056259973 | True |
| 51 | tigon | 0.05 | reference*(1+relative_margin)-IOT | 0.2414498236611383 | True |
| 52 | tigon | 0.05 | IOT-reference+absolute_margin | 0.0029452571736283 | False |
| 53 | tigon | 0.03 | IOT-reference+absolute_margin | -0.0020179820179819 | False |
| 54 | wot | 0.05 | reference*(1+relative_margin)-IOT | 0.0993112729161704 | True |
| 55 | wot | 0.05 | reference*(1+relative_margin)-IOT | 0.2440361656301059 | True |
| 56 | wot | 0.05 | IOT-reference+absolute_margin | 0.0457358233591534 | False |
| 57 | wot | 0.03 | IOT-reference+absolute_margin | 0.0073143523143523 | False |
| 58 | PERSIST-no-IOT | 0.05 | reference*(1+relative_margin)-IOT | 6.078825342938821 | True |
| 59 | PERSIST-no-IOT | 0.05 | reference*(1+relative_margin)-IOT | 0.0901484354134841 | True |
| 60 | PERSIST-no-IOT | 0.03 | IOT-reference+absolute_margin | 0.2071408300761732 | True |

**Analysis group 3**

Fields blank in this group: scenario, metric, paired_n, comparator, margin, score_definition, mean_margin_score, noninferior.

Fields 1–5; records matched by Record.

| **Record** | **benchmark** | **positive_favors** | **mean_difference** | **ci95_lower** | **ci95_upper** |
| --- | --- | --- | --- | --- | --- |
| 61 | B2_lineage_transition_output_proxy | UOT-IOT | 0.0110059757596878 | 0.0039436974235691 | 0.0182019679575644 |
| 62 | B2_lineage_transition_output_proxy | UOT-IOT | 0.0007003341550357 | 0.0003014806834681 | 0.0011411135479025 |
| 63 | B2_lineage_transition_output_proxy | UOT-IOT | 0.1196616362994399 | 0.0122762240403488 | 0.2270470485585309 |

Fields 6–7; records matched by Record.

| **Record** | **comparison** | **paired_dataset_n** |
| --- | --- | --- |
| 61 | UOT-IOT_minus_mioflow | 4.0 |
| 62 | UOT-IOT_minus_prescient | 4.0 |
| 63 | UOT-IOT_minus_tigon | 4.0 |

**Extended Data Table 5 | Direction stability and interpretability.**

Source: ED_Table_5_direction_interpretability.csv. Field names and values retain the source terminology; blank fields indicate values not supplied in the source. Record numbers identify rows in the source file.

**Analysis group 1**

Fields blank in this group: dataset, method, direction_pairwise_cosine, direction_pairwise_cosine_min, direction_seed_variance, direction_sign_agreement, direction_type, seed_n, pairwise_cosine_mean, pairwise_cosine_min, pairwise_cosine_variance, training_cohort, validation_cohort, site, direction_status, gain, bootstrap_replicates, ci_excludes_zero, relative_gain.

Fields 1–5; records matched by Record.

| **Record** | **benchmark** | **scenario** | **metric** | **positive_favors** | **paired_n** |
| --- | --- | --- | --- | --- | --- |
| 1 | B1_known_truth | equivalent_fit_multiple_parameters | heldout_coupling_relative_frobenius | soft_iot | 15.0 |
| 2 | B1_known_truth | equivalent_fit_multiple_parameters | coefficient_cosine | soft_iot | 15.0 |
| 3 | B1_known_truth | equivalent_fit_multiple_parameters | coefficient_sign_accuracy | soft_iot | 15.0 |
| 4 | B1_known_truth | equivalent_fit_multiple_parameters | pure_column_minimum_curvature | soft_iot | 15.0 |
| 5 | B1_known_truth | equivalent_fit_multiple_parameters | restart_direction_variance | soft_iot | 15.0 |
| 6 | B1_known_truth | hard_marginal_flat_direction | heldout_coupling_relative_frobenius | soft_iot | 15.0 |
| 7 | B1_known_truth | hard_marginal_flat_direction | coefficient_cosine | soft_iot | 15.0 |
| 8 | B1_known_truth | hard_marginal_flat_direction | coefficient_sign_accuracy | soft_iot | 15.0 |
| 9 | B1_known_truth | hard_marginal_flat_direction | pure_column_minimum_curvature | soft_iot | 15.0 |
| 10 | B1_known_truth | hard_marginal_flat_direction | restart_direction_variance | soft_iot | 15.0 |
| 11 | B1_known_truth | identifiable_soft_marginal | heldout_coupling_relative_frobenius | soft_iot | 15.0 |
| 12 | B1_known_truth | identifiable_soft_marginal | coefficient_cosine | soft_iot | 15.0 |
| 13 | B1_known_truth | identifiable_soft_marginal | coefficient_sign_accuracy | soft_iot | 15.0 |
| 14 | B1_known_truth | identifiable_soft_marginal | pure_column_minimum_curvature | soft_iot | 15.0 |
| 15 | B1_known_truth | identifiable_soft_marginal | restart_direction_variance | soft_iot | 15.0 |
| 16 | B1_known_truth | nonlinear_or_hidden_confounding_misspecification | heldout_coupling_relative_frobenius | soft_iot | 15.0 |
| 17 | B1_known_truth | nonlinear_or_hidden_confounding_misspecification | coefficient_cosine | soft_iot | 15.0 |
| 18 | B1_known_truth | nonlinear_or_hidden_confounding_misspecification | coefficient_sign_accuracy | soft_iot | 15.0 |
| 19 | B1_known_truth | nonlinear_or_hidden_confounding_misspecification | pure_column_minimum_curvature | soft_iot | 15.0 |
| 20 | B1_known_truth | nonlinear_or_hidden_confounding_misspecification | restart_direction_variance | soft_iot | 15.0 |
| 21 | B1_known_truth | observation_noise_and_dropout | heldout_coupling_relative_frobenius | soft_iot | 15.0 |
| 22 | B1_known_truth | observation_noise_and_dropout | coefficient_cosine | soft_iot | 15.0 |
| 23 | B1_known_truth | observation_noise_and_dropout | coefficient_sign_accuracy | soft_iot | 15.0 |
| 24 | B1_known_truth | observation_noise_and_dropout | pure_column_minimum_curvature | soft_iot | 15.0 |
| 25 | B1_known_truth | observation_noise_and_dropout | restart_direction_variance | soft_iot | 15.0 |
| 26 | B1_known_truth | shared_plus_context_offset | heldout_coupling_relative_frobenius | soft_iot | 15.0 |
| 27 | B1_known_truth | shared_plus_context_offset | coefficient_cosine | soft_iot | 15.0 |
| 28 | B1_known_truth | shared_plus_context_offset | coefficient_sign_accuracy | soft_iot | 15.0 |
| 29 | B1_known_truth | shared_plus_context_offset | pure_column_minimum_curvature | soft_iot | 15.0 |
| 30 | B1_known_truth | shared_plus_context_offset | restart_direction_variance | soft_iot | 15.0 |

Fields 6–8; records matched by Record.

| **Record** | **mean_difference** | **ci95_lower** | **ci95_upper** |
| --- | --- | --- | --- |
| 1 | 0.0288593975486667 | 0.0167000624886133 | 0.0430744361132459 |
| 2 | 0.916553331106456 | 0.652352621106563 | 1.1781892314519191 |
| 3 | 0.45 | 0.2833333333333333 | 0.6166666666666667 |
| 4 | 0.2199842290551145 | 0.1792142365294088 | 0.2603262079866238 |
| 5 | 0.199830487323871 | 0.1951443068159652 | 0.2042973395561561 |
| 6 | 0.028908779328022 | 0.0191151808582945 | 0.0390590112133629 |
| 7 | 0.6749596933186999 | 0.5129515338141685 | 0.8414419213574108 |
| 8 | 0.3111111111111111 | 0.2111111111111111 | 0.411111111111111 |
| 9 | 0.249500294646765 | 0.2118359757200908 | 0.2839642311713852 |
| 10 | 0.1044142801317334 | 0.0881728024412423 | 0.1200933727832787 |
| 11 | 0.028908779328022 | 0.0191151808582945 | 0.0390590112133629 |
| 12 | 0.6749596933186999 | 0.5129515338141685 | 0.8414419213574108 |
| 13 | 0.3111111111111111 | 0.2111111111111111 | 0.411111111111111 |
| 14 | 0.249500294646765 | 0.2118359757200908 | 0.2839642311713852 |
| 15 | 0.1044142801317334 | 0.0881728024412423 | 0.1200933727832787 |
| 16 | -0.0046658890356185 | -0.0092000935083892 | 0.0002528778612028 |
| 17 | 0.4598756258613382 | 0.3098769897627247 | 0.6265084739995905 |
| 18 | 0.1555555555555555 | 0.0555555555555555 | 0.2666666666666666 |
| 19 | 0.2390954236671424 | 0.2089216723540528 | 0.2682463760480165 |
| 20 | 0.0843929004872588 | 0.071608779446144 | 0.0970209557038204 |
| 21 | 0.1241463088251642 | 0.090039839477343 | 0.1609383323319466 |
| 22 | 0.5947089279809432 | 0.4542354903754507 | 0.732048299352083 |
| 23 | 0.2222222222222222 | 0.1333333333333334 | 0.3 |
| 24 | 0.2503089619132481 | 0.213495366672145 | 0.2833454308892591 |
| 25 | 0.1030124041835771 | 0.0860665835686519 | 0.1196129225041015 |
| 26 | -0.083403944491157 | -0.1091640855889779 | -0.0550596932477628 |
| 27 | 0.6937446660531419 | 0.5063854725215005 | 0.8846555590470682 |
| 28 | 0.3444444444444444 | 0.2444444444444444 | 0.4444444444444445 |
| 29 | 0.2165772909161016 | 0.1826948299576969 | 0.2493914377092269 |
| 30 | 0.1045198532558373 | 0.0880907373081598 | 0.1206534119893146 |

**Analysis group 2**

Fields blank in this group: scenario, metric, positive_favors, paired_n, mean_difference, ci95_lower, ci95_upper, direction_type, seed_n, pairwise_cosine_mean, pairwise_cosine_min, pairwise_cosine_variance, training_cohort, validation_cohort, site, direction_status, gain, bootstrap_replicates, ci_excludes_zero, relative_gain.

Fields 1–5; records matched by Record.

| **Record** | **benchmark** | **dataset** | **method** | **direction_pairwise_cosine** | **direction_pairwise_cosine_min** |
| --- | --- | --- | --- | --- | --- |
| 31 | B2_lineage_transition | gse140802_t2_t16 | UOT-IOT | 0.999189304252851 | 0.9972169170573716 |
| 32 | B2_lineage_transition | gse140802_t2_t9 | UOT-IOT | 0.9985569175451944 | 0.9963128669912582 |
| 33 | B2_lineage_transition | gse239651_transition_panel | UOT-IOT | 0.9999999999996774 | 0.9999999999990508 |
| 34 | B2_lineage_transition | macsgestalt_transition_panel | UOT-IOT | 0.999999999999972 | 0.9999999999999384 |

Fields 6–7; records matched by Record.

| **Record** | **direction_seed_variance** | **direction_sign_agreement** |
| --- | --- | --- |
| 31 | 0.0001013369683936 | 0.95 |
| 32 | 0.0001803853068507 | 1.0 |
| 33 | 4.608330652804224e-14 | 1.0 |
| 34 | 5.560000779658263e-15 | 1.0 |

**Analysis group 3**

Fields blank in this group: scenario, metric, positive_favors, paired_n, mean_difference, ci95_lower, ci95_upper, direction_pairwise_cosine, direction_pairwise_cosine_min, direction_seed_variance, direction_sign_agreement, training_cohort, validation_cohort, site, direction_status, gain, bootstrap_replicates, ci_excludes_zero, relative_gain.

Fields 1–5; records matched by Record.

| **Record** | **benchmark** | **dataset** | **method** | **direction_type** | **seed_n** |
| --- | --- | --- | --- | --- | --- |
| 35 | B2_lineage_transition_output_proxy | gse140802_t2_t16 | mioflow | output_change_proxy | 5.0 |
| 36 | B2_lineage_transition_output_proxy | gse140802_t2_t16 | prescient | output_change_proxy | 5.0 |
| 37 | B2_lineage_transition_output_proxy | gse140802_t2_t16 | tigon | output_change_proxy | 5.0 |
| 38 | B2_lineage_transition_output_proxy | gse140802_t2_t16 | UOT-IOT | output_change_proxy | 5.0 |
| 39 | B2_lineage_transition_output_proxy | gse140802_t2_t9 | mioflow | output_change_proxy | 5.0 |
| 40 | B2_lineage_transition_output_proxy | gse140802_t2_t9 | prescient | output_change_proxy | 5.0 |
| 41 | B2_lineage_transition_output_proxy | gse140802_t2_t9 | tigon | output_change_proxy | 5.0 |
| 42 | B2_lineage_transition_output_proxy | gse140802_t2_t9 | UOT-IOT | output_change_proxy | 5.0 |
| 43 | B2_lineage_transition_output_proxy | gse239651_transition_panel | mioflow | output_change_proxy | 5.0 |
| 44 | B2_lineage_transition_output_proxy | gse239651_transition_panel | prescient | output_change_proxy | 5.0 |
| 45 | B2_lineage_transition_output_proxy | gse239651_transition_panel | tigon | output_change_proxy | 5.0 |
| 46 | B2_lineage_transition_output_proxy | gse239651_transition_panel | UOT-IOT | output_change_proxy | 5.0 |
| 47 | B2_lineage_transition_output_proxy | macsgestalt_transition_panel | mioflow | output_change_proxy | 5.0 |
| 48 | B2_lineage_transition_output_proxy | macsgestalt_transition_panel | prescient | output_change_proxy | 5.0 |
| 49 | B2_lineage_transition_output_proxy | macsgestalt_transition_panel | tigon | output_change_proxy | 5.0 |
| 50 | B2_lineage_transition_output_proxy | macsgestalt_transition_panel | UOT-IOT | output_change_proxy | 5.0 |

Fields 6–8; records matched by Record.

| **Record** | **pairwise_cosine_mean** | **pairwise_cosine_min** | **pairwise_cosine_variance** |
| --- | --- | --- | --- |
| 35 | 0.9867831219113228 | 0.9620641010947786 | 0.0001312567081836 |
| 36 | 0.9996784168693672 | 0.9992688034897268 | 4.6528386527618354e-08 |
| 37 | 0.811179784426215 | 0.6751141051349906 | 0.010822910713117 |
| 38 | 0.9999999999999948 | 0.9999999999999898 | 1.1756218768085357e-29 |
| 39 | 0.9785732579589712 | 0.9335445146305332 | 0.0004315078109701 |
| 40 | 0.9992159151004486 | 0.99782839768188 | 4.1568343052165014e-07 |
| 41 | 0.7347261184567122 | 0.5990222727199811 | 0.0161151300449865 |
| 42 | 0.9999999999999944 | 0.9999999999999892 | 1.0205887961296839e-29 |
| 43 | 0.991472354292795 | 0.9797715455910626 | 2.6772836779552747e-05 |
| 44 | 0.999718621763674 | 0.9994089979443004 | 2.2108592849093744e-08 |
| 45 | 0.9848122044126048 | 0.9633524685369584 | 0.0001108688234851 |
| 46 | 0.9999999999999828 | 0.9999999999999653 | 8.623646635251988e-29 |
| 47 | 0.999147362798084 | 0.9981557161886372 | 2.9937227770014384e-07 |
| 48 | 0.9985857096462916 | 0.9962378084695191 | 1.6779070795147743e-06 |
| 49 | 0.9906353475066324 | 0.9754842138450092 | 0.0001123355780941 |
| 50 | 0.9999999999999524 | 0.9999999999998828 | 1.2563185126721335e-27 |

**Analysis group 4**

Fields blank in this group: scenario, positive_favors, paired_n, mean_difference, dataset, method, direction_pairwise_cosine, direction_pairwise_cosine_min, direction_seed_variance, direction_sign_agreement, direction_type, seed_n, pairwise_cosine_mean, pairwise_cosine_min, pairwise_cosine_variance.

Fields 1–5; records matched by Record.

| **Record** | **benchmark** | **metric** | **ci95_lower** | **ci95_upper** | **training_cohort** |
| --- | --- | --- | --- | --- | --- |
| 51 | external_frozen_direction | conditional_transition_mae_gain_over_independence | 0.0280030501463581 | 0.0298753725638859 | GSE163558 |
| 52 | external_frozen_direction | conditional_transition_mae_gain_over_independence | 0.0207568095326046 | 0.0239218660063596 | GSE163558 |
| 53 | external_frozen_direction | conditional_transition_mae_gain_over_independence | 0.0289964311641579 | 0.0313483896185037 | GSE163558 |
| 54 | external_frozen_direction | conditional_transition_mae_gain_over_independence | 0.0409769540077706 | 0.0469881721565458 | GSE163558 |
| 55 | external_frozen_direction | conditional_transition_mae_gain_over_independence | 0.0710046599141826 | 0.0761329001343414 | GSE163558 |
| 56 | external_frozen_direction | conditional_transition_mae_gain_over_independence | 0.0549188589190246 | 0.0622782911157637 | GSE163558 |

Fields 6–10; records matched by Record.

| **Record** | **validation_cohort** | **site** | **direction_status** | **gain** | **bootstrap_replicates** |
| --- | --- | --- | --- | --- | --- |
| 51 | GSE246662 | GSM7874175_LM1 | frozen_before_validation | 0.0290196646248589 | 200.0 |
| 52 | GSE246662 | GSM7874176_LM2 | frozen_before_validation | 0.0223080073324266 | 200.0 |
| 53 | GSE246662 | GSM7874177_LM3 | frozen_before_validation | 0.0301460531965383 | 200.0 |
| 54 | GSE183904 | s19 | frozen_before_validation | 0.0437538300366153 | 200.0 |
| 55 | GSE183904 | s20 | frozen_before_validation | 0.0735746918829006 | 200.0 |
| 56 | GSE183904 | s38 | frozen_before_validation | 0.0586592208897275 | 200.0 |

Fields 11–12; records matched by Record.

| **Record** | **ci_excludes_zero** | **relative_gain** |
| --- | --- | --- |
| 51 | True | 0.2082106538216941 |
| 52 | True | 0.1621098916859902 |
| 53 | True | 0.1986557010601491 |
| 54 | True | 0.2872568510641332 |
| 55 | True | 0.4609673918350048 |
| 56 | True | 0.3801399287265014 |

**Extended Data Table 6 | Persistence and detection performance.**

Source: ED_Table_6_detection_boundary.csv. Field names and values retain the source terminology; blank fields indicate values not supplied in the source. Record numbers identify rows in the source file.

| **Record** | **partition** | **metric** | **role** | **method** | **value** |
| --- | --- | --- | --- | --- | --- |
| 1 | development_outer_oof | Brier | PERSIST-IOT | PERSIST-IOT | 0.050092494660085 |
| 2 | development_outer_oof | Brier | Best supervised | extra_trees | 0.045277149520772 |
| 3 | development_outer_oof | AUPRC | PERSIST-IOT | PERSIST-IOT | 0.7806524464438694 |
| 4 | development_outer_oof | AUPRC | Best supervised | random_forest | 0.809823426221381 |
| 5 | locked_external_e1 | Brier | PERSIST-IOT | PERSIST-IOT | 0.0297555056253869 |
| 6 | locked_external_e1 | Brier | Best supervised | hist_gradient_boosting | 0.0310343873785891 |
| 7 | locked_external_e1 | AUPRC | PERSIST-IOT | PERSIST-IOT | 0.7155979268605477 |
| 8 | locked_external_e1 | AUPRC | Best supervised | hist_gradient_boosting | 0.7327494279065814 |

**Paired Brier bootstrap comparisons**

Source: ED_Table_6b_brier_paired_bootstrap.csv. Differences are PERSIST-IOT minus the supervised comparator; bootstrap_replicates = 10000 for both partitions.

| **partition** | **dataset_id** | **comparator** | **difference_persist_minus_supervised** | **ci_low** | **ci_high** |
| --- | --- | --- | --- | --- | --- |
| development_outer_oof | GSE239651_expt1 | extra_trees | 0.004815345139313 | 0.0026908434239634 | 0.0071931092434346 |
| locked_external_e1 | GSE239651_expt2 | hist_gradient_boosting | -0.0012788817532021 | -0.0021882271048234 | -0.0003812870161306 |
